# A Measure of Transcriptional Dyscoordination for Quantifying Aging in Single Cells

**DOI:** 10.64898/2026.01.24.701460

**Authors:** Yilin Yang, Paul R. Hess, Xinyi E. Chen, Sijia Huang, Marcos G. Teneche, Hanzhi Wang, Karl N. Miller, Andrew E. Davis, Charlene Miciano, Kelly Yichen Li, Sainath Mamde, Kevin Yip, Bing Ren, Qian Yang, Elizabeth Smoot, Allen Wang, Bradley Johnson, Alexander C. Huang, Parker Wilson, Peter D. Adams, Nancy R. Zhang

## Abstract

Aging reshapes tissues through changes in cellular composition, coordinated transcriptional reprogramming, and loss of transcriptional coordination. Whereas the first two have been characterized across aging tissues, the third remains difficult to quantify. We introduce an unsupervised, first-principles framework for measuring transcriptional dyscoordination in single cells as deviation from a learned, predictable structure accounting for technical noise. Orthogonal validation links transcriptional dyscoordination to classical intrinsic noise and distinguishes it from coordinated change. In controlled perturbations, dyscoordination rises after genotoxic injury and senescence induction, then falls following senolytic depletion. Across mouse, rat, and human tissues, dyscoordination increases with chronological age, especially in regenerative compartments. In human T cells, dyscoordination increases with clonal expansion and effector function yet declines within persisting clones after checkpoint blockade. Cross-modal analyses further link dyscoordination to chromatin-based mitotic age and genome instability. These results identify loss of transcriptional coordination as a distinct and dynamic feature of cellular aging.

## Introduction

Cellular aging is widely believed to arise from the gradual accumulation of molecular damage^1–6^. According to the cellular damage hypothesis, damage to the genome propagates disorganization across molecular layers, impairing the coordination of normal cellular functions. DNA damage accumulates even in non-dividing cells such as neurons^7–9^ and muscle fibers^10,11^, and evidence shows that these lesions, along with persistent chromatin alterations triggered by DNA repair, contribute to widespread dysregulation in gene expression with age^12–15^. At the epigenetic level, age-related loss of chromatin accessibility and disruption of repressive domains further amplify transcriptional disarray, with aging cells exhibiting increasing variability in gene regulation across tissues and individuals^13,16–22^. These observations motivate a fundamental question: how does the accumulated molecular disruption of aging manifest in the transcriptome of individual cells?

At single-cell resolution, age-associated transcriptomic remodeling can be considered along at least three nonexclusive dimensions. First, tissues undergo changes in cellular composition, as particular populations expand, contract, or are replaced with age^13,23–25^. Second, cells within a defined population can undergo coordinated transcriptional reprogramming, producing shifts in mean gene expression and cellular state^13,26,27^. Third, cells can exhibit a loss of transcriptional coordination, in which the expression of individual genes becomes less predictable from the coordinated expression structure of the cell.

These phenomena are conceptually distinct: compositional change alters which cells are present, coordinated reprogramming alters the transcriptional state occupied by those cells, and dyscoordination reflects reduced precision with which that state is executed. The first two dimensions have been extensively characterized through single-cell atlas, cell composition, and differential expression analyses^23–25,28–37^. Here, we focus on the third dimension, which remains considerably more difficult to quantify at single-cell resolution.

Loss of transcriptional coordination has long been proposed as a feature of aging, but its extent and biological significance remain unresolved^14,21,23,29,38–45^. At the bulk sequencing level, several reports have suggested that gene expression becomes more variable with age^40,41,46–48^, but others have failed to detect a consistent signal^49–51^, leaving the extent and significance of transcriptional dyscoordination an open question. Single-cell RNA sequencing has opened the door to more granular assessments of transcriptional noise, allowing recent studies to trace age-related increase in variability from various perspectives. One common strategy is to cluster cells into putatively homogeneous states and compute gene-level or cell-level variances within each cluster^28,30–32,43,52,53^. However, this approach depends critically on the resolution of clustering: too coarse and biologically distinct states may be merged, while too fine and technical noise may dominate the variance estimate. On the other hand, multiple approaches estimate cell-level transcriptomic variances or identify aging-related features through carefully curated aging-related gene sets or data collections^38,42,44,54–60^. In parallel, a growing number of methods quantify biological aging by measuring stochastic epigenetic variation in DNA methylation data using predefined sets of CpG sites^61–70^. Because these approaches depend on specific cell state definitions, feature sets, or reference datasets, their performance can vary substantially across cellular contexts. Indeed, recent benchmarking studies have found poor agreement among transcriptional variability measures across aging datasets and tissues^39,53^. Thus, a central challenge is to distinguish genuine loss of transcriptional coordination from coordinated changes in cell identity or state, while accounting for the substantial technical variation inherent to single-cell RNA sequencing.

To quantify this complementary dimension of transcriptomic aging, we developed an unsupervised measure of transcriptional dyscoordination for single-cell RNA-sequencing data. Rather than replacing analyses of cellular composition or coordinated gene-expression change, the method investigates a distinct question: how precisely does a cell conform to the predictable expression structure learned from the population? We first establish the meaning of this quantity through controlled simulations and allele-specific analyses, then test its responsiveness to experimentally induced damage and senescence. We next apply the framework across regenerative compartments, lineage-resolved human T cells, and aging kidney and liver, where matched chromatin measurements provide independent readouts of mitotic history and somatic genome instability. At the gene level, we distinguish loss of expression precision from coordinated activation of programs associated with high-dyscoordination cells. Together, these analyses position dyscoordination as a complementary, scalable dimension of cellular aging rather than a comprehensive measure of all age-associated transcriptional change.

## Results

### Defining and quantifying transcriptional dyscoordination

We start with a first-principles definition of transcriptional dyscoordination that bypasses clustering and predefined markers by exploiting predictable gene–gene relationships across cells. Conceptually, coordinated transcriptional states occupy a low-dimensional structure in high-dimensional expression space (Figure 1A). Computationally, however, we do not explicitly estimate a classical geometric manifold or rely on a particular low-dimensional embedding. Instead, for each gene, we estimate the expression expected from the rest of the transcriptome using a regularized prediction model. Coordinated differences among cell types, differentiation states, and other shared programs are represented in this predictable expression component. Crucially, this prediction is applied to latent biological expression after modeling the technical sampling variation inherent to single-cell RNA sequencing. Transcriptional dyscoordination then quantifies the residual biological variation that is not predicted by this shared expression structure.

**Figure 1.**
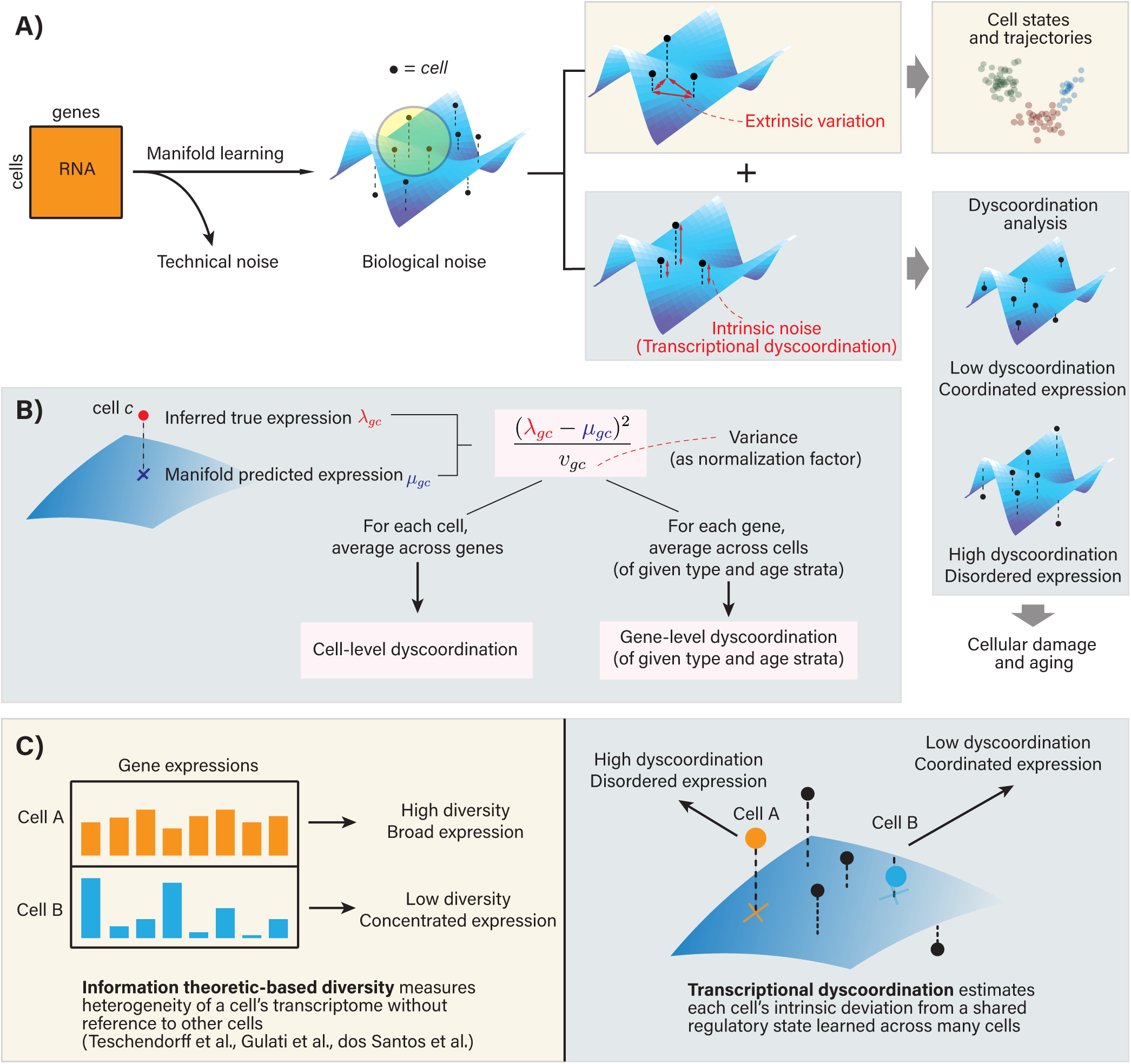
Defining and quantifying transcriptional dyscoordination. **A)** Conceptual framework for defining transcriptional dyscoordination. A low-dimensional manifold (blue surfaces) provides geometric intuition for the coordinated expression structure across cells (black dots). Variation along this structure represents coordinated state differences, whereas residual biological deviations from the predictable expression reference, after accounting for technical sampling variation, contribute to transcriptional dyscoordination. **B)** For each gene *g* and cell *c*, a variance-normalized squared residual deviation between inferred expression (*λ̂_gc_*) and predictable expression estimate (*µ̂gc*) iscomputed. Averaging them across genes yields cell-level dyscoordination. Averaging them across cells yields gene-level dyscoordination. **C)** Differences between transcriptional dyscoordination and information theoretic-based diversity measures.

To distinguish biological dyscoordination from technical sampling variation, we build on the statistical measurement model developed in SAVER, which explicitly models UMI sampling noise and has been extensively validated for recovery of biological expression variation from sparse single-cell RNA-sequencing data^72^. SAVER combines a count-based sampling model with regularized gene-wise prediction and Bayesian shrinkage, allowing the predictable biological component to be estimated without treating sparse count fluctuations as biological variability. Thus, technical noise modeling is an integral part of the dyscoordination estimator, rather than a post hoc correction applied to its output^71,72^. For each cell, we define cell-level transcriptional dyscoordination as the squared deviation from its regularized predictable-expression estimate, averaged over genes and adjusted against a gene- and cell-specific expected variance term. Similarly, gene-level dyscoordination is computed as the average squared deviation from the predictable-expression estimate across cells, standardized against the expected variance for that gene, again after accounting for technical noise (Figure 1B). We then conduct dyscoordination comparisons within the same annotated cell type across age or condition strata, asking whether cells of that type become more dyscoordinated under the condition of interest.

Note that absolute dyscoordination levels are not interpreted as directly comparable across transcriptionally distinct cell types (Methods). Throughout the manuscript, we use the term “expression precision” as the inverse interpretation of this residual magnitude: smaller variance-scaled deviations indicate greater precision relative to the predictable-expression reference, whereas larger deviations indicate loss of precision.

To demonstrate how this framework distinguishes coordinated transcriptional reprogramming from loss of transcriptional coordination, we performed an in-silico spike-in experiment using real single-cell RNA-sequencing datasets as the substrate.

Specifically, we perturbed real data by adding either a strong low-rank expression program in which a set of genes changed coherently across cells, a mean-preserving increase in independent gene-wise variability, or both perturbations simultaneously. The coordinated expression program produced extensive differential expression but essentially no increase in dyscoordination, and it was recovered predominantly in the fitted predictable expression component rather than the residual. Conversely, increasing gene-wise variability while preserving mean expression increased dyscoordination without producing the corresponding coordinated expression shift. When both changes were introduced, the framework recovered both components. These results illustrate that transcriptional dyscoordination is not a measure of transcriptional change in the general sense: coordinated changes in cellular state are incorporated into the predictable expression structure when sufficiently represented in the data, whereas loss of expression coordination contributes to the transcriptional dyscoordination measure (Supplementary Note 1, Supplementary Figure 1-2).

Information-theoretic notions of entropy have also been used to characterize transcriptional heterogeneity in single cells. Measures such as signaling entropy (SCENT)^73^, transcriptional diversity score of differentiation potential (CytoTRACE)^74^, and Shannon-based transcriptomic entropy computed from bulk tissue profiles quantify the distributional diversity of an individual cell’s transcriptome^75^, or how broadly transcripts are spread across genes, without referring to other cells. By contrast, rather than summarizing the shape of a single cell’s distribution, transcriptional dyscoordination measures how strongly each cell deviates from the predictable expression structure learned from the population (Figure 1C). Because information-theoretic entropy depends only on a cell’s own transcriptome, it largely tracks cellular identity and differentiation state, whereas transcriptional dyscoordination is designed to account for coordinated differences through the learned predictable-expression reference. The two therefore capture fundamentally different aspects of a cell’s transcriptome and, as we demonstrate later, can exhibit opposing age-associated trends.

We systematically evaluated several potential confounders across the datasets analyzed in this study. By regressing out a single global LOESS fit of dyscoordination against total counts per cell, we verified that all reported age- and condition-associated dyscoordination changes persist in the depth-adjusted residuals. We also subsampled each age group to a common cell number and found dyscoordination estimates to remain stable with age-associated differences preserved, indicating robustness to differing cell counts across strata. Furthermore, group-level dyscoordination distributions were not driven by individual biological replicates, and cell cycle activity contributed only weakly to cell-level dyscoordination, consistent with our spike-in experiment that showed that sufficiently coordinated programs were absorbed into the predictable expression structure. Together, these analyses indicate that the reported dyscoordination patterns are not explained by these common technical covariates and capture reproducible biological variation (Supplementary Note 2). Relevant analyses are noted alongside each result section below.

### Transcriptional dyscoordination aligns with classical measures of intrinsic transcriptional noise

In pioneering studies in the early 2000’s, Elowitz et al. articulated the distinction between intrinsic transcriptional “noise” and extrinsic transcriptional variation^76^. In their seminal dual-reporter experiments, two identically regulated fluorescent proteins (e.g., GFP and YFP) were expressed from separate genomic loci in single cells, and the per-cell difference in their expression levels was used to quantify intrinsic transcriptional variation. In this way, they defined intrinsic transcriptional variation as stochastic fluctuations arising from the randomness of transcriptional processes in an otherwise homogeneous intracellular environment. While single-cell RNA sequencing lacks the temporal resolution or controlled reporter design of these experiments, we mimicked the dual-reporter design through allele-specific gene expression analysis (Figure 2A).

**Figure 2.**
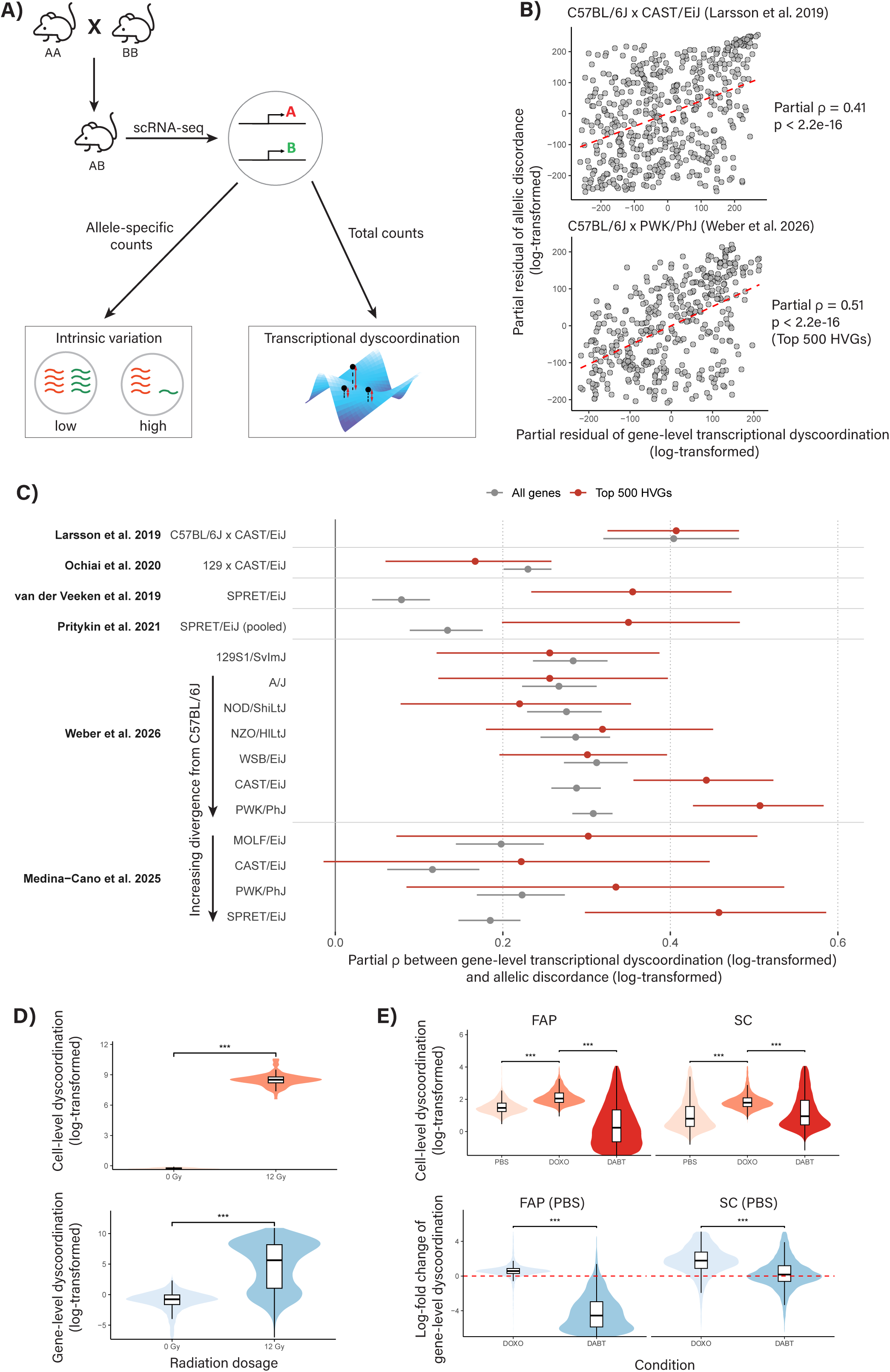
Biological validation of transcriptional dyscoordination through intrinsic noise, cellular damage, and senescence. **A)** In F1 hybrid mice, intrinsic variation in allele-specific counts is estimated through allelic discordance and in total counts is estimated through gene-level dyscoordination. **B)** Scatter plot of partial residuals of log-transformed allelic discordance vs. partial residuals of log-transformed gene-level dyscoordination for two representative F1 hybrid mice experiments showing that transcriptional dyscoordination aligns with classical measures of intrinsic transcriptional noise. **C)** Partial Spearman’s correlation between log-transformed gene-level dyscoordination and log-transformed allelic discordance across 6 F1 hybrid mice experiments over all covered genes (grey) and top 500 HVGs (red). Whiskers indicate the bootstrap 95% confidence intervals and the solid vertical line marks ρ = 0. Crosses within the multi-cross datasets are ordered by their genome-wide sequence divergence from the reference strain (C57BL/6J) from top to bottom. **D)** Log-transformed cell-(left) and gene-level (right) dyscoordination of ESCC cells exposed to 0 Gy and 12 Gy doses of ionizing radiation. **E)** Log-transformed cell-level (left) and gene-level (right) dyscoordination of fibro-adipogenic progenitors and satellite cells sorted from mouse skeletal muscle under three conditions: vehicle control (PBS), doxorubicin (DOXO), and doxorubicin followed by the senolytic ABT-263 (DABT).

We extracted allele-specific reads from six scRNA-seq datasets of F1 hybrid mice^77–82^, which encompass small, homogeneous stem cell benchmarks and large, heterogeneous differentiated cell atlases, and treated each allele of a given gene as a replicate of identically regulated units (Supplementary Table 1). Some loci exhibit allele-specific biases reproducible across cells, which were adjusted through allele-specific mean subtraction. After this adjustment, we defined allelic discordance for each gene in each cell as the squared deviation of the cell’s allelic expression balance from the gene’s baseline allelic ratio, normalized by the binomial sampling variance, of which the genome-wide pattern serves as a proxy for intrinsic noise in the classical sense. Mirroring our derivation of the two manifold-based transcriptional dyscoordination measures, we summarized allelic discordance at the gene level by averaging across cells within a stratum and at the cell level by averaging across a cell’s informative genes (Methods). Note that allelic discordance is computed from the difference between alleles and can only be applied to genes with heterozygous loci in the sequenced region, whereas transcriptional dyscoordination uses only the sum of alleles and can be applied universally.

We first compared these measures at the gene level. Across all crosses of the six datasets, gene-level allelic discordance showed a consistently positive, expression-controlled partial correlation with gene-level dyscoordination (Figure 2B-2C, Supplementary Figure 3-7). In most datasets, this correlation becomes more prominent when we restrict to the top 500 highly variable genes (HVGs), consistent with intrinsic noise being most reliably estimated where transcriptional variability is the highest. Critically, the strength of this correlation also closely tracked how genetically distinct a cell’s two haplotypes are. Within the two multi-cross datasets, each cross pairs C57BL/6J with a different second strain, resulting in a wide range of genome-wide sequence divergence between the two parental haplotypes. As this sequence divergence increased, the correlation between dyscoordination and discordance also strengthened, providing an internal validation that the observed concordance reflects genuine biological signal rather than a spurious association (Figure 2C, Supplementary Figure 8).

At the cell level, we further examined whether cells with the greatest dyscoordination are also those whose alleles are most discordant. Cell-level allelic measurements are inherently noisy, as each cell’s estimate is derived from only a few hundred well-covered heterozygous genes. Nevertheless, cell-level dyscoordination and allelic discordance were still significantly positively associated in the more deeply sequenced datasets (e.g., Weber et al. 2026), and the strength of these associations scaled tightly with each cell’s measurement reliability, indicating that the gene-level relationship is recapitulated across individual cells but attenuated by limited reliability (Methods, Supplementary Figure 9-10). Collectively, these findings show that transcriptional dyscoordination aligns with an independent proxy for classical intrinsic transcriptional noise, providing orthogonal biological validation of the measure. At the same time, this correspondence does not imply that all residual variation captured by dyscoordination is stochastic, as rare or incompletely modeled biological programs may also contribute to the residual.

### Transcriptional dyscoordination increases with induced damage and senescence

Before interrogating the heterogeneous and subtle changes expected in natural aging, we first tested whether transcriptional dyscoordination responds to controlled, experimentally induced perturbations of cellular integrity, where transcriptional disruption is expected to be pronounced. Two complementary laboratory scenarios were examined: acute genotoxic injury from radiation, and drug-induced cellular senescence. Ionizing radiation is a well-characterized inducer of DNA and chromatin damage and provides a controlled perturbation to assess the sensitivity of transcriptomic measures to cellular injury. We used an scRNA-seq dataset on esophageal squamous cell carcinoma (ESCC) cells exposed to 0 or 12 gray radiation doses^83^. Because these are transformed cells that already carry substantial transcriptional and genomic alteration, and because a 12 Gy dose engages several coherent responses including cell cycle arrest and apoptosis, we treat this experiment as an extreme positive control for severe cellular injury rather than as a model of physiological aging. As shown in Figure 2D, both cell-level and gene-level dyscoordination significantly increase in 12 Gy treated cells from 0 Gy treated cells. This finding supports the view that transcriptional dyscoordination reflects damage-induced breakdown of gene expression coordination: as DNA lesions and chromatin disorganization accumulate, gene expression becomes increasingly stochastic and decoupled from coordinated regulatory programs.

We next examined cellular senescence, a stable, stress-induced state that accumulates with age and is frequently a downstream consequence of cellular damage. In an scRNA-seq dataset of fibro-adipogenic progenitors (FAP) and satellite cells (SC) sorted from mouse skeletal muscle, cells were profiled under three conditions: phosphate-buffered saline as vehicle control (PBS), doxorubicin, which inflicts DNA double-strand breaks and drives cells into senescence through a sustained DNA damage response (DOXO), and doxorubicin followed by the senolytic ABT-263 (navitoclax), which inhibits the pro-survival proteins that senescent cells depend on, selectively triggers their apoptosis and thereby depletes them from the population (DABT)^84^. This dataset is particularly informative because the same primary cell types are examined across senescence induction and subsequent senolytic depletion. Cells from the three conditions are broadly intermixed in the transcriptomic embedding (Supplementary Figure 11), but we do not interpret this as evidence that senescence lacks coordinated transcriptional changes. Rather, the design provides a controlled setting to ask whether dyscoordination changes bidirectionally as a damage- and senescence-enriched population is induced and then depleted. Consistent with this prediction, DOXO cells showed significantly elevated cell- and gene-level dyscoordination relative to controls in both cell types, and this elevation was significantly reduced following senolytic clearance (Figure 2E). Together, these two controlled perturbations show that transcriptional dyscoordination increases after acute genotoxic injury and senescence induction and is reduced following senolytic depletion, supporting its interpretation as a biologically responsive dimension of cellular state.

### Transcriptional dyscoordination increases with age in regenerative cell compartments

In the hematopoietic system, age-associated loss of lymphoid progenitors, myeloid skewing, and transcriptional remodeling of stem and progenitor populations are well established^85–87^. Here, we ask whether cells within these populations also lose transcriptional coordination with age. We first applied the dyscoordination framework to single-cell RNA-sequencing data from the bone marrow of Tabula Muris Senis^24^. Notably, UMAP visualizations indicate that transcriptomic embeddings for most cell types, including multiple hematopoietic lineage cell types, are intermixed across various age strata (Supplementary Figure 12). This absence of age-associated clustering underscores the challenge of identifying aging signatures in heterogeneous and dynamic tissues like bone marrow. We computed the manifold-based cell-level and gene-level transcriptional dyscoordination for all 18 annotated cell types spanning the hematopoietic differentiation hierarchy. Gene- and cell-level dyscoordination measures were estimated separately for each cell type, allowing us to dissect aging dynamics for cells of varying differentiation potential (Supplementary Figure 13-14). In particular, hematopoietic precursor cells (HPCs) showed a striking, monotonic increase in transcriptional dyscoordination with chronological age at both the cell- and gene-levels (ρ = 0.397, p < 2.2 x 10^-16^, Figure 3A).

**Figure 3.**
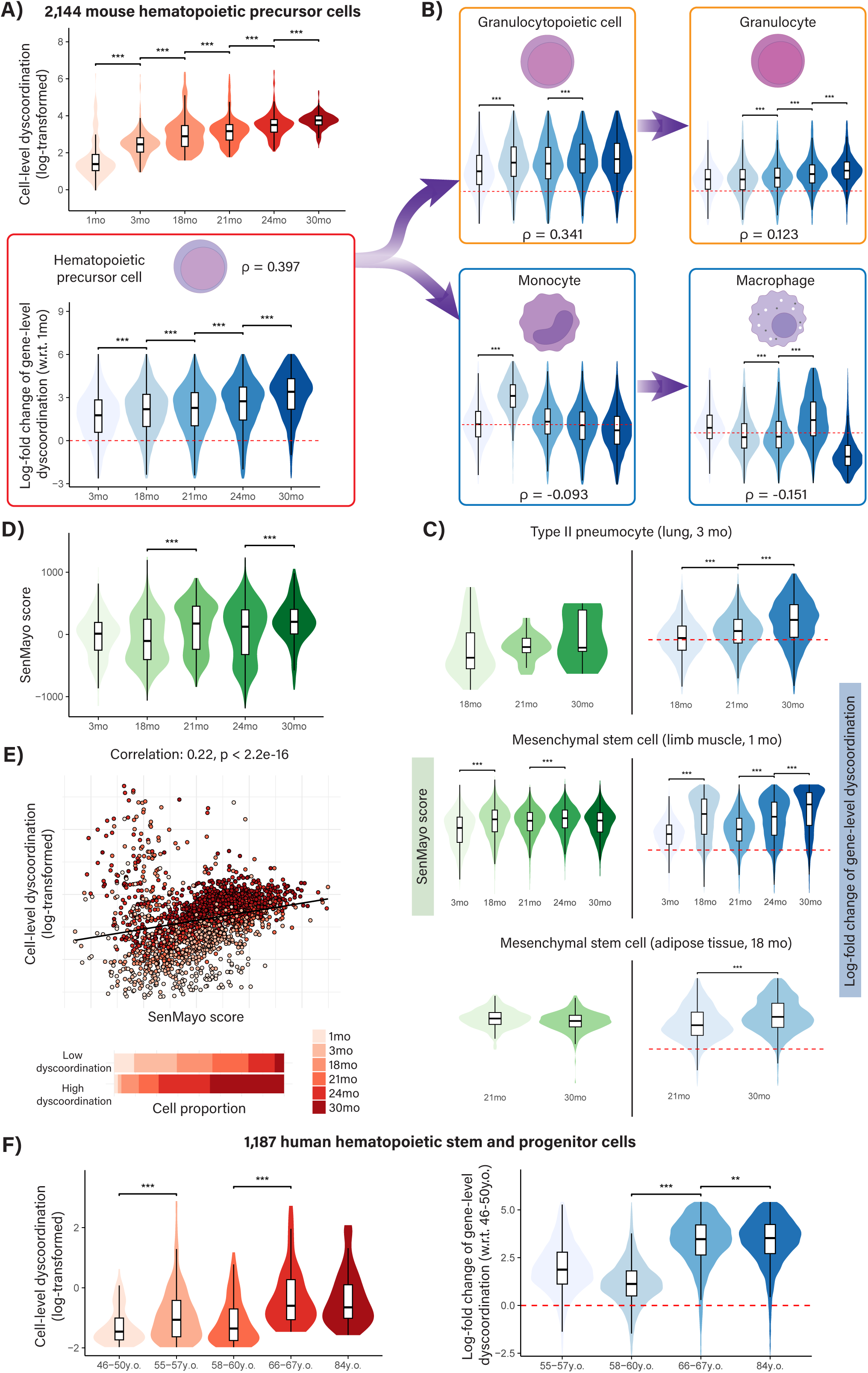
Transcriptional dyscoordination in regenerative cell compartments. **A)** Violin and box plots showing log-transformed cell-level dyscoordination and log-fold change of gene-level dyscoordination with respect to 1-month-old mice by age group in 2,144 hematopoietic precursor cells, along with Spearman’s ρ and corresponding p-values to indicate their associations with age. Values in the top and bottom 2.5% quantiles are considered outliers and are removed from the illustration. Red dashed line denotes zero log-fold change. Pairwise comparisons between age groups were tested using one-sided Wilcoxon rank-sum test (alternative hypothesis: older age group has higher dyscoordination). Significance levels are annotated above each comparison (* p < 0.05, ** p < 0.01, *** p < 0.001). **B)** Violin and box plots showing log-fold change of gene-level dyscoordination with respect to 1-month-old mice by age group across hematopoietic lineages. **C)** Violin and box plots showing SenMayo score (left) and log-fold change of gene-level dyscoordination relative to the baseline age group (right) for selected stem cell populations. Baseline age groups are indicated in parentheses of each subplot title. **D)** Violin and box plots showing SenMayo score of hematopoietic precursor cells. **E)** Scatter plot showing correlation between log-transformed cell-level dyscoordination and SenMayo score of hematopoietic precursor cells. Black solid line denotes the best-fit regression line, with cells above it or below it denoted as low or high dyscoordination. Age proportions for low and high-dyscoordination cells are shown in the stacked bar plots (bottom). **F)** Violin and box plots showing log-transformed cell-level dyscoordination and log-fold change of gene-level dyscoordination with respect to 46 to 50-year-old donors in 1,187 human hematopoietic stem and progenitor cells. Values in the top and bottom 2.5% quantiles are considered outliers and are removed from the illustration.

This trend remained robust to sequencing depth (ρ = 0.584, p < 2.2 x 10^-16^), was fairly consistent across individual mouse samples, and only weakly correlated with variations in cell cycle states, indicating that our observation was not primarily confounded by common technical covariates (Supplementary Note 2, Supplementary Figure 15-18). Intriguingly, the strength of the dyscoordination–age association declined progressively along the hematopoietic differentiation axis (Figure 3B): granulopoietic progenitors retained a positive trend (ρ = 0.34, p < 2.2 x 10^-16^), whereas more terminal populations such as monocytes and macrophages exhibited no consistent age-related increase in dyscoordination. This pattern aligns with previous observations that mutational burden accumulates primarily in long-lived hematopoietic stem and multipotent progenitor cells while terminally differentiated granulocytes experienced minimal mutations^88^. The absence of an age-associated rise in dyscoordination in these populations does not indicate that they are unaffected by aging, since cells can still age through compositional change or coordinated reprogramming without losing expression precision.

To compare with existing measures of transcriptional variance, we applied published methods that quantify transcriptional variability. Specifically, we calculated Euclidean distances of each cell to the average expressions of its cell type^28^, Euclidean distances to the tissue-wide average restricted to invariant genes^28^, and Scallop scores that quantify the stability of each cell’s cluster assignment with resampling^53^. None of these approaches revealed any notable age-related shifts, consistent with the interpretation that they conflate extrinsic and intrinsic sources of variation and thus cannot directly capture dyscoordination (Supplementary Figure 19-21).

In Figure 1C, we drew a conceptual distinction between transcriptional dyscoordination and information-theoretic diversity measures. Here, we examine that difference empirically by considering two such measures that summarize the diversity of a single cell’s own transcriptome: signaling entropy (SCENT)^73^ and potency score (CytoTRACE)^74^. Briefly, both measures capture how broadly transcripts are distributed within a cell and therefore primarily estimate differentiation potency (Methods). As expected, we observed a moderate, though not monotonic, age-related decrease in both SCENT signaling entropy and CytoTRACE potency score across most bone marrow cell types, with the most prominent decline in progenitor cells, such as HPCs, granulocytopoietic cells, and megakaryocyte-erythroid progenitor cells, indicating a substantial age-associated decline in estimated differentiation potential in these cell types (Supplementary Figure 22-23). In line with these opposing trends, cell-level dyscoordination showed modest negative associations with both measures across all cell types (Supplementary Figure 24-25). Thus, transcriptional dyscoordination and information-theoretic diversity measures are related but fundamentally different in what they quantify: dyscoordination tracks age-associated loss of transcriptional coordination while diversity measures capture the decline in differentiation potential with age.

We extended the analysis to additional tissues profiled in the aging mouse atlas, including lungs, limb muscle, and adipose tissue. Across these tissues, we observed a consistent increase in transcriptional dyscoordination with age in their stem cell compartments (Figure 3C). However, in more differentiated or specialized cell types, there was no consistent relationship between dyscoordination and age (Supplementary Figure 26-31). Specifically, Figure 3C illustrates representative trends in four progenitor populations: hematopoietic stem cells in bone marrow, type II pneumocytes in lung, and mesenchymal stem cells in limb and adipose tissue.

Having established this aging signature across mouse stem cells, we asked whether it is also conserved in human cells by examining dyscoordination changes in a single-cell human bone marrow atlas^89^. As in the mouse, human bone marrow cells did not segregate by donor age in the transcriptomic space (Supplementary Figure 32). Nevertheless, at both the cell- and gene-levels, hematopoietic stem and progenitor cells (HSPCs) showed a significant increase in transcriptional dyscoordination with age, particularly across the older cohorts, recapitulating the mouse HSC results (Figure 3F, Supplementary Figure 33-34). This observation reinforces that aging-related gain in transcriptional dyscoordination is a conserved feature of regenerative cell compartments across species.

### Comparison with SenMayo highlights complementary age-associated information

While literature lacks reliable gene-level markers of senescence, SenMayo serves as a multi-gene composite score that can be computed at the single cell level and that has shown utility in hematopoietic aging^54^. As shown in Figure 3C-3D, SenMayo scores increase with age in hematopoietic precursor cells and correlate strongly with transcriptional dyscoordination, yet this is not universal across progenitor cell types from other tissues. This is expected, as SenMayo was trained primarily on cells from the hematopoietic compartment.

We further examined the relationship between transcriptional dyscoordination and the SenMayo score by performing direct cell-by-cell comparison for the hematopoietic stem cells, where both measures performed well (Figure 3E). As expected, the two metrics were significantly correlated, consistent with their shared association with chronological age (*r* = 0.22, *p* < 2.2 x 10^-16^). Importantly, cells with low SenMayo score but high transcriptional dyscoordination predominantly came from old mice. Within each SenMayo score strata, we observed a significant association between age and transcriptional dyscoordination: among cells with similar SenMayo scores, those with higher dyscoordination primarily originated from 24- and 30-month-old mice, the oldest cohorts in the dataset. We also see this effect for type II pneumocytes in lungs, the one other tissue where the two measures are highly correlated (Figure 3C, Supplementary Figure 35-37). This enrichment of high-dyscoordination cells from older mice after stratifying by SenMayo score indicates that transcriptional dyscoordination contains age-associated information not fully captured by the SenMayo score. These findings underscore the potential of dyscoordination to capture transcriptomic disarray in aging cells without reliance on designated senescence markers.

### TCR clonality reveals a dynamic axis of T cell-associated transcriptional dyscoordination

T cell populations are extensively remodeled in aging and disease, through changes in subset composition, clonal expansion, and coordinated differentiation and exhaustion programs (Supplementary Note 3)^33,36,90–92^. Paired TCR information provides a unique opportunity to distinguish these structured changes from loss of transcriptional coordination by linking cells through their clonal history. Cells sharing the same T cell receptor (TCR) clonotype are expected to derive from a common antigen-experienced clone, and clone size provides a quantitative readout of clonal expansion. We therefore analyzed four human T cell datasets with matched single-cell transcriptomes and TCR clonotypes: tumor-infiltrating CD8^+^ T cells from glioma^93^, peripheral CD8^+^ T cells from melanoma patients undergoing combination immune checkpoint blockade (anti-PD-1 and anti-CTLA-4)^94^, and T cells from two healthy blood aging cohorts, Terekhova et al. 2023 and Wang et al. 2025^33,36^. Together, these datasets allowed us to examine whether transcriptional dyscoordination in T cells is associated with clonal expansion, donor age, tumor context, and therapeutic perturbation (Figure 4A).

**Figure 4.**
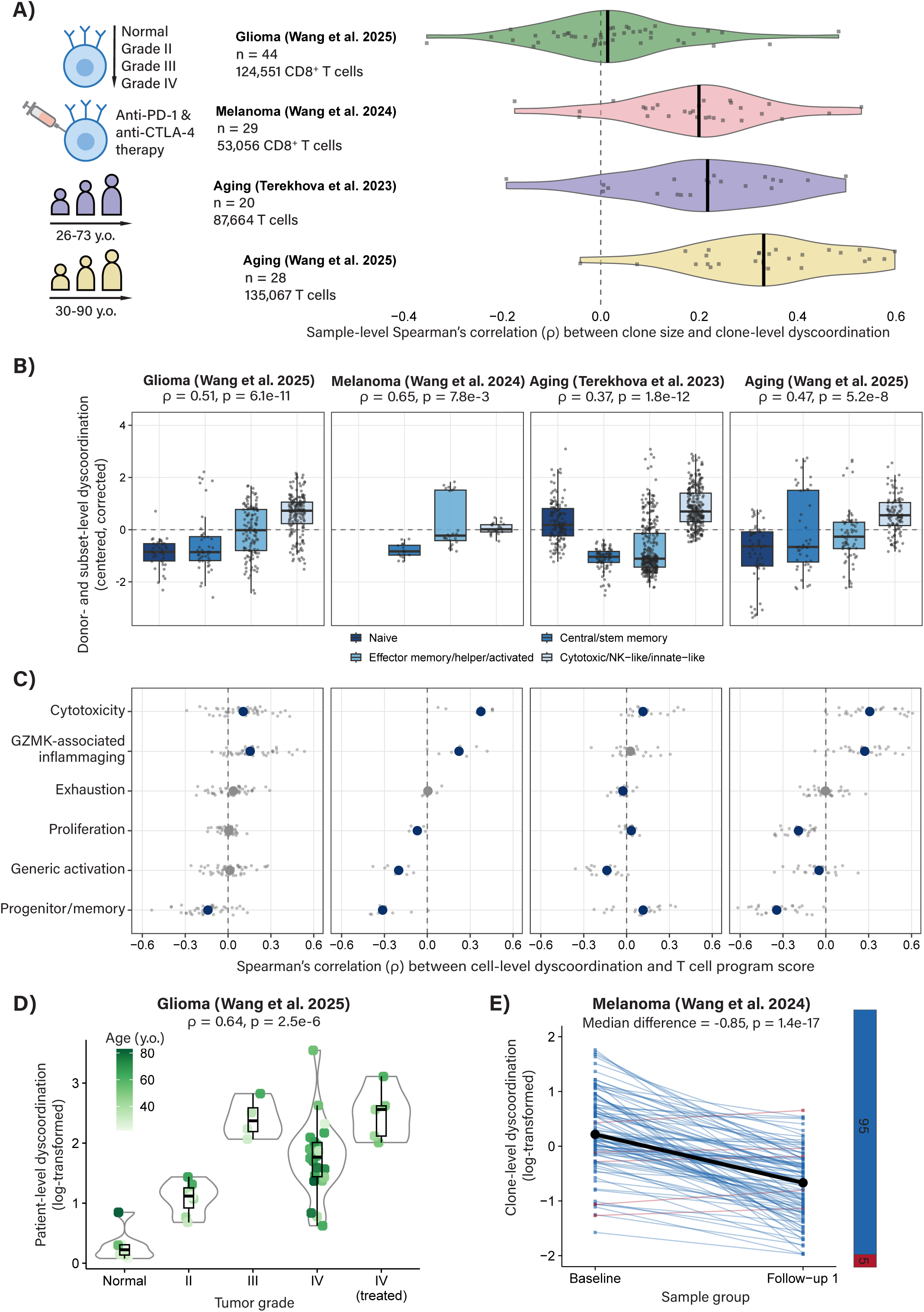
Transcriptional dyscoordination in TCR clones. **A)** Experimental design and violin and box plots showing individual-level Spearman’s correlation between clone size and clone-level dyscoordination across four human T cell datasets: 124,551 glioma CD8^+^ T cells across tumor grades, 53,056 treated melanoma CD8^+^ T cells across timepoints after therapy, 87,664 T cells from 26-to 73-year-old healthy human donors, and 135,067 T cells from 30-to 90-year-old healthy human donors. Each dot represents an individual (Melanoma: individual × timepoint), black bars denote medians, and the dashed line denotes ρ = 0. **B)** Box plots showing transcriptional dyscoordination across four coarse T cell subset groups in each of the four datasets. Each point is one donor-by-subset median of depth-corrected cell-level dyscoordination centered within each donor. Spearman’s correlations and their *p*-values are indicated above each panel. **C)** Dot plots showing donor-level Spearman’s correlations between cell-level dyscoordination and 6 T cell gene program scores. Gray dots in the background represent donor-level correlations, and colored points represent median correlations across donors, colored by whether the donor-level signed-rank test shows nominal significance (blue: significant; gray: insignificant). **D)** Violin and box plots showing patient-level dyscoordination by tumor grade in glioma T cells, with each point indicating one patient colored by age. Spearman’s correlation and its *p*-value are indicated above. **E)** Line plots showing changes in within-clone dyscoordination of treated melanoma T cells from baseline to the first follow-up after immune checkpoint blockade, with one line per clone and color indicating direction (blue: decrease; red: increase). The black line denotes the median trajectory. Median per-clone difference and paired Wilcoxon *p*-value are indicated above.

For each dataset, we first quantified cell- and gene-level transcriptional dyscoordination within the relevant T cell subtypes and study strata, including donor age, glioma grade, and time after immune checkpoint blockade (Supplementary Figures 38-49). Dyscoordination estimates were robust to sequencing depth and were only weakly related to cell cycle activity (Supplementary Note 2, Supplementary Figures 50-65). We then defined TCR clones within each donor by the TCR beta sequence and computed clone-level dyscoordination as the average cell-level dyscoordination of cells within each clone.

Because both T cell state composition and baseline dyscoordination vary substantially across individuals, we tested the association between clone size and dyscoordination primarily within each donor or patient. Across all four cohorts, larger clones tended to exhibit higher dyscoordination within individuals (Figure 4A-4B, Supplementary Figures 66-69). This association was observed robustly across donors, in both cancer and healthy aging cohorts, indicating that the association between clonal expansion and transcriptional dyscoordination is a general phenomenon not restricted to cancer.

Intriguingly, in the PBMC lifespan cohort (Wang et al. 2025), the pooled clone-size-dyscoordination association was nearly absent at the cohort level (*ρ* = 0.020, *p* = 2.0 x 10^-1^), whereas stratification by donor consistently recovered a positive association (average *ρ* = 0.337, positive in 96% of donors; Figure 4A). Thus, between-individual differences can obscure a reproducible within-individual association between clonal expansion and transcriptional dyscoordination.

Individual TCR clones, however, can contain cells spanning multiple transcriptional states. We therefore next asked whether dyscoordination itself differs systematically across T cell states, irrespective of clone-size stratification. Despite substantial differences in annotation schemes across the four datasets, T cell subset identity was strongly associated with dyscoordination in every cohort (*p* ≤ 2.3 x 10^-8^ across all four datasets; Figure 4B). For cross-dataset comparison, we grouped annotated subsets into broad categories representing naive, central/stem memory, effector memory/helper/activated, and cytotoxic/NK-like/innate-like states (Supplementary Table 2). Cytotoxic and several innate-like populations generally showed higher dyscoordination, whereas many CD4 helper and central memory populations showed lower dyscoordination. Detailed subset rankings revealed that this pattern did not follow a simple differentiation hierarchy: naive cells were not uniformly the least dyscoordinated, with tissue-resident, GZMK-associated, and other effector populations varying substantially across datasets (Supplementary Figure 70-73), and such orderings across subset groups remained unchanged without within-donor centering or depth correction (Supplementary Figure 74). Thus, T cell state itself represents a major axis of transcriptional dyscoordination, with cytotoxic and effector-associated populations showing particularly prominent increases.

These state-wise differences also help clarify the association between clonal expansion and dyscoordination. Expanded clones are enriched in antigen-experienced T-cell compartments, and the aggregate clone-size association can therefore arise in part from differences in state composition. When clone-size–dyscoordination associations were examined within annotated states, selected disease populations retained significant associations, such as GZMK-expressing, tissue-resident memory T cells in glioma (*ρ* = 0.259, *p* = 8.2 x 10^-6^) and activated and effector clones showed significant within-state associations in treated melanoma (*ρ* = 0.184, *p* = 6.8 x 10^-7^; *ρ* = 0.178, *p* = 3.7 x 10^-3^), whereas within-state associations were largely absent in the two healthy blood cohorts (Supplementary Figures 75-78). Specifically, in the Terekhova et al. 2023 study, only the CD4 Th1/Th17 subset showed a significant within-state association, and no individual state reached significance in the Wang et al. 2025 cohort. Thus, the relationship between clonal expansion and dyscoordination reflects both the enrichment of expanded clones in high-dyscoordination states and, in selected disease-associated states, an additional within-state association between expansion and loss of transcriptional coordination.

To characterize the transcriptional programs associated with these state differences, we next examined six predefined T cell programs encompassing cytotoxicity, GZMK-associated inflammaging, exhaustion, proliferation, generic activation, and progenitor/memory states (Supplementary Table 2). For each donor, we correlated cell-level dyscoordination with the corresponding program score and then compared the direction and magnitude of these associations across datasets (Figure 4C), with the same ranking of programs obtained without global depth correction (Supplementary Figure 79). Cytotoxicity showed the most consistent positive association with dyscoordination, with positive donor-median correlations in all four cohorts (median *ρ* = 0.11-0.38). The GZMK-associated inflammaging program was also positively associated with dyscoordination in three of the four datasets, whereas the progenitor/memory program showed the opposite relationship in three. By contrast, exhaustion, proliferation, and generic activation showed less consistent associations across datasets. These findings reinforce the subset-level analysis by identifying cytotoxic effector state as the most reproducible transcriptional correlate of high dyscoordination. However, they do not establish that cytotoxic differentiation causes dyscoordination. Instead, loss of transcriptional coordination is preferentially associated with particular functional T cell states rather than with activation or proliferation in general.

We next asked whether transcriptional dyscoordination is additionally shaped by the tissue environment in which T cells reside. Glioma provides a useful setting because tumor grade reflects increasing malignancy and microenvironmental stress. In tumor-infiltrating T cells from glioma, patient-level dyscoordination significantly increased with tumor grade, particularly across the earlier grades (*ρ* = 0.643, *p* = 2.5 x 10^-6^, Figure 4D). This association persisted after adjustment for patient age and clonal expansion (Supplementary Figure 80), indicating that higher-grade tumors are associated with increased T cell transcriptional dyscoordination beyond differences attributable to clonal expansion alone. The gradient was present within individual T cell subsets rather than reflecting a shift in subset composition, with a significantly positive patient-level association between tumor grade and dyscoordination in 9 of 11 subsets (Supplementary Figure 39-40). Thus, the malignant tissue environment provides an additional independent axis associated with loss of transcriptional coordination.

Finally, we used longitudinal melanoma samples collected before and after immune checkpoint blockade to determine whether T cell transcriptional dyscoordination can be remodeled by combination therapy. At both the cell- and gene-levels, transcriptional dyscoordination dropped sharply after treatment initiation across most cell types, with negative median patient-level changes observed in all nine annotated subsets, indicating that the drop was not confined to compositional shift in any particular subset (Supplementary Figure 42-43). Still, such a decrease could in principle reflect a compositional turnover, such as loss of high-dyscoordination clones or expansion of low-dyscoordination clones. To distinguish these possibilities, we tracked TCR clones present at both baseline and the first follow-up timepoint (week 3) and decomposed the clone-level dyscoordination shift (-0.56) into within-clone, between-clone, new-clone, and lost-clone components, weighted by clone size. The within-clone component was the dominant net contributor (-0.48), where persisting clones showed a marked reduction in their own dyscoordination after therapy (median difference = -0.85, *p* = 1.4 x 10^-17^, Figure 4E). This reduction in dyscoordination was independent of clone size, implying that it reflects a general tightening across clones rather than an effect driven by a few highly expanded ones (*ρ* = -0.14, *p* = 0.15, Supplementary Figure 81). On the other hand, compositional turnover contributed little overall, as the influx of new clones (-0.68) was largely offset by the loss of existing clones (+0.57), and re-weighting of persistent clones (between-clone) was negligible (+0.02, Supplementary Figure 82). These results suggest that immune checkpoint blockade primarily decreases T cell dyscoordination by tightening the transcriptional coordination within persisting clones. Although immune checkpoint blockade also remodels the clonal composition of the repertoire, this compositional turnover does not itself account for the decrease in dyscoordination.

Overall, these analyses show that transcriptional dyscoordination in T cells jointly reflects clonal history and cellular state. Expanded clones tend to be more dyscoordinated, and dyscoordination varies across T cell subsets with cytotoxic effector states showing the most reproducible elevation across datasets. In cancer, dyscoordination is further associated with tumor grade and can decrease within the same TCR clone following immune checkpoint blockade. Thus, rather than representing a fixed property of clonal identity or differentiation state, transcriptional dyscoordination captures a dynamic feature of T cell state that reflects clonal expansion, functional phenotype, environmental context, and therapeutic perturbation.

### Rat kidney aging: transcriptional dyscoordination reveals segment-specific decline in proximal tubule cells

Kidney aging is accompanied by well-described changes in cellular composition and coordinated within-cell-type transcriptional programs involving metabolism, mitochondrial function, inflammation, and injury responses^45,95^. Here, we investigate whether cells within an anatomically defined cellular compartment become less transcriptionally coordinated with age. We applied the transcriptional dyscoordination framework to aging rat kidney, where we used a combination of DEFND-seq and 10x Multiome to simultaneously profile transcriptome and chromatin accessibility within single cells. 22 kidney samples were collected from 18 rats at 16, 30, 56, and 82 weeks of age, and our analysis focused on the 49,405 proximal tubule (PT) cells (30,593 after quality control) identified by clustering and annotation using canonical markers. PT cells from all age groups were broadly intermixed in the transcriptomic embedding (Figure 5A) despite substantial age-associated changes detectable in conventional differential expression analysis (Supplementary Note 3).

**Figure 5.**
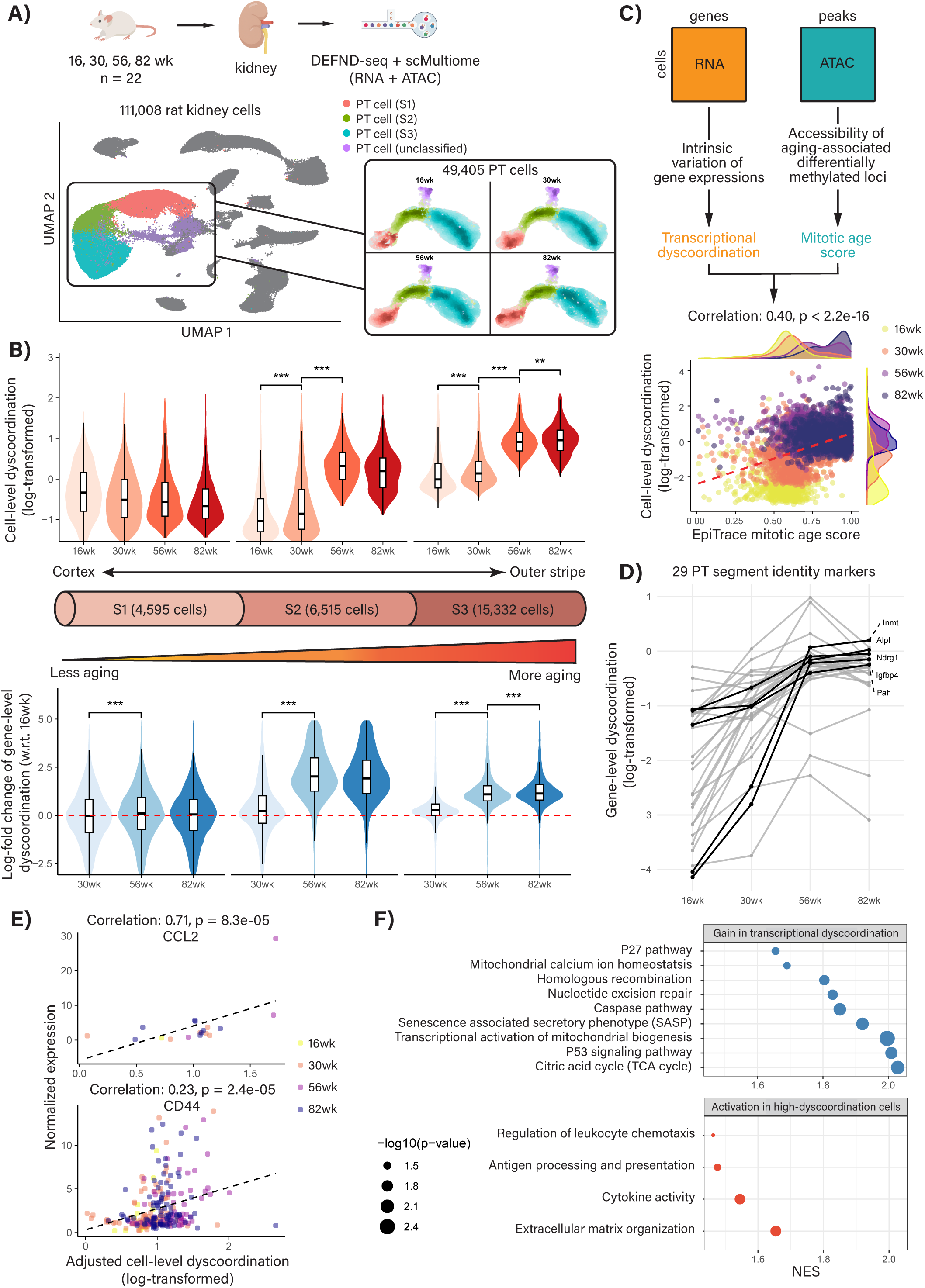
Transcriptional dyscoordination of PT cells in aging rat kidney. **A)** Experimental design and UMAP visualization of 111,008 kidney cells with DEFND-seq or scMultiome profiles from 16-to 82-week-old rats. Cell types and ages of 49,405 PT cells are highlighted in the UMAP. **B)** Violin and box plots showing log-transformed cell-level dyscoordination and log-fold change of gene-level dyscoordination with respect to 16-week-old rats by age group for S1 to S3 PT cells after quality control (26,442 cells). **C)** In parallel with transcriptional dyscoordination, EpiTrace mitotic age score is computed by quantifying accessibility of aging-associated differentially methylated loci. Scatter plot shows the correlation between log-transformed cell-level dyscoordination and EpiTrace mitotic age score of PT cells, with marginal density plots by age group to the side. Red dashed line denotes the best-fit regression line. **D)** Line plots showing log-transformed gene-level dyscoordination across age groups for 29 PT segment identity markers. **E)** Scatter plots showing correlations between normalized expressions of two inflammation and stress-related markers, *CCL2* and *CD44*, and log-transformed cell-level dyscoordination adjusted for total counts of each cell. Black dashed line denotes the best-fit regression line. **F)** Significantly enriched pathways for genes with gain in transcriptional dyscoordination (top) and activation in high-dyscoordination cells (bottom). Dot size denotes –log10(p-value) and x-axis denotes normalized enrichment score (NES).

Despite the lack of overt clustering by age, there is a significant increase in transcriptional dyscoordination with age in PT cells, both at the cell- and gene-levels (Supplementary Figure 83), which remained robust after global adjustment for sequencing depth and after subsampling each age group to a common cell number. The increase of dyscoordination with age was reproducible across biological replicates and was only weakly related to cell cycle state (Supplementary Note 2, Supplementary Figure 84-87). The aging trend is particularly significant between 16 to 30 weeks, and between 30 to 56 weeks.

The proximal tubule can be anatomically and functionally divided into three segments: segment 1 (S1), segment 2 (S2), and segment 3 (S3), each characterized by distinct marker genes and physiological roles. PT cells can be assigned to different segments based on segment-specific marker genes. Upon stratifying cells by segment (Figure 5B), we found that the age-associated increase in transcriptional dyscoordination was pronounced in segments 2 and 3, but not in 1. By contrast, published methods estimating transcriptional variability did not capture this aging trend (Supplementary Figure 88-90). The two information-theoretic entropy measures, which estimate differentiation potency rather than transcriptional coordination, decreased with age in PT cells (Supplementary Figure 91). Thus, the age-associated increase in transcriptional dyscoordination is segment-dependent, with stronger and more consistent changes in S2 and S3 than in S1.

These findings align with existing literature indicating differential vulnerability among proximal tubule segments. S2 and S3 segments are more prone to ischemic injury and nephrotoxic damage, reflecting their higher metabolic activity and exposure to reabsorption stressors^96–100^. In contrast, S1 segments exhibit greater resilience due to their distinct protective mechanisms and cellular environments: located more proximal to the cortical labyrinth, S1 cells receive comparably more oxygen supply than S2 and S3 cells to counter their high metabolic demand, thereby reducing ischemic injury risk. Furthermore, S3 segments have been shown to possess regenerative capacity in response to recurring injuries^101–103^. The stronger age-associated dyscoordination in S2 and S3 is therefore consistent with known segment-specific differences in injury response, although our analysis does not directly measure susceptibility to injury.

We hypothesized that increased transcriptional dyscoordination in the aging rat kidney should be associated with molecular drift at the level of chromatin organization. To test this, we leveraged the matched chromatin accessibility data generated alongside the cellular transcriptomes and applied EpiTrace, a recently published method for estimating cellular mitotic age from chromatin accessibility profiles^70^. EpiTrace computes a global aging score based on the accessibility of loci previously identified as differentially methylated with age in bulk tissue methylation studies, offering a chromatin-derived readout of replicative history. The original EpiTrace study provided strong evidence that this score could reliably stratify cells by mitotic age across diverse contexts.

We compared EpiTrace mitotic age estimates with transcriptional dyscoordination scores at single-cell resolution (Figure 5C, Supplementary Figure 92). As shown in Figure 5C, the two measures are significantly correlated across biological age groups (*r* = 0.40, *p* < 2.2 x 10^-16^), despite being derived from entirely distinct molecular modalities and conceptual frameworks. Even among PT cells derived from organisms of the same chronological age, particularly at 16 and 30 weeks, transcriptional dyscoordination and EpiTrace scores remain significantly associated, suggesting that the two measures capture partially overlapping aspects of cellular aging (Supplementary Figure 93). This moderate cross-modal association provides convergent evidence that transcriptional dyscoordination relates to a chromatin-derived dimension of cellular aging, while the two measures remain biologically distinct.

### Two modes of gene association with transcriptional dyscoordination: loss of expression precision and activation in high-dyscoordination cells

To further interpret the biological correlates of transcriptional dyscoordination, we turned our attention to identifying dyscoordination-associated genes and pathways. In this analysis, we asked two questions for each gene: (1) does the gene exhibit an increase in transcriptional dyscoordination with age, indicating that its expression becomes less predictable across cells, and (2) is the gene’s expression correlated with cell dyscoordination level, indicating that its activation is linked to cellular response to transcriptional disarray. These two modes of association capture different biological phenomena and should not be conflated. We hypothesize that genes involved in maintaining stable cellular identities, such as PT segment markers, should lose regulatory precision over time and therefore exhibit increased gene-level dyscoordination with age. In contrast, genes involved in inflammation and senescence pathways are often induced in response to damage or stress, and thus for these genes, we do not necessarily expect an increase in their gene-level dyscoordination. Instead, we postulate that their expression levels should positively correlate with cell-level dyscoordination scores, as cells with high transcriptional dyscoordination are more likely to exhibit hallmarks of senescence. Thus, the two analyses distinguish loss of expression precision within a gene from coordinated activation of gene programs in dyscoordinated cellular states, providing complementary views of how aging-associated transcriptional changes relate to dyscoordination.

In line with the first question, we curated a set of 29 PT segment identity markers and observed significant age-dependent increases in their gene-level dyscoordination, particularly across 16, 30, and 56 weeks (Supplementary Table 3). This observation indicates that PT cells progressively lose segment fidelity during initial stages of aging (Figure 5D). Addressing the second question, we identified inflammation and stress-related markers whose normalized expression levels showed significantly positive correlations with cell-level dyscoordination scores, after adjusting for total count differences across cells and multiple testing. Two examples include *CCL2*, a chemokine secreted by stressed or injured PT cells to mediate inflammation (*r* = 0.71, *p* = 8.3 x 10^-5^)^104^, and *CD44*, a fibrosis-related marker that characterizes maladaptive, high-dyscoordination states of PT cells in chronic kidney diseases (*r* = 0.23, *p* = 2.4 x 10^-5^)^105^, highlighting the coordinated activation of inflammation and stress-response programs in high-dyscoordination cellular states (Figure 5E). Together, these findings suggest that transcriptional dyscoordination, whether assessed at the gene level or cell level, captures key aging-related signatures of PT cells: the breakdown of segment distinctions and the activation of pro-inflammatory pathways.

Beyond these marker-level observations, we next systematically examined pathway-level associations with transcriptional dyscoordination in PT cells (Figure 5F). We ranked all genes by the magnitude of their age-dependent changes in dyscoordination using Spearman’s ρ and identified 5,022 genes with significantly positive changes. Pathway enrichment analyses revealed strong representation of apoptosis and senescence-related pathways such as p27 and p53 signaling, the caspase pathway, and the senescence-associated secretory phenotype (SASP); mitochondrial pathways such as calcium ion homeostasis, biogenesis, and the TCA cycle; and DNA repair processes, including nucleotide excision repair and homologous recombination (Supplementary Table 4). The involvement of these pathways suggests that the loss of expression precision not only intersects directly with canonical aging hallmarks but also captures key molecular alterations such as mitochondrial dysregulation and genome instability. At the cell level, using correlation values for 994 genes whose expression levels were significantly associated with transcriptional dyscoordination (Supplementary Figure 94), we found enrichment in pathways related to immune activation and extracellular remodeling, including antigen processing and presentation, cytokine activity, leukocyte chemotaxis, and extracellular matrix organization (Supplementary Table 5). These results indicate that disordered cellular states are characterized by coordinated activation of inflammatory and fibrotic responses, consistent with the emergence of maladaptive programs during aging in PT cells.

### Human kidney aging: transcriptional dyscoordination converges with somatic genomic instability in aging human tubular epithelium

Existing studies have likewise shown that aging of the human kidney is accompanied by changes in tubular cell states, immune-cell representation, and coordinated metabolic and injury-response programs^45,106,107^. We next asked whether the dyscoordination observed in rat proximal tubule generalizes to human tubular epithelium. For this purpose, we analyzed a single-nucleus Multiome atlas of human kidneys comprising 134,458 nuclei sampled from 32 healthy donors spanning 30 to 79 years of age (Supplementary Figure 95). We focused on three major tubular epithelial populations: the proximal tubule (PT, 48,397 nuclei), comprising proximal convoluted tubule (PCT; S1 and S2) and proximal straight tubule (PST; S3), the thick ascending limb (TAL, 30,214 nuclei), and the distal convoluted tubule (DCT, 24,386 nuclei). These cell types were prioritized in this analysis, because they arise from a shared nephron-progenitor lineage and represent successive tubular segments most frequently implicated in chronic kidney disease and age-related decline. In all three compartments, cells across age groups are intermixed in the low-dimensional embedding (Figure 6A).

**Figure 6.**
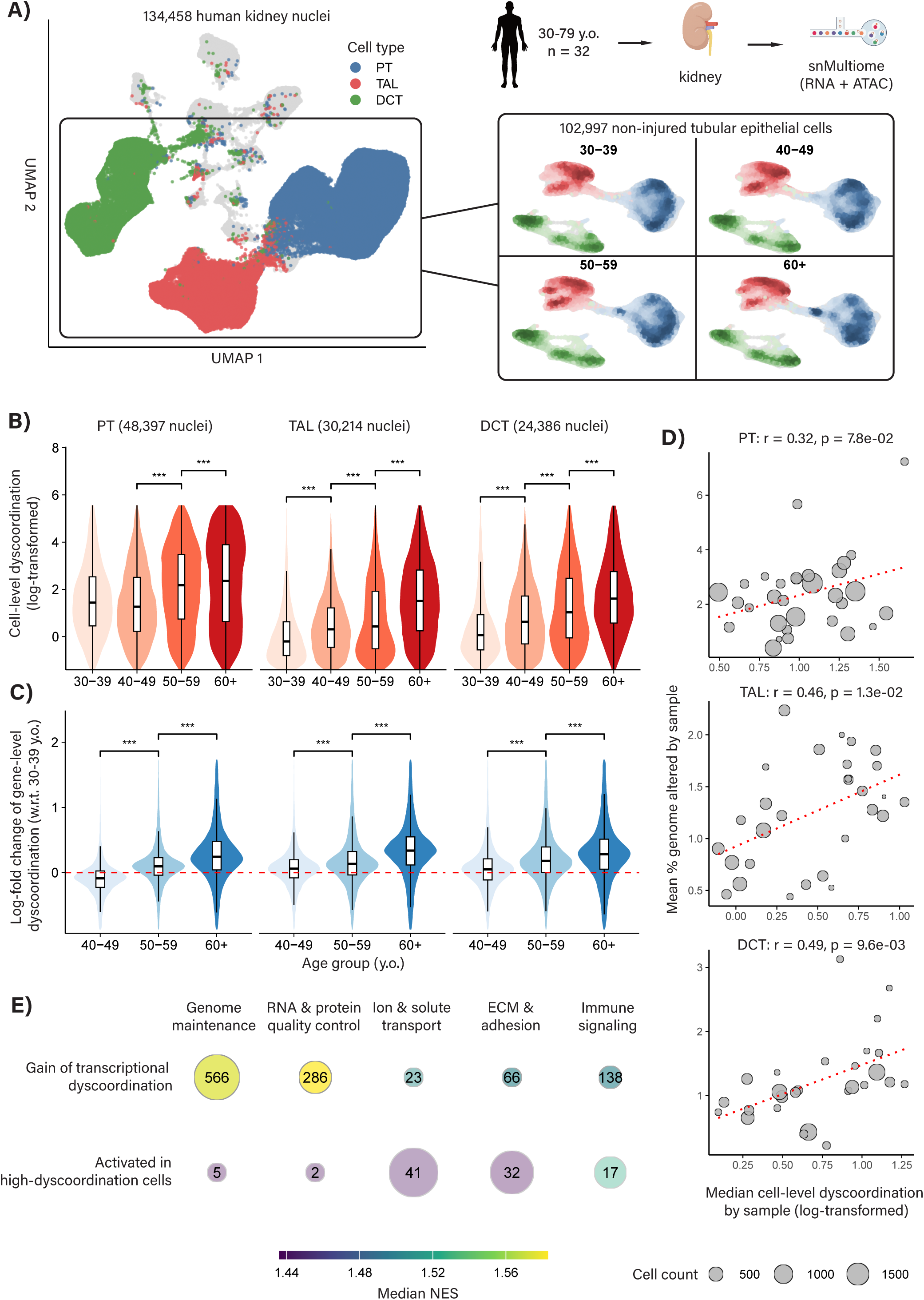
Transcriptional dyscoordination of tubular epithelial cells in aging human kidney. **A)** Experimental design and UMAP visualization of 134,458 kidney nuclei with snMultiome profiles from 30-79-year-old healthy human donors. Cell types and ages of 102,997 tubular epithelial cells are highlighted in the UMAP. **B)** Violin and box plots showing log-transformed cell-level dyscoordination by age group for PT, TAL, DCT cells after quality control. **C)** Violin and box plots showing log-fold change of gene-level dyscoordination with respect to 30-39-year-old donors. **D)** CHASM estimates somatic genome instability by detecting mCAs in single-cell chromatin accessibility. Scatter plots showing the donor-level association between donor-median log-transformed cell-level dyscoordination and donor-mean log-transformed percentage of genome altered. Red dotted lines denote the best-fit regression line. **E)** Functional themes of significantly enriched pathways for genes with gain in transcriptional dyscoordination (top) and activation in high-dyscoordination cells (bottom). Size of and the number within each bubble denotes the number of pathways in that theme, filled color denotes the median NES across pathways in that theme, and opacity denotes the fraction of the theme’s FDR-significant pathways.

Across all three tubular epithelial cell types, transcriptional dyscoordination increased significantly with age at both the cell and gene levels (Figure 6B-6C). Unlike the rat proximal tubule, where age-associated rise was confined to specific tubular segments, age-associated dyscoordination increased broadly across PT, TAL, and DCT, implying a broad, segment-independent degradation of transcriptional coordination within the human tubular epithelium (Supplementary Figure 96-97). This broad gain remained robust to sequencing depth, was reproducible across donors, and was only weakly related to cell cycle activity, indicating that measured technical covariates do not account for it (Supplementary Note 2, Supplementary Figure 98-101). For comparison, we applied transcriptional variability and diversity measures to the same cells and found no significant change across age groups except for SCENT signaling entropy, which declined modestly with age (Supplementary Figure 102-106). This decline was weaker and less consistent than the dyscoordination increase, indicating that the information-theoretic potency measures do not reliably capture signatures of aging in more heterogeneous human tissue.

We next asked whether the observed loss of transcriptional coordination is associated with somatic genome instability. To examine this, we applied CHASM, a method for detecting mosaic chromosomal copy number alterations (mCAs) in single-cell chromatin accessibility data^108^. As cells with mCAs in non-neoplastic tissue are rare and have not undergone clonal expansion, CHASM tests each cell’s copy number state against a cell type-matched background to distinguish true alterations from technical noise with high specificity. For each nucleus, CHASM summarizes mCA burden as the proportion of the genome affected by copy number alterations, providing a DNA-level readout of somatic genome instability. Thus, we were able to compare transcriptional dyscoordination of 43,018 nuclei with each nucleus’s corresponding genome instability measures from CHASM in the 3 tubular epithelial cell types. As mCAs are rare clonal events, we assessed this association at the donor level by aggregating all nuclei with paired profiles and retaining donors with at least 50 nuclei in each cell type (31, 29, and 27 donors for PT, TAL, and DCT, respectively). Donor-median cell-level dyscoordination and donor-mean percentage of genome altered showed significant positive association in TAL and DCT (*r* = 0.46, *p* = 1.3 x 10-2; *r* = 0.49, *p* = 9.6 x 10-3) and marginally significant positive association in PT (*r* = 0.32, *p* = 7.8 x 10-2; Figure 6D). Therefore, donors whose tubular epithelial cells were more transcriptionally disordered also carried more somatic genome damage, coupling purely expression-based transcriptional dyscoordination to an independent, chromatin-derived measure of genome instability.

We next applied the same two-mode gene and pathway analysis used in the rat kidney to examine whether the biological correlates of dyscoordination were conserved in human tubular epithelium. As in rat PT cells, we distinguished genes whose expression became less predictable with age from genes whose expression was preferentially elevated in highly dyscoordinated cells. Among 24,924 genes evaluated for age-associated changes, 3,549 showed significant increases in gene-level dyscoordination and were enriched for pathways involved in genome maintenance and RNA/protein quality control, including heterochromatin organization, mitotic spindle checkpoint, p53 signaling, and ubiquitin-dependent proteolysis (Figure 6E, Supplementary Table 6)^109,110^. This extends our findings in rat kidney, where genes gaining dyscoordination were also enriched for DNA repair and other cellular maintenance pathways, but in the human kidney the connection to genome integrity was especially prominent. Notably, this enrichment converged with the independent CHASM analysis: donors with higher transcriptional dyscoordination also carried greater somatic mCA burden, providing a cross-modal link between loss of transcriptional coordination and genome instability.

The second mode of association was also strikingly concordant with the rat kidney. Genes whose expression was elevated in high-dyscoordination cells were enriched for ion and solute transport, extracellular-matrix and adhesion programs, and immune signaling, including pathways related to Th1 responses, integrin interactions, and collagen-containing extracellular matrix (Supplementary Table 7). In rat PT cells, high-dyscoordination states were similarly associated with inflammatory and extracellular-remodeling programs. Thus, across species, genes that lose expression precision with age and genes activated in dyscoordinated cells define two reproducible but distinct biological signatures: disruption of cellular maintenance programs and coordinated activation of inflammatory and fibrotic responses^111,112^. Because the human donors were histologically normal, the enrichment of fibrosis-associated programs in highly dyscoordinated tubular cells further suggests that these transcriptional changes can be detected before overt tissue pathology.

Together, these results identify a conserved pattern of age-associated transcriptional dyscoordination across rat and human kidney, with an important difference in its anatomical distribution: the increase is segment-selective within rat PT but broader across human PT, TAL, and DCT. Across both species, genes that lose expression precision with age are enriched for cellular-maintenance programs, whereas highly dyscoordinated cells activate inflammatory and extracellular-remodeling programs. The human kidney Multiome data further connect this transcriptional phenotype to somatic genome instability, as dyscoordination is associated both with CHASM-inferred mCA burden and with disorder of genome-maintenance pathways.

### Liver aging: transcriptional dyscoordination reveals zone-specific vulnerability of the regenerative compartment

Liver aging remodels both tissue composition and hepatocyte transcriptional programs, including pathways involved in metabolism, inflammation, and regeneration^113,114^. Against this background of coordinated age-associated change, we investigated whether hepatocytes also lose transcriptional coordination within defined cellular states. To jointly profile transcriptional and chromatin-level aging, we performed 10x single-cell Multiome analysis on liver tissue collected from young (4 months) and old (22 months) mice (Figure 7A). After quality control, we obtained 183,005 high-quality liver cells across both age groups. Our analysis focused on the 66,816 high-quality hepatocytes identified through clustering and marker-based annotation.

**Figure 7.**
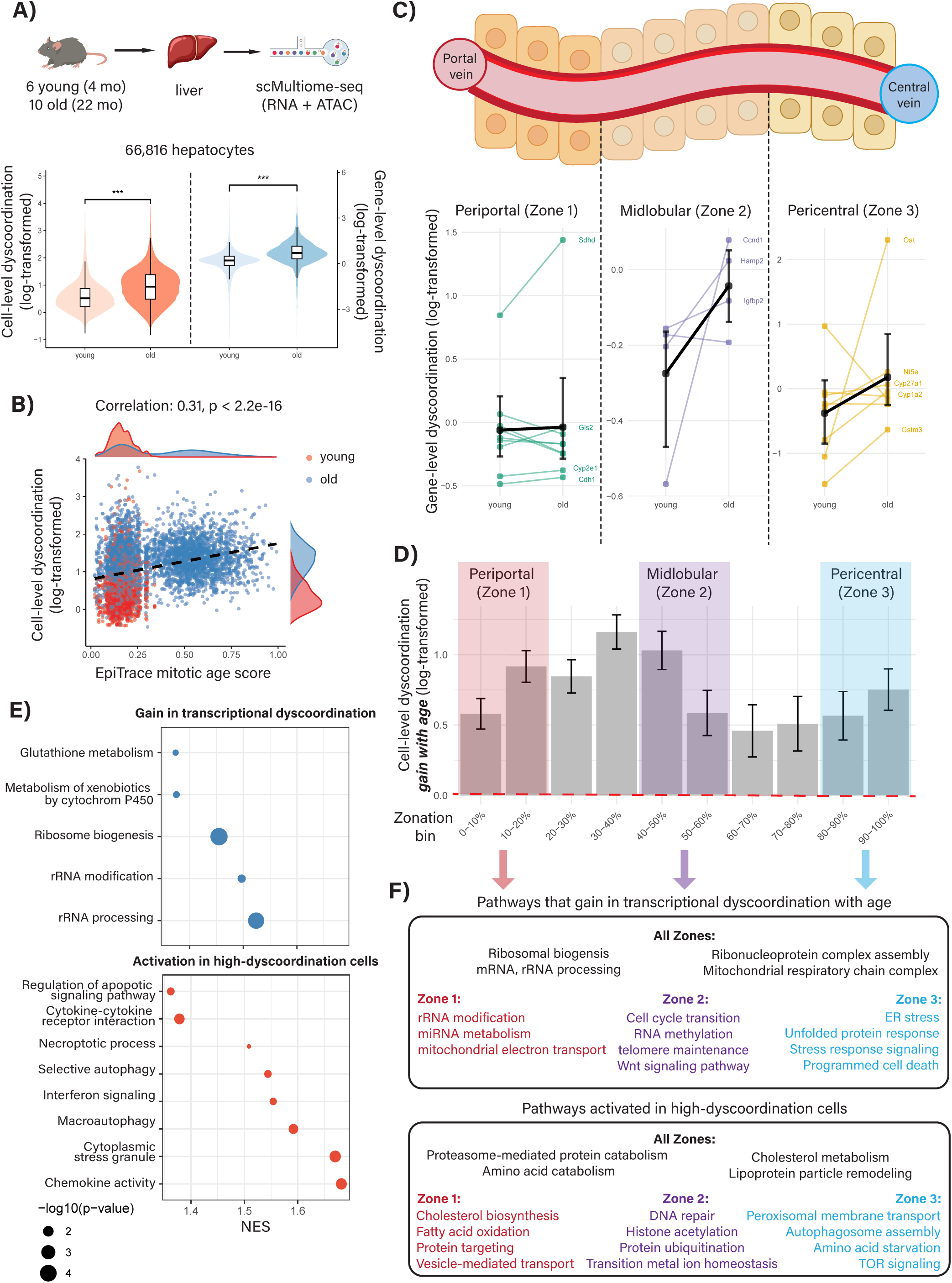
Transcriptional dyscoordination of hepatocytes in aging mouse liver. **A)** Experimental design and violin and box plots showing log-transformed cell-level and gene level dyscoordination of 66,816 hepatocytes with scMultiome profiles from young (4 month) and old (22 month) mice. **B)** Scatter plot showing correlation between log-transformed cell-level dyscoordination and EpiTrace mitotic age score of hepatocytes, with marginal density plots by age group to the side. Black dashed line denotes the best-fit regression line. **C)** Schematic of hepatic zonation including periportal (zone 1), midlobular (zone 2), and pericentral zones (zone 3) and line plots showing log-transformed gene-level dyscoordination of 21 zonation identity markers in their respective zones. Genes with age-related dyscoordination increase are labeled. Black dots indicate mean dyscoordination and I-shaped bars indicate 95% confidence intervals. **D)** Histograms showing the log-transformed mean cell-level dyscoordination gain with age across zonation bins, with bins corresponding to each hepatic zone highlighted. **E)** Significantly enriched pathways for genes with gain in transcriptional dyscoordination (top) and activation in high-dyscoordination cells (bottom). **F)** Summary of significantly enriched pathways for genes with zone-specific gain in transcriptional dyscoordination (top) and activation in zone-specific high-dyscoordination cells (bottom) across zone 1-3.

Standard transcriptomic embeddings did not segregate hepatocytes into discrete young and old populations (Supplementary Figure 107), as aging-related transcriptomic changes are subtle and distributed. Despite this, transcriptional dyscoordination, quantified at both the gene and cell levels, was significantly elevated in hepatocytes from old mice relative to young controls (*p* < 2.2 x 10^-16^, Figure 7A). This change persisted after global correction for sequencing depth and after subsampling age groups to a common cell number, was reproducible across mice, and was only weakly related to cell cycle activity (Supplementary Note 2, Supplementary Figure 108-111). As for the kidney, we further computed mitotic age by applying EpiTrace to the cell-matched chromatin accessibility data (Supplementary Figure 112). EpiTrace scores were significantly positively correlated with transcriptional dyscoordination (Figure 7B), echoing the alignment observed in the aging kidney. Nevertheless, only 1 of the 3 published transcriptional variability methods were able to capture the elevated transcriptional variance (Supplementary Figure 113-115). The two information-theoretic entropy measures likewise decreased with age in hepatocytes (Supplementary Figure 116-117).

Despite its strong statistical significance (*r* = 0.31, *p* < 2.2 x 10^-16^), the correlation between transcriptional dyscoordination and EpiTrace’s estimate of mitotic age is moderate, reflecting the difference in what they capture. Transcriptional dyscoordination increases uniformly across hepatocytes from old mice. In contrast, EpiTrace scores in old mice exhibit a bimodal distribution, revealing a subpopulation of hepatocytes with elevated mitotic age while another remains comparable to that of young mice. As hepatocytes are largely post-mitotic, this bimodal pattern likely reflects partial regenerative activity in aging livers: the subpopulation with noticeably higher EpiTrace scores may correspond to cells that have undergone replication during regeneration, whereas the remainder retain mitotic histories similar to young hepatocytes. Thus, the two measures capture distinct aspects of the aging process: EpiTrace reflects cumulative mitotic history, while transcriptional dyscoordination captures a broader dysregulation of gene expression independent of cell division.

The liver is organized into lobules with hepatocytes arranged along defined spatial zones that perform distinct metabolic and detoxification functions, starting at the periportal region (zone 1), extending through the midlobular region (zone 2), and ending at the pericentral region (zone 3). We explored how aging affects hepatocyte spatial identity by analyzing a curated set of 21 gene markers for zone identity (Supplementary Table 8). Gene-level transcriptional dyscoordination estimates revealed zone-specific aging effects: pericentral and midlobular markers showed the strongest increases in dyscoordination with age, while most periportal markers remained relatively stable (Figure 7C). A notable exception among periportal markers was *SDHD*, a key gene in mitochondrial oxidative metabolism, which exhibited substantial increase in dyscoordination in old cells. These results suggest that hepatocyte zonal specialization becomes increasingly unstable with age, particularly in zones 2 and 3.

To further assess the change in hepatocyte zonal integrity with age, we assigned each cell a zonation score defined as the ratio of total pericentral marker expression to the combined expression of pericentral and periportal markers. As expected, zone 1 (periportal) and zone 3 (pericentral) markers showed high expressions near 0 and 1, respectively (Supplementary Figure 118). Zone 2 (midlobular) markers, which are not used in the zonation score calculation, are low near 0 and 1 and high in the middle, serving as an independent validation that this score effectively captures the zonation gradient (Supplementary Figure 119). We then ranked all cells by their zonation score and divided them into 10 equal sized bins. Then, we averaged the cell-level dyscoordination for all cells in each zonation bin, separately for young and old mice. Across both age groups, we observed that the mean cell-level dyscoordination and the proportion of high-dyscoordination cells (top 5%) varied systematically across the zonation axis and tracked closely with one another (Supplementary Figure 120). Within each zonation bin, old cells consistently showed significantly higher dyscoordination than their young counterparts, and this difference was not attributable to changes in cell proportions, which remained close to 50:50 (Supplementary Figure 121). Interestingly, the magnitude of this age-related dyscoordination gain exhibits significant spatial variations: moderate in the periportal zone, elevating to a peak in the midlobular zone, declining towards pericentral, then peaking again at the terminal of the pericentral zone (zone 3) (Figure 7D). These patterns highlight the spatial differences in age-associated transcriptional dyscoordination, which will be examined in greater detail in the subsequent zone-specific analysis.

To examine the association between transcriptional dyscoordination and the aging-related processes of senescence and inflammation, we curated 138 genes enriched in senescence and pro-inflammation processes and assessed the correlation of their expression with cell-level dyscoordination, following the same approach applied to aging kidney PT cells (Supplementary Table 8). In total, 29 (21.0%) of the markers showed significant positive correlations, which is significantly higher than the background frequency of 8.8%, and the frequency among the zonation markers genes (4.7%). As in the aging rat kidney, this is the second of the two gene-level questions defined above and is distinct from the analysis of zonation markers in Figure 7C: that analysis asked which genes lose expression precision with age, whereas this one asks which genes are activated in high-dyscoordination cells.

The large number of senescence- and pro-inflammation genes with significantly positive associations with cell-level dyscoordination supports a close connection between increased transcriptional dyscoordination and activation of stress and senescence programs in hepatocytes (Supplementary Figure 122). Example genes include *AXL*, a cell differentiation- and carcinogenesis-associated gene whose expression is heightened in parallel to chronic liver disease progression^115^, *CCND2*, a cell cycle regulator that was previously found to increase its expression in senescent hepatocytes^116^, *IL1B*, which promotes hepatic inflammation through inflammasome activation in mice^117^, and *MMPS*, which is involved in extracellular matrix remodeling and contributes to hepatic fibrosis and apoptosis during injury^118^. Overall, genes whose expression levels increased in high-dyscoordination cells (Supplementary Figure 123) were enriched for inflammatory and immune signaling pathways (e.g., cytokine–cytokine receptor interaction, chemokine signaling, interferon response), cell death programs (e.g., apoptotic signaling, necroptosis), autophagy-related processes (e.g., macroautophagy, selective autophagy), and RNA stress responses (e.g., cytoplasmic stress granule formation) (Figure 7E, Supplementary Table 9). These findings support the conclusion that hepatocytes with elevated transcriptional dyscoordination engage stress-response and inflammatory programs characteristic of dysfunctional or damaged cellular states.

In contrast, genes with the greatest increase in transcriptional dyscoordination with age were enriched in RNA and ribosome-related processes (e.g., rRNA processing and modification, ribosome biogenesis) and metabolism (e.g., glutathione metabolism, metabolism of xenobiotics by cytochrome P450) (Figure 7E, Supplementary Table 10). The increased dyscoordination of RNA and ribosome pathways suggests reduced precision of proteostasis with age. The increased dyscoordination in antioxidant and detoxification pathways points to a breakdown in the coordination of hepatocellular defense mechanisms, potentially rendering aged hepatocytes more susceptible to oxidative stress and xenobiotic toxicity.

### Liver aging: zone-specific gene dyscoordination and functional alterations with age

Using the zonation scores computed for each cell, we next examined how aging-related changes differ across spatial zones of the liver lobule. We first computed zone-specific gene-level dyscoordination to identify genes exhibiting the greatest loss of transcriptional coordination with age. Notably, genes with the largest dyscoordination increases were not necessarily those highly expressed ones, indicating that age-related transcriptional variability is not only driven by expression level (Supplementary Figure 124). We also extracted differentially expressed genes between high and low dyscoordination cells within each zone to characterize zone-specific functional reprogramming accompanying transcriptional dysregulation. Below, we highlight genes identified through both approaches.

Starting with the periportal zone, aging hepatocytes showed greater dyscoordination across genes involved in rRNA maturation and modification, miRNA metabolism, and mitochondrial electron transport, reflecting instability in the transcriptional and translational machinery that coordinates RNA processing and mitochondrial function. Notably, *ASS1*, involved in protein catabolism and hepatic lipid metabolism^114,119,120^, showed the greatest age-related dyscoordination increase in periportal hepatocytes (log2FC = 0.47), suggesting heightened transcriptional variability in core anabolic programs. In parallel, high-dyscoordination periportal hepatocytes exhibited activation of pathways in energy production and oxidative metabolism, including steroid and cholesterol biosynthesis, fatty acid oxidation, amino acid (e.g., L-serine, tryptophan) metabolism, and carbohydrate homeostasis. Enrichment of protein synthesis and trafficking processes, such as translation initiation, protein targeting, COPII-coated vesicle budding, and retrograde vesicle-mediated transport, were also observed in periportal hepatocytes likely as an adaptive compensation towards rising metabolic demand (Figure 7F, Supplementary Tables 11 and 12).

Pericentral hepatocytes also displayed a widespread metabolic stress profile. At the gene level, increase in transcriptional dyscoordination was primarily associated with endoplasmic reticulum (ER) signatures, including organization, vesicle-mediated transport, protein localization and exit, stress response, and unfolded protein response (UPR). As the cellular center of protein folding, modification, and secretion, ER is expected to withstand a particularly heavy metabolic load in the pericentral zone that specializes in xenobiotic metabolism and detoxification. Upon ER stress in the form of accumulation of unfolded and misfolded proteins, UPR is triggered to restore its homeostasis and promote cell survival^121^. Moreover, we observed elevated activity of pathways regulating integrated stress response signaling, apoptotic processes, and programmed necrotic cell death, suggesting activation of broader cellular damage response programs. Examples of genes with age-related dyscoordination increase include *GLUL* (log2FC = 0.26), a gene involved in glutamine synthesis and ammonia detoxification^122,123^, and *IQGAP2* (log2FC = 0.86), a gene linked to cellular apoptosis and suppression of hepatocellular carcinoma^124^. In parallel, activation of stress-response programs was evident in high-dyscoordination pericentral hepatocytes. For instance, peroxisome-associated programs, including protein import, membrane transport, and organization, were prominently enriched. As a key oxidative organelle, peroxisome mediates metabolism of fatty acids and detoxification of reactive oxygen species (ROS) to reduce oxidative stress and maintain redox homeostasis^125,126^. Autophagic processes such as autophagosome assembly, lysosomal and nuclear microautophagy, and late nucleophagy were also upregulated, reflecting elevated cellular recycling in response to damaged proteins and organelles. Concurrent enrichment of amino acid starvation response and target of rapamycin (TOR) signaling pathways further indicates adaptive nutrient sensing and energy conservation under the high metabolic burden of aging pericentral hepatocytes. Together, these findings indicate an uneven functional engagement of detoxification and metabolic programs resulting from an imbalanced cellular stress specific to the pericentral cells (Figure 7F, Supplementary Tables 11 and 12).

Finally, our data showed that the midlobular zone, a major compartment implicated in homeostatic hepatocyte renewal, was most strikingly altered. We found that genes with increased transcriptional dyscoordination were associated with tissue regeneration, cell cycle transition, telomere maintenance, and multiple RNA regulatory mechanisms, such as RNA methylation, translation, and mRNA splicing. For example, we detected significant age-related dyscoordination increase in *AXIN2* (log2FC = 4.74), a WNT-responsive gene that contributes to liver homeostasis and repair^127,128^. As WNT signaling is normally enriched in pericentral hepatocytes^129^, its increase in dyscoordination in midlobular zone is consistent with our finding of age-related loss of zone identity. Increased transcriptional dyscoordination with age was also observed in *TERT* (log2FC = 1.35), a critical gene for regenerative capacity through maintenance of metabolic balance and telomere integrity^130–132^. Correspondingly, pathway enrichment analysis revealed that high-dyscoordination midlobular hepatocytes were enriched in pathways related to genome maintenance and transcriptional regulation (e.g., DNA repair, histone acetylation, transcription, and RNA processing), proteostasis and post-translational control (e.g., protein ubiquitination, stabilization, modification, and phosphorylation), and stress response and cellular homeostasis (e.g., ROS response, acetyl-CoA metabolism, and transition metal ion homeostasis). These pathways focus on maintaining genomic integrity, protein quality, and metabolic balance, which are core processes underpinning the regenerative capacity of midlobular hepatocytes. Together, these results demonstrate that aged zone 2 hepatocytes exhibit alteration of programs linked to regenerative capacity, consistent with age-associated vulnerability of this compartment (Figure 7F, Supplementary Tables 11 and 12).

In summary, trends observed across the three spatial zones of hepatocytes reflect their underlying functional specialization. Affected pathways in zone 1 and zone 3 hepatocytes align with longstanding evidence that they experience distinct but significant stressors with age. Periportal cells engage in oxidative metabolism and protein synthesis and operate under oxygen-rich conditions that favor metabolic rigidity and transcriptional stability, whereas pericentral cells are exposed to hypoxia and high concentrations of xenobiotic metabolites, rendering them prone to injury and stress responses. In parallel, the dyscoordination peak in the midlobular zone echoes our findings in the bone marrow, where long-lived precursor populations such as hematopoietic stem cells show the most prominent age-associated increases in dyscoordination. In the liver, midlobular (zone 2) hepatocytes have been shown to contribute disproportionately to tissue renewal throughout homeostasis, and our results raised the possibility that this regenerative compartment is particularly vulnerable to transcriptional disorganization with age^131,133^.

## Discussion

Together, these findings establish transcriptional dyscoordination as a reproducible measure of loss of predictable expression coordination that connects the classical concept of intrinsic transcriptional noise^76^ to modern single-cell transcriptomic data. Across perturbational, lineage-resolved, and aging datasets, the measure responds to induced damage and senolytic depletion, varies with T cell clonal and environmental context, and shows moderate cross-modal associations with chromatin-based mitotic age and somatic genome instability. Unlike existing approaches, transcriptional dyscoordination does not rely on curated gene markers or supervised training, instead offering a first-principles-based method for quantifying this dimension of cellular aging across diverse cell types without retraining. As single-cell atlases continue to grow, transcriptional dyscoordination may serve as a scalable and broadly applicable tool to map where loss of transcriptional coordination accumulates across tissues, lineages, and conditions.

An important implication of this work is that transcriptional dyscoordination offers a principled way to prioritize cell populations most vulnerable to aging-related dysfunction. Across tissues, we observed that dyscoordination highlights compartments known to exhibit early molecular disruption or functional decline with age, even when they are not easily identifiable by conventional marker-based approaches. We therefore view dyscoordination as a marker-free way to prioritize cellular contexts for biological interpretation, rather than as a direct measure of tissue-level vulnerability or function.

A consistent theme across tissues is the heightened dyscoordination observed in regenerative compartments (hematopoietic stem cells, segment 3 proximal tubule cells, zone 2 hepatocytes, and mesenchymal stem cells) relative to more terminally differentiated cells. These compartments share sustained roles in tissue maintenance or regenerative response and may therefore experience prolonged exposure to molecular damage and repeated demands on transcriptional control. Our observation that these same cells exhibit the largest increases in transcriptional dyscoordination suggests that the very systems responsible for tissue maintenance may be particularly susceptible to stochastic dysregulation. While this may reflect their intrinsic vulnerability, it may also point to selective pressure for tighter transcriptional control in differentiated cells.

Disentangling these effects, and understanding whether increased dyscoordination causally impairs regenerative potential, remains an important avenue for future research.

In stem cell populations, increased dyscoordination may reflect age-related degradation of regulatory systems that normally maintain transcriptional fidelity. However, the same accumulated damage can also fuel clonal evolution, which intensifies with age^134^. Such clonal dynamics give rise to genetically distinct lineages that can propagate into mature blood cells, a phenomenon clinically recognized as clonal hematopoiesis of indeterminate potential (CHIP)^135^. Structured transcriptional heterogeneity is conceptually distinct from dyscoordination. If a mutation, differentiation program, or environmental response induces a sufficiently represented and coherent expression change across cells, that program is expected to enter the predictable expression component rather than substantially increase dyscoordination (Supplementary Figure 125). Rare or weakly represented cellular programs may be incompletely learned and can contribute to the residual, as demonstrated by our simulation analysis. We therefore interpret transcriptional dyscoordination as a measure of loss of predictable expression coordination, rather than as a pure or exhaustive measure of molecular stochasticity.

Our findings also clarify how transcriptional dyscoordination relates to information-theoretic entropy measures such as SCENT signaling entropy and CytoTRACE potency score. As both measures summarize the diversity of an individual cell’s own expression distribution, they primarily quantify differentiation potency rather than transcriptional dysregulation. Consistent with this distinction, the two moved in opposite directions with age across multiple tissue environments, and notably this opposition was sharpest in the long-lived stem and progenitor cells. This trend reflects that diversity measures are informative where clear differentiation gradients exist and are most responsive to aging in progenitor populations whose potency erodes over time, whereas in terminally differentiated cells their signal becomes diluted. By contrast, transcriptional dyscoordination can rise with age in both progenitor and differentiated populations, with the strongest and most consistent effects observed in regenerative compartments. The negative correlation between the two measures in regenerative compartments thus reflects a co-occurrence of two aging effects rather than a shared underlying quantity, as these measures capture fundamentally different aspects of cellular aging from dyscoordination.

Transcriptional dyscoordination is not a fixed record of accumulated damage. In primary mouse muscle stem and progenitor cells, dyscoordination rose after doxorubicin-induced damage and fell again after senolytic clearance with ABT-263, so the measure tracks the induction and removal of a damaged cellular state rather than chronological time alone. In peripheral T cells sampled before and after combination immune checkpoint blockade, the decrease in dyscoordination was carried by clones that persisted across timepoints rather than by turnover of the repertoire, indicating that a lineage-defined population can regain transcriptional coordination within weeks. Whether the restoration of coordination contributes to therapeutic benefit, or merely accompanies it, cannot be determined from these data. These observations nonetheless establish that dyscoordination is a modifiable property of living cells and therefore a plausible readout for interventions intended to restore cellular fidelity.

At the gene level, we provide a conceptual framework for classifying aging-associated transcriptional changes along two axes: genes whose expression becomes more disordered with age (i.e., increasing gene-level dyscoordination), and genes whose expression levels are associated with the dyscoordination of the cell (i.e., activated in high-dyscoordination cells). These categories capture distinct biological processes. The former reflects a breakdown in regulatory precision, and we find it enriched among genes defining cell identity such as zonation markers in the liver. The latter likely represents genes activated in response to cellular damage or dysregulation, including those involved in inflammation, stress response, and cell death. These axes are not redundant: they identify genes with distinct roles in aging biology and provide complementary entry points for understanding how aging reshapes our cells. Together, these complementary analyses separate loss of expression precision from coordinated response programs and provide distinct entry points for interpreting how aging reshapes cellular regulation.

Transcriptional dyscoordination should therefore be interpreted as a model- and context-referenced measure, not as a universal aging ruler or a pure measure of molecular stochasticity. The present implementation specifically uses the UMI measurement model and regularized gene-wise prediction from SAVER, and alternative reference models may produce different residual quantities. Rare or weakly represented coherent programs can be incompletely learned and enter the residual, and absolute dyscoordination values are not intended for direct comparison across transcriptionally distinct cell types. Cross-modal associations with EpiTrace and somatic genome instability are also convergent rather than causal. Accordingly, dyscoordination complements, rather than replaces, analyses of cell type composition, coordinated differential expression, and functional aging phenotypes.

## Acknowledgements

This work was supported by joint DMS/NIGMS grant DMS-2245575 and NIH grant 1R56AG081351 and 5R01GM149671 (Y.Y. and N.R.Z.), and NIH grant U54 AG079758 (M.G.T., K.N.M, A.E.D., C.M., K.Y.L., S.M., K.Y., B.R., Q.Y., E.S., A.W., P.D.A.)

## Author Contributions

Conceptualization: N.R.Z.

Method development: Y.Y., P.R.H., N.R.Z.

Data analysis and interpretation: Y.Y., X.E.C., P.R.H., S.H., N.R.Z.

Data generation and preprocessing: M.G.T., H.W., K.N.M., A.E.D., C.M., K.Y.L., S.M., K.Y., Q.Y., E.S., A.W.

Consultation and guidance: B.J., P.W., P.D.A., A.C.H

Manuscript writing: Y.Y., M.G.T., P.D.A., N.R.Z.

Supervision: P.W., B.R., P.D.A., N.R.Z.

## Code and Data Availability

All codes implementing transcriptional dyscoordination, together with analysis scripts reproducing figures shown in this study, are available at https://github.com/EddieYang1222/transcriptional-dyscoordination.

Rat kidney and mouse liver single-cell Multiome datasets generated for this study will also be deposited. All other datasets analyzed here are previously published and cited in Methods.

## Methods

### Data preprocessing

The computation of transcriptional dyscoordination takes as input a gene by cell raw UMI count matrix generated from single-cell RNA sequencing. By the nature of scRNA-seq data, many genes have nonzero counts in only few or zero cells. These extremely sparsely observed genes are not expected to be informative and are thus excluded from our analysis. As data quality varies across datasets, exact details for quality control are provided in the later data sections. Before computing transcriptional dyscoordination, we apply a library size normalization to each cell, where a size factor is computed, defined as the cell’s total UMI count divided by the mean total UMI count across all cells, and the count for each gene is divided by this size factor. For the aging rat kidney and aging mouse liver datasets generated in this study, clusters were identified with the `FindClusters` function from `Seurat`^136^ and assigned by marker-based biological interpretation using cluster-level differential expression and established canonical markers rather than through automated label transfer alone. For rest of the studies, cell type annotations were taken directly from each original study when provided.

### Transcriptional dyscoordination

Let *Y_gc_* denote the observed, library-size normalized UMI count for gene *g* in cell *c*. Previous works have shown that UMI-based scRNA-seq data have technical noise that can be approximated by a Poisson^137,138^ distribution with a library size scaling factor, and thus, in line with previous expression models^72,139–142^, we assume that *Y_gc_*, conditioned on the true expression *λ_gc_* of gene *g* in cell *c*, has distribution:

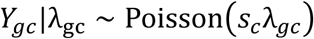

where *s_c_* is the cell-specific size factor. We don’t observe the true expression (*λ_gc_*), but as described in the main text, variations in *λ_gc_* across all cells can be decomposed into two components: predictable coordinated (extrinsic) variation, which is represented by the learned expression reference, and residual biological (intrinsic) variation, which is not explained by that predictable structure. To model this, we use a Gamma prior on *λ_gc_*:

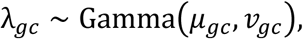

where *μ_gc_* denotes the prior mean and *ν_gc_* denotes the prior variance. We assume that *μ_gc_* is the “predictable” component of gene expression, while the deviation in *λ_gc_* from *μ_gc_* is the unpredictable component. We denote the estimate of *μ_gc_* from the data by *µ̂_gc_*, i.e., *µ̂_gc_* is the regularized predictable expression reference underlying our geometric manifold interpretation. In the current implementation, we estimate this predictable expression reference using SAVER^72^. SAVER predicts each gene from genome-wide expression using penalized regression, selects the degree of shrinkage through cross-validation, and explicitly models UMI sampling variation. Then, given *µ̂_gc_*, the true expression of gene *g* in cell *c* can be estimated via its Bayesian posterior mean, which essentially evaluates to a weighted average of the prior mean *µ̂_gc_* and the observed data *Y_gc_*^72^. Given both {*λ̂_gc_*} and {*µ̂_gc_*}, transcriptional dyscoordination quantifies the deviations {*λ̂_gc_* − *µ̂_gc_*}, which are summarized at the gene or cell level. Since the *expected* deviations vary according to *μ_gc_*, we need to standardize the observed deviations based on some variance model. Thus, we introduce a variance-scaled squared deviation, *δ_gc_*:

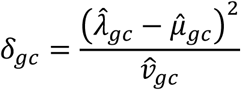

This definition partially relies on the SAVER-based predictable expression model, and alternative reference models may yield different residual quantities and are not assumed to produce equivalent dyscoordination estimates. Here, *v̂_gc_* serves as a normalization factor that adjusts for baseline variability expected of each gene in each cell. The estimation of *v_gc_* is based on one of three candidate models: constant coefficient of variation (CV), constant Fano factor (FF), and constant variance, as described below:

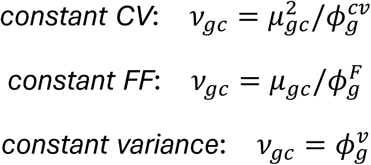

Marginal likelihood of the observed counts, given *µ̂_gc_*, is calculated for each model, and the one that attains the highest likelihood is chosen. Then, *v̂_gc_* is computed based on this model and the maximum likelihood estimate of the appropriate dispersion parameter, *φ̂_g_*. The predictable expression reference *µ̂_gc_* and the baseline variance *v̂_gc_* are both computed through models fitted on the full cell population to provide a common reference for evaluating residual biological variation within the analyzed population. Examples of how these parameters are computed are introduced in later sections.

Finally, we summarize the variance-scaled squared deviation *δ_gc_* into cell- and gene-level transcriptional dyscoordination measures. At the cell level, dyscoordination is calculated by averaging *δ_gc_* across genes while fixing *c*:

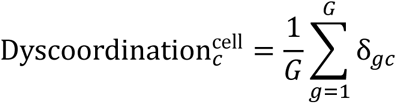

This reflects the degree to which cell *c*’s transcriptome as a whole deviates from its predictable expression reference. At the gene level, dyscoordination is calculated by averaging residuals across cells within a pre-defined stratum *S* (e.g., a given cell type in a given age group) to quantify the residual biological variability of each gene within the specified subpopulation *S*:

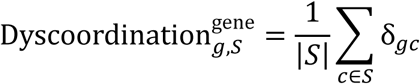

To ensure reliable estimation, gene-level dyscoordination is only calculated for strata *S* containing at least 10 cells in our analyses. Overall, cell-level dyscoordination allows direct assessment of transcriptional disorder for a given cell, while gene-level dyscoordination enables identification of genes and relevant pathways that exhibit increase in dyscoordination within specific cell strata. By nature, gene-level dyscoordination is a stratum-specific score, and log-fold changes relative to a baseline age group are used only for comparative display in downstream analyses.

### Allele-specific dyscoordination analysis

We analyzed 6 F1 hybrid mouse single-cell RNA-seq datasets in which a gene’s two alleles serve as cis-matched replicate reporters and preprocessed to remove genes with low total counts^77–82^. In each dataset, we obtained allelic-specific counts at heterozygous SNPs, aggregated them to genes, and summed the counts of two alleles into a total expression matrix on which transcriptional dyscoordination was computed. For gene g in cell c, let *A_gc_* and *B_gc_* be the UMI counts assigned to its two alleles, and *N_gc_* = *A_gc_* + *B_gc_*. Let *p_g_* = median 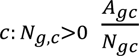 be the gene’s typical allele fraction so that constitutive allelic imbalance is absorbed into the reference baseline. We then define allelic discordance as:

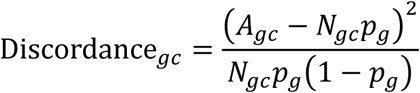

where the numerator is the squared deviation of the observed allele count from the expected value and the denominator is the binomial variance of *A_gc_* used for scaling, so that discordance has an expectation of 1 under pure binomial sampling. Discordance was summarized at the gene level by averaging over all cells within each stratum and the cell level by averaging over a cell’s qualifying heterozygous genes. Gene-level associations between dyscoordination and discordance were quantified by an expression-controlled partial Spearman’s correlation. Specifically, we first log-transformed and rank-transformed gene-level transcriptional dyscoordination, allelic discordance, and mean predicted expression by the manifold. Then, ranked dyscoordination and discordance were separately regressed on the ranked mean expression, and the Pearson correlation between the two residual vectors was recorded across all genes and the top 500 HVGs. For the genetic divergence analysis, each cross was assigned an approximate genome-wide number of SNVs separating its non-reference strain from C57BL/6J, obtained from the Mouse Genomes Project, the same variant catalog used to call the heterozygous sites for allele counting. Allele counts were processed with N-masked realignment for the two datasets prone to reference mapping bias (van der Veeken et al. 2019, Weber et al. 2026) and with standard alignment otherwise. At the cell level, both measures were made comparable before averaging. Within each gene, qualifying entries (at least 5 allele-informative reads, 0.05 < *p_g_* < 0.95) were first converted to normal scores and residualized on their log-transformed allelic sequencing depth and the cell’s log-transformed library size then averaged for each cell over the entire gene panel. As each single cell yields only few informative entries, we also pooled each cell’s score over its 15 nearest neighbors in the principal component space with re-residualization on the neighborhood-averaged depth and entry count. Cell-level associations were then quantified by a partial Spearman’s correlation adjusting for library size, entry count, and cell type, with 95% confidence intervals acquired from 500 cell bootstraps and significance evaluated from a permutation null that shuffles cells within depth deciles before pooling. To interpret the weak cell-level associations, we estimated each measure’s split-half reliability by randomly splitting each cell’s genes into two halves, scoring the cell from each half, and correlating the two halves across cells (Spearman-Brown correction). Reliability ceiling, defined as the geometric mean of the two reliabilities, represents the maximum correlation attainable given the measurement noise, with which the observed cell-level associations scaled across analysis configurations.

### Induced cellular damage and senescence dyscoordination analysis

For cellular damage, single-cell RNA-seq (Smart-seq2) count matrices were downloaded for 133 KYSE-180 ESCC cells exposed to different doses of ionizing radiation (41 with 0 Gy, 92 with 12 Gy)^83^. All cells and genes passed the pre-processing and were retained. Gene- and cell-level transcriptional dyscoordination were separately computed for 0 Gy and 12 Gy populations.

For cellular senescence, single-cell RNA-seq of 52,997 FACS-sorted FAP (29,214 cells) and SC (23,783 cells) from mouse skeletal muscle were obtained across three conditions: phosphate-buffered saline as vehicle control (PBS), doxorubicin-induced senescence (DOXO), and doxorubicin followed by the senolytic ABT-263 (navitoclax) to deplete senescent cells (DABT). After quality control, gene- and cell-level transcriptional dyscoordination were computed within each cell type in each condition.

### Tabula Muris Senis dyscoordination analysis

Raw droplet-based gene count matrices were downloaded from Tabula Muris Senis, a single-cell transcriptomic atlas of aging mouse tissues^24^. Selected tissues for our analysis include bone marrow, lung, limb muscle, and adipose tissue, with cells sampled at 1, 3, 18, 21, 24, and 30 months of age. Annotations for cell type and age groups were directly taken from the accompanying metadata. In the original study, extensive quality control has already been conducted across all tissues for the raw count matrices: genes not expressed in at least 3 cells and then cells without at least 250 detected genes were removed. Additionally, cells with fewer than 2500 total UMI counts were excluded in droplet-based libraries. Therefore, we only performed one additional quality control step to make sure that dyscoordination calculation applies only to expressed genes: across all cells within each tissue, genes with zero total counts were removed.

To compute transcriptional dyscoordination, we estimated the predictable-expression reference *μ_gc_* and the baseline variance *ν_gc_* based on models fit on combined cell types within each tissue. The latent true expression λ*_gc_* was inferred separately for each cell type × age group stratum (e.g., 18-month-old hematopoietic precursor cells). Cell-level dyscoordination was calculated by averaging the residuals across all retained genes in each cell, while gene-level dyscoordination was calculated once for each stratum in each gene. Note that some of the strata were missing, since not all cell types were present across all ages and strata containing fewer than 10 cells were also excluded.

Three clustering-based measures used for comparison with transcriptional dyscoordination were computed with the decibel (version 1.0) and Scallop (version 1.3.0) R packages^53^. Briefly, we used decibel to compute Euclidean distance from each cell to its cell type expression average and Euclidean distance from each cell to its tissue-wide expression average using a set of invariant genes. Invariant genes are selected by their coefficients of variation, as described in the package documentation. Scallop is used to compute a membership score, which is the frequency with which each cell is assigned to its most frequently assigned cluster label. This score quantifies transcriptional stability and is therefore interpreted in the opposite direction from variability or dyscoordination measures.

Two single-cell transcriptional diversity measures, SCENT^73^ and CytoTRACE^74^, were employed to illustrate fundamental differences between transcriptional dyscoordination and information-theoretic entropy methods. Briefly, SCENT integrates a cell’s gene expression with a protein-protein interaction (PPI) network and computes the entropy rate of a random walk over the network, with higher values reflecting the promiscuous signaling state of undifferentiated cells. CytoTRACE instead derives a potency score from the number of distinctly expressed genes per cell, treating a more diverse transcriptome as a hallmark of lower differentiation. Signaling entropy was computed with `SCENT` R package and differentiation potency score was computed with `CytoTRACE` R package (version 0.3.3). For signaling entropy, we normalized the raw count matrix, mapped mouse genes (or rat genes in following analyses) to their human orthologs with `biomaRt`^143^, and integrated the resulting matrix with the package’s built-in PPI network using `DoIntegPPI`. Signaling entropy was then computed with `CompSRana`, with higher values indicating transcriptomic nature of undifferentiated cells. For differential potency score, we used the default algorithm of `CytoTRACE` function, which orders cells by the number of distinctly expressed genes refined against a co-varying gene signature to produce a score between 0 and 1, with higher scores indicating less differentiated cells. Both measures were computed once per cell and compared against cell-level dyscoordination across cell types and age groups.

SenMayo is a set of 125 gene signatures proposed as robust cross-tissue and cross-species markers of cellular senescence and SASP activity^54^. In our analysis, SenMayo scores were calculated by extracting the inferred true expressions of these genes and computing enrichment scores using the `escape` R package (version 2.4.0)^144^. These scores were then compared against cell-level dyscoordination across stem cell populations in each tissue.

### Aging human bone marrow dyscoordination analysis

We analyzed single-cell RNA-seq data of bone marrow mononuclear cells from healthy human donors spanning early adulthood to old age^89^. In total, 87,349 cells from 25 samples in 21 donors (age 24-84) were annotated into 18 cell types and subsequently studied. To remove the confounding effect of sequencing depth, three donors in the 58-60-year-old range (samples A, G, P) with over 50% higher mean depths than the cross-age median were excluded. To make all age groups evenly distributed and comparable, we focused on the 12 donors of at least 46-year-old and re-classified cells into 5 age groups (46-50, 55-57, 58-60, 66-67, and 84-year-old). For quality control, we removed cells with fewer than 200 detected genes or fewer than 500 total UMI counts, as well as genes expressed in less than 1% of cells. In total, we retained 55,319 cells and 10,465 genes for downstream analysis.

### TCR clone dyscoordination analysis

We downloaded four human T cell datasets with matched TCR labels: glioma CD8^+^ T cells (124,551 cells from 44 patients of tumor grade I, II, III, IV, and IV (treated))^93^, melanoma CD8^+^ T cells treated with anti-PD1 and anti-CTLA4 combination therapy (53,056 cells from 8 samples at baseline and 3 follow-up timepoints)^94^, and two healthy aging cohorts (87,664 T cells from 20 donors (2 males + 2 females for each age bin, age 26-73)^33^ and 135,067 T cells from 28 donors (age 30-90))^36^. For quality control, we removed cells with fewer than 200 detected genes or fewer than 500 total UMI counts, as well as genes expressed in less than 0.1% of cells. Clonal identities were defined within each patient/donor by TCR-β CDR3, and clone-level dyscoordination were computed as the mean cell-level dyscoordination across all cells belonging to the same clone. For subset-specific clonality analysis, only cell types with at least 20 clones of at least 2 cells were retained. For longitudinal clonality analysis, only clones with at least 3 cells present at either the baseline or the first follow-up timepoint were analyzed.

For each donor and annotated T cell subset, cells were summarized by the median depth-corrected cell-level dyscoordination, which required at least 10 cells per donor and subset. Subsets present in fewer than 5 donors were dropped. Depth correction uses the global LOESS residual described in Supplementary Note 2. Each donor-by-subset median was then centered within its donor by subtracting that donor’s mean across its own subsets, which removes the donor’s overall level and effects from between-donor covariates, including sequencing depth, age, disease grade, and batch. Whether dyscoordination differs across subsets at all was tested by a Friedman rank test on the complete donor-by-subset matrix, blocking on donor and restricted to subsets present in at least 80% of donors. For cross-dataset comparison, subsets were assigned to four ordered groups: (1) naive, (2) central or stem memory, (3) effector memory, helper or activated, and (4) cytotoxic, NK-like or innate-like, with full assignment list in Supplementary Table 2. Other regulatory, interferon-high, proliferating, double-negative and NME1+ subsets were ranked but left ungrouped as distinct lineages or transient states. This grouping is a coarse summary that mixes lineage, differentiation state, and effector function and does not describe a developmental trajectory. The ordered trend was quantified with Spearman’s correlation between group index and centered subset median across all donor-by-subset combinations and tested within donor by a Wilcoxon signed-rank test on the per-donor difference between the last group and first two groups.

Then, we scored six T cell-associated gene programs for each cell: cytotoxicity, GZMK-associated inflammaging, exhaustion, progenitor/memory, proliferation, and generic activation. Scores were computed with the `AddModuleScore` function from `Seurat` for treated melanoma, Terekhova et al. 2023 and Wang et al. 2025 aging cohorts. For glioma, whose expression matrix is stored sparsely, an equivalent mean of gene-level z-scores was used without control gene bins, so its coefficients are not strictly on the same scale as those of the other three datasets. For each donor with at least 100 cells, cell-level dyscoordination was correlated with each program score by Spearman’s correlation. Values were summarized into median across donors and tested against zero by a Wilcoxon signed-rank test.

To ask whether tumor grade and post-therapy shifts act within subsets rather than through subset composition changes, the same donor-by-subset medians were used. For glioma, each subset’s median was correlated across patients with ordered tumor grade by Spearman’s correlation and controlled by Benjamini-Hochberg procedure across subsets. For treated melanoma, each patient’s baseline value was compared to the mean of that patient’s later timepoints within the same subset, and the per-subset medians of these paired differences were reported.

### Aging rat kidney dyscoordination analysis

We extracted DEFND-seq and 10X scMultiome data for 22 kidney samples from 18 rats aged 16, 30, 56, and 82 weeks. Standard preprocessing through `Seurat` was performed to annotate 24 cell types across all 111,008 cells. For quality control, we removed cells with fewer than 250 detected genes or fewer than 2500 total UMI counts, as well as genes expressed in fewer than 5 cells. All analyses were conducted over the subset of 26,442 post-QC PT cells assigned to segments S1, S2, and S3. Transcriptional dyscoordination and clustering-based variance estimates for comparison were computed using the procedure described in the previous section.

EpiTrace is a method that estimates mitotic age of single cells by measuring chromatin accessibility in a set of reference “clock-like” genomic loci, as these loci are characterized by decreased heterogeneity in accessibility during cellular aging^70^. For the 4 Multiome libraries (12,075 PT cells), we used the paired ATAC data and selected the “iterative” option to compute mitotic age score. This option iteratively searches for additional loci showing best correlations between their accessibility and estimated mitotic age, starting from the initial reference set of clock-like differentially methylated loci. Since the original method used reference loci from human, we performed liftover from human genome (hg18) to rat genome (rn7) using the `easylift` R package (version 1.6.0)^145^ before applying the method.

To search for pathways enriched in genes with gains in transcriptional dyscoordination and with activation in high-dyscoordination cells, we performed gene set enrichment analysis with the `fgsea` R package (version 1.34.2)^146^. Pathway databases used include KEGG^147^, Gene Ontology^148^, and MSigDB Canonical Pathways^149^. For pathways related to genes with gains in transcriptional dyscoordination, we ranked all genes by their Spearman’s *ρ* values across age groups. For pathways related to genes with activation in high-dyscoordination cells, we ranked all genes by the correlation values between their log-normalized expressions and cell-level dyscoordination across all cells. In both cases, enrichment p-values were adjusted for multiple testing using the Benjamini-Hochberg procedure. Pathways with an FDR-adjusted p-value ≤ 0.05 were considered significant.

### Aging human kidney dyscoordination analysis

We analyzed a 10X snMultiome human kidney atlas, comprising 134,458 nuclei from 32 histologically normal donors (30-79 years old). For quality control, we removed nuclei with fewer than 250 detected genes or fewer than 2500 total UMI counts, as well as genes expressed in less than 0.1% of cells. Donor ages were grouped into 30-39, 40-49, 50-59, and 60+ bins. Analyses were restricted to the 3 tubular epithelial populations previously reported to carry significant somatic chromosomal alteration burden in non-injured kidneys. Transcriptional dyscoordination and pathways related to genes with gains in transcriptional dyscoordination and activation in high-dyscoordination cells were computed following the same procedure as for the aging rat kidney.

Somatic mosaic chromosomal alterations (mCAs) were inferred from the snATAC-seq using CHASM, which detects the rare, non-clonal copy number alterations (CNAs) found in normal tissue by estimating a cell-matched “no-CNA” baseline directly from the data rather than from a designated normal population^108^. To compare transcriptional dyscoordination with CHASM-derived mCA burden, we extracted paired RNA and ATAC profiles from 43,018 nuclei across the 3 cell types and aggregated to the donor level, where cell-level dyscoordination donors in the top and bottom 10% quantiles were removed and donors with at least 50 paired nuclei in each cell type were retained. Pearson correlation was computed between median cell-level dyscoordination and mean percentage of genome altered at the donor level.

### Aging mouse liver dyscoordination analysis

We collected 10X scMultiome data from 6 young-mouse liver samples and 10 old-mouse liver samples, with equal representation of males and females within each age group. Standard preprocessing through Seurat was performed for cell type annotation, where all cells annotated as hepatocytes were retained for downstream analysis. For quality control, we first identified and removed all doublets using the `DoubletFinder` R package (version 2.0.4)^150^. Then, we applied the same procedure as before by removing cells with fewer than 250 detected genes or fewer than 2500 total UMI counts, as well as genes expressed in fewer than 5 cells, leaving 66,816 hepatocytes with adequate transcriptome coverage.

Transcriptional dyscoordination, clustering-based variance estimates for comparison, and pathways related to genes with gains in transcriptional dyscoordination and activation in high-dyscoordination cells were all computed following the same procedure as for the aging rat kidney.

Zonation scores were calculated for all cells as the ratio of total pericentral marker expression to the combined expression of pericentral and periportal markers. Cells with strict 0 or 1 zonation scores are removed, and the rest are divided into 10 equal-sized bins, where we considered 0-20% bins to be periportal, 40-60% bins to be midlobular, and 80-100% bins to be pericentral. To find zone-specific genes with gain in transcriptional dyscoordination, we computed gene-level dyscoordination for all genes expressed by young and old cells within each zone separately and retained all genes with positive log-fold changes between the age groups. To find zone-specific genes with activation in high-dyscoordination cells, we extracted differentially expressed genes between low (bottom 20%) and high (top 20%) dyscoordination cells within each zone. Identified genes were then searched for their enriched pathways using the `enrichR` R package (version 3.2)^151^. KEGG^147^ and Gene Ontology^148^ were used as databases. Similarly, enrichment p-values were adjusted for multiple testing using the Benjamini-Hochberg procedure. Pathways with an FDR-adjusted p-value ≤ 0.05 were considered significant.

## Supplementary

**Supplementary Note 1-3 and Supplementary Figure 1-125 are kept together in the Transcriptional_dyscoordination_supplementary_material.docx.**

**Supplementary Table 1 List of single-cell RNA datasets used in the allele-specific expression analysis.**

Key information of the datasets used, including source, biological system, cross, and cell and gene counts, are recorded.

**Supplementary Table 2 List of T cell subset group assignments and program genes.**

Sheet 1 includes group assignments for all subsets across the four T cell datasets. Sheet 2 includes genes used to score each of the six T cell-associated transcriptional programs.

**Supplementary Table 3 List of PT segment identity markers.**

All 29 PT segment identity markers are recorded.

**Supplementary Table 4 Genes with significant dyscoordination changes across ages and their enriched pathways in rat PT cells.**

Sheet 1 includes all genes ranked by their Spearman’s ρ values across age groups of rat PT cells. Sheet 2 includes significantly enriched pathways identified with these genes.

**Supplementary Table 5 Genes significantly correlated with cell-level dyscoordination and their enriched pathways in rat PT cells.**

Sheet 1 includes all genes significantly correlated with cell-level dyscoordination of rat PT cells. Sheet 2 includes significantly enriched pathways identified with these genes.

**Supplementary Table 6 Genes with significant dyscoordination changes across ages and their enriched pathways in human tubular epithelial cells.**

Sheet 1 includes all genes ranked by their Spearman’s ρ values across age groups of human tubular epithelial cells. Sheet 2 includes significantly enriched pathways identified with these genes.

**Supplementary Table 7 Genes significantly correlated with cell-level dyscoordination and their enriched pathways in human tubular epithelial cells.**

Sheet 1 includes all genes significantly correlated with cell-level dyscoordination of human tubular epithelial cells. Sheet 2 includes significantly enriched pathways identified with these genes.

**Supplementary Table 8 Hepatocyte zonation markers and senescence and pro-inflammation markers.**

Sheet 1 records all 21 hepatocyte zonation markers (9 periportal, 4 midlobular, 8 pericentral). Sheet 2 records all curated 138 senescence and pro-inflammation markers.

**Supplementary Table 9 Genes significantly correlated with cell-level dyscoordination and their enriched pathways in hepatocytes.**

Sheet 1 includes all genes significantly correlated with cell-level dyscoordination of hepatocytes. Sheet 2 includes significantly enriched pathways identified with these genes.

**Supplementary Table 10 Genes with significant dyscoordination changes across ages and their enriched pathways in hepatocytes.**

Sheet 1 includes all genes ranked by their log-fold changes in gene-level dyscoordination across old vs. young hepatocytes. Sheet 2 includes significantly enriched pathways identified with these genes.

**Supplementary Table 11 Genes with significant zone-specific increases in dyscoordination across ages and their enriched pathways in hepatocytes.**

Sheet 1 includes all genes with positive log-fold changes in gene-level dyscoordination across old vs. young hepatocytes in each zone. Sheet 2 includes significantly enriched pathways identified with these genes by zone.

**Supplementary Table 12 Genes significantly correlated with zone-specific cell-level dyscoordination and their enriched pathways in hepatocytes.**

Sheet 1 includes all genes significantly positively correlated with cell-level dyscoordination of hepatocytes in each zone. Sheet 2 includes significantly enriched pathways identified with these genes by zone.

### Declaration of generative AI and AI-assisted technologies in the manuscript preparation process

During the preparation of this work the authors used ChatGPT 5 and 5.1 in order to refine the writing of the article. After using this tool/service, the authors reviewed and edited the content as needed and took full responsibility for the content of the published article.

