## Supplementary Notes and Figures for "A Measure of Transcriptional Dyscoordination for Quantifying Aging in Single Cells"

**Supplementary Note 1 In-silico spike-ins**

Analyses in the main text rely on real datasets, in which the true amount of transcriptional disorder is unknown. To test the measure against a known ground truth, we used an in-silico spike-in applied directly to real single-cell counts from rat kidney PT cells (n = 8,000), mouse liver hepatocytes (n = 7,999), and mouse bone marrow HPCs (n = 2,144), which retains the noise structure of genuine data while fully controlling the perturbation. For rat kidney and mouse liver substrates, we restricted to the cells in the young animals, whose cells have not yet accumulated substantial age-related dyscoordination and therefore provide a clean null baseline against which the injected perturbations are measured. We first constructed a baseline true expression profile for every cell by smoothing the observed expression over its 20 nearest neighbors in the expression space, which served as the ground truth onto which perturbations are added. This baseline was built with cell-cell similarity deliberately and differed from the gene-gene regression the measure itself uses, so that the truth is defined in one way and measured in another to prevent circularity. Within each substrate (dataset), cells were randomly divided into 4 arms sharing the same library sizes and gene panel (the 5,000 most detected genes with detection fraction ≥ 0.1): an unperturbed control (40%), a coordinated arm carrying a perfectly coordinated (rank one) program across 150 spike-in genes, in which a single per-cell magnitude shifts all program genes together (20%), a dyscoordinated arm receiving mean-preserving noise applied independently to each of the same 150 genes above, which increases their variability without changing the mean expression (20%), and a combined arm carrying both perturbations (20%). These perturbations were applied to the latent baseline expression, and counts were regenerated with multinomial resampling at each cell's original depth to match the arms on sequencing depth by construction. Within each biological replicate, transcriptional dyscoordination was then computed with a single manifold fit for all arms, since a separate fit per arm would allow each arm to re-learn its own perturbation. Each arm was summarized by the ratio of its mean dyscoordination to that of the control and evaluated across 10 replicates per substrate.

Across substrates, transcriptional dyscoordination responded specifically to genuinely uncoordinated variation (Supplementary Figure 1). The coordinated program rendered all 150 spike-in genes differentially expressed and shifted mean expression on the spike-in genes roughly 2.7-fold, yet left dyscoordination essentially at the control level (dyscoordination ratio: 1.01-1.02), indicating that the manifold absorbed this strong cell state shift into its predictable component rather than scoring it as disorder. The dyscoordinated arm did the reverse, raising dyscoordination substantially (1.06-1.16) while rendering far fewer genes differentially expressed, and the combined arm recovered both signals (1.20-1.32). It is worth noting that sensitivity diverged on mouse bone marrow

HPCs, where cells bearing mean-preserving noise only exhibited a dyscoordination ratio of 1.06 and were significant in 3 of 10 replicates. This is expected, as its manifold is learned from roughly only a quarter as many observations as the other two populations, making the control expression noisier and the ratio more compressed.

The manifold can absorb a coordinated program only if it can properly learn that program from the data. We therefore tested the measure's learnability by varying how many cells carry the program and how many genes it spans and concluded that absorption primarily depends on the former (Supplementary Figure 2). Specifically, when the program was present in fewer than 10% of cells, there were too few cells for the regression to learn, and it leaked into the residual as a small false-positive dyscoordination signal, while at 20% of cells or more it was learned and fully absorbed. In comparison, widening the coordinated program from 30 to 600 genes produced no such breakdown at any width, while the dyscoordinated arm showed the expected rise in dyscoordination ratio as more genes were perturbed. Rarity across cells therefore defeats manifold learning, whereas narrowness across genes does not. Note that within the 5,000 gene panel, dyscoordination is rather depth-dependent, so every interpretation made here is on the depth-balanced arms. Together, these simulations confirm that transcriptional dyscoordination reports genuine uncoordinated variability of single cells rather than coordinated differential expression.

**Supplementary Note 2 Technical robustness of transcriptional dyscoordination.**

As transcriptional dyscoordination is derived from single-cell count data, its magnitude could in principle reflect technical properties of the assay, including sequencing depth, cell cycle activity, and batch structure, or to the number of cells available to estimate each stratum, rather than genuine differences in intrinsic transcriptional variability. In this note, we evaluated each of these possibilities holistically across all datasets primarily analyzed in this study.

**Sequencing depth and cell count**

As with any measure derived from single-cell counts, cell-level dyscoordination is unavoidably dependent on sequencing depth (the total UMI count per cell) to some extent. To confirm that the observed age- and condition-associated dyscoordination shifts in the main text are not attributable to depth, we fitted a single global LOESS of log-transformed cell-level dyscoordination against sequencing depth, removed the depth-predicted component for each cell, and asked whether the biological signal survived in the residuals. This is a global, depth-only correction rather than a cell type-specific or multi-covariate fit, because the latter also removes genuine biological structure between cell types or states. After this adjustment, all observed trends were essentially unchanged across all 8 datasets primarily analyzed in the study (mouse bone marrow cells, 4 T cell datasets, rat kidney PT cells, human kidney tubular epithelial cells, and mouse liver hepatocytes; Supplementary Figure 15, 50-53, 84, 98, 108). Specifically, the reported age-associated increases in cell-level dyscoordination remained significant, indicating that they reflect true biology rather than depth-confounded artifact.

Another concern is that strata containing more cells could be estimated more precisely and therefore yield systematically different values. We addressed this in rat kidney PT cells and mouse liver hepatocytes by running a subsampling experiment and reading it from two perspectives. Within each age group, we recomputed its dyscoordination from random subsamples of increasing size from 100 to 5,000 cells, summarizing each subsample by the median cell-level dyscoordination across 5 independent draws. First, reading along subsample size for an age group, the dyscoordination estimate did not decline as more cells were added. Instead, it plateaued above a few hundred cells, indicating that a group's cell count does not systematically bias its dyscoordination downward. Second, reading across age groups at a matched subsample size, the age-associated dyscoordination increase was fully preserved in both datasets. These analyses demonstrate that the reported aging trend in transcriptional dyscoordination is not an artifact of differing cell counts or estimation precision but a genuine biological signal (Supplementary Figure 85, 109).

**Biological batches and replicates**

To understand whether the age- and condition-associated gradients were driven by individual samples, we examined the distribution of cell-level dyscoordination separately for every biological replicate (e.g., donor, patient, animal, or sequencing library) in each of the 9 datasets analyzed above. Within the same age or condition group, sample-wise distributions were mutually consistent, and no single sample accounted for the group-level trend alone (Supplementary Figure 16, 54-57, 86, 99, 110). The observed gradients are therefore inherent properties of the biological groups, not of batch structure. One exception should be noted. In the aging human T cell cohort of Wang et al. 2025, 8 donors of the 30-40-year-old group (sample 29 to 36) were profiled as a block and share a uniform offset of roughly 2 to 3 logarithmic units above donors in every other age band across all subsets (Supplementary Figure 57). As donor age and processing order are confounded in that dataset, its age-level axis is not interpreted.

**Cell-cycle effect**

We assigned each cell an S phase and G2/M score using the `CellCycleScoring` function from `Seurat` and computed the Spearman's correlation between each score and cell-level dyscoordination within every cell type of the analyzed datasets. Correlations were weak in magnitude and inconsistent in sign across most cell types, with modest positive ones only in a selection of highly proliferative, effector-like populations, such as terminal effector CD8+ T cells in glioma (Supplementary Figure 17, 18, 58-65, 87, 100, 101, 111). As a result, cell cycle activity is not a uniform driver of transcriptional dyscoordination, which is expected by the measure's construction: as cell cycle variation is usually a coordinated transcriptional program shared across cells, it is absorbed into the structured component of the expression (manifold) and thus largely excluded from the residual that defines dyscoordination.

**Summary**

Across every dataset examined, the reported age- and condition-associated dyscoordination patterns remained stable after the sequencing-depth adjustment and were not driven by cell count per stratum, individual biological replicates, or cell-cycle state in the analyses above. These results support interpreting the reported biological differences as reproducible variation not explained by these measured technical factors.

**Supplementary Note 3 Conventional age-associated changes in cellular composition and mean expression**

To distinguish transcriptional dyscoordination from conventional forms of transcriptomic aging, we separately analyzed age-associated changes in cell type composition and within-cell type mean gene expression across the principal aging datasets. These analyses were performed independently of the dyscoordination framework, using the biological sample as the unit of replication. They were used to address two complementary questions: composition analysis investigates whether the abundance of annotated populations changes with age, whereas differential expression analysis investigates whether the mean transcriptional state changes within a defined population.

Across the solid-tissue datasets, we observed extensive within-cell type mean expression remodeling even in settings where the available sample sizes provided limited evidence for cell type abundance changes, and rat kidney was a particularly clear example. In the blood cohorts, age-associated compositional remodeling was more prominent, including marked depletion of naive CD8 T cell compartments. Null composition tests are interpreted relative to the resolution available at the biological-sample level rather than as evidence that no change occurs.

These conventional analyses reinforce the three-part framework used throughout the manuscript: aging can alter which cells are present, coordinately reprogram the mean state of cells within a population, and reduce the precision with which a predictable transcriptional state is executed. The first two effects are appropriately studied with composition and differential-expression analyses, while transcriptional dyscoordination is designed specifically to quantify the third. Accordingly, a flat dyscoordination trajectory is not interpreted as absence of aging, and activation of inflammatory, SASP, or stress-response genes in high-dyscoordination cells is treated as coordinated biology associated with a dyscoordinated state rather than as stochasticity of those programs themselves.

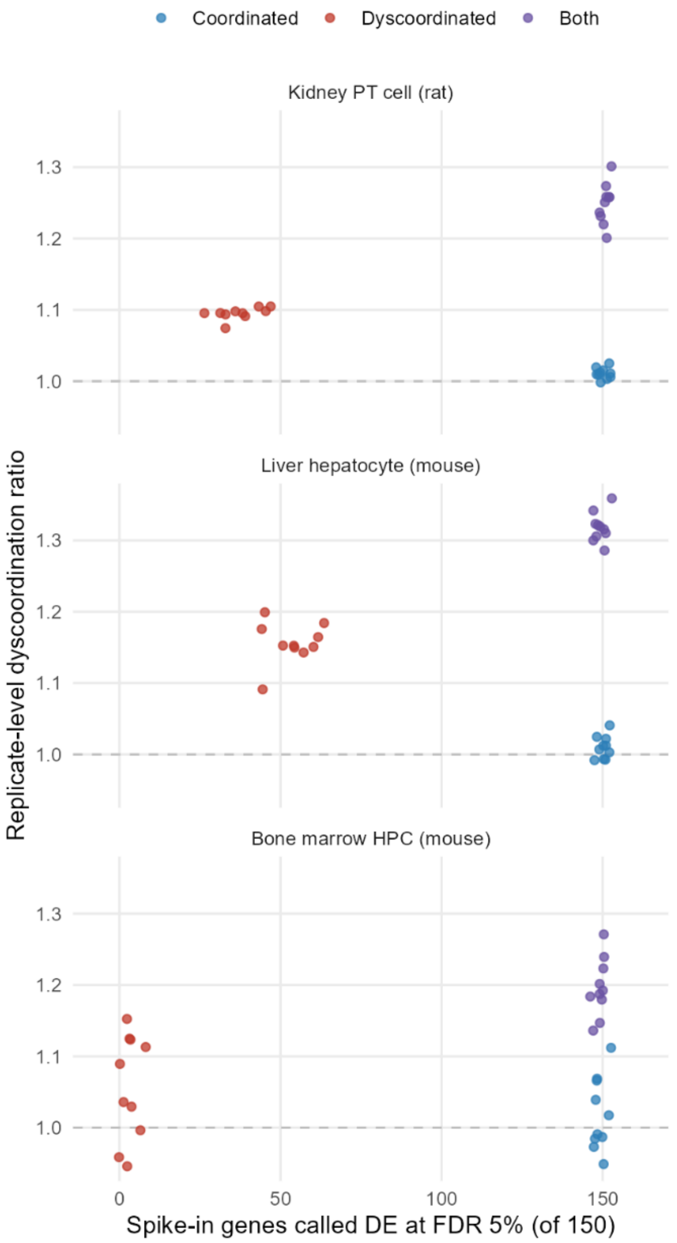


**Supplementary Figure 1 Replicate-level dyscoordination ratio and spike-in DE gene count across substrates.**

Dot plots show the mean spike-in DE gene count (x-axis) and dyscoordination ratio (y-axis) across 10 replicates in 3 substrates: rat kidney PT cells (top), mouse liver hepatocytes (middle), and mouse bone marrow HPCs (bottom). Horizontal and vertical bars show the 95% confidence intervals across replicates for the two quantities, and the dashed lines mark the control baseline (ratio = 1). Each plotted point represents one replicate within a perturbation arm; there are 10 replicates per perturbation arm (30 non-control perturbation replicates per substrate).

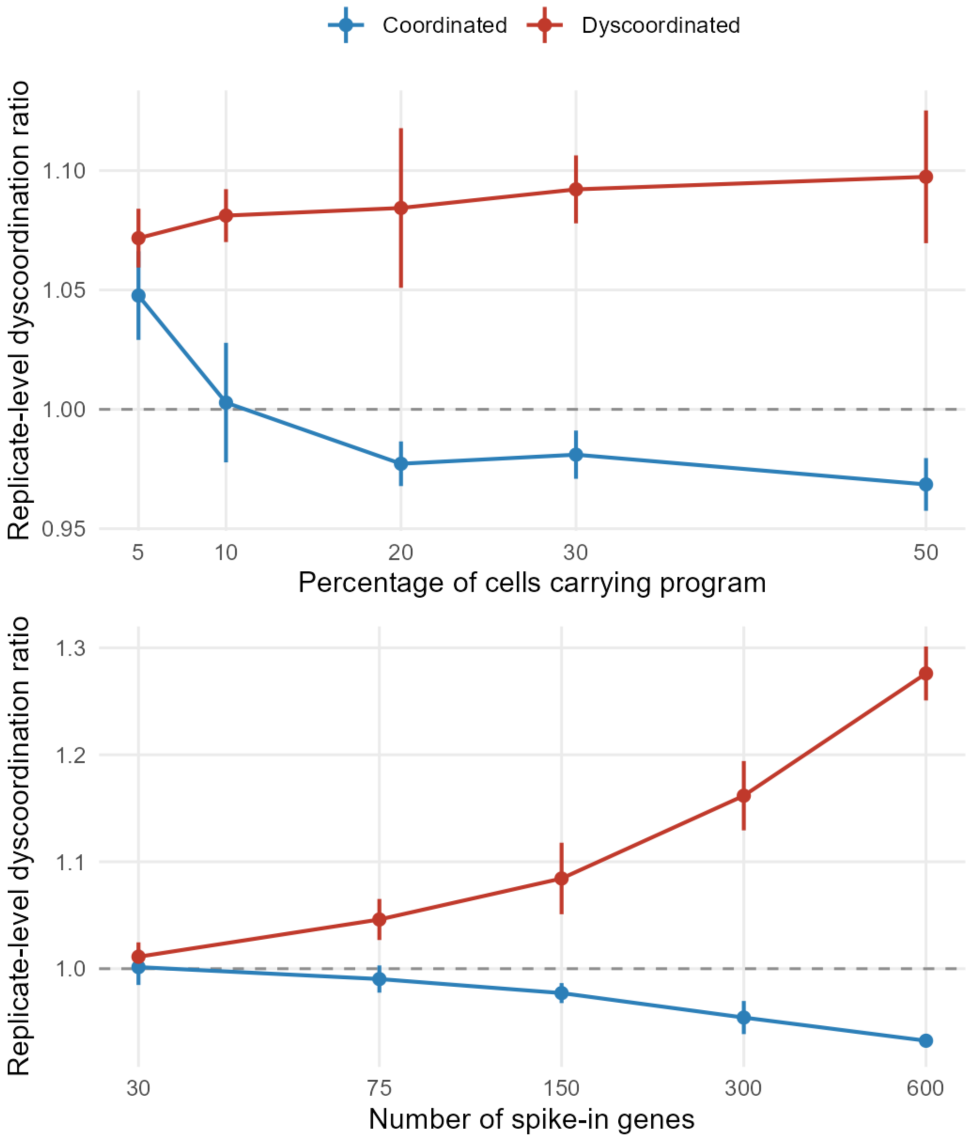


**Supplementary Figure 2 Correlation between dyscoordination ratio and percentage of cells carrying program and number of spike-in genes.**

Dot plots show the replicate-level dyscoordination ratio across different percentage of cells carrying program (top) and number of spike-in genes (bottom). Vertical bars show the 95% confidence intervals of each ratio, and color indicates coordinated (blue) or dyscoordinated (red) arm.

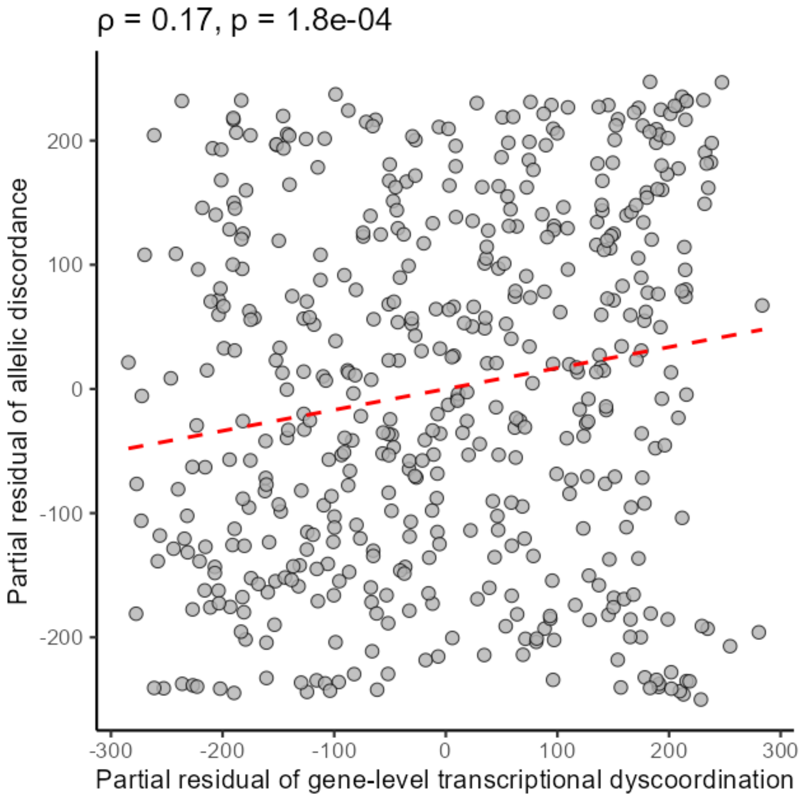


**Supplementary Figure 3 Correlation between partial residuals of gene-level dyscoordination and allelic discordance in 129 x CAST/EiJ mice (Ochiai et al. 2020).**

Scatter plot shows the relationship between partial residuals of log-transformed gene-level dyscoordination and allelic discordance across top 500 highly variable genes detected on both alleles in 129 x CAST/EiJ mice. The red dotted line indicates linear regression fits. Spearman correlation coefficients and corresponding p-values are shown in the top left corner.

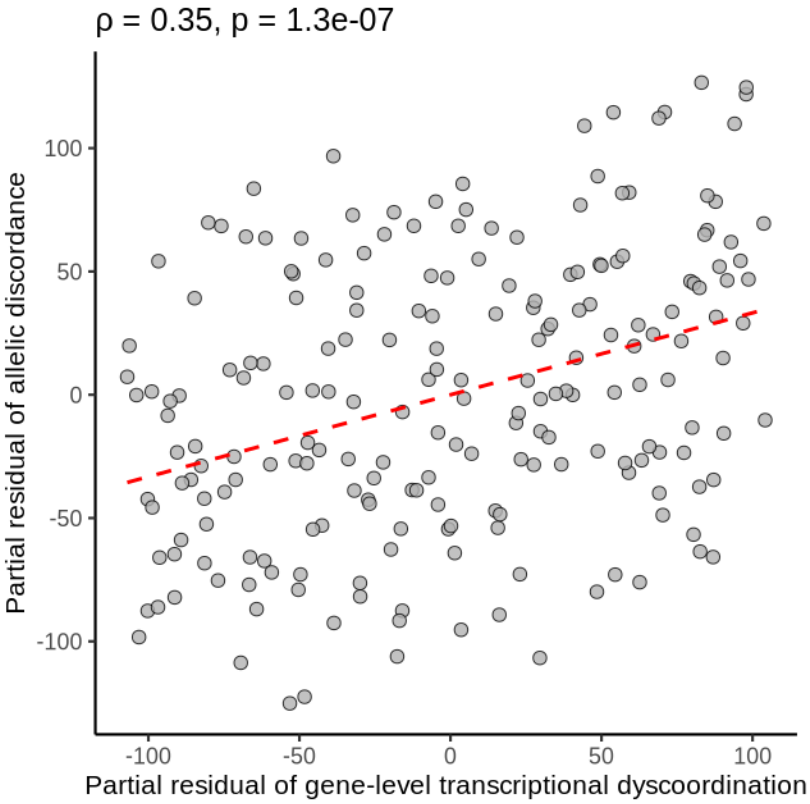


**Supplementary Figure 4 Correlation between partial residuals of gene-level dyscoordination and allelic discordance in SPRET/EiJ x C57BL/EiJ mice (van der Veeken et al. 2019).**

Scatter plot shows the relationship between partial residuals of log-transformed gene-level dyscoordination and allelic discordance across top 500 highly variable genes detected on both alleles in SPRET/EiJ x C57BL/EiJ mice. The red dotted line indicates linear regression fits. Spearman correlation coefficients and corresponding p-values are shown in the top left corner.

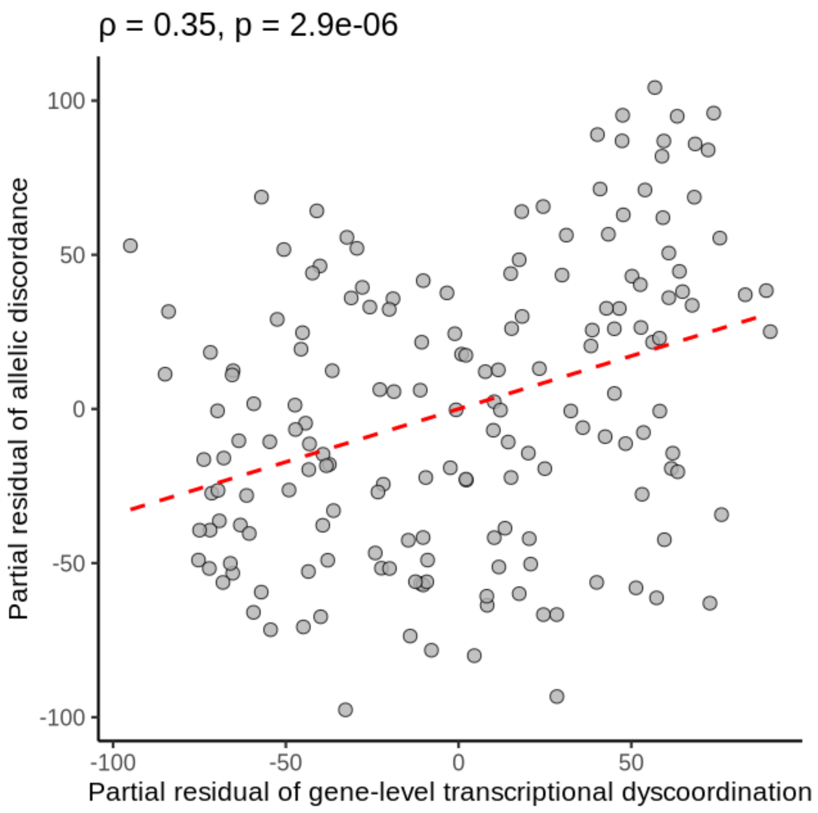


**Supplementary Figure 5 Correlation between partial residuals of gene-level dyscoordination and allelic discordance in pooled SPRET/EiJ x C57BL/EiJ mice (Pritykin et al. 2021).**

Scatter plot shows the relationship between partial residuals of log-transformed gene-level dyscoordination and allelic discordance across top 500 highly variable genes detected on both alleles in pooled SPRET/EiJ x C57BL/EiJ mice. The red dotted line indicates linear regression fits. Spearman correlation coefficients and corresponding p-values are shown in the top left corner.

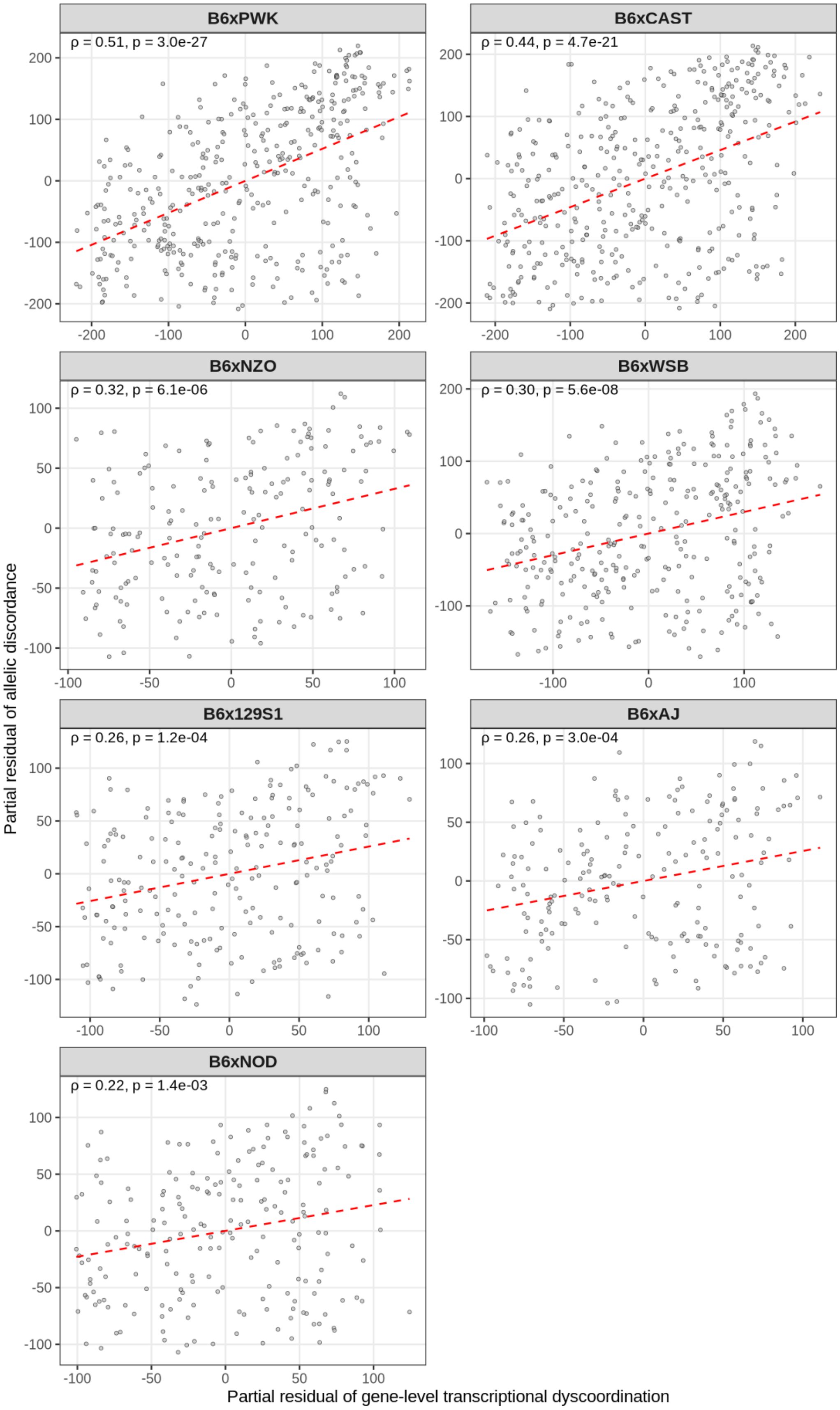
B6xPWK
ρ = 0.51, p = 3.0e-27
B6xCAST
ρ = 0.44, p = 4.7e-21
B6xNZO
ρ = 0.32, p = 6.1e-06
B6xWSB
ρ = 0.30, p = 5.6e-08
B6x129S1
ρ = 0.26, p = 1.2e-04
B6xAJ
ρ = 0.26, p = 3.0e-04
B6xNOD
ρ = 0.22, p = 1.4e-03
Partial residual of allelic discordance
Partial residual of gene-level transcriptional dyscoordination

**Supplementary Figure 6 Correlation between partial residuals of gene-level dyscoordination and allelic discordance in multi-cross F1 hybrid mice (Weber et al. 2026).**

Scatter plots show the relationship between partial residuals of log-transformed gene-level dyscoordination and allelic discordance across top 500 highly variable genes detected on both alleles in multi-cross F1 hybrid mice (Weber et al. 2026). Red dotted lines indicate linear regression fits. Spearman correlation coefficients and corresponding p-values are shown in the top left corner of each panel.

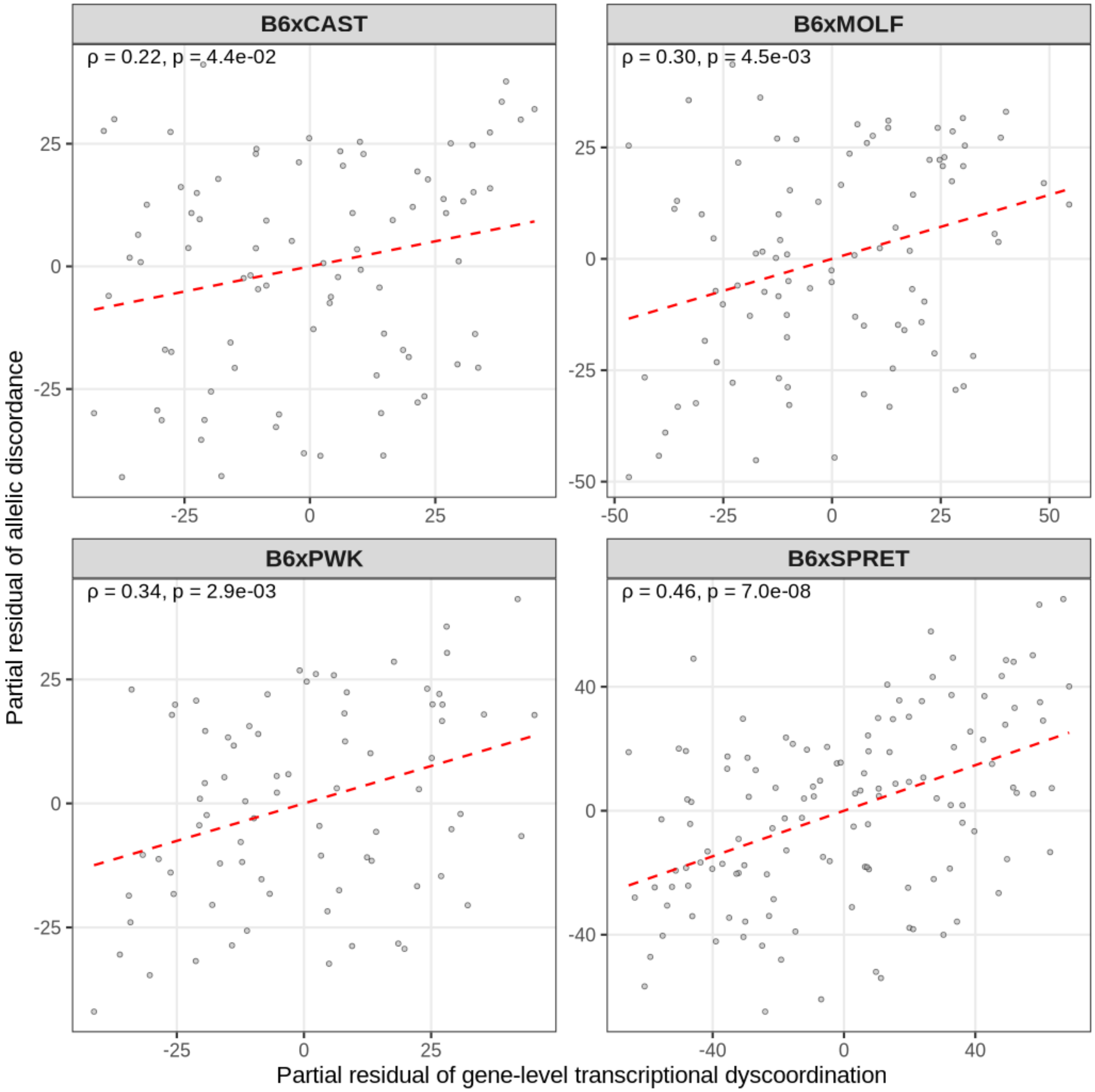


**Supplementary Figure 7 Correlation between partial residuals of gene-level dyscoordination and allelic discordance in multi-cross F1 hybrid mice (Medina-Cano et al. 2025).**

Scatter plots show the relationship between partial residuals of log-transformed gene-level dyscoordination and allelic discordance across top 500 highly variable genes detected on both alleles in multi-cross F1 hybrid mice (Medina-Cano et al. 2025). Red dotted lines indicate linear regression fits. Spearman correlation coefficients and corresponding p-values are shown in the top left corner of each panel.

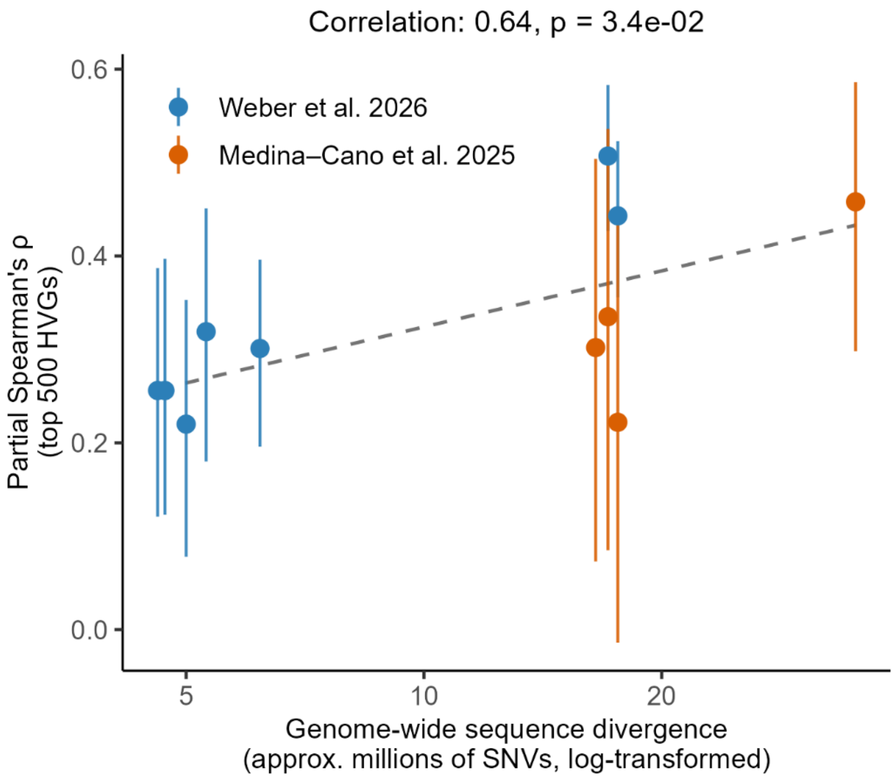


**Supplementary Figure 8 Correlation between partial Spearman's correlation and genome-wide sequence divergence.**

Scatter plot shows correlation between partial Spearman's correlation between gene-level dyscoordination and allelic discordance for top 500 HVGs and genome-wide sequence divergence for the two multi-cross F1 hybrid mice experiments. Black dotted line denotes the best-fit regression line. Spearman correlation coefficient and corresponding p-value are shown in the top left corner.

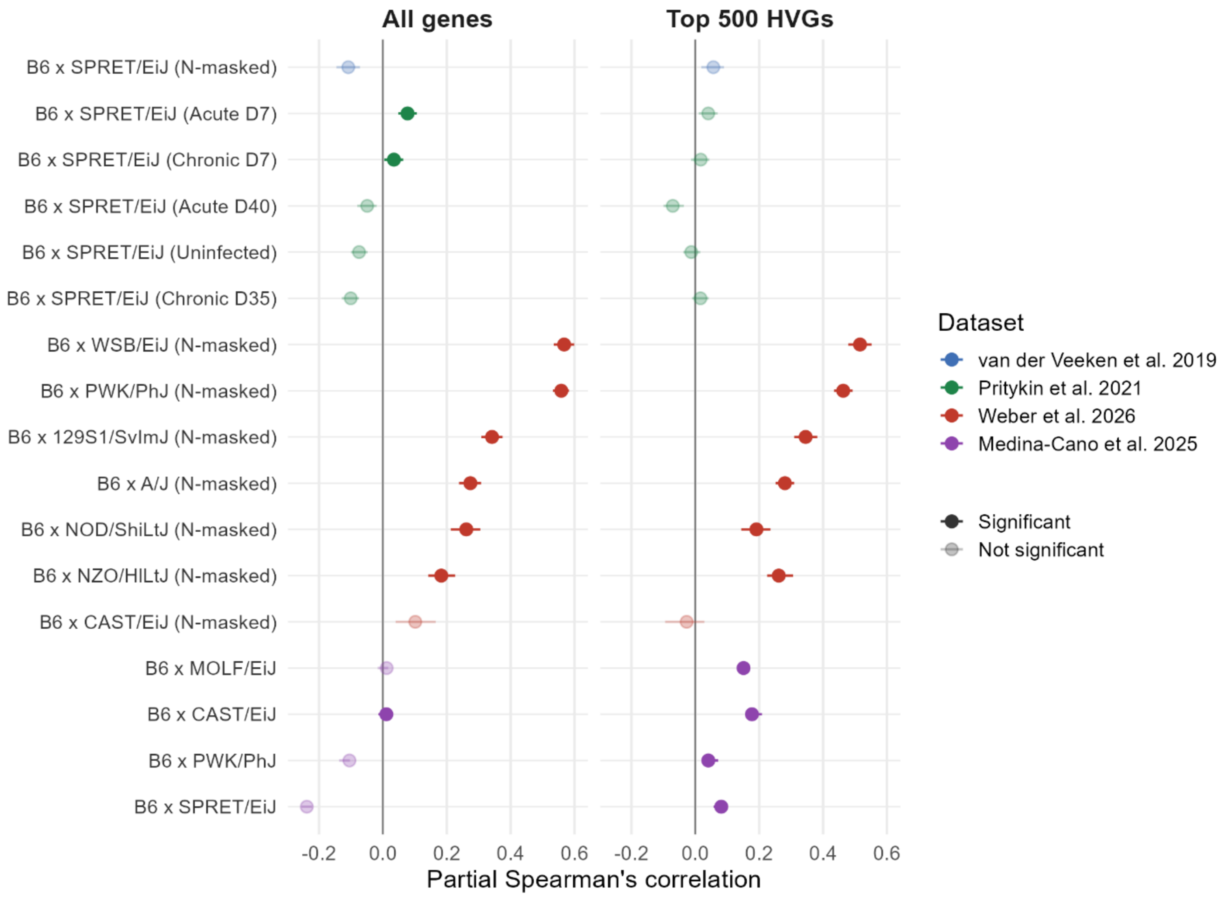


**Supplementary Figure 9 Correlation between partial residuals of cell-level dyscoordination and allelic discordance in F1 hybrid mice.**

Dot plots show the correlation between partial residuals of log-transformed cell-level dyscoordination and allelic discordance, computed after averaging each cell's score over its 15 nearest neighbors (pooling with k = 15), across all genes and top 500 HVGs in multi-cross F1 hybrid mice. Each row represents one cross (unit), with points indicating the correlation values and horizontal bars indicating the corresponding 95% confidence intervals. Color denotes dataset, and opaque points are significant under a permutation null ($p \leq 0.05$) while faded points are not. For crosses in van der Veeken et al. 2019 and Weber et al. 2026, allelic counts were obtained from N-masked realignment due to susceptibility to reference-mapping bias.

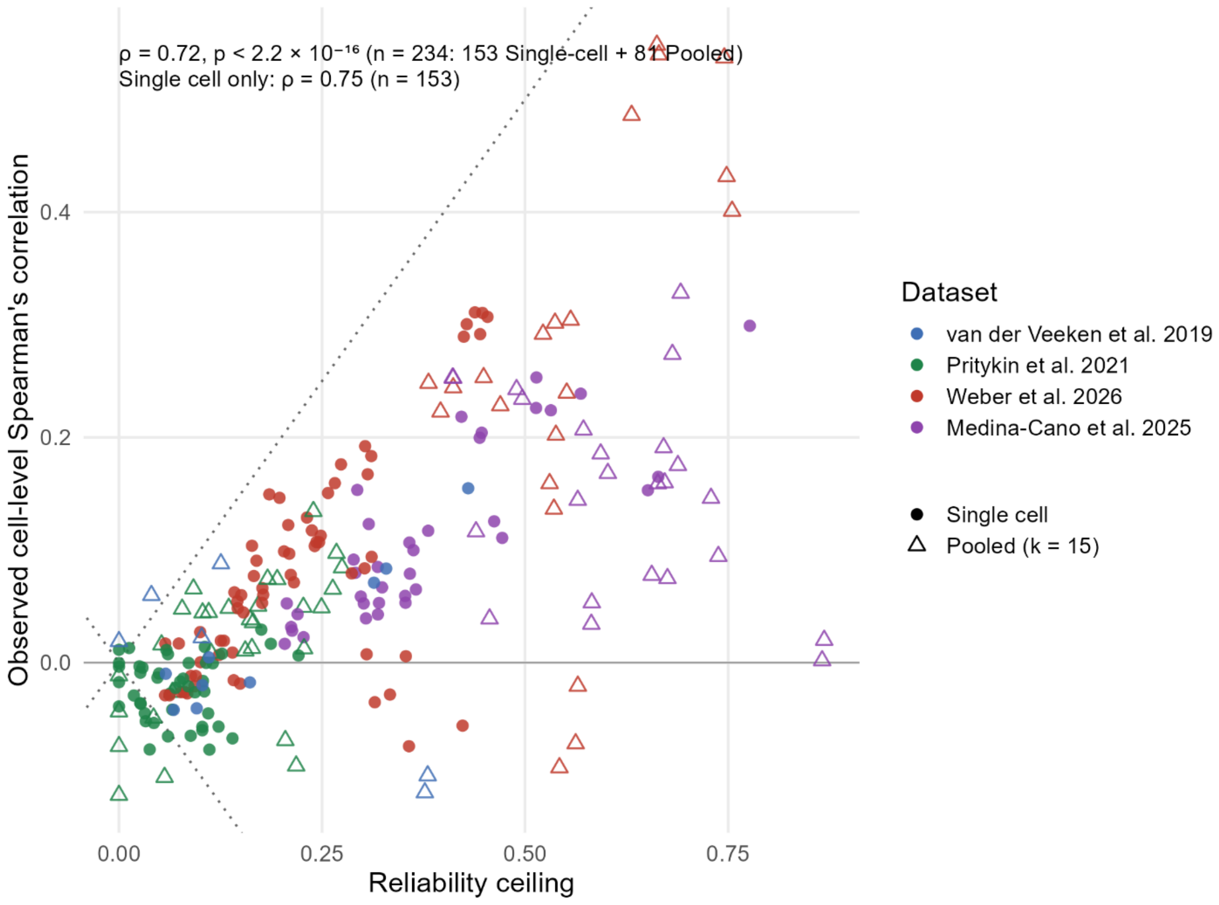


**Supplementary Figure 10 Correlation between reliability ceiling and observed cell-level Spearman's correlation.**

Scatter plot shows the correlation between reliability ceiling, or the geometric mean of the split-half reliabilities of the allelic discordance and transcriptional dyscoordination, and observed cell-level Spearman's correlation. Each point represents one analysis configuration as the combination of dataset, cross (unit), and gene panel evaluated at either single-cell resolution ($k = 1$, circular dots) or with neighborhood pooling ($k = 15$, triangular dots). Color denotes dataset, and the dotted lines mark the attenuation bounds, or the maximum correlation magnitude attainable under the shrinkage of true correlation. Spearman correlation coefficients and corresponding p-values for all configurations and single-cell ones are shown in the top left corner.

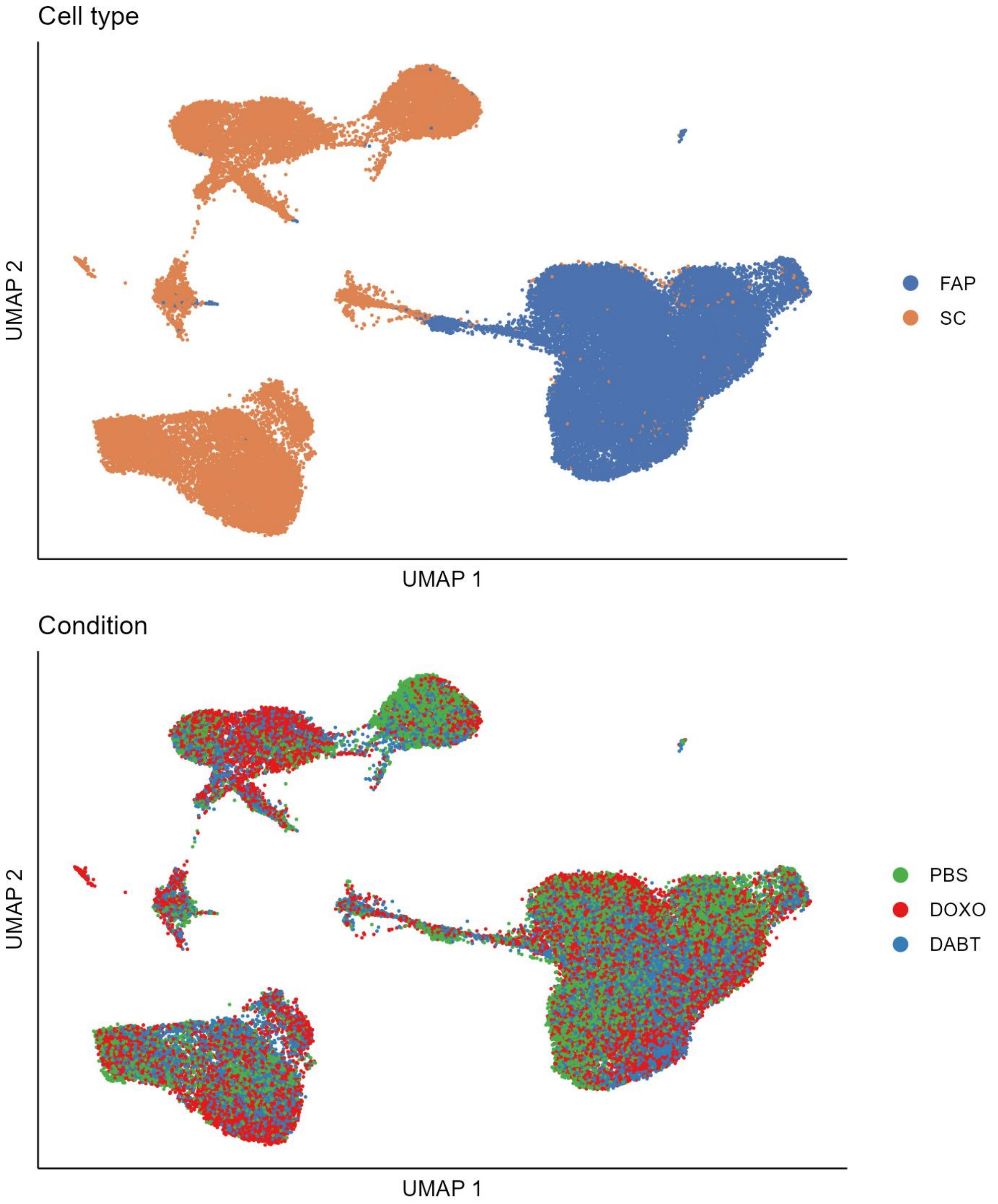


**Supplementary Figure 11 UMAP visualization of induced senescence in mouse muscle stem cells.**

Cells are embedded using the first two UMAP dimensions and colored by annotated cell type (top) and condition (bottom).

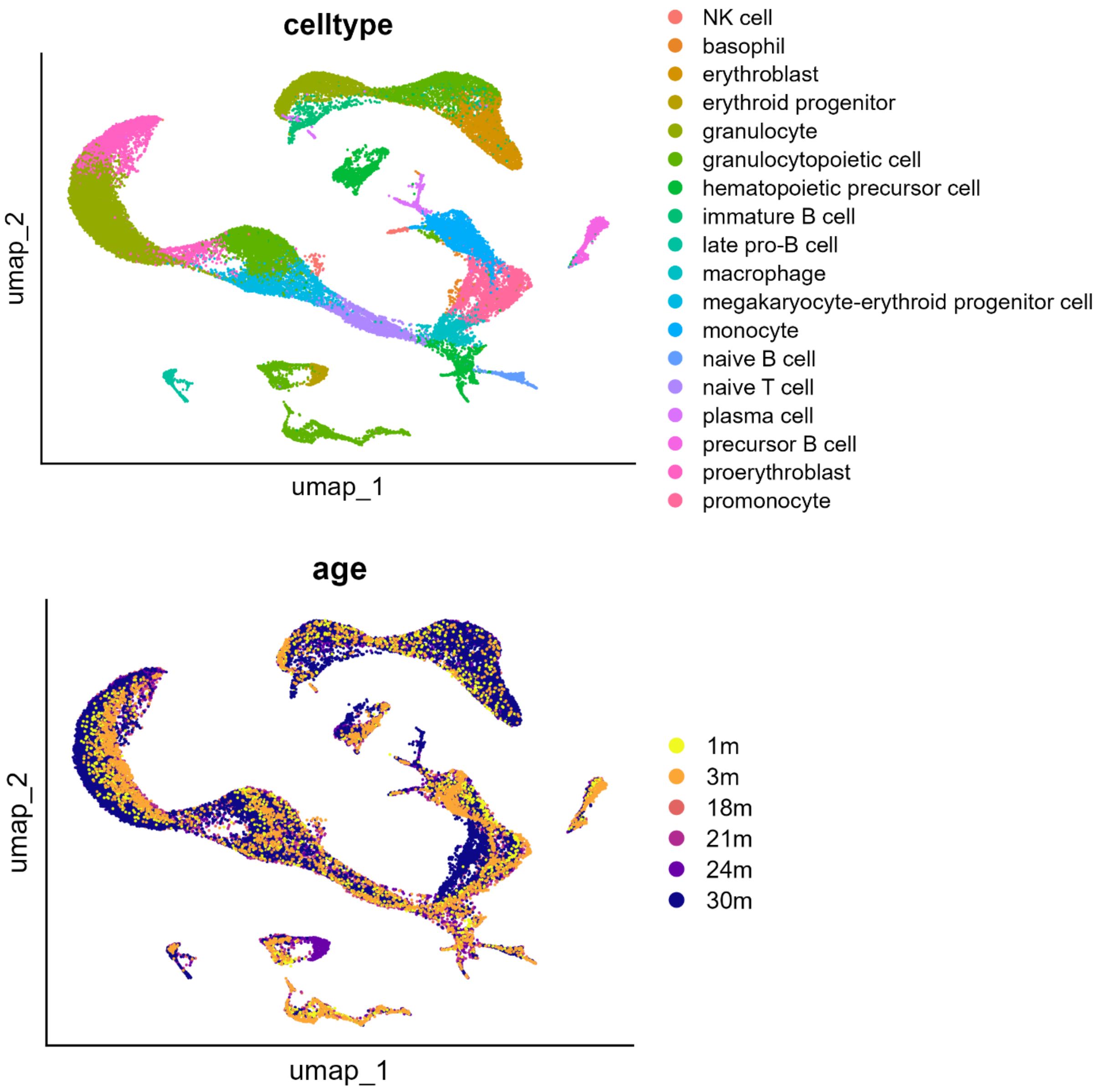


**Supplementary Figure 12 UMAP visualization of bone marrow cells from Tabula Muris Senis.**

Cells are embedded using the first two UMAP dimensions and colored by annotated cell type (top) and age group (bottom).

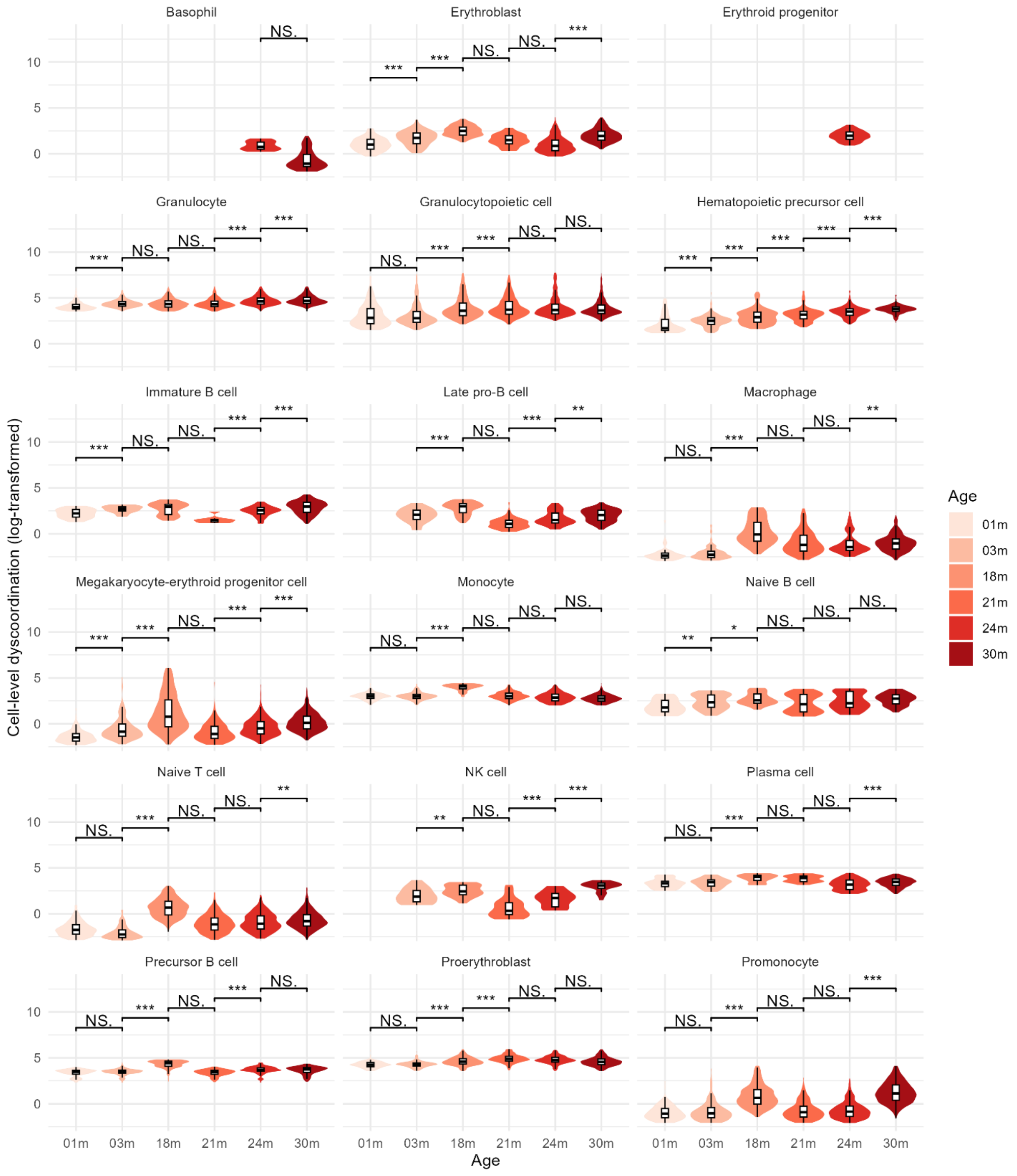
Basophil
Erythroblast
Erythroid progenitor
Granulocyte
Granulocytopoietic cell
Hematopoietic precursor cell
Immature B cell
Late pro-B cell
Macrophage
Megakaryocyte-erythroid progenitor cell
Monocyte
Naive B cell
Naive T cell
NK cell
Plasma cell
Precursor B cell
Proerythroblast
Promonocyte
Cell-level dyscoordination (log-transformed)
Age
01m
03m
18m
21m
24m
30m
NS.
***
**
*

**Supplementary Figure 13 Distribution of cell-level dyscoordination in mouse bone marrow cells.**

Violin and box plots show the distribution of log-transformed cell-level dyscoordination values separately for each cell type and age group. Values in the top and bottom 2.5% quantiles are considered outliers and are removed from the illustration. Pairwise comparisons between age groups were tested using one-sided Wilcoxon rank-sum test (alternative hypothesis: older age group has higher dyscoordination). Significance levels are annotated above each comparison (NS = not significant, * $p < 0.05$, ** $p < 0.01$, *** $p < 0.001$).

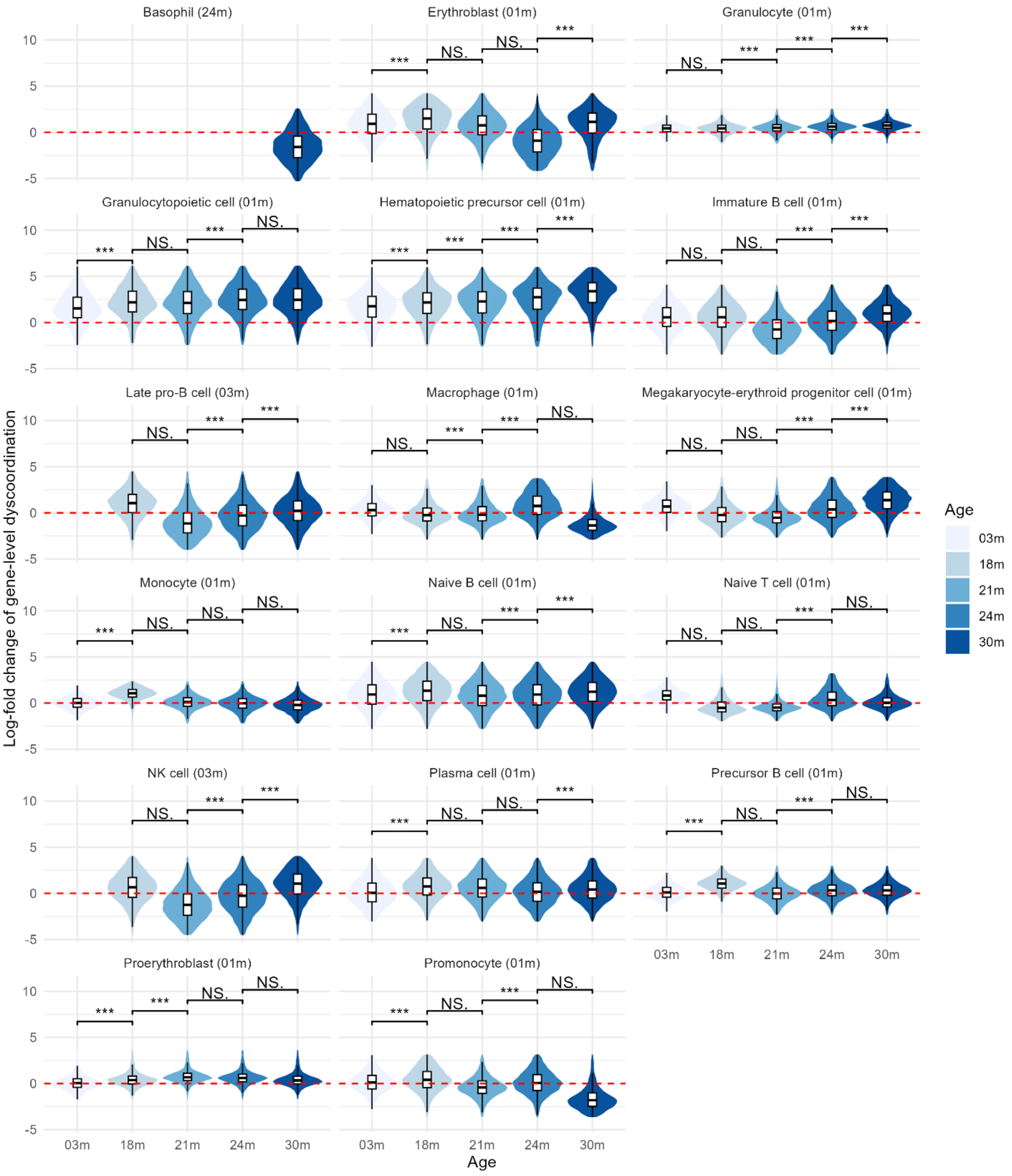

Basophil (24m)
Erythroblast (01m)
Granulocyte (01m)
Granulocytopoietic cell (01m)
Hematopoietic precursor cell (01m)
Immature B cell (01m)
Late pro-B cell (03m)
Macrophage (01m)
Megakaryocyte-erythroid progenitor cell (01m)
Monocyte (01m)
Naive B cell (01m)
Naive T cell (01m)
NK cell (03m)
Plasma cell (01m)
Precursor B cell (01m)
Proerythroblast (01m)
Promonocyte (01m)
NS.
***
10
5
0
-5
03m
18m
21m
24m
30m
Age
Log-fold change of gene-level dyscoordination
Age
03m
18m
21m
24m
30m

**Supplementary Figure 14 Distribution of gene-level dyscoordination in mouse bone marrow cells.**

Violin and box plots show the distribution of $\log_2$-fold changes in dyscoordination relative to the baseline age group. Values are displayed separately for each cell type and age group, with the baseline age group indicated in parentheses in each subplot title. Values in the top and bottom 2.5% quantiles are considered outliers and are removed from the illustration. Red dashed line denotes zero log-fold change. Pairwise comparisons between age groups were tested using one-sided Wilcoxon rank-sum test (alternative hypothesis: older age group has higher dyscoordination). Significance levels are annotated above each comparison (NS = not significant, * $p < 0.05$, ** $p < 0.01$, *** $p < 0.001$).

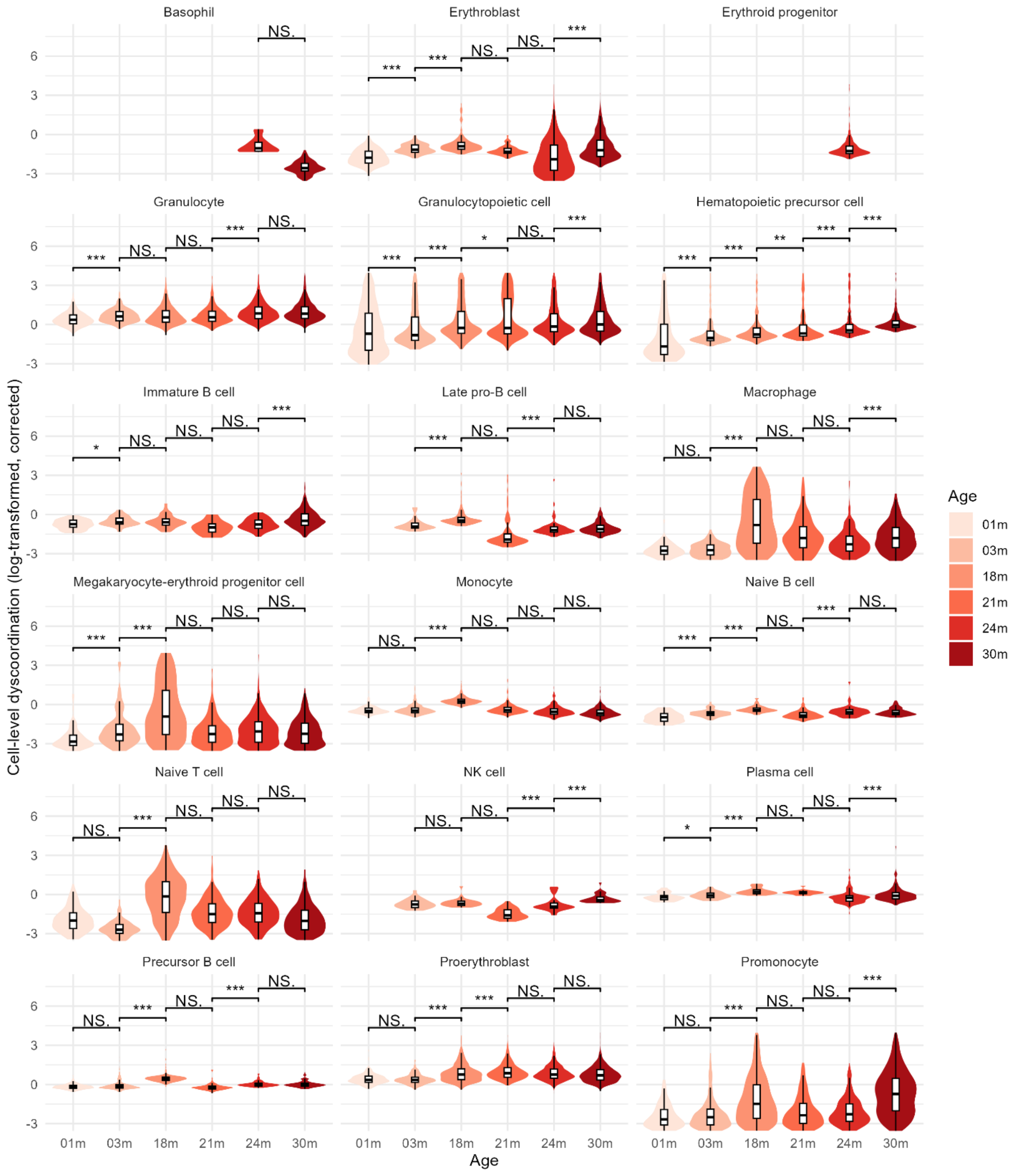

Basophil
Erythroblast
Erythroid progenitor
Granulocyte
Granulocytopoietic cell
Hematopoietic precursor cell
Immature B cell
Late pro-B cell
Macrophage
Megakaryocyte-erythroid progenitor cell
Monocyte
Naive B cell
Naive T cell
NK cell
Plasma cell
Precursor B cell
Proerythroblast
Promonocyte
Cell-level dyscoordination (log-transformed, corrected)
Age
01m
03m
18m
21m
24m
30m
NS.
***
**
*

**Supplementary Figure 15 Distribution of corrected cell-level dyscoordination in mouse bone marrow cells.**

Violin and box plots show the distribution of log-transformed and depth-corrected cell-level dyscoordination values separately for each cell type and age group. Values in the top and bottom 2.5% quantiles are considered outliers and are removed from the illustration. Pairwise comparisons between age groups were tested using one-sided Wilcoxon rank-sum test (alternative hypothesis: older age group has higher dyscoordination). Significance levels are annotated above each comparison (NS = not significant, * $p < 0.05$, ** $p < 0.01$, *** $p < 0.001$).

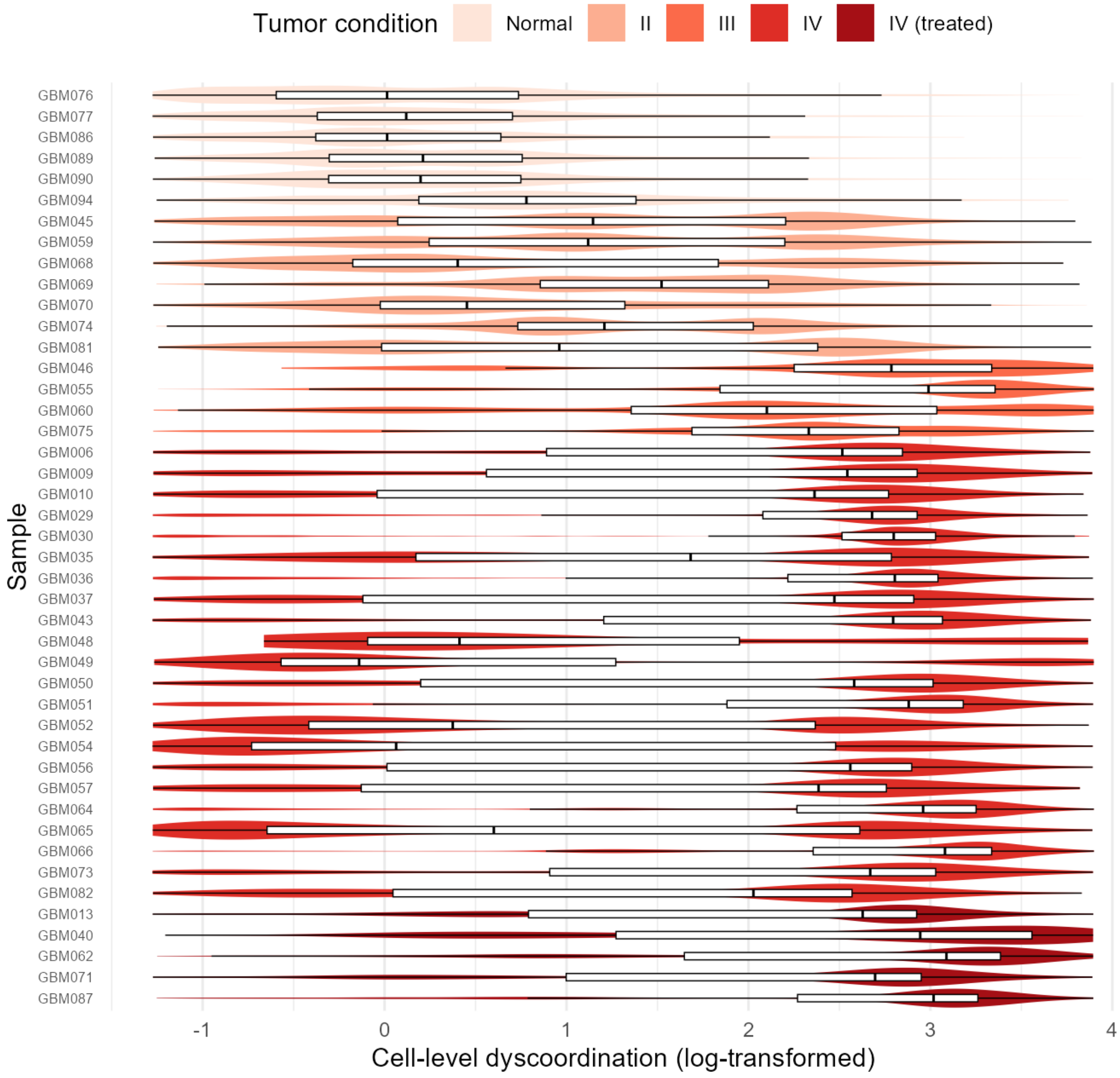


**Supplementary Figure 16 Distribution of cell-level dyscoordination across biological replicates in mouse bone marrow cells.**

Violin and box plots show the distribution of log-transformed cell-level dyscoordination for each individual sequencing library, ordered and colored by age group. Values in the top and bottom 2.5% quantiles are considered outliers and are removed from the illustration.

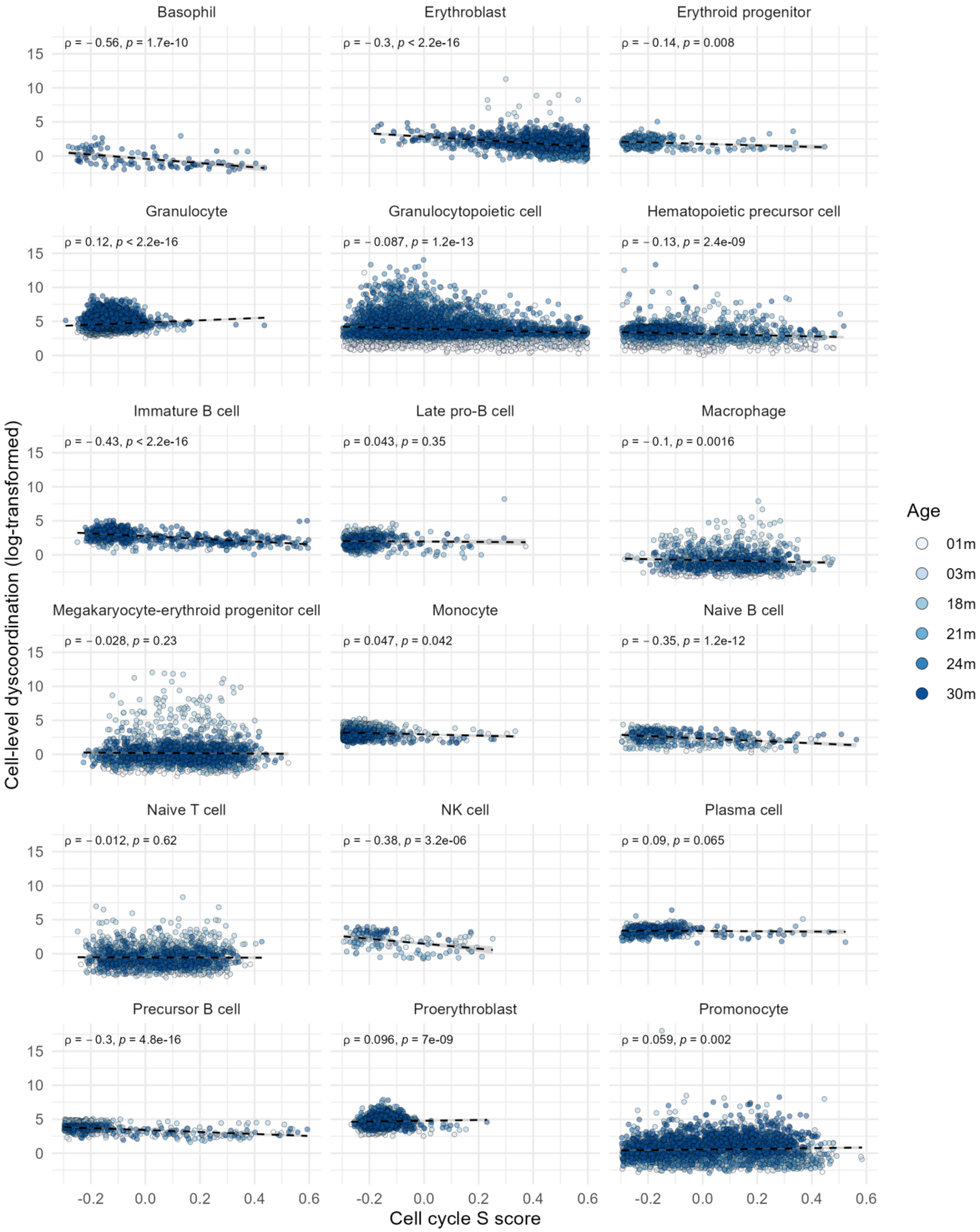

Basophil
ρ = − 0.56, p = 1.7e-10
Erythroblast
ρ = − 0.3, p < 2.2e-16
Erythroid progenitor
ρ = − 0.14, p = 0.008
Granulocyte
ρ = 0.12, p < 2.2e-16
Granulocytopoietic cell
ρ = − 0.087, p = 1.2e-13
Hematopoietic precursor cell
ρ = − 0.13, p = 2.4e-09
Immature B cell
ρ = − 0.43, p < 2.2e-16
Late pro-B cell
ρ = 0.043, p = 0.35
Macrophage
ρ = − 0.1, p = 0.0016
Megakaryocyte-erythroid progenitor cell
ρ = − 0.028, p = 0.23
Monocyte
ρ = 0.047, p = 0.042
Naive B cell
ρ = − 0.35, p = 1.2e-12
Naive T cell
ρ = − 0.012, p = 0.62
NK cell
ρ = − 0.38, p = 3.2e-06
Plasma cell
ρ = 0.09, p = 0.065
Precursor B cell
ρ = − 0.3, p = 4.8e-16
Proerythroblast
ρ = 0.096, p = 7e-09
Promonocyte
ρ = 0.059, p = 0.002
Cell-level dyscoordination (log-transformed)
Cell cycle S score
Age
01m
03m
18m
21m
24m
30m

**Supplementary Figure 17 Correlation between cell cycle S phase score and cell-level dyscoordination in mouse bone marrow cells.**

Scatter plots show the relationship between the cell cycle S phase score and log-transformed cell-level dyscoordination for each cell type. Black dashed lines indicate linear regression fits. Spearman's correlation coefficients and p-values are shown on the top left corner of each panel.

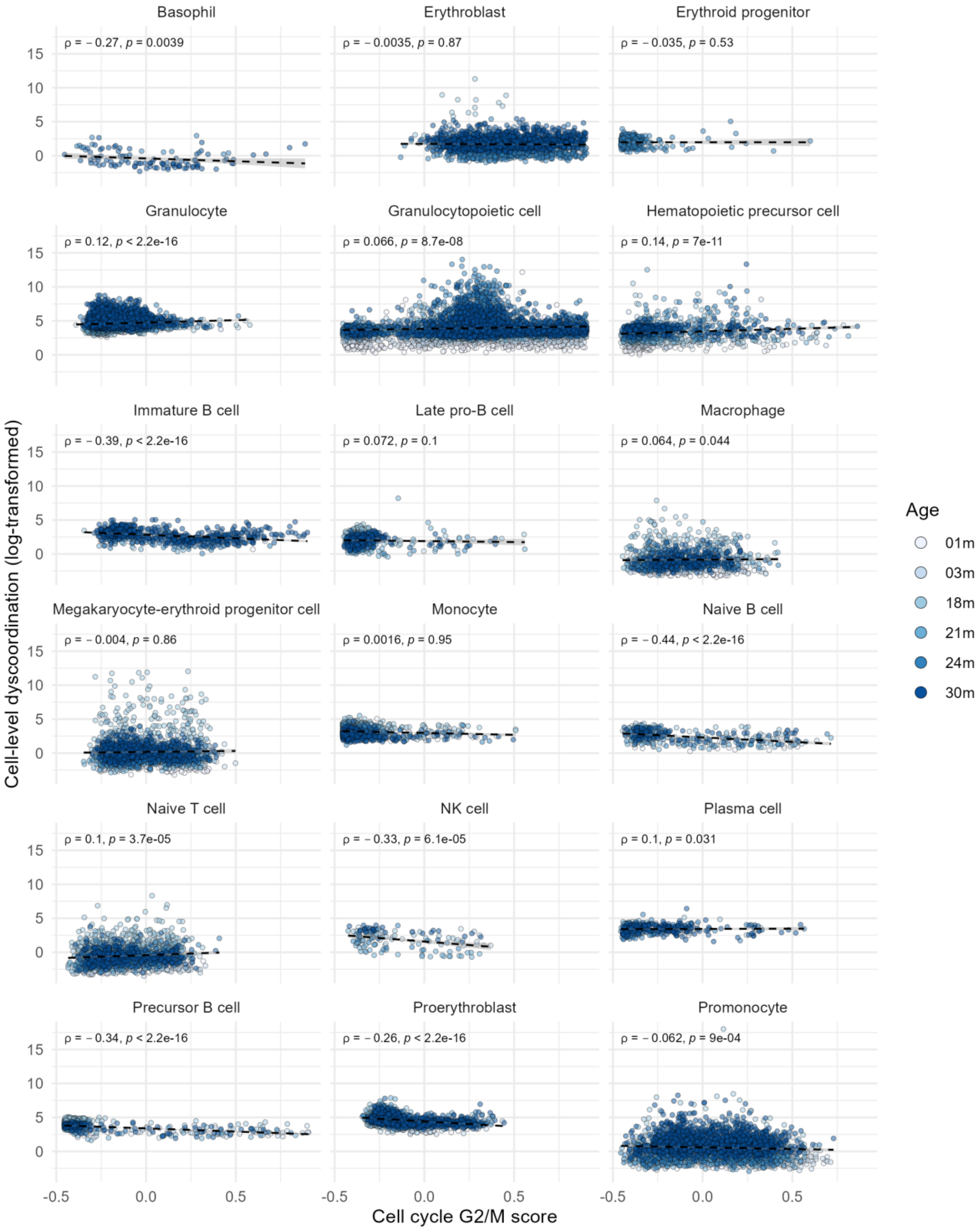

Basophil
ρ = −0.27, p = 0.0039
Erythroblast
ρ = −0.0035, p = 0.87
Erythroid progenitor
ρ = −0.035, p = 0.53
Granulocyte
ρ = 0.12, p < 2.2e-16
Granulocytopoietic cell
ρ = 0.066, p = 8.7e-08
Hematopoietic precursor cell
ρ = 0.14, p = 7e-11
Immature B cell
ρ = −0.39, p < 2.2e-16
Late pro-B cell
ρ = 0.072, p = 0.1
Macrophage
ρ = 0.064, p = 0.044
Megakaryocyte-erythroid progenitor cell
ρ = −0.004, p = 0.86
Monocyte
ρ = 0.0016, p = 0.95
Naive B cell
ρ = −0.44, p < 2.2e-16
Naive T cell
ρ = 0.1, p = 3.7e-05
NK cell
ρ = −0.33, p = 6.1e-05
Plasma cell
ρ = 0.1, p = 0.031
Precursor B cell
ρ = −0.34, p < 2.2e-16
Proerythroblast
ρ = −0.26, p < 2.2e-16
Promonocyte
ρ = −0.062, p = 9e-04
Cell-level dyscoordination (log-transformed)
Cell cycle G2/M score
Age
01m
03m
18m
21m
24m
30m

**Supplementary Figure 18 Correlation between cell cycle G2/M score and cell-level dyscoordination in mouse bone marrow cells.**

Scatter plots show the relationship between the cell cycle G2/M phase score and log-transformed cell-level dyscoordination for each cell type. Black dashed lines indicate linear regression fits. Spearman's correlation coefficients and p-values are shown on the top left corner of each panel.

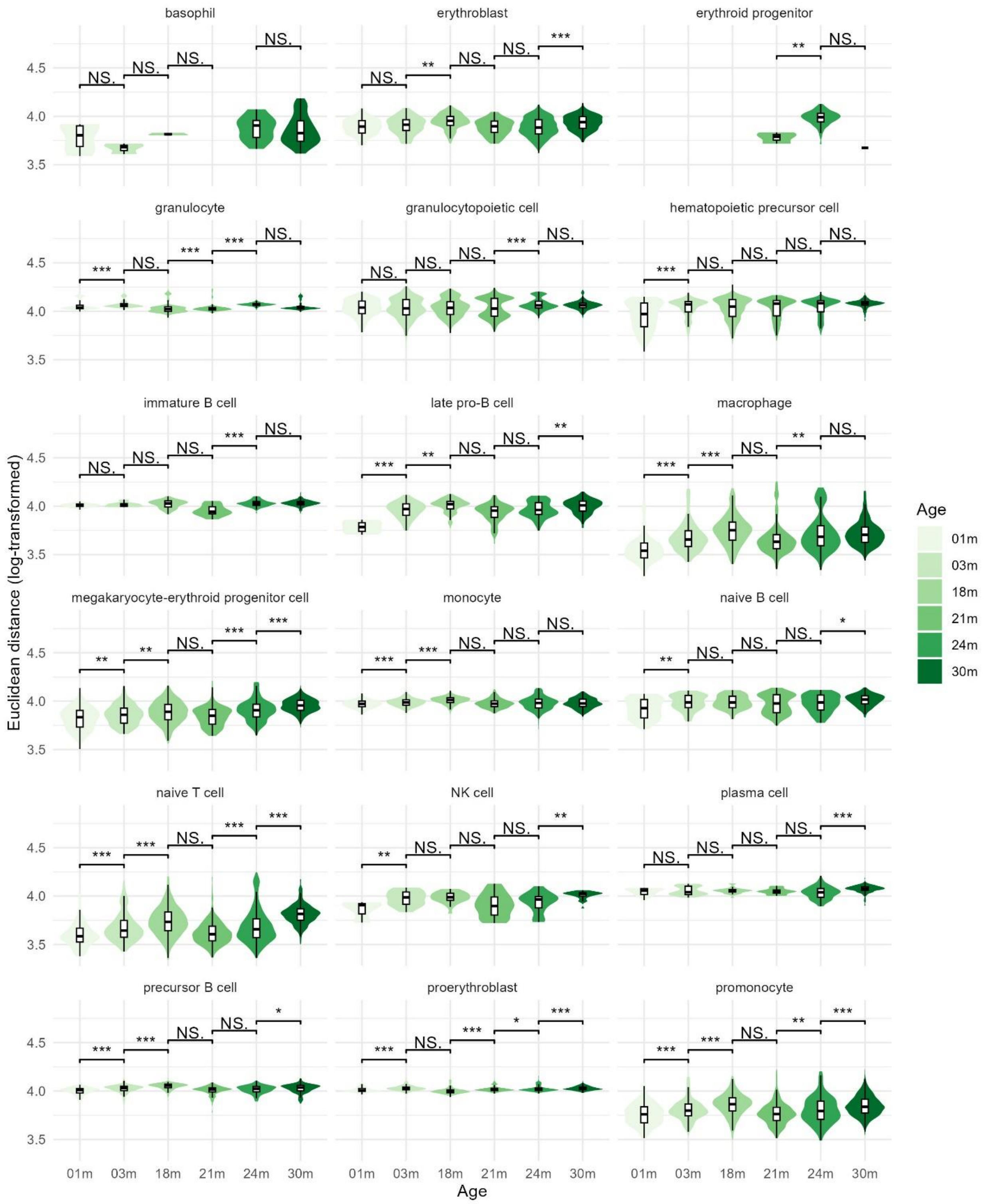

basophil
erythroblast
erythroid progenitor
granulocyte
granulocytopoietic cell
hematopoietic precursor cell
immature B cell
late pro-B cell
macrophage
megakaryocyte-erythroid progenitor cell
monocyte
naive B cell
naive T cell
NK cell
plasma cell
precursor B cell
proerythroblast
promonocyte
Euclidean distance (log-transformed)
Age
01m
03m
18m
21m
24m
30m
NS.

**Supplementary Figure 19 Distribution of Euclidean distance to cell type average in mouse bone marrow cells.**

Violin and box plots show the distribution of log-transformed Euclidean distance to cell type average across mouse bone marrow cell types and age groups. This method is adopted from decibel R package, which computes the Euclidean distance between each cell and the mean expression of its cell type across all genes. Pairwise comparisons between age groups were tested using one-sided Wilcoxon rank-sum test (alternative hypothesis: older age group has higher distance). Significance levels are annotated above each comparison (NS = not significant, * $p < 0.05$, ** $p < 0.01$, *** $p < 0.001$).

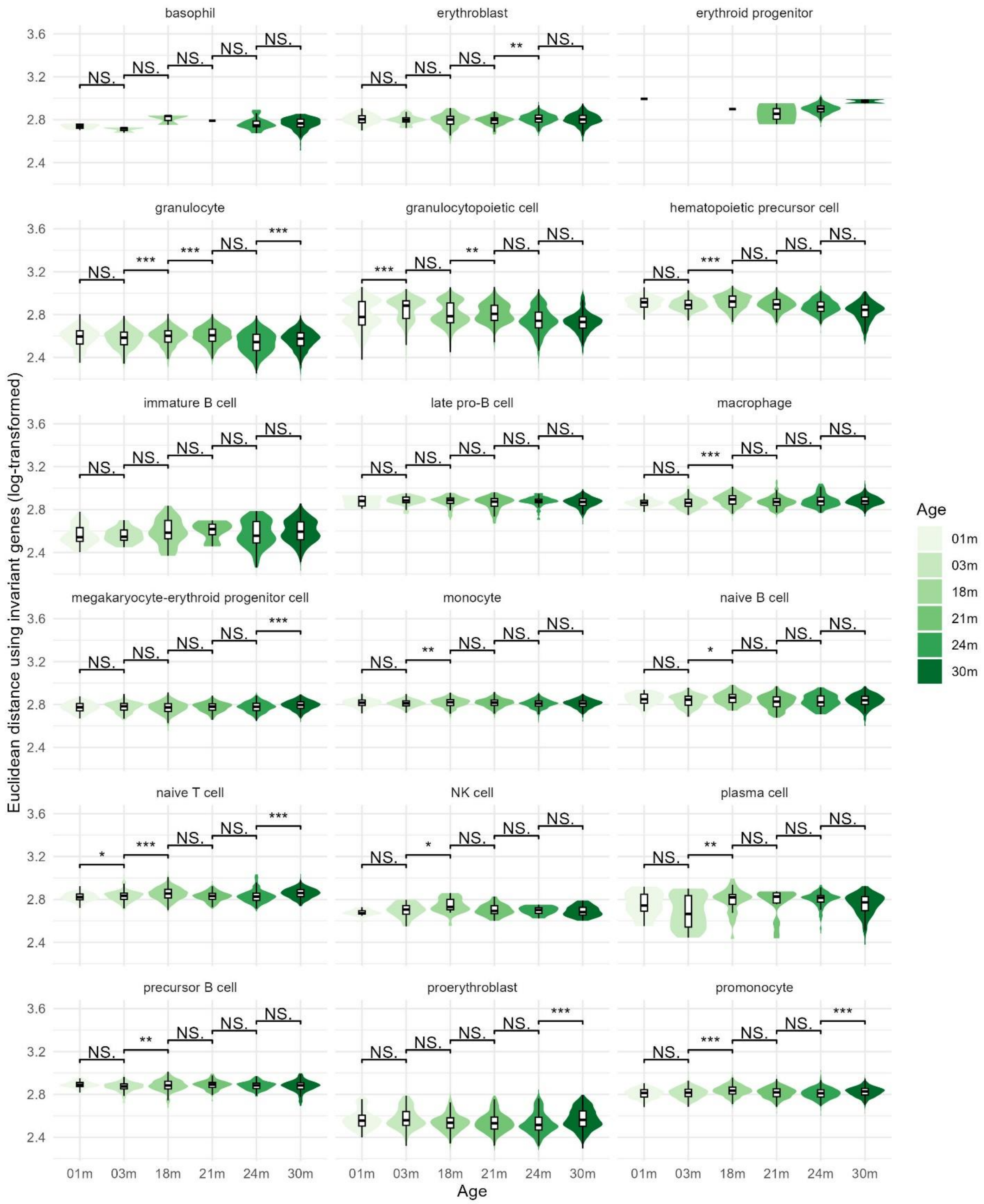
basophil
erythroblast
erythroid progenitor
granulocyte
granulocytopoietic cell
hematopoietic precursor cell
immature B cell
late pro-B cell
macrophage
megakaryocyte-erythroid progenitor cell
monocyte
naive B cell
naive T cell
NK cell
plasma cell
precursor B cell
proerythroblast
promonocyte
Euclidean distance using invariant genes (log-transformed)
Age
Age
01m
03m
18m
21m
24m
30m

**Supplementary Figure 20 Distribution of Euclidean distance to tissue average using invariant genes in mouse bone marrow cells.**

Violin and box plots show the distribution of log-transformed Euclidean distance to tissue average using invariant genes across mouse bone marrow cell types and age groups. This method is adopted from decibel R package, which computes the Euclidean distance between each cell and the mean expression of all cells in the tissue across all genes using a set of invariant genes. Details for how invariant genes are selected can be found in the package documentation. Pairwise comparisons between age groups were tested using one-sided Wilcoxon rank-sum test (alternative hypothesis: older age group has higher distance). Significance levels are annotated above each comparison (NS = not significant, * $p < 0.05$, ** $p < 0.01$, *** $p < 0.001$).

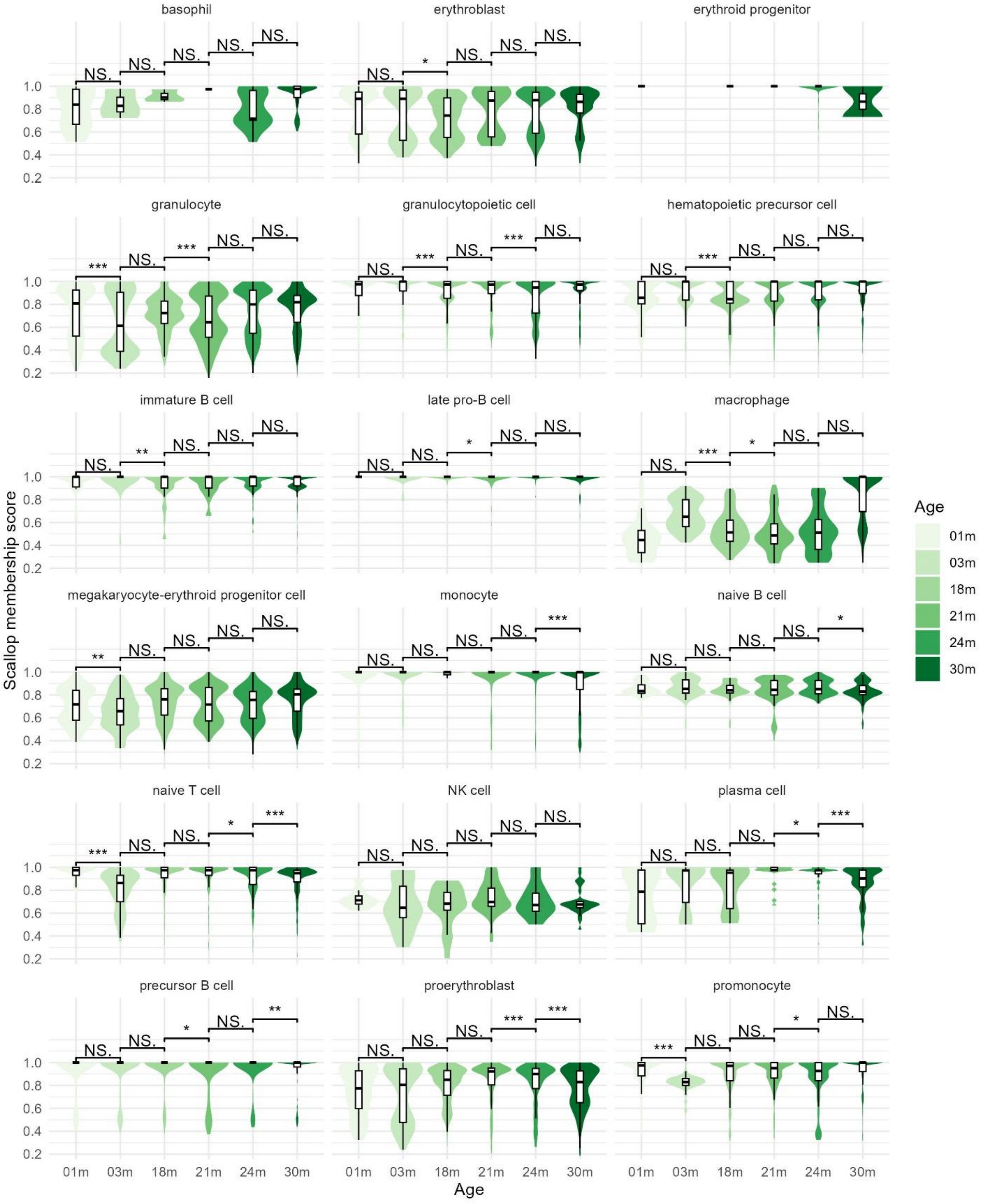
basophil
erythroblast
erythroid progenitor
granulocyte
granulocytopoietic cell
hematopoietic precursor cell
immature B cell
late pro-B cell
macrophage
megakaryocyte-erythroid progenitor cell
monocyte
naive B cell
naive T cell
NK cell
plasma cell
precursor B cell
proerythroblast
promonocyte
Scallop membership score
Age
01m
03m
18m
21m
24m
30m

**Supplementary Figure 21 Distribution of Scallop membership score in mouse bone marrow cells.**

Violin and box plots show the distribution of Scallop membership score across mouse bone marrow cell types and age groups. Scallop computes it as the frequency with which each cell is assigned to its most frequently assigned cluster. Since this metric reflects transcriptional stability, it is inversely related to our transcriptional dyscoordination measure. Pairwise comparisons between age groups were tested using one-sided Wilcoxon rank-sum test (alternative hypothesis: older age group has lower membership score). Significance levels are annotated above each comparison (NS = not significant, * $p < 0.05$, ** $p < 0.01$, *** $p < 0.001$).

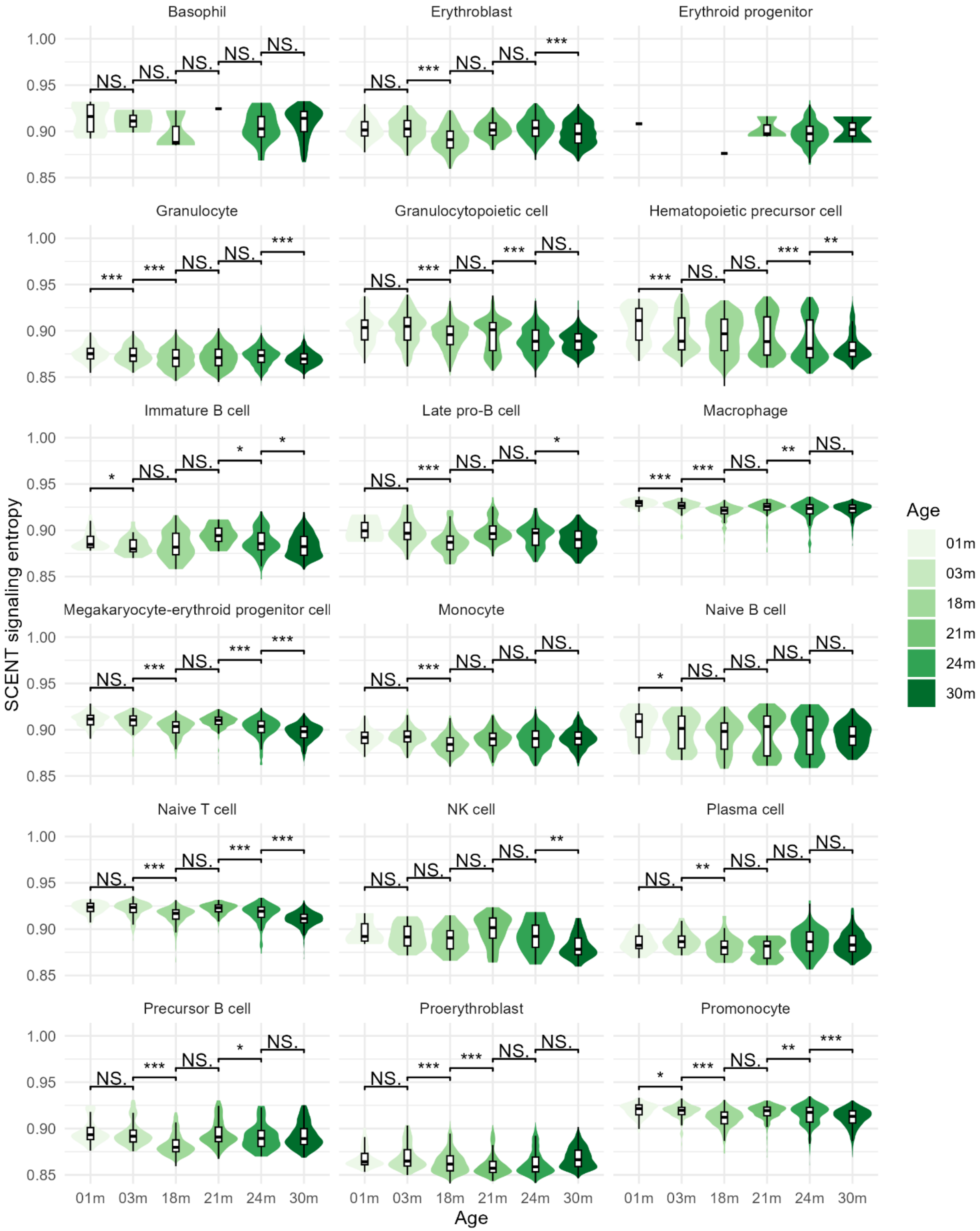
Basophil
Erythroblast
Erythroid progenitor
Granulocyte
Granulocytopoietic cell
Hematopoietic precursor cell
Immature B cell
Late pro-B cell
Macrophage
Megakaryocyte-erythroid progenitor cell
Monocyte
Naive B cell
Naive T cell
NK cell
Plasma cell
Precursor B cell
Proerythroblast
Promonocyte
SCENT signaling entropy
Age
01m
03m
18m
21m
24m
30m

**Supplementary Figure 22 Distribution of SCENT signaling entropy in mouse bone marrow cells.**

Violin and box plots show the distribution of SCENT signaling entropy across mouse bone marrow cell types and age groups. SCENT computes it by integrating each cell's gene expression with a PPI network and calculating the entropy rate of a random walk over the network. Since this metric reflects differentiation potency, it is inversely related to our transcriptional dyscoordination measure. Pairwise comparisons between age groups were tested using one-sided Wilcoxon rank-sum test (alternative hypothesis: older age group has lower signaling entropy). Significance levels are annotated above each comparison (NS = not significant, * $p < 0.05$, ** $p < 0.01$, *** $p < 0.001$).

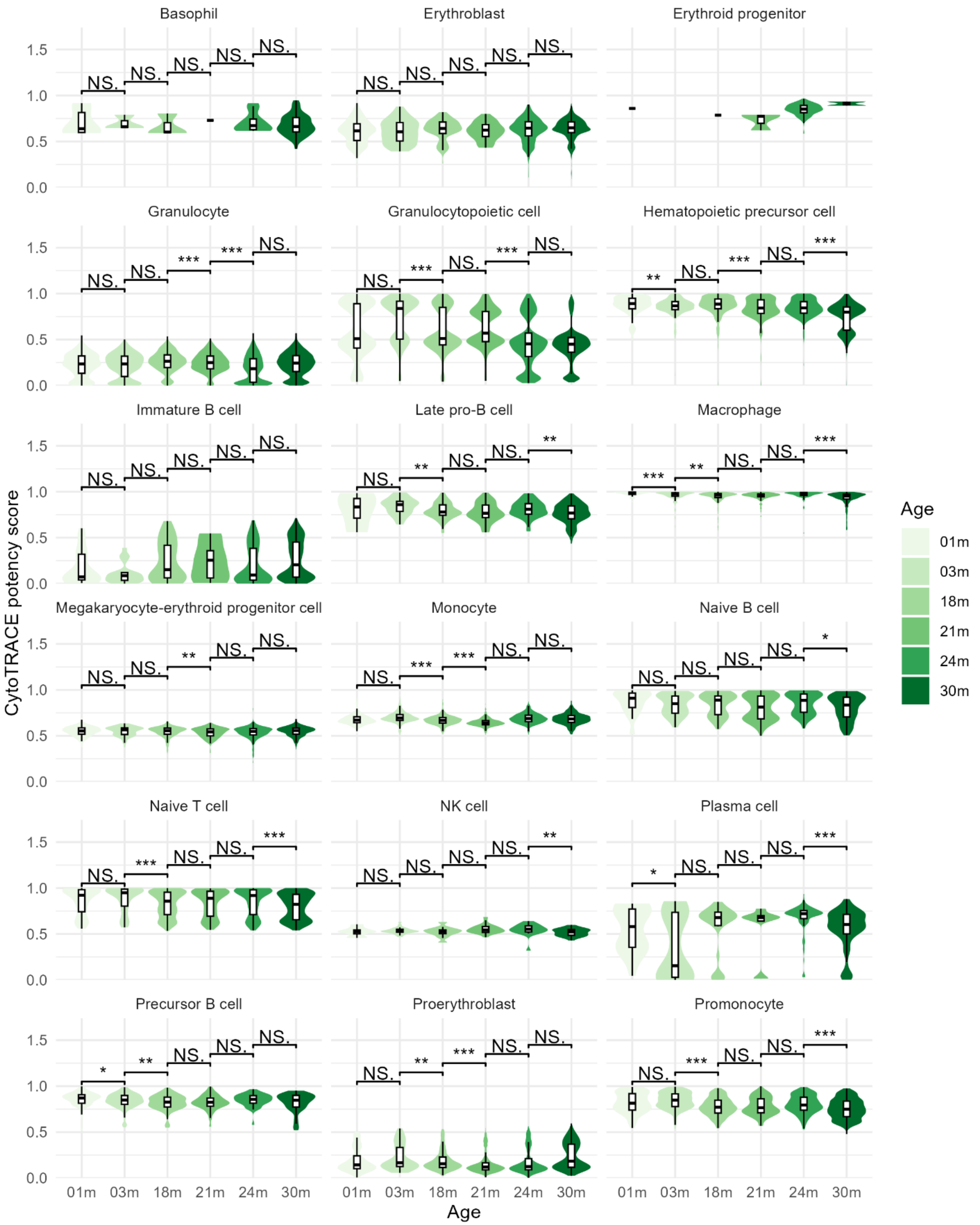

Basophil
Erythroblast
Erythroid progenitor
Granulocyte
Granulocytopoietic cell
Hematopoietic precursor cell
Immature B cell
Late pro-B cell
Macrophage
Megakaryocyte-erythroid progenitor cell
Monocyte
Naive B cell
Naive T cell
NK cell
Plasma cell
Precursor B cell
Proerythroblast
Promonocyte
CytoTRACE potency score
Age
01m
03m
18m
21m
24m
30m

**Supplementary Figure 23 Distribution of CytoTRACE potency score in mouse bone marrow cells.**

Violin and box plots show the distribution of CytoTRACE potency score across mouse bone marrow cell types and age groups. CytoTRACE computes it from the number of distinctly expressed genes per cell, with more diverse transcriptome indicating higher potency. Since this metric reflects differentiation potency, it is inversely related to our transcriptional dyscoordination measure. Pairwise comparisons between age groups were tested using one-sided Wilcoxon rank-sum test (alternative hypothesis: older age group has lower potency score). Significance levels are annotated above each comparison (NS = not significant, * $p < 0.05$, ** $p < 0.01$, *** $p < 0.001$).

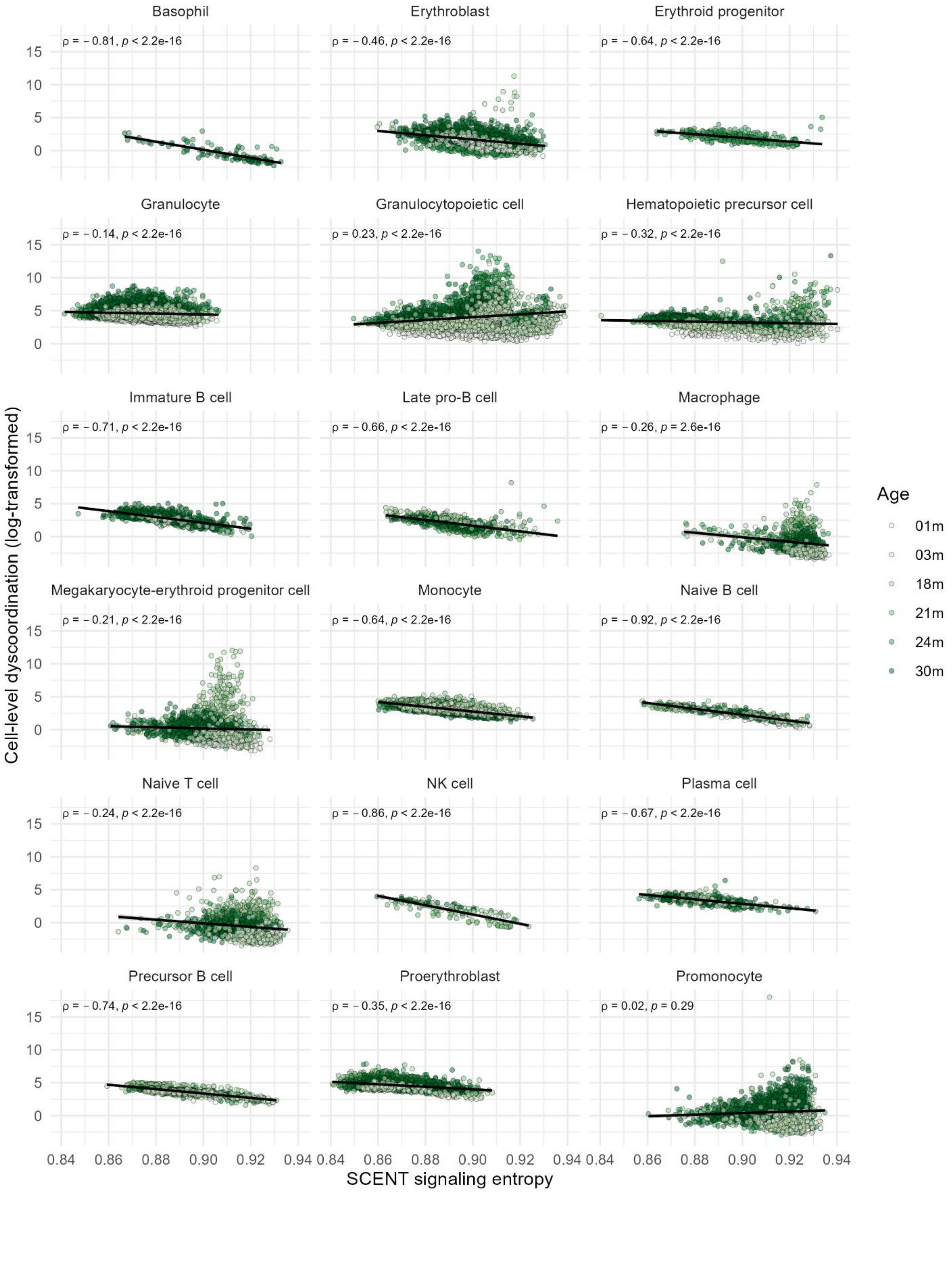
Basophil
ρ = − 0.81, p < 2.2e-16
Erythroblast
ρ = − 0.46, p < 2.2e-16
Erythroid progenitor
ρ = − 0.64, p < 2.2e-16
Granulocyte
ρ = − 0.14, p < 2.2e-16
Granulocytopoietic cell
ρ = 0.23, p < 2.2e-16
Hematopoietic precursor cell
ρ = − 0.32, p < 2.2e-16
Immature B cell
ρ = − 0.71, p < 2.2e-16
Late pro-B cell
ρ = − 0.66, p < 2.2e-16
Macrophage
ρ = − 0.26, p = 2.6e-16
Megakaryocyte-erythroid progenitor cell
ρ = − 0.21, p < 2.2e-16
Monocyte
ρ = − 0.64, p < 2.2e-16
Naive B cell
ρ = − 0.92, p < 2.2e-16
Naive T cell
ρ = − 0.24, p < 2.2e-16
NK cell
ρ = − 0.86, p < 2.2e-16
Plasma cell
ρ = − 0.67, p < 2.2e-16
Precursor B cell
ρ = − 0.74, p < 2.2e-16
Proerythroblast
ρ = − 0.35, p < 2.2e-16
Promonocyte
ρ = 0.02, p = 0.29
Cell-level dyscoordination (log-transformed)
SCENT signaling entropy
Age
01m
03m
18m
21m
24m
30m

**Supplementary Figure 24 Correlation between SCENT signaling entropy and cell-level dyscoordination in mouse bone marrow cells.**

Scatter plots show the relationship between SCENT signaling entropy and log-transformed cell-level dyscoordination across cell types and age groups. Black lines indicate linear regression fits. Spearman correlation coefficients and corresponding p-values are shown in the top left corner of each panel.

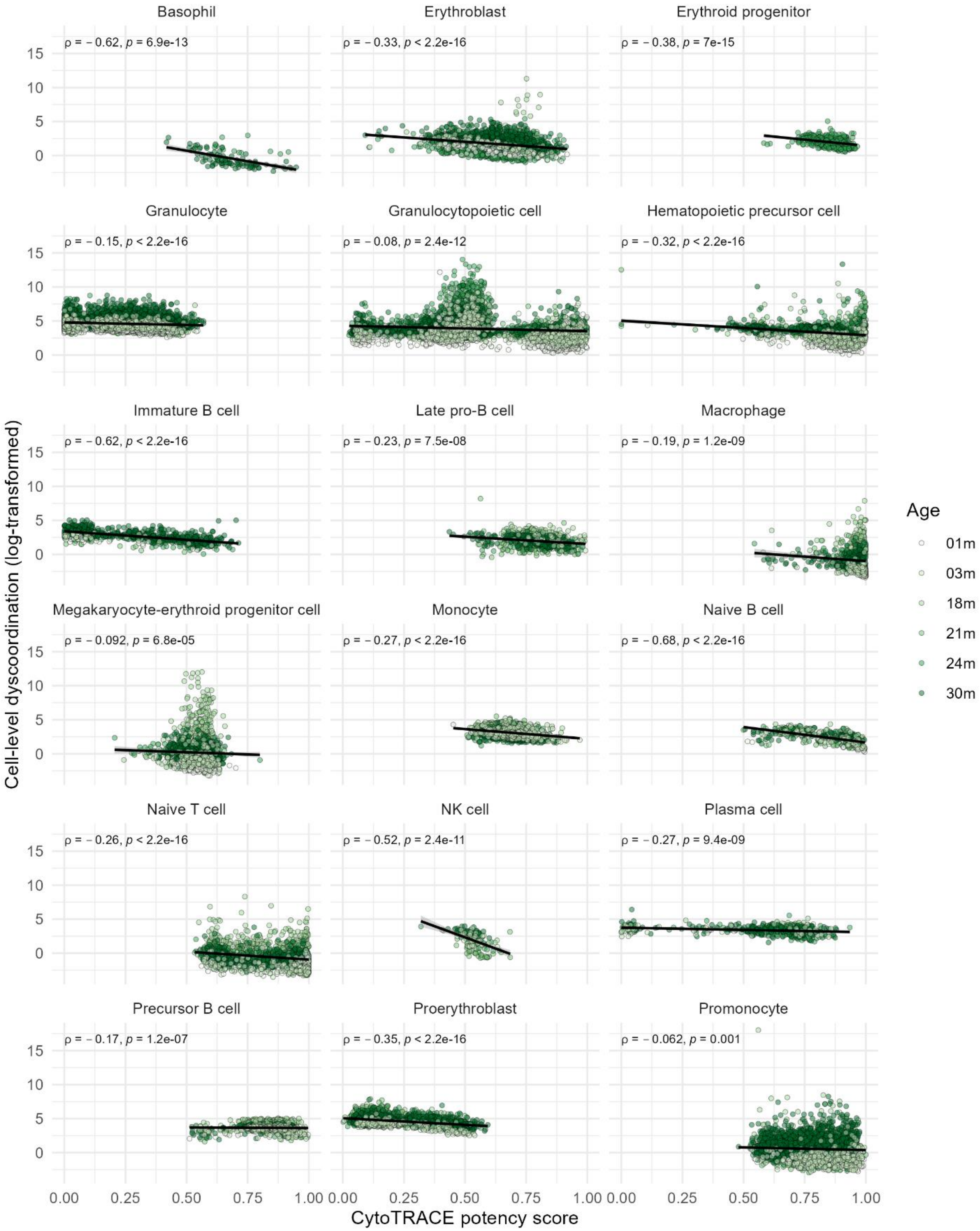

Basophil
ρ = − 0.62, p = 6.9e-13
Erythroblast
ρ = − 0.33, p < 2.2e-16
Erythroid progenitor
ρ = − 0.38, p = 7e-15
Granulocyte
ρ = − 0.15, p < 2.2e-16
Granulocytopoietic cell
ρ = − 0.08, p = 2.4e-12
Hematopoietic precursor cell
ρ = − 0.32, p < 2.2e-16
Immature B cell
ρ = − 0.62, p < 2.2e-16
Late pro-B cell
ρ = − 0.23, p = 7.5e-08
Macrophage
ρ = − 0.19, p = 1.2e-09
Megakaryocyte-erythroid progenitor cell
ρ = − 0.092, p = 6.8e-05
Monocyte
ρ = − 0.27, p < 2.2e-16
Naive B cell
ρ = − 0.68, p < 2.2e-16
Naive T cell
ρ = − 0.26, p < 2.2e-16
NK cell
ρ = − 0.52, p = 2.4e-11
Plasma cell
ρ = − 0.27, p = 9.4e-09
Precursor B cell
ρ = − 0.17, p = 1.2e-07
Proerythroblast
ρ = − 0.35, p < 2.2e-16
Promonocyte
ρ = − 0.062, p = 0.001
Cell-level dyscoordination (log-transformed)
CytoTRACE potency score
0.00 0.25 0.50 0.75 1.00
0 5 10 15
Age
01m
03m
18m
21m
24m
30m

**Supplementary Figure 25 Correlation between CytoTRACE potency score and cell-level dyscoordination in mouse bone marrow cells.**

Scatter plots show the relationship between CytoTRACE potency score and log-transformed cell-level dyscoordination across cell types and age groups. Black lines indicate linear regression fits. Spearman correlation coefficients and corresponding p-values are shown in the top left corner of each panel.

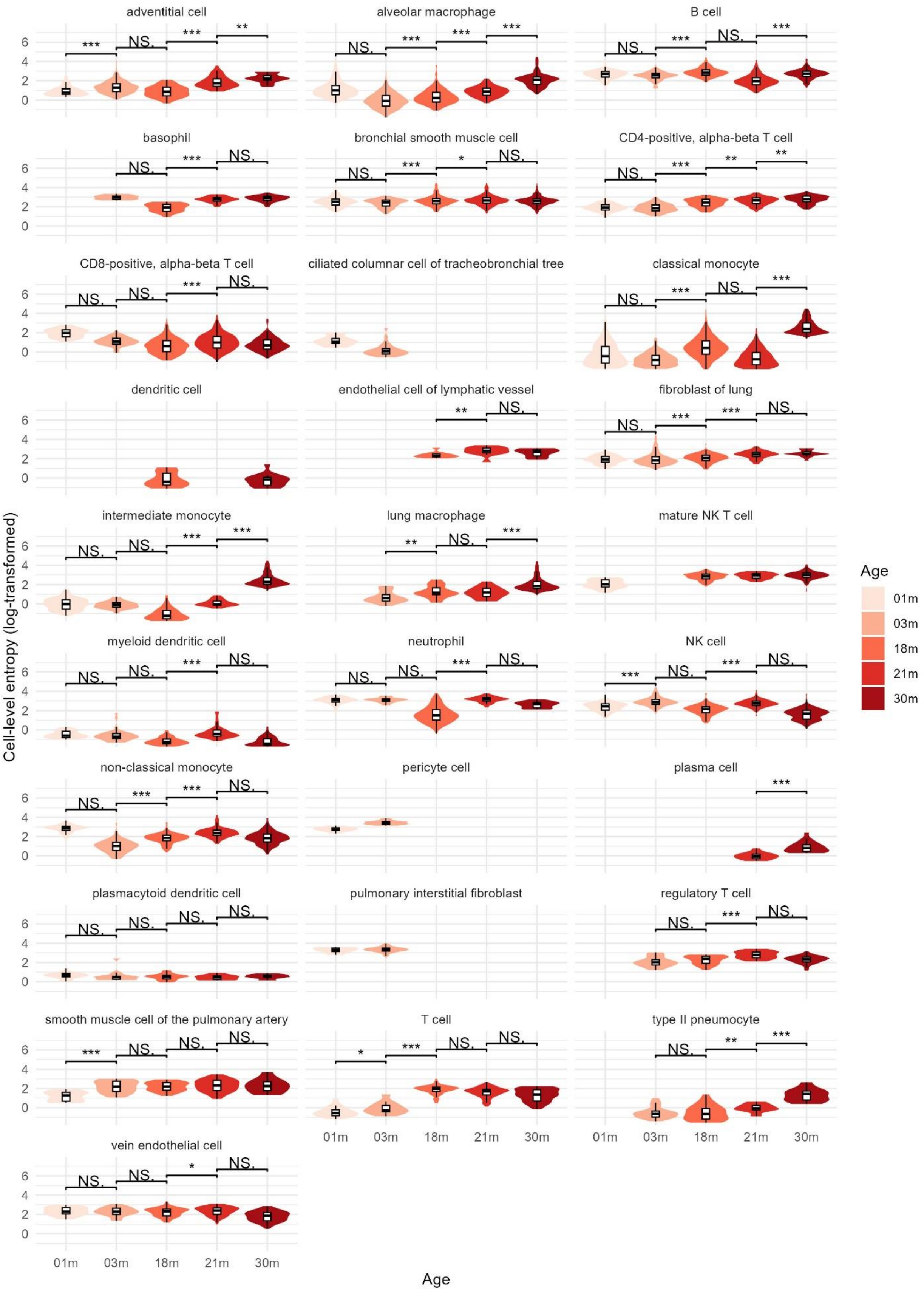
adventitial cell
alveolar macrophage
B cell
basophil
bronchial smooth muscle cell
CD4-positive, alpha-beta T cell
CD8-positive, alpha-beta T cell
ciliated columnar cell of tracheobronchial tree
classical monocyte
dendritic cell
endothelial cell of lymphatic vessel
fibroblast of lung
intermediate monocyte
lung macrophage
mature NK T cell
myeloid dendritic cell
neutrophil
NK cell
non-classical monocyte
pericyte cell
plasma cell
plasmacytoid dendritic cell
pulmonary interstitial fibroblast
regulatory T cell
smooth muscle cell of the pulmonary artery
T cell
type II pneumocyte
vein endothelial cell
Cell-level entropy (log-transformed)
Age
01m
03m
18m
21m
30m

**Supplementary Figure 26 Distribution of cell-level dyscoordination in lung cells.**

Violin and box plots show the distribution of log-transformed cell-level dyscoordination values separately for each cell type and age group. Values in the top 5% and bottom 1% quantiles are considered outliers and are removed from the illustration. Pairwise comparisons between age groups were tested using one-sided Wilcoxon rank-sum test (alternative hypothesis: older age group has higher dyscoordination). Significance levels are annotated above each comparison (NS = not significant, * $p < 0.05$, ** $p < 0.01$, *** $p < 0.001$).

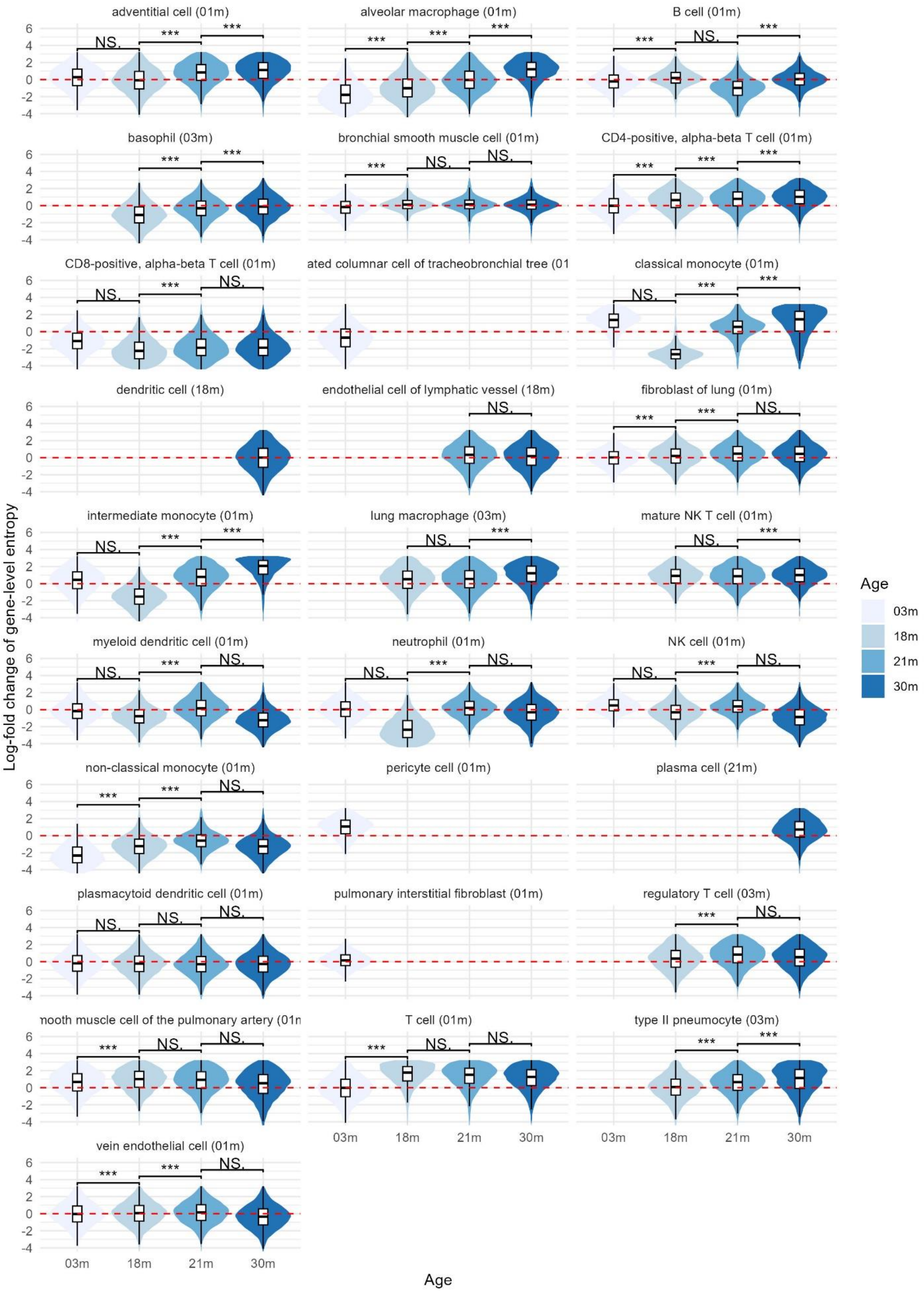

adventitial cell (01m)
alveolar macrophage (01m)
B cell (01m)
basophil (03m)
bronchial smooth muscle cell (01m)
CD4-positive, alpha-beta T cell (01m)
CD8-positive, alpha-beta T cell (01m)
ated columnar cell of tracheobronchial tree (01
classical monocyte (01m)
dendritic cell (18m)
endothelial cell of lymphatic vessel (18m)
fibroblast of lung (01m)
intermediate monocyte (01m)
lung macrophage (03m)
mature NK T cell (01m)
myeloid dendritic cell (01m)
neutrophil (01m)
NK cell (01m)
non-classical monocyte (01m)
pericyte cell (01m)
plasma cell (21m)
plasmacytoid dendritic cell (01m)
pulmonary interstitial fibroblast (01m)
regulatory T cell (03m)
nooth muscle cell of the pulmonary artery (01n
T cell (01m)
type II pneumocyte (03m)
vein endothelial cell (01m)
Log-fold change of gene-level entropy
Age
03m
18m
21m
30m

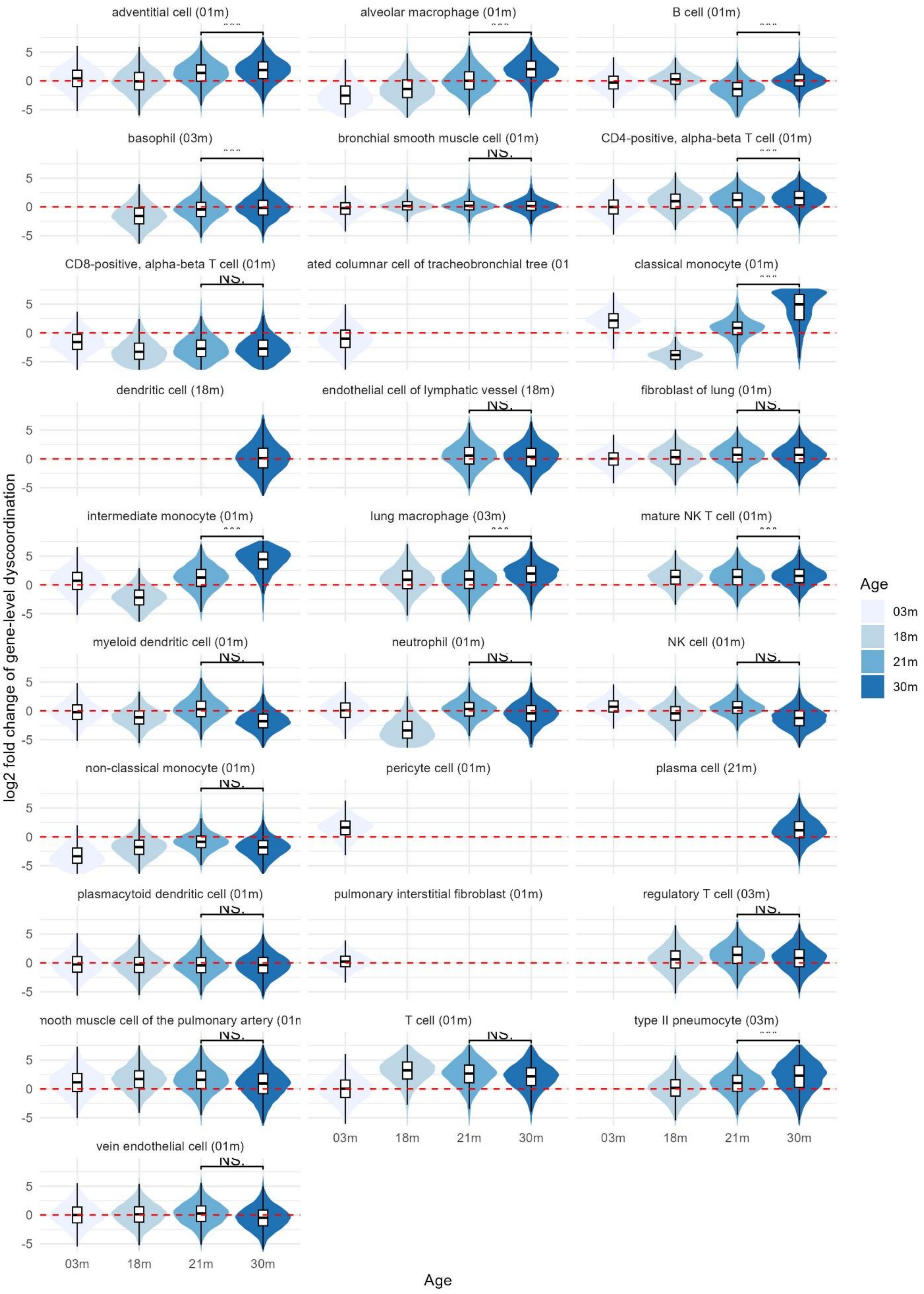

adventitial cell (01m)
alveolar macrophage (01m)
B cell (01m)
basophil (03m)
bronchial smooth muscle cell (01m)
CD4-positive, alpha-beta T cell (01m)
CD8-positive, alpha-beta T cell (01m)
ated columnar cell of tracheobronchial tree (01
classical monocyte (01m)
dendritic cell (18m)
endothelial cell of lymphatic vessel (18m)
fibroblast of lung (01m)
intermediate monocyte (01m)
lung macrophage (03m)
mature NK T cell (01m)
myeloid dendritic cell (01m)
neutrophil (01m)
NK cell (01m)
non-classical monocyte (01m)
pericyte cell (01m)
plasma cell (21m)
plasmacytoid dendritic cell (01m)
pulmonary interstitial fibroblast (01m)
regulatory T cell (03m)
nooth muscle cell of the pulmonary artery (01n
T cell (01m)
type II pneumocyte (03m)
vein endothelial cell (01m)
NS.
5
0
-5
03m
18m
21m
30m
log2 fold change of gene-level dyscoordination
Age
Age
03m
18m
21m
30m

**Supplementary Figure 27 Distribution of gene-level dyscoordination in lung cells.**

Violin and box plots show the distribution of log-fold changes in dyscoordination relative to the baseline age group. Values are displayed separately for each cell type and age group, with the baseline age group indicated in parentheses in each subplot title. Values in the top 5% and bottom 1% quantiles are considered outliers and are removed from the illustration. Red dashed line denotes zero log-fold change. Pairwise comparisons between age groups were tested using one-sided Wilcoxon rank-sum test (alternative hypothesis: older age group has higher dyscoordination). Significance levels are annotated above each comparison (NS = not significant, * $p < 0.05$, ** $p < 0.01$, *** $p < 0.001$).

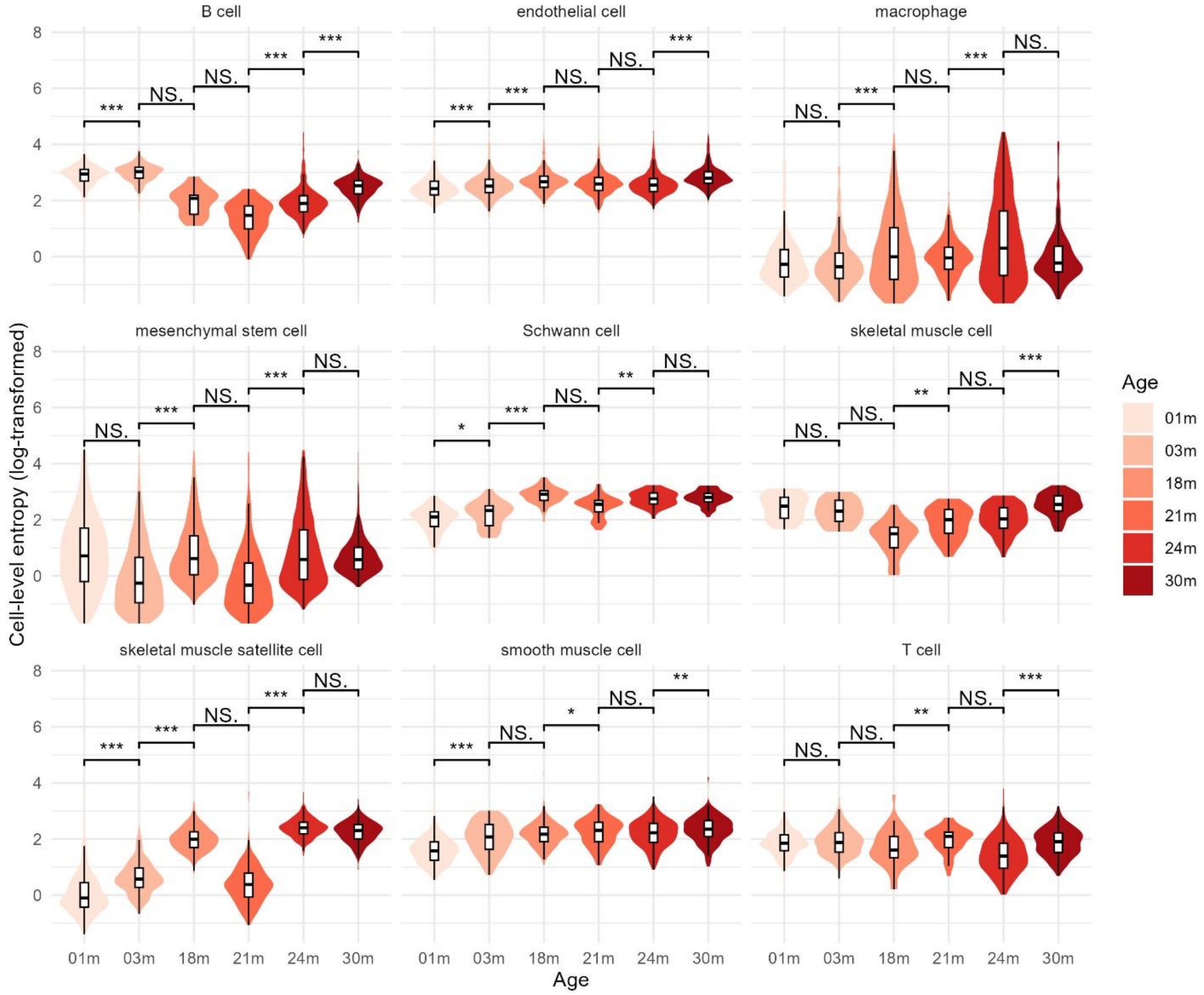


**Supplementary Figure 28 Distribution of cell-level dyscoordination in limb muscle cells.**

Violin and box plots show the distribution of log-transformed cell-level dyscoordination values separately for each cell type and age group. Values in the top and bottom 1% quantiles are considered outliers and are removed from the illustration. Pairwise comparisons between age groups were tested using one-sided Wilcoxon rank-sum test (alternative hypothesis: older age group has higher dyscoordination). Significance levels are annotated above each comparison (NS = not significant, * $p < 0.05$, ** $p < 0.01$, *** $p < 0.001$).

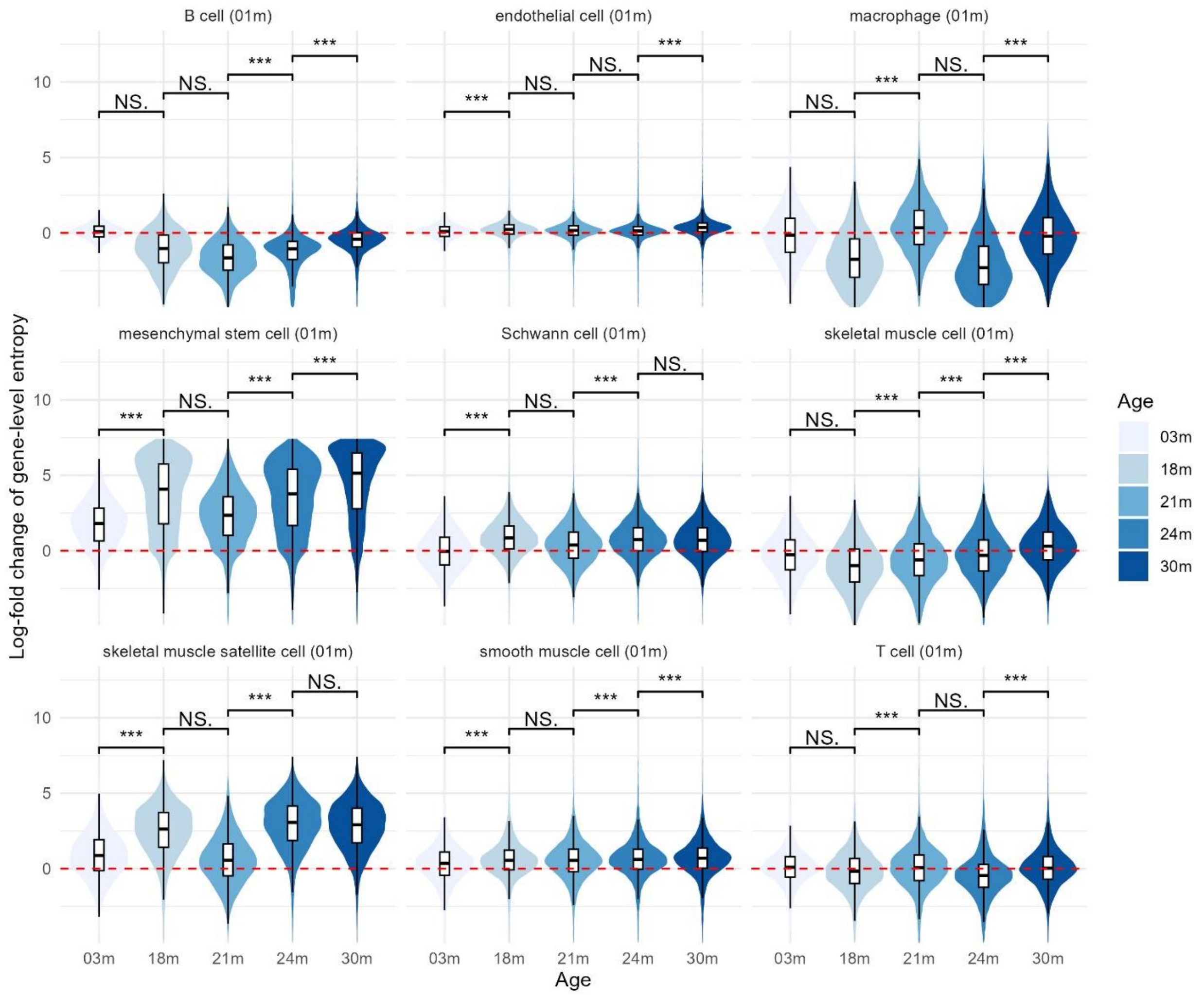


**Supplementary Figure 29 Distribution of gene-level dyscoordination in limb muscle cells.**

Violin and box plots show the distribution of log-fold changes in dyscoordination relative to the baseline age group. Values are displayed separately for each cell type and age group, with the baseline age group indicated in parentheses in each subplot title. Values in the top and bottom 1% quantiles are considered outliers and are removed from the illustration. Red dashed line denotes zero log-fold change. Pairwise comparisons between age groups were tested using one-sided Wilcoxon rank-sum test (alternative hypothesis: older age group has higher dyscoordination). Significance levels are annotated above each comparison (NS = not significant, * p < 0.05, ** p < 0.01, *** p < 0.001).

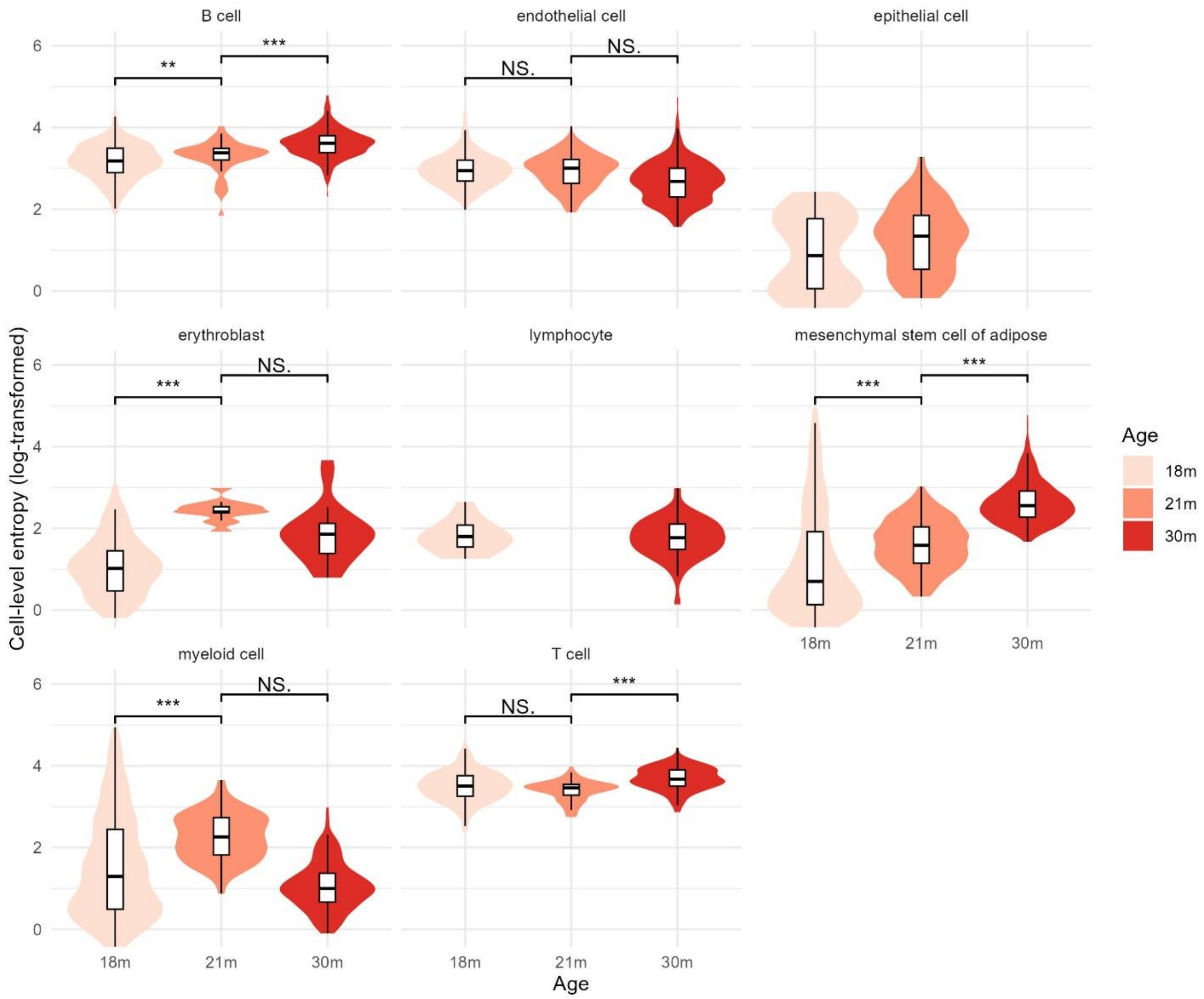


**Supplementary Figure 30 Distribution of cell-level dyscoordination in adipose tissue cells.**

Violin and box plots show the distribution of log-transformed cell-level dyscoordination values separately for each cell type and age group. Values in the top and bottom 1% quantiles are considered outliers and are removed from the illustration. Pairwise comparisons between age groups were tested using one-sided Wilcoxon rank-sum test (alternative hypothesis: older age group has higher dyscoordination). Significance levels are annotated above each comparison (NS = not significant, * $p < 0.05$, ** $p < 0.01$, *** $p < 0.001$).

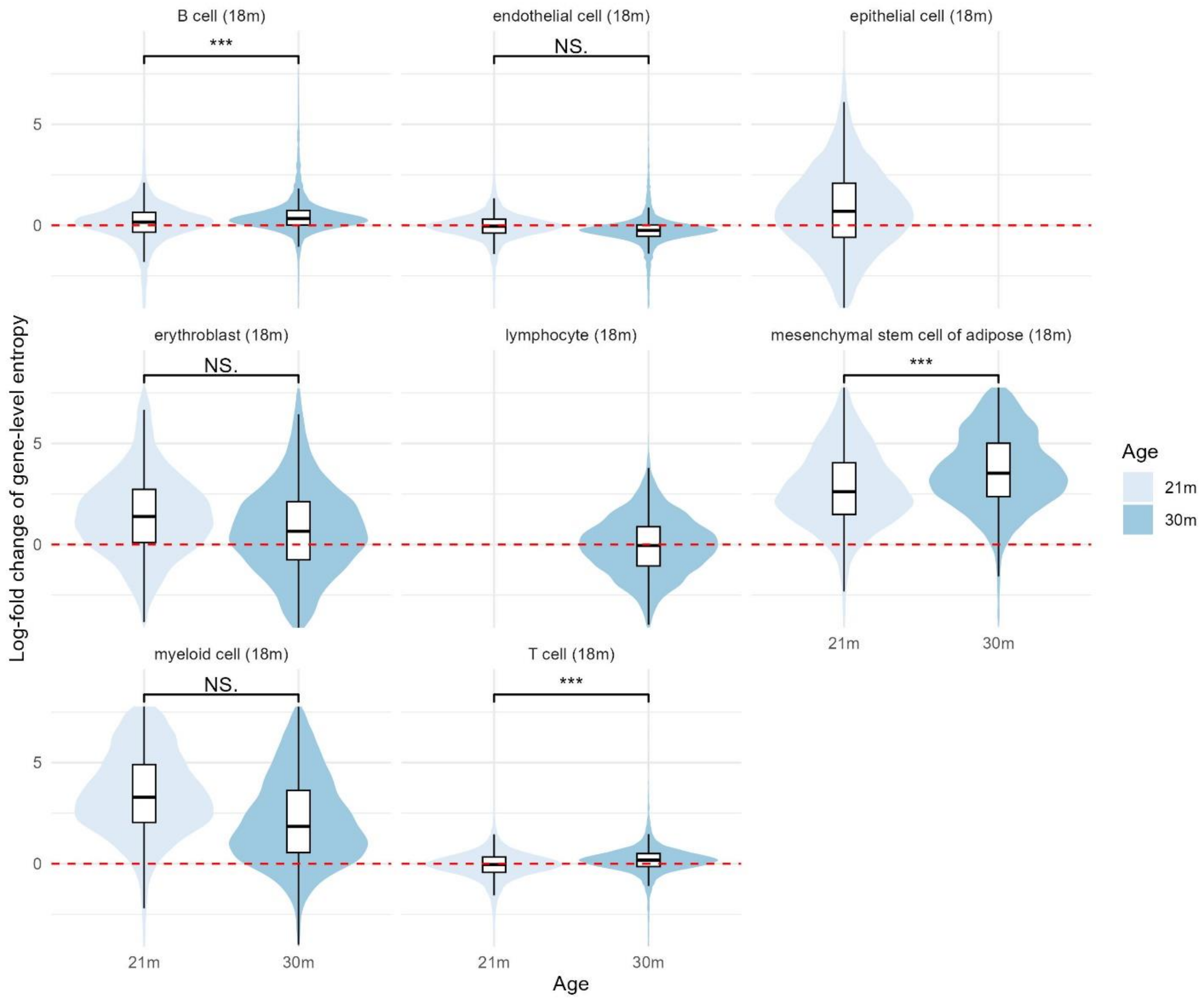


**Supplementary Figure 31 Distribution of gene-level dyscoordination in adipose tissue cells.**

Violin and box plots show the distribution of log-fold changes in dyscoordination relative to the baseline age group. Values are displayed separately for each cell type and age group, with the baseline age group indicated in parentheses in each subplot title. Values in the top and bottom 1% quantiles are considered outliers and are removed from the illustration. Red dashed line denotes zero log-fold change. Pairwise comparisons between age groups were tested using one-sided Wilcoxon rank-sum test (alternative hypothesis: older age group has higher dyscoordination). Significance levels are annotated above each comparison (NS = not significant, * $p < 0.05$, ** $p < 0.01$, *** $p < 0.001$).

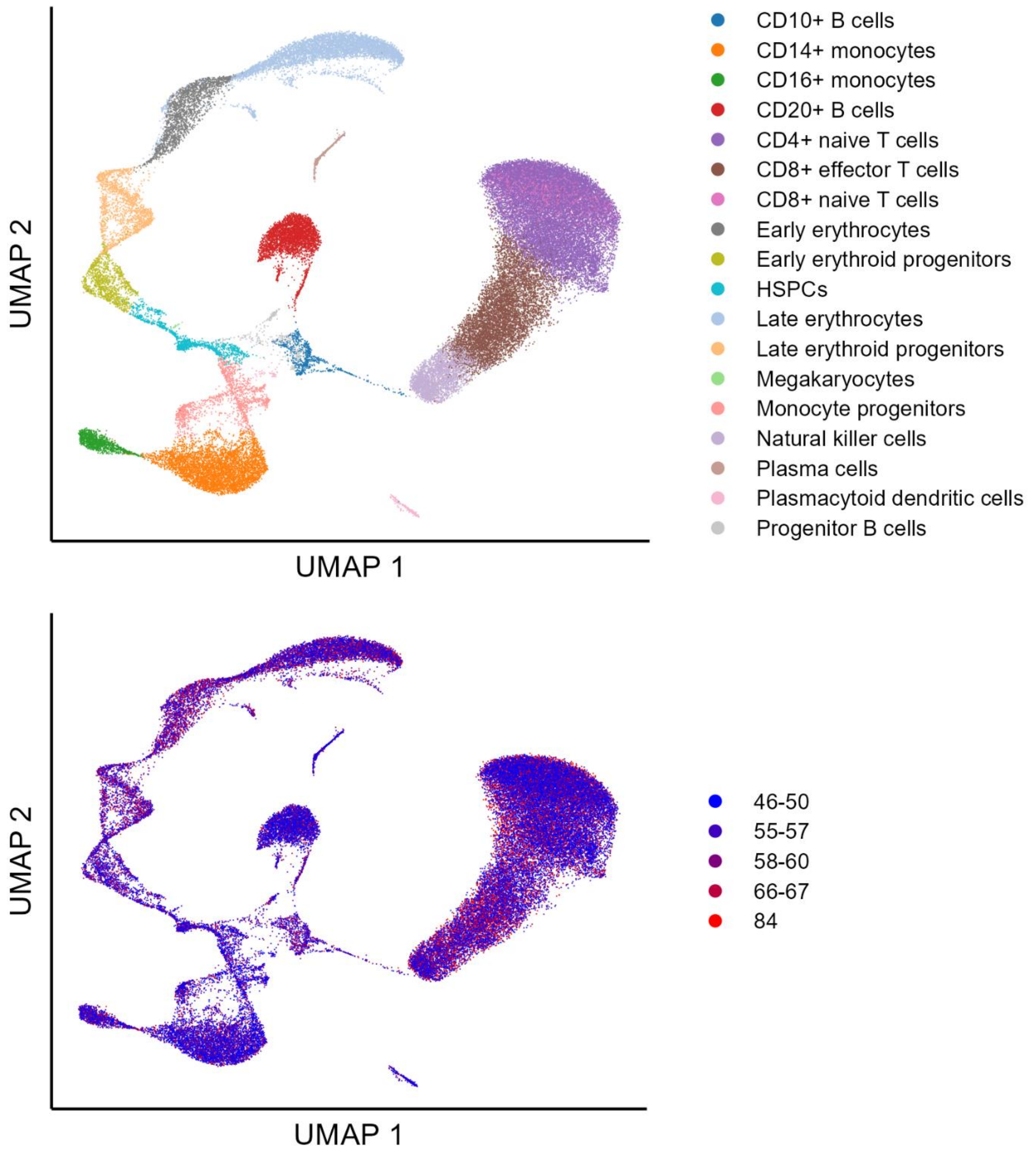


**Supplementary Figure 32 UMAP visualization of aging human bone marrow cells.**

Cells are embedded using the first two UMAP dimensions and colored by annotated cell type (top) and age group (bottom).

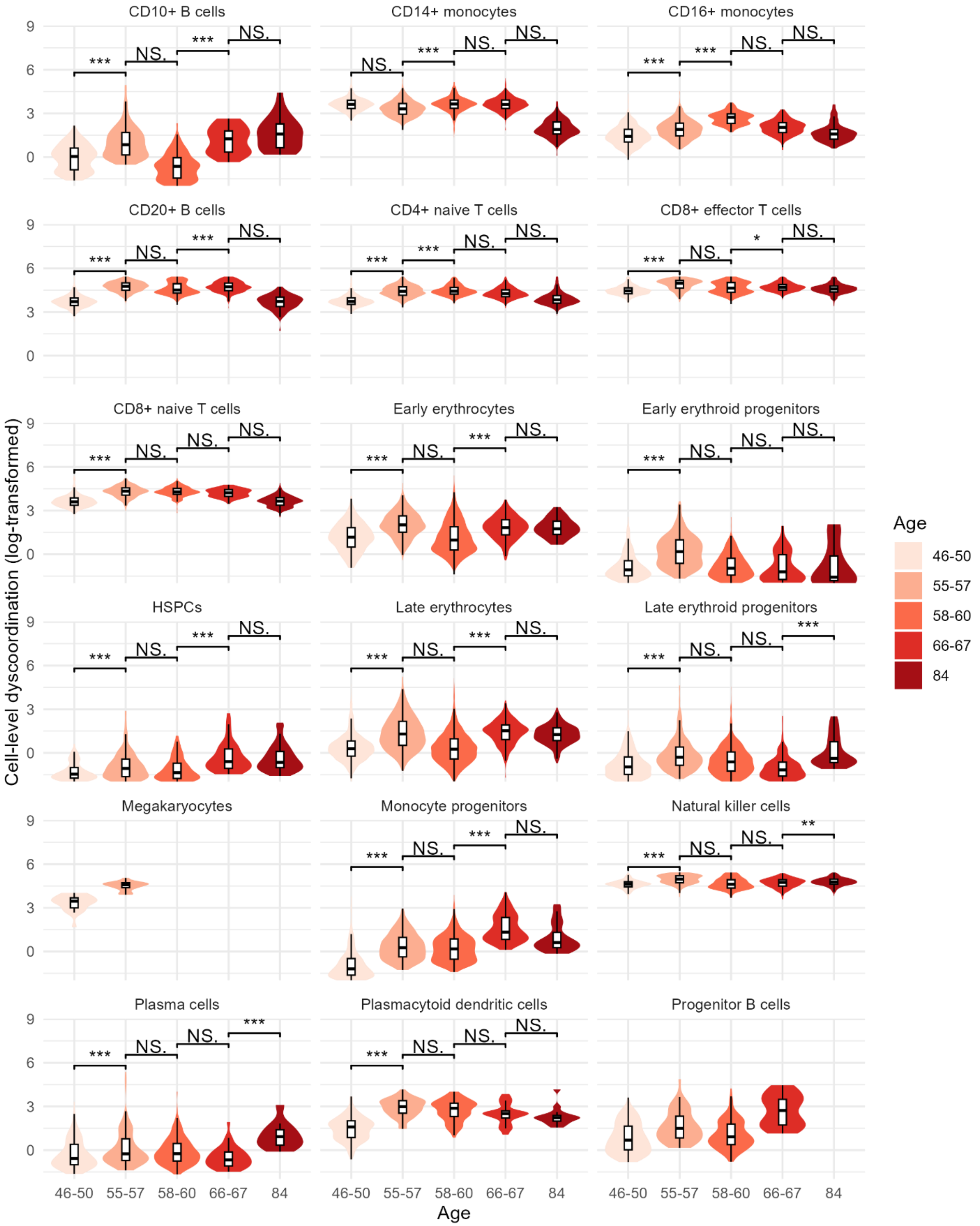

CD10+ B cells
CD14+ monocytes
CD16+ monocytes
CD20+ B cells
CD4+ naive T cells
CD8+ effector T cells
CD8+ naive T cells
Early erythrocytes
Early erythroid progenitors
HSPCs
Late erythrocytes
Late erythroid progenitors
Megakaryocytes
Monocyte progenitors
Natural killer cells
Plasma cells
Plasmacytoid dendritic cells
Progenitor B cells
Cell-level dyscoordination (log-transformed)
Age
46-50
55-57
58-60
66-67
84

**Supplementary Figure 33 Distribution of cell-level dyscoordination in human bone marrow cells.**

Violin and box plots show the distribution of log-transformed cell-level dyscoordination values separately for each cell type and age group. Values in the top and bottom 2.5% quantiles are considered outliers and are removed from the illustration. Pairwise comparisons between age groups were tested using one-sided Wilcoxon rank-sum test (alternative hypothesis: older age group has higher dyscoordination). Significance levels are annotated above each comparison (NS = not significant, * $p < 0.05$, ** $p < 0.01$, *** $p < 0.001$).

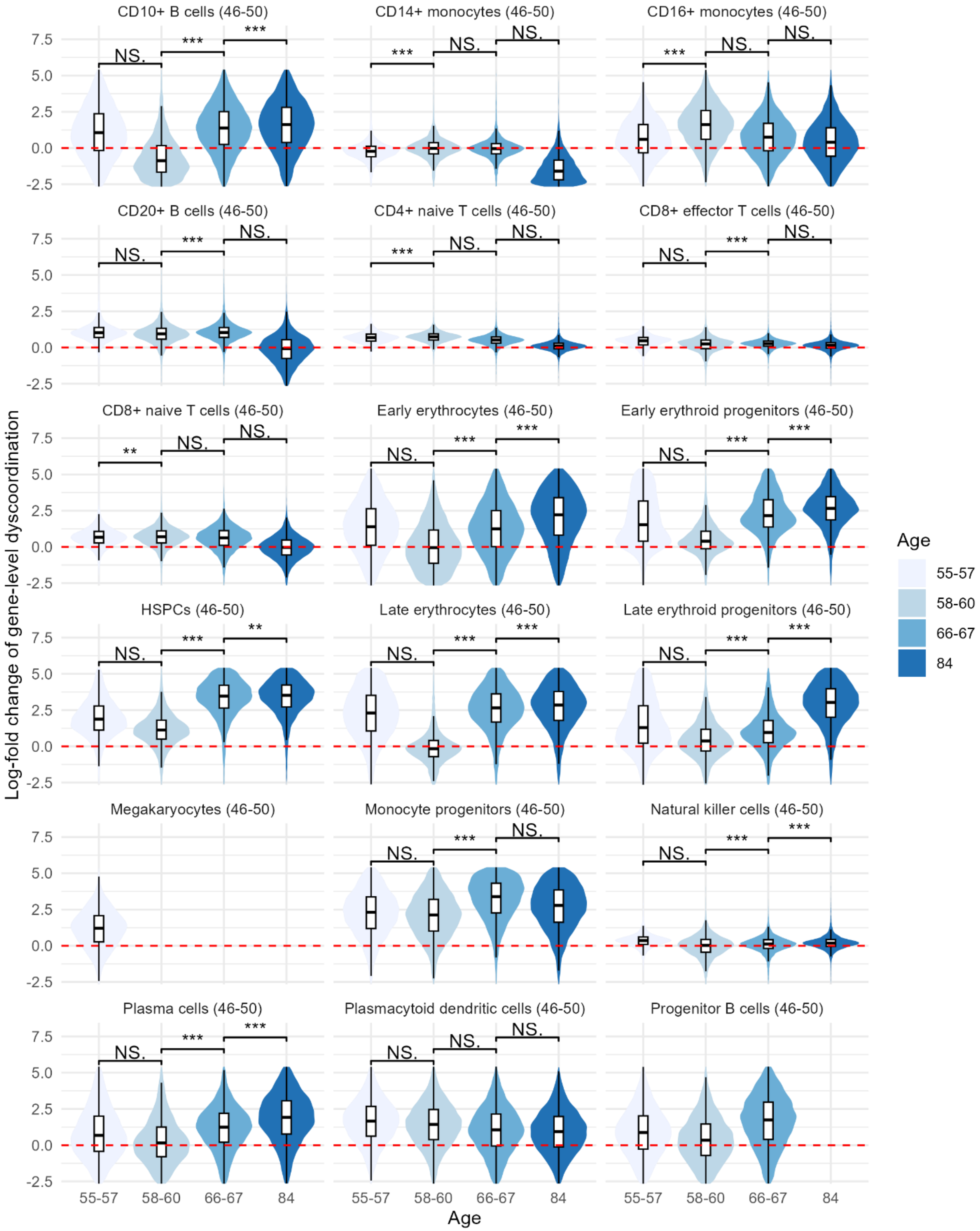

CD10+ B cells (46-50)
CD14+ monocytes (46-50)
CD16+ monocytes (46-50)
CD20+ B cells (46-50)
CD4+ naive T cells (46-50)
CD8+ effector T cells (46-50)
CD8+ naive T cells (46-50)
Early erythrocytes (46-50)
Early erythroid progenitors (46-50)
HSPCs (46-50)
Late erythrocytes (46-50)
Late erythroid progenitors (46-50)
Megakaryocytes (46-50)
Monocyte progenitors (46-50)
Natural killer cells (46-50)
Plasma cells (46-50)
Plasmacytoid dendritic cells (46-50)
Progenitor B cells (46-50)
Log-fold change of gene-level dyscoordination
Age
55-57
58-60
66-67
84

**Supplementary Figure 34 Distribution of gene-level dyscoordination in human bone marrow cells.**

Violin and box plots show the distribution of log-fold changes in dyscoordination relative to the baseline age group. Values are displayed separately for each cell type and age group, with the baseline age group indicated in parentheses in each subplot title. Values in the top and bottom 2.5% quantiles are considered outliers and are removed from the illustration. Red dashed line denotes zero log-fold change. Pairwise comparisons between age groups were tested using one-sided Wilcoxon rank-sum test (alternative hypothesis: older age group has higher dyscoordination). Significance levels are annotated above each comparison (NS = not significant, * $p < 0.05$, ** $p < 0.01$, *** $p < 0.001$)

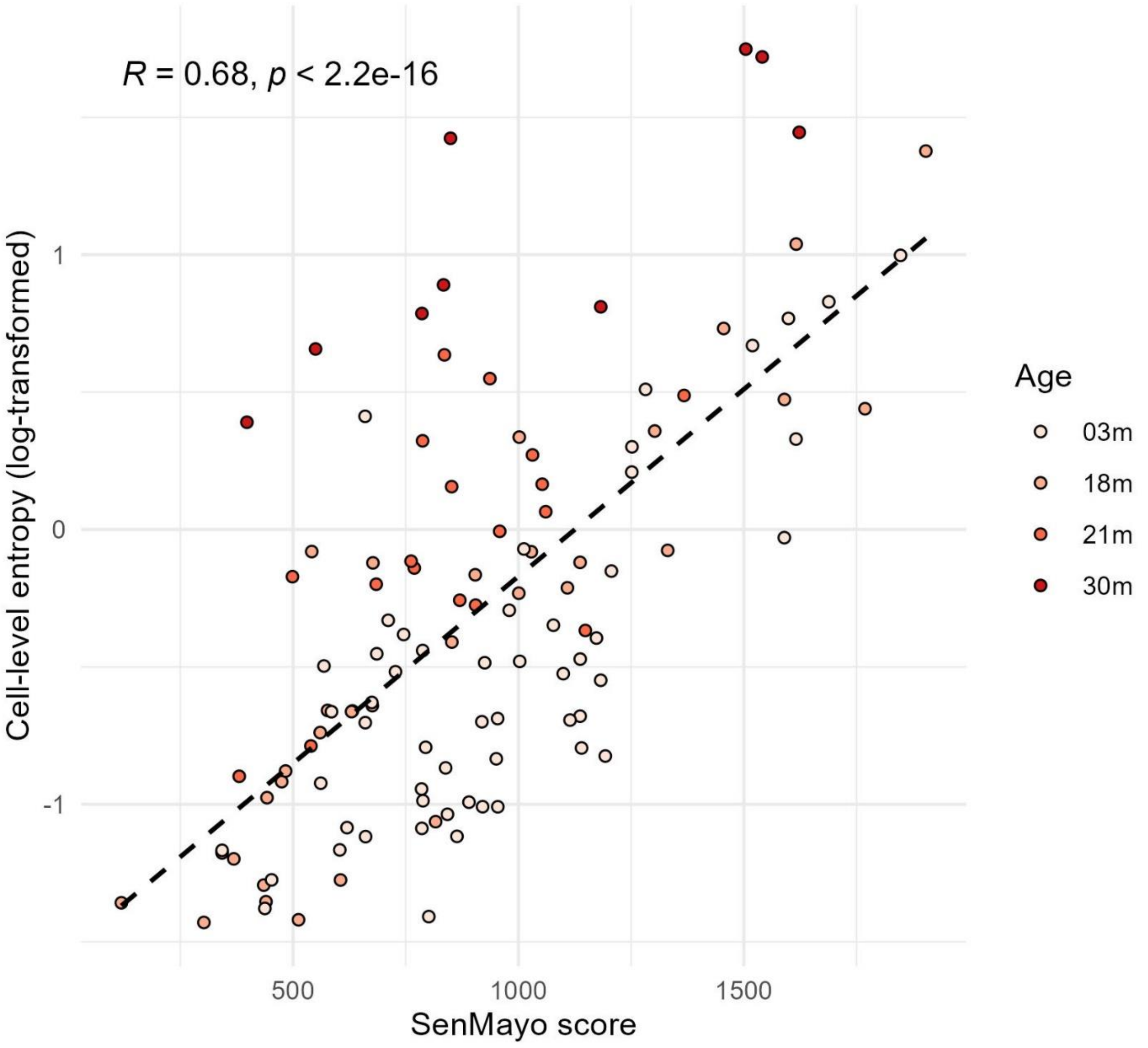


**Supplementary Figure 35 Correlation between SenMayo score and cell-level dyscoordination in type II pneumocytes (lung).**

Scatter plots show the relationship between SenMayo score and log-transformed cell-level dyscoordination across age groups. Black dashed lines indicate linear regression fits. Pearson correlation coefficients and corresponding p-values are shown in the top left corner.

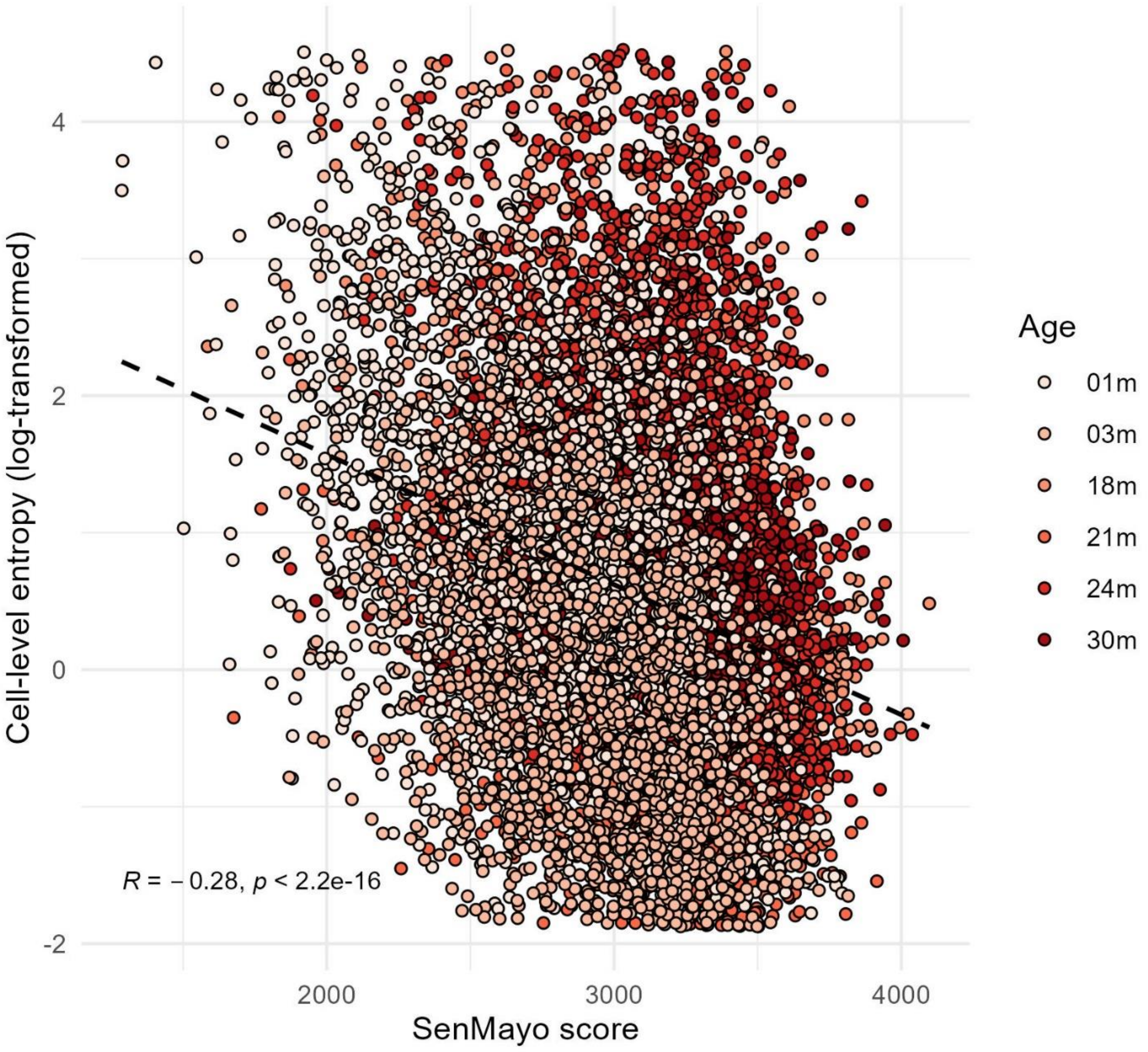


**Supplementary Figure 36 Correlation between SenMayo score and cell-level dyscoordination in mesenchymal stem cell (limb muscle).**

Scatter plots show the relationship between SenMayo score and log-transformed cell-level dyscoordination across age groups. Black dashed lines indicate linear regression fits. Pearson correlation coefficients and corresponding p-values are shown in the bottom left corner.

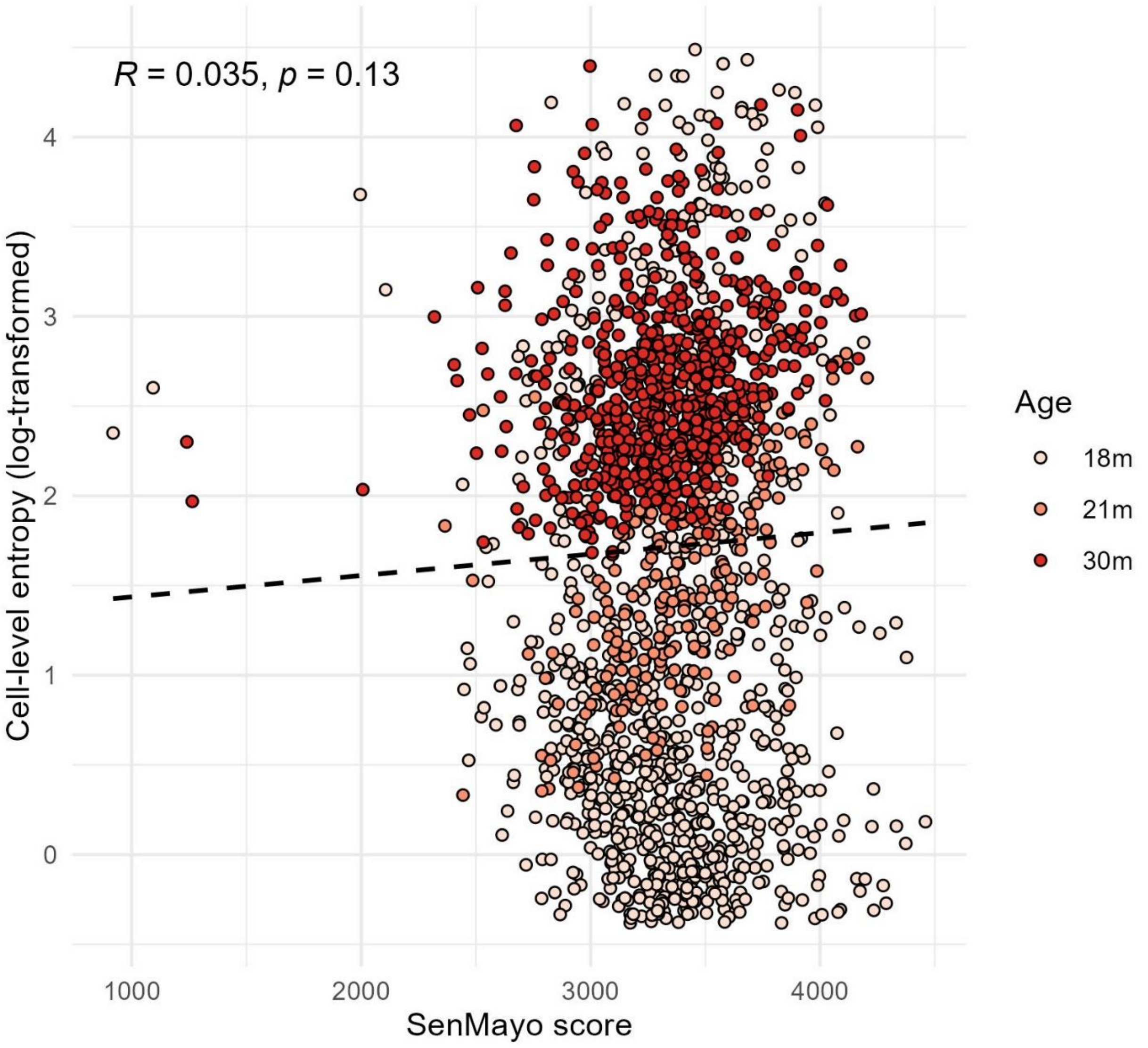


**Supplementary Figure 37 Correlation between SenMayo score and cell-level dyscoordination in mesenchymal stem cell (adipose tissue).**

Scatter plots show the relationship between SenMayo score and log-transformed cell-level dyscoordination across age groups. Black dashed lines indicate linear regression fits. Pearson correlation coefficients and corresponding p-values are shown in the top left corner.

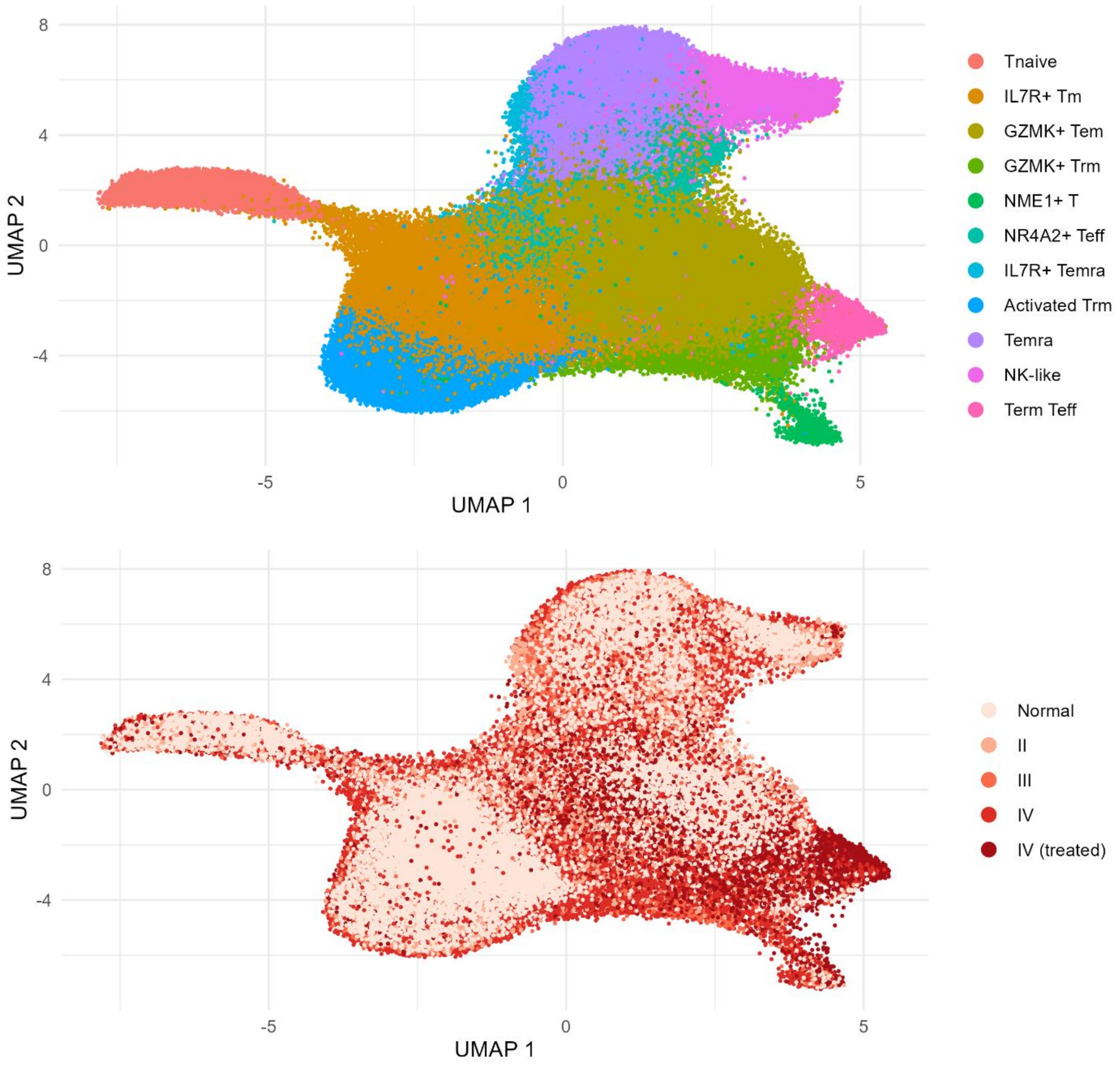


**Supplementary Figure 38 UMAP visualization of glioma CD8⁺ T cells (Wang et al. 2025).**

Cells are embedded using the first two UMAP dimensions and colored by annotated T cell subset (top) and tumor condition (bottom).

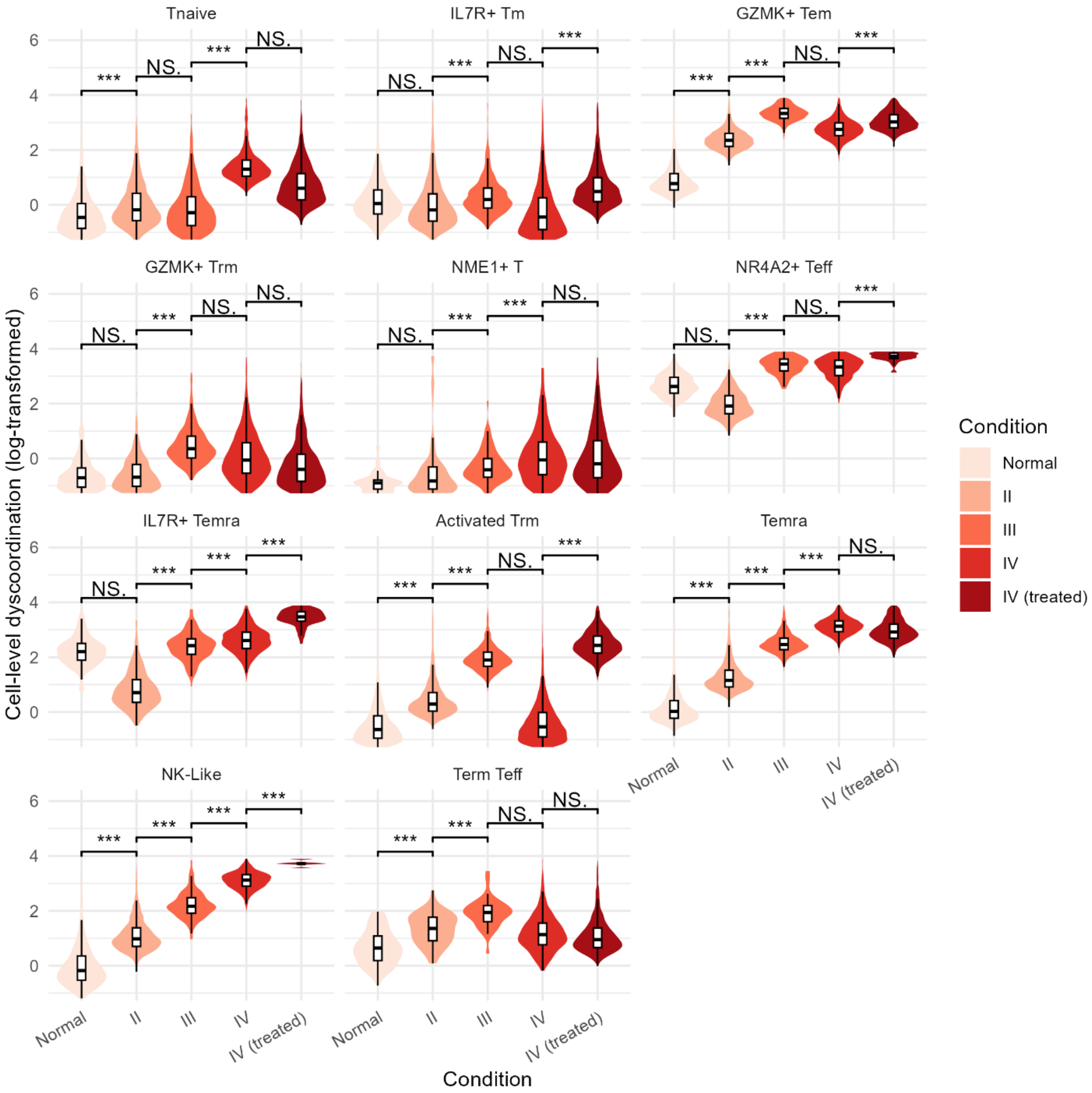


**Supplementary Figure 39 Distribution of cell-level dyscoordination in glioma CD8⁺ T cells.**

Violin and box plots show the distribution of log-transformed cell-level dyscoordination values separately for each subset and tumor condition. Values in the top and bottom 2.5% quantiles are considered outliers and are removed from the illustration. Pairwise comparisons between tumor conditions were tested using one-sided Wilcoxon rank-sum test (alternative hypothesis: more severe tumor condition has higher dyscoordination). Significance levels are annotated above each comparison (NS = not significant, * p < 0.05, ** p < 0.01, *** p < 0.001).

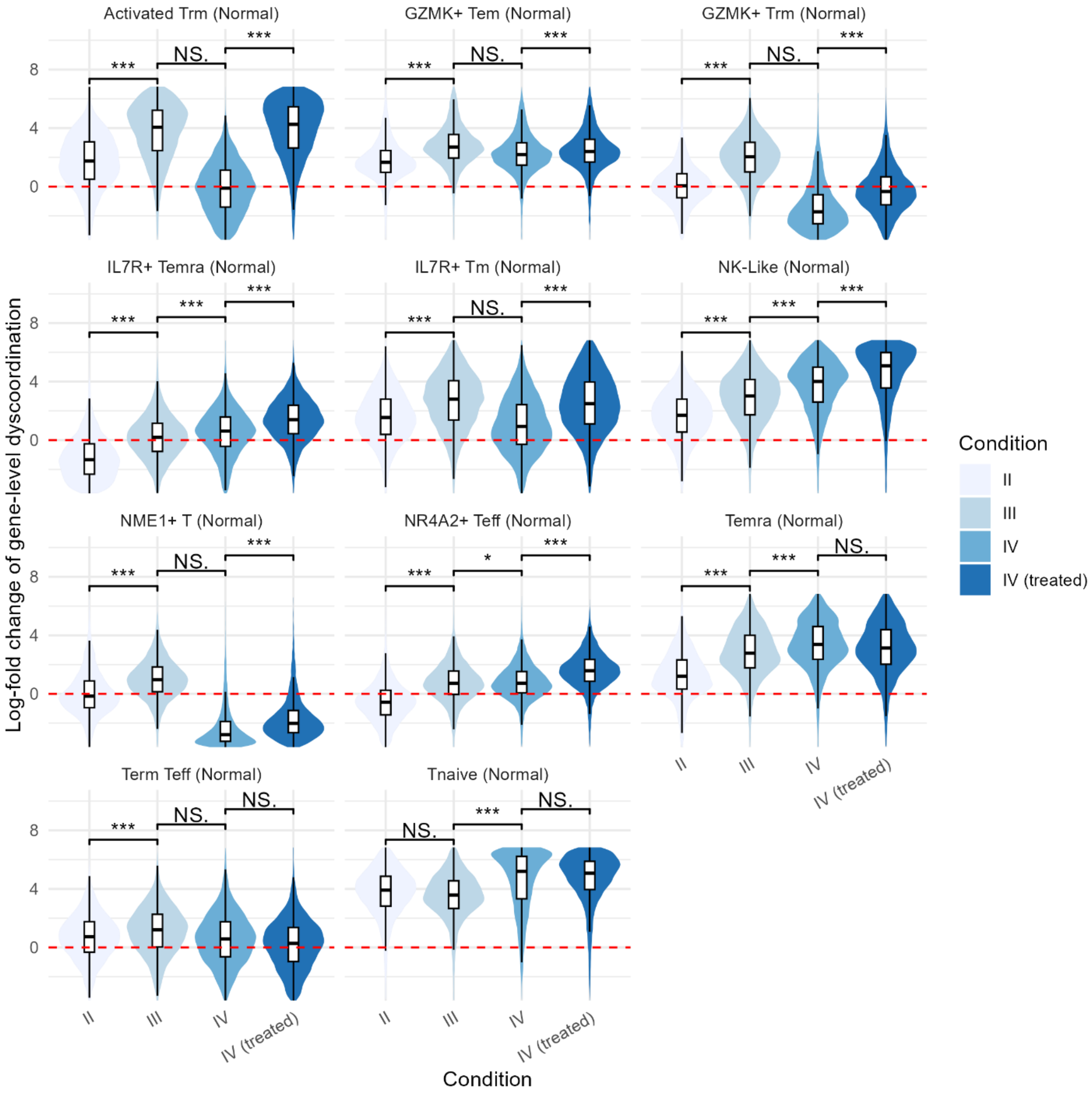

Activated Trm (Normal)
GZMK+ Tem (Normal)
GZMK+ Trm (Normal)
IL7R+ Temra (Normal)
IL7R+ Tm (Normal)
NK-Like (Normal)
NME1+ T (Normal)
NR4A2+ Teff (Normal)
Temra (Normal)
Term Teff (Normal)
Tnaive (Normal)
***
NS.
*
Log-fold change of gene-level dyscoordination
Condition
II
III
IV
IV (treated)

**Supplementary Figure 40 Distribution of gene-level dyscoordination in glioma $CD8^+$ T cells.**

Violin and box plots show the distribution of log-fold changes in dyscoordination relative to the baseline tumor condition. Values are displayed separately for each subset and tumor condition, with the baseline tumor condition indicated in parentheses in each subplot title. Values in the top and bottom 2.5% quantiles are considered outliers and are removed from the illustration. Red dashed line denotes zero log-fold change. Pairwise comparisons between tumor conditions were tested using one-sided Wilcoxon rank-sum test (alternative hypothesis: more severe tumor condition has higher dyscoordination). Significance levels are annotated above each comparison (NS = not significant, * $p < 0.05$, ** $p < 0.01$, *** $p < 0.001$).

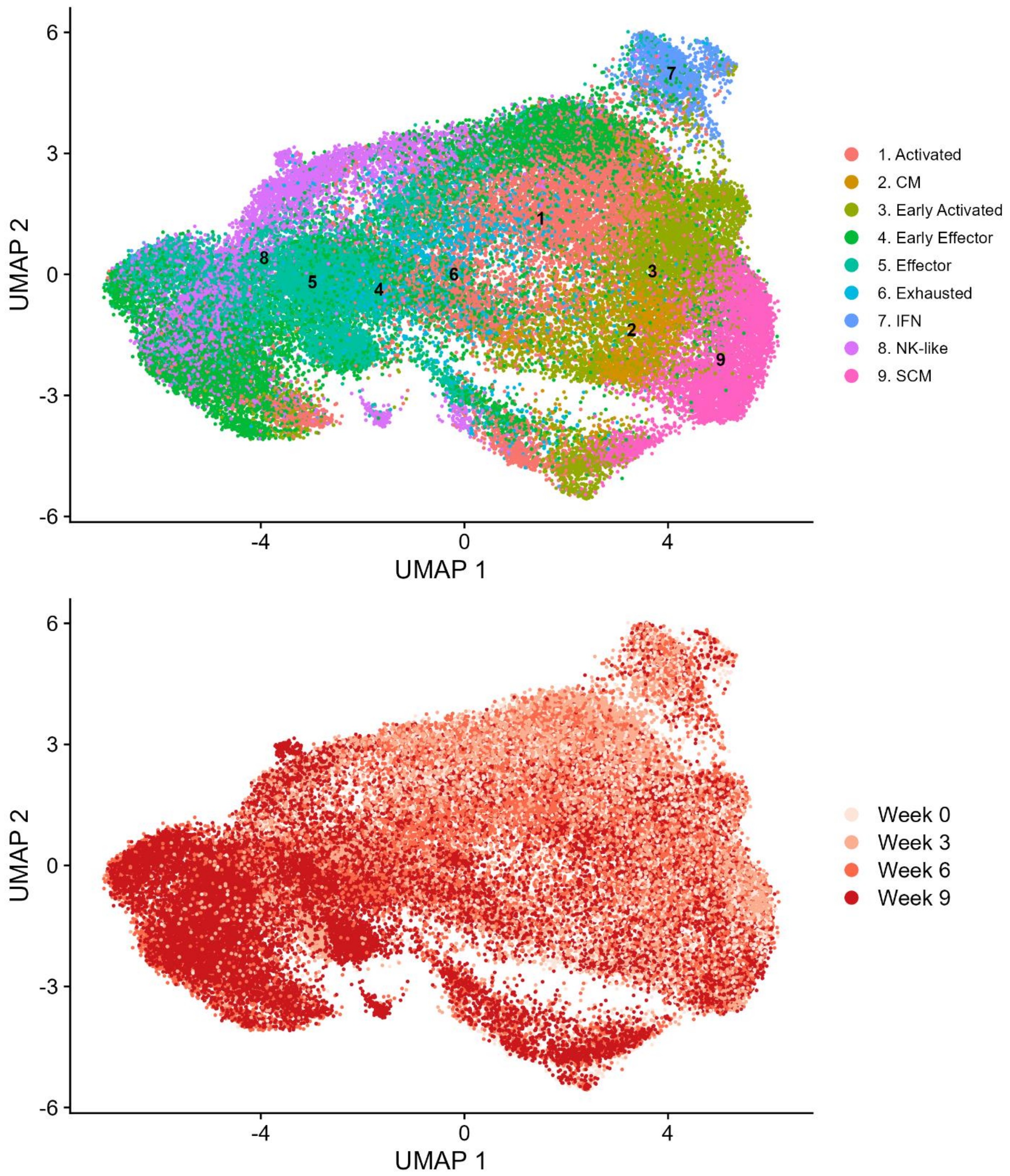


**Supplementary Figure 41 UMAP visualization of treated melanoma CD8+ T cells (Wang et al. 2024).**

Cells are embedded using the first two UMAP dimensions and colored by annotated T cell subset (top) and timepoint after combination therapy (bottom).

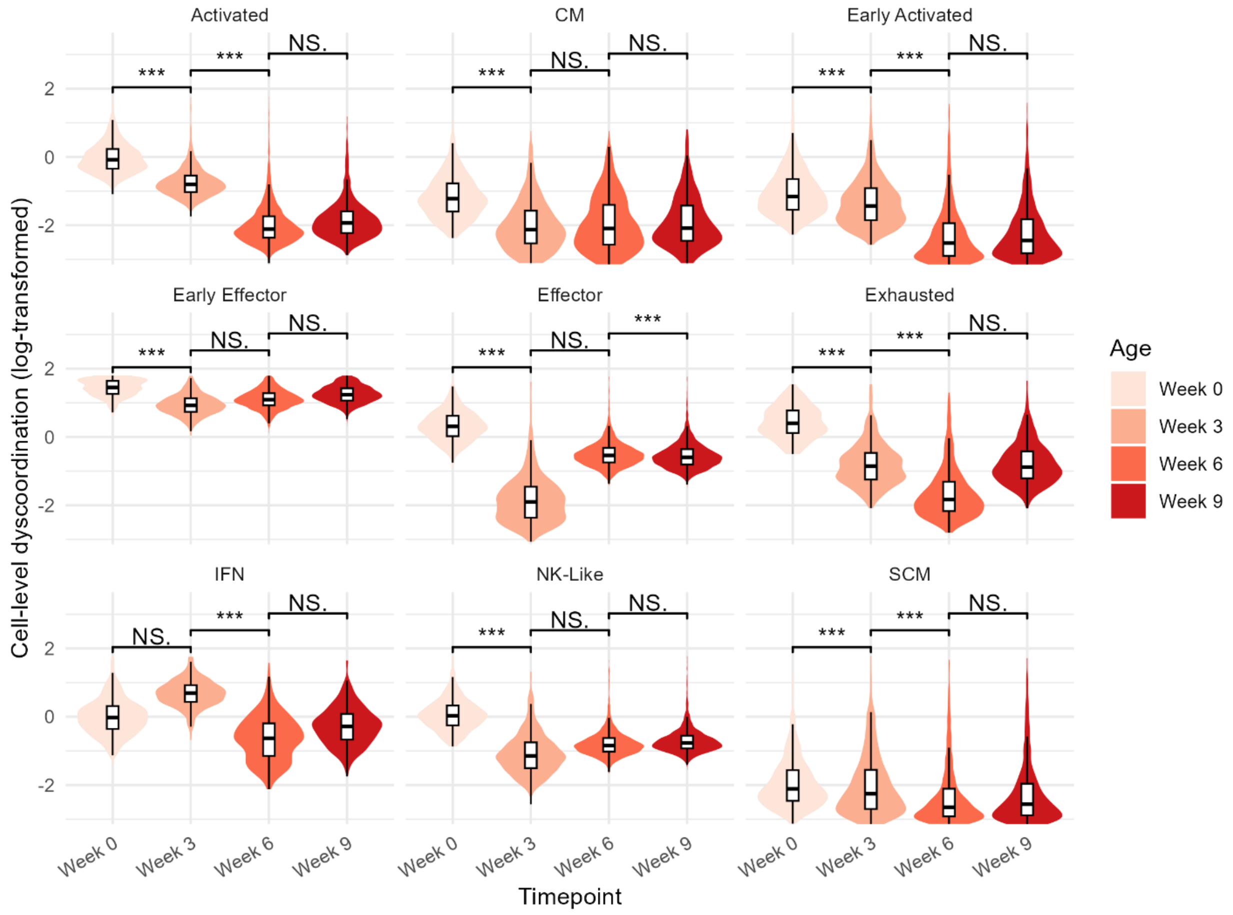


**Supplementary Figure 42 Distribution of cell-level dyscoordination in treated melanoma CD8⁺ T cells.**

Violin and box plots show the distribution of log-transformed cell-level dyscoordination values separately for each subset and timepoint after therapy. Values in the top and bottom 2.5% quantiles are considered outliers and are removed from the illustration. Pairwise comparisons between timepoints were tested using one-sided Wilcoxon rank-sum test (alternative hypothesis: later timepoint has lower dyscoordination). Significance levels are annotated above each comparison (NS = not significant, * $p < 0.05$, ** $p < 0.01$, *** $p < 0.001$).

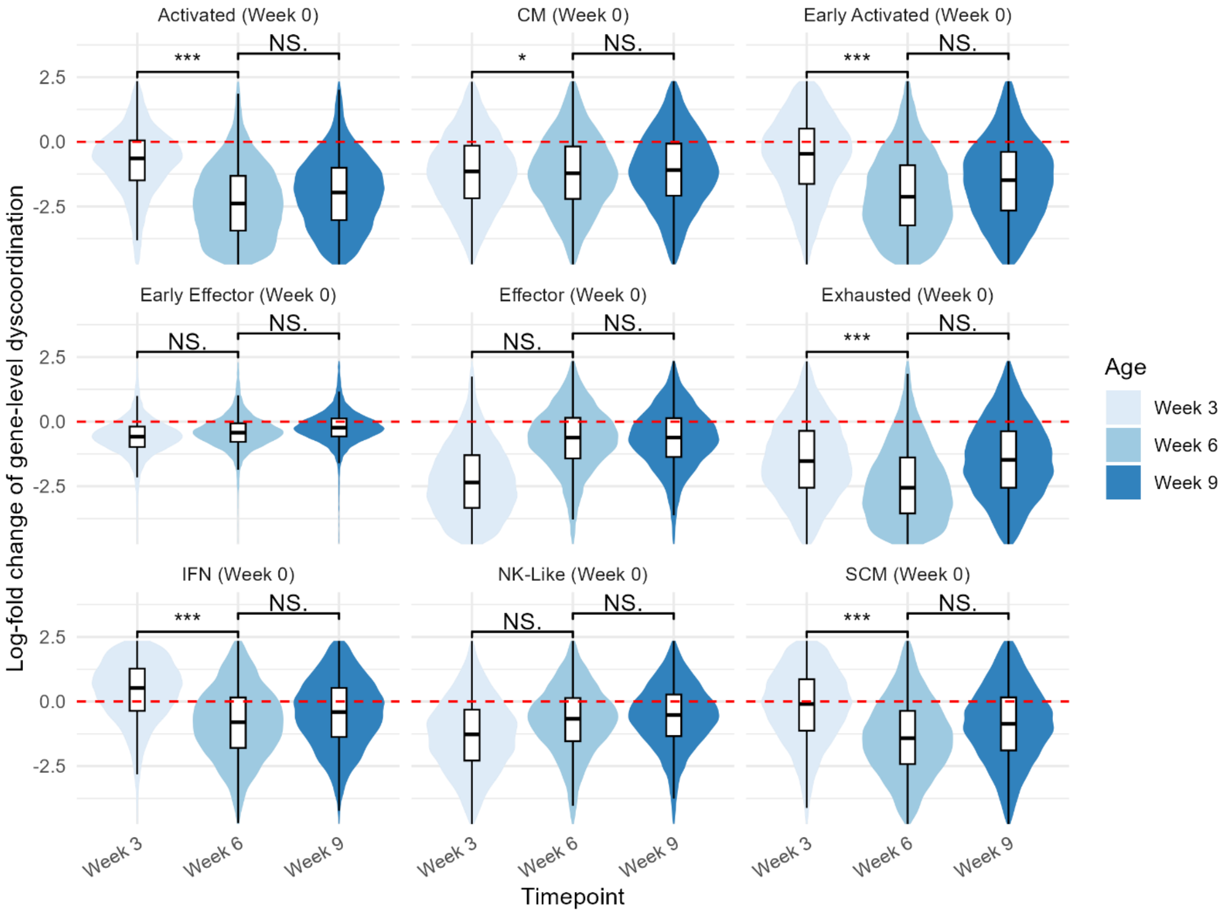


**Supplementary Figure 43 Distribution of gene-level dyscoordination in treated melanoma CD8$^+$ T cells.**

Violin and box plots show the distribution of log-fold changes in dyscoordination relative to the baseline timepoint after therapy. Values are displayed separately for each subset and timepoint after therapy, with the baseline timepoint after therapy indicated in parentheses in each subplot title. Values in the top and bottom 2.5% quantiles are considered outliers and are removed from the illustration. Red dashed line denotes zero log-fold change. Pairwise comparisons between timepoints were tested using one-sided Wilcoxon rank-sum test (alternative hypothesis: later timepoint has lower dyscoordination). Significance levels are annotated above each comparison (NS = not significant, * $p < 0.05$, ** $p < 0.01$, *** $p < 0.001$).

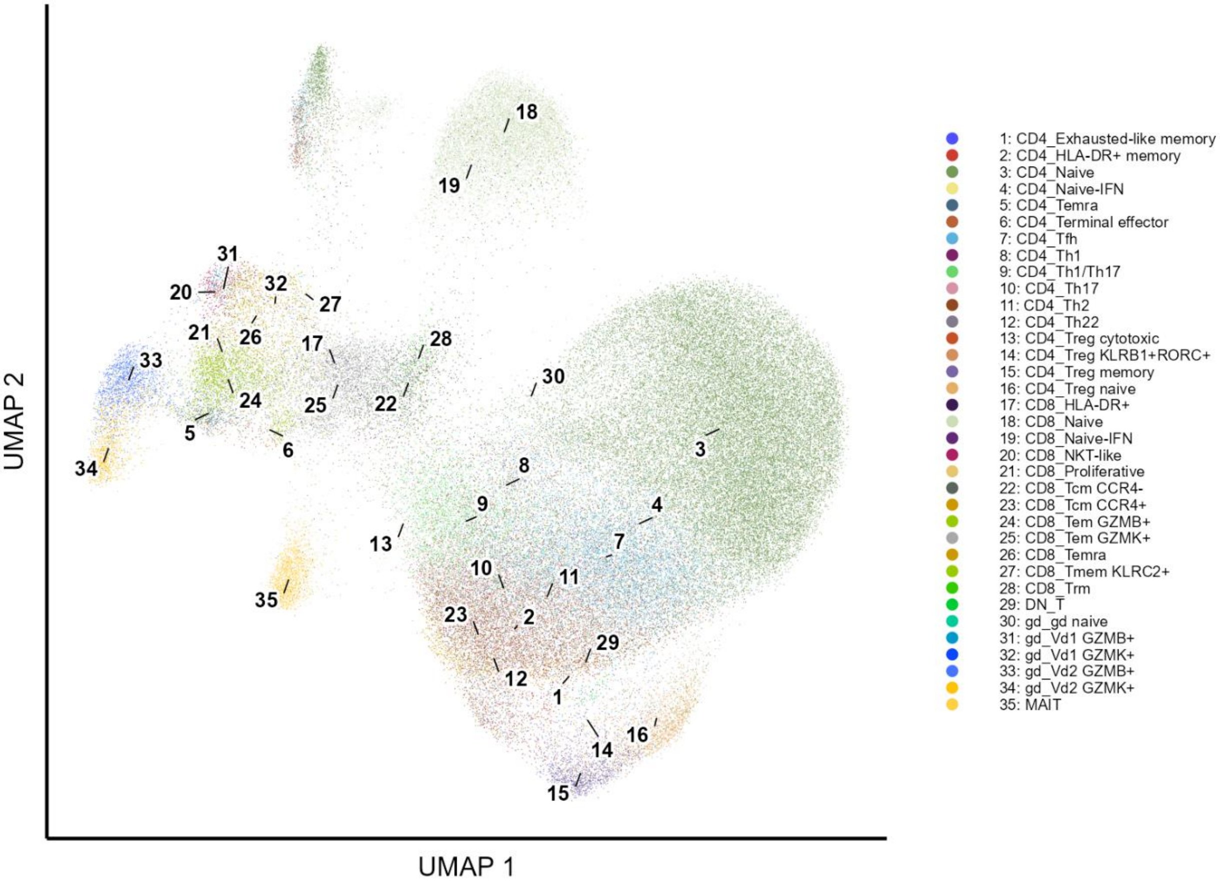
UMAP 2
UMAP 1
1: CD4_Exhausted-like memory
2: CD4_HLA-DR+ memory
3: CD4_Naive
4: CD4_Naive-IFN
5: CD4_Temra
6: CD4_Terminal effector
7: CD4_Tfh
8: CD4_Th1
9: CD4_Th1/Th17
10: CD4_Th17
11: CD4_Th2
12: CD4_Th22
13: CD4_Treg cytotoxic
14: CD4_Treg KLRB1+RORC+
15: CD4_Treg memory
16: CD4_Treg naive
17: CD8_HLA-DR+
18: CD8_Naive
19: CD8_Naive-IFN
20: CD8_NKT-like
21: CD8_Proliferative
22: CD8_Tcm CCR4-
23: CD8_Tcm CCR4+
24: CD8_Tem GZMB+
25: CD8_Tem GZMK+
26: CD8_Temra
27: CD8_Tmem KLRC2+
28: CD8_Trm
29: DN_T
30: gd_gd naive
31: gd_Vd1 GZMB+
32: gd_Vd1 GZMK+
33: gd_Vd2 GZMB+
34: gd_Vd2 GZMK+
35: MAIT

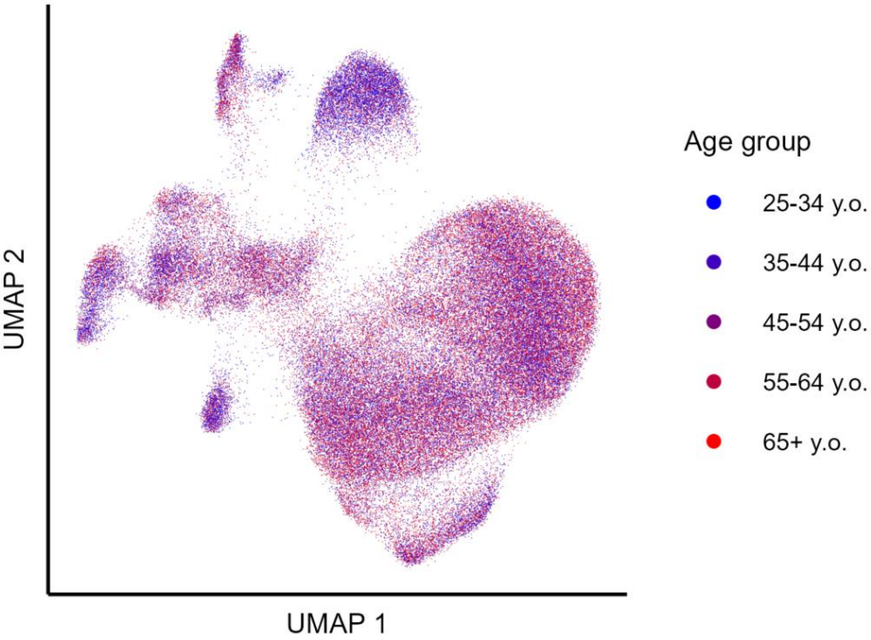
UMAP 2
UMAP 1
Age group
25-34 y.o.
35-44 y.o.
45-54 y.o.
55-64 y.o.
65+ y.o.

**Supplementary Figure 44 UMAP visualization of aging human blood T cells (Terekhova et al. 2023).**

Cells are embedded using the first two UMAP dimensions and colored by annotated T cell subset (top) and age group (bottom).

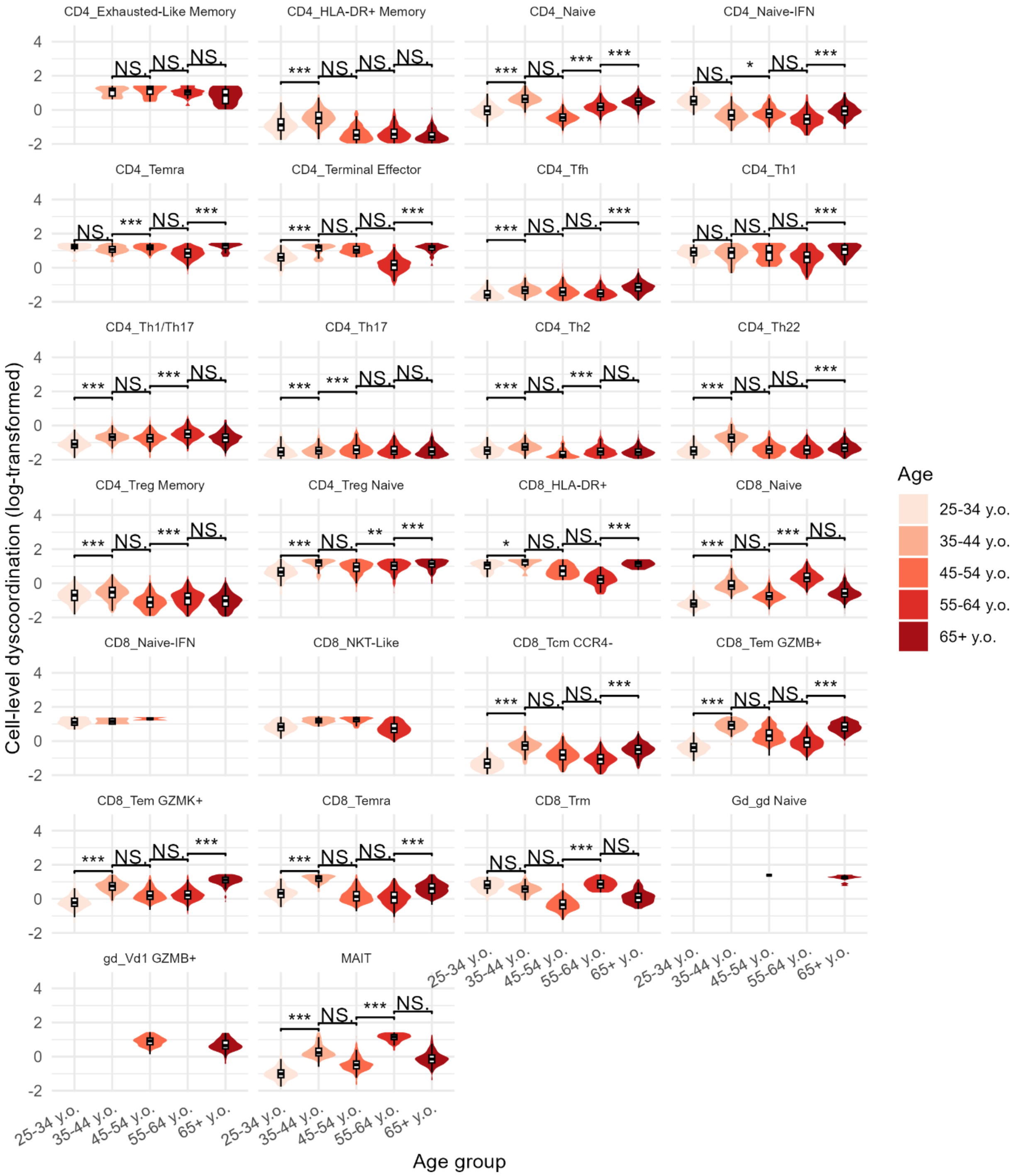
CD4_Exhausted-Like Memory
CD4_HLA-DR+ Memory
CD4_Naive
CD4_Naive-IFN
CD4_Temra
CD4_Terminal Effector
CD4_Tfh
CD4_Th1
CD4_Th1/Th17
CD4_Th17
CD4_Th2
CD4_Th22
CD4_Treg Memory
CD4_Treg Naive
CD8_HLA-DR+
CD8_Naive
CD8_Naive-IFN
CD8_NKT-Like
CD8_Tcm CCR4-
CD8_Tem GZMB+
CD8_Tem GZMK+
CD8_Temra
CD8_Trm
Gd_gd Naive
gd_Vd1 GZMB+
MAIT
Cell-level dyscoordination (log-transformed)
Age group
25-34 y.o.
35-44 y.o.
45-54 y.o.
55-64 y.o.
65+ y.o.
Age

**Supplementary Figure 45 Distribution of cell-level dyscoordination in aging human T cells (Terekhova et al. 2023).**

Violin and box plots show the distribution of log-transformed cell-level dyscoordination values separately for each subset and age group. Values in the top and bottom 2.5% quantiles are considered outliers and are removed from the illustration. Pairwise comparisons between age groups were tested using one-sided Wilcoxon rank-sum test (alternative hypothesis: older age group has higher dyscoordination). Significance levels are annotated above each comparison (NS = not significant, * $p < 0.05$, ** $p < 0.01$, *** $p < 0.001$).

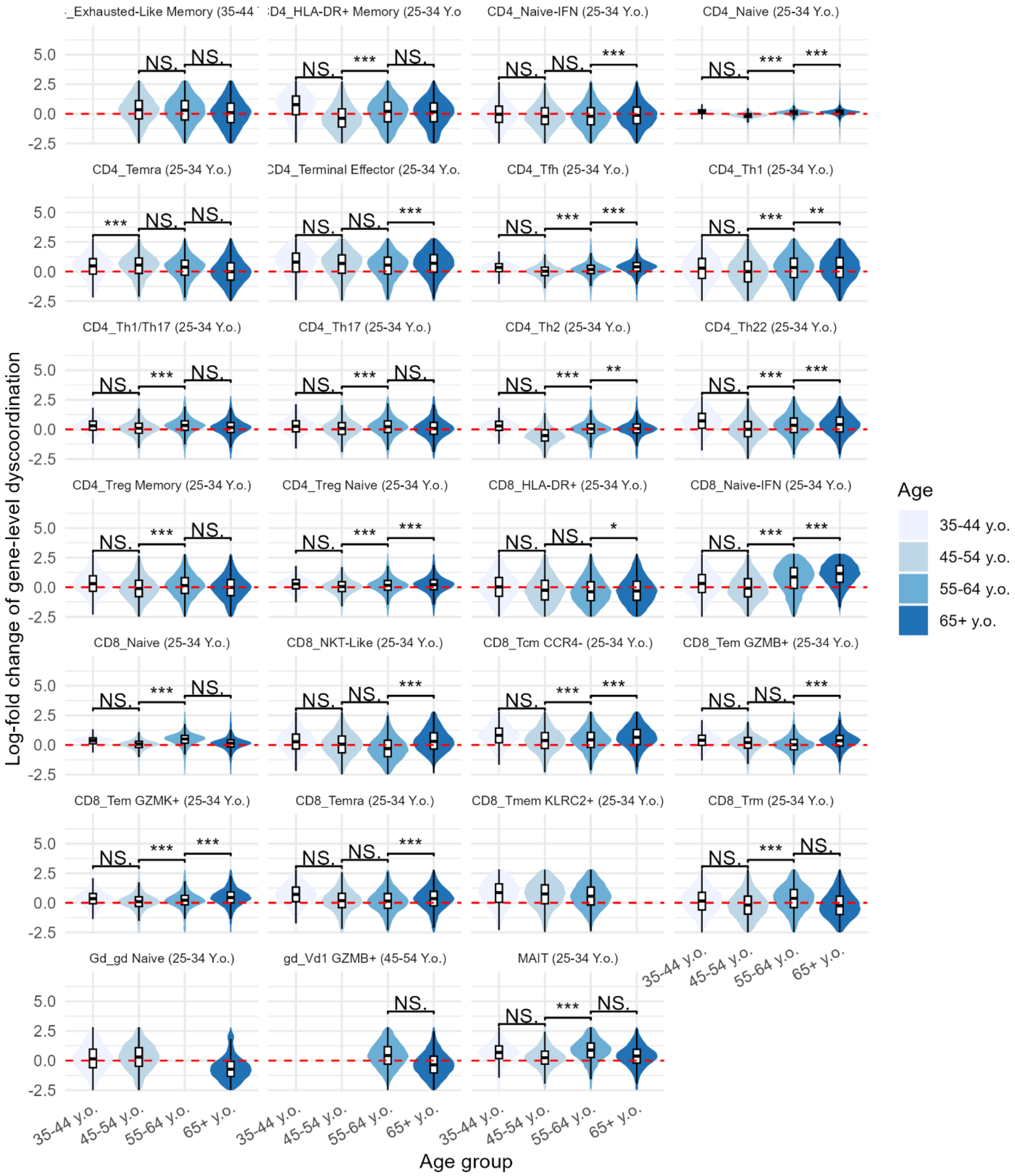

_Exhausted-Like Memory (35-44 Y
D4_HLA-DR+ Memory (25-34 Y.o
CD4_Naive-IFN (25-34 Y.o.)
CD4_Naive (25-34 Y.o.)
CD4_Temra (25-34 Y.o.)
D4_Terminal Effector (25-34 Y.o.
CD4_Tfh (25-34 Y.o.)
CD4_Th1 (25-34 Y.o.)
CD4_Th1/Th17 (25-34 Y.o.)
CD4_Th17 (25-34 Y.o.)
CD4_Th2 (25-34 Y.o.)
CD4_Th22 (25-34 Y.o.)
CD4_Treg Memory (25-34 Y.o.)
CD4_Treg Naive (25-34 Y.o.)
CD8_HLA-DR+ (25-34 Y.o.)
CD8_Naive-IFN (25-34 Y.o.)
CD8_Naive (25-34 Y.o.)
CD8_NKT-Like (25-34 Y.o.)
CD8_Tcm CCR4- (25-34 Y.o.)
CD8_Tem GZMB+ (25-34 Y.o.)
CD8_Tem GZMK+ (25-34 Y.o.)
CD8_Temra (25-34 Y.o.)
CD8_Tmem KLRC2+ (25-34 Y.o.)
CD8_Trm (25-34 Y.o.)
Gd_gd Naive (25-34 Y.o.)
gd_Vd1 GZMB+ (45-54 Y.o.)
MAIT (25-34 Y.o.)
NS.
***
**
*
5.0
2.5
0.0
-2.5
Log-fold change of gene-level dyscoordination
35-44 y.o.
45-54 y.o.
55-64 y.o.
65+ y.o.
Age group
Age

**Supplementary Figure 46 Distribution of gene-level dyscoordination in aging human T cells (Terekhova et al. 2023).**

Violin and box plots show the distribution of log-fold changes in dyscoordination relative to the baseline age group. Values are displayed separately for each subset and age group, with the baseline age group indicated in parentheses in each subplot title. Values in the top and bottom 2.5% quantiles are considered outliers and are removed from the illustration. Red dashed line denotes zero log-fold change. Pairwise comparisons between age groups were tested using one-sided Wilcoxon rank-sum test (alternative hypothesis: older age group has higher dyscoordination). Significance levels are annotated above each comparison (NS = not significant, * $p < 0.05$, ** $p < 0.01$, *** $p < 0.001$).

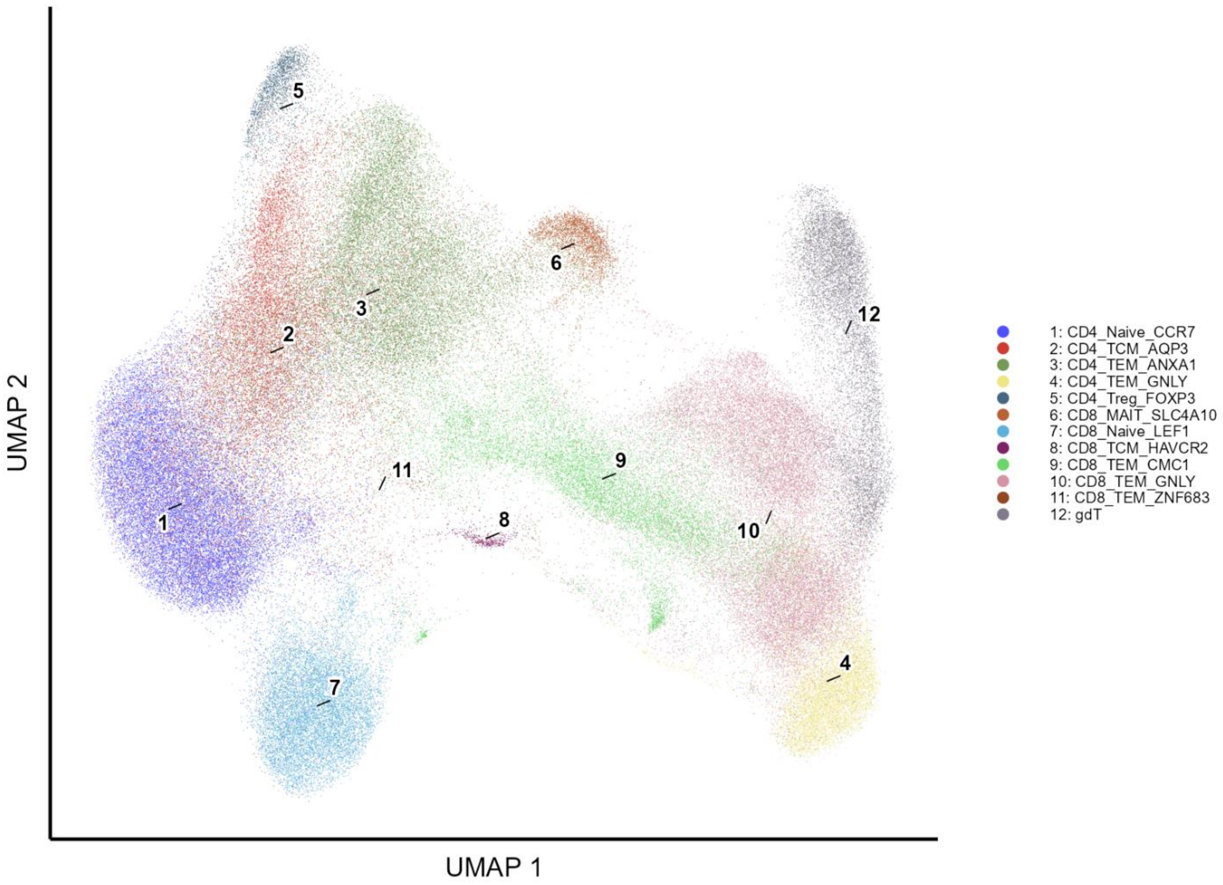
5
6
3
2
12
11
9
1
8
10
7
4
UMAP 2
UMAP 1
1: CD4_Naive_CCR7
2: CD4_TCM_AQP3
3: CD4_TEM_ANXA1
4: CD4_TEM_GNLY
5: CD4_Treg_FOXP3
6: CD8_MAIT_SLC4A10
7: CD8_Naive_LEF1
8: CD8_TCM_HAVCR2
9: CD8_TEM_CMC1
10: CD8_TEM_GNLY
11: CD8_TEM_ZNF683
12: gdT

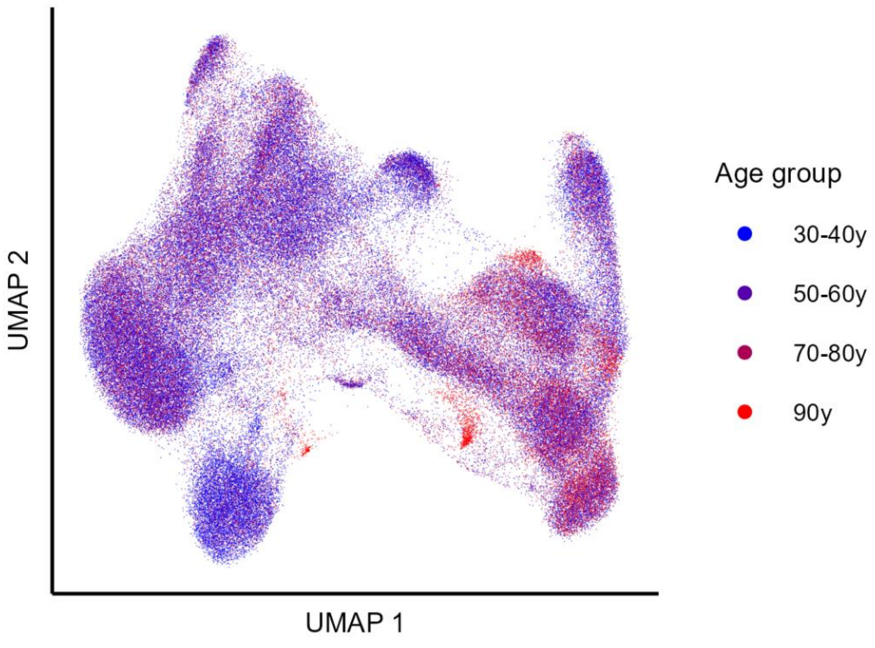
UMAP 2
UMAP 1
Age group
30-40y
50-60y
70-80y
90y

**Supplementary Figure 47 UMAP visualization of aging human blood T cells (Wang et al. 2025).**

Cells are embedded using the first two UMAP dimensions and colored by annotated T cell subset (top) and age group (bottom).

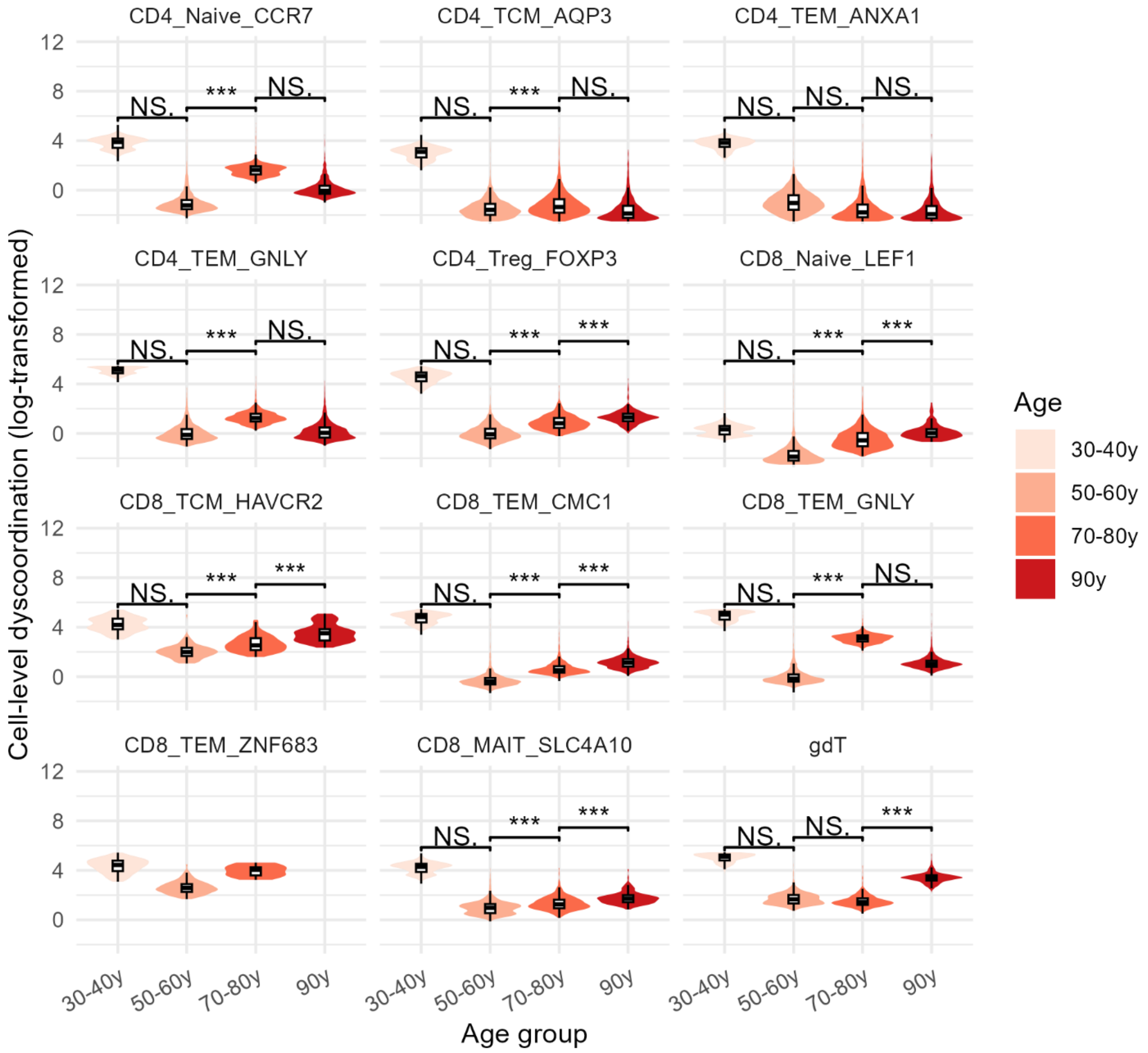


**Supplementary Figure 48 Distribution of cell-level dyscoordination in aging human T cells (Wang et al. 2025).**

Violin and box plots show the distribution of log-transformed cell-level dyscoordination values separately for each subset and age group. Values in the top and bottom 2.5% quantiles are considered outliers and are removed from the illustration. Pairwise comparisons between age groups were tested using one-sided Wilcoxon rank-sum test (alternative hypothesis: older age group has higher dyscoordination). Significance levels are annotated above each comparison (NS = not significant, * $p < 0.05$, ** $p < 0.01$, *** $p < 0.001$).

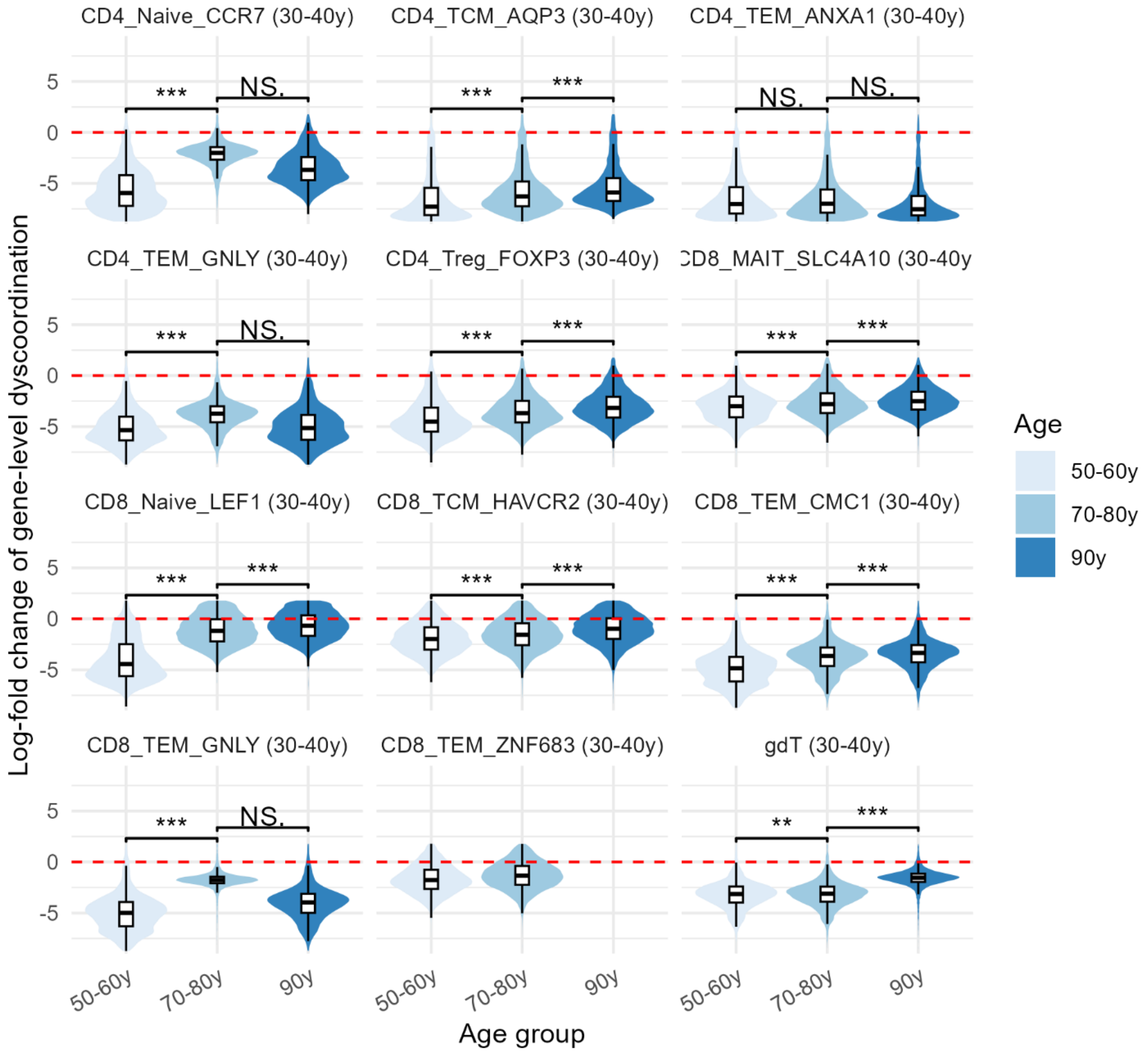


**Supplementary Figure 49 Distribution of gene-level dyscoordination in aging human T cells (Wang et al. 2025).**

Violin and box plots show the distribution of log-fold changes in dyscoordination relative to the baseline age group. Values are displayed separately for each subset and age group, with the baseline age group indicated in parentheses in each subplot title. Values in the top and bottom 2.5% quantiles are considered outliers and are removed from the illustration. Red dashed line denotes zero log-fold change. Pairwise comparisons between age groups were tested using one-sided Wilcoxon rank-sum test (alternative hypothesis: older age group has higher dyscoordination). Significance levels are annotated above each comparison (NS = not significant, * $p < 0.05$, ** $p < 0.01$, *** $p < 0.001$).

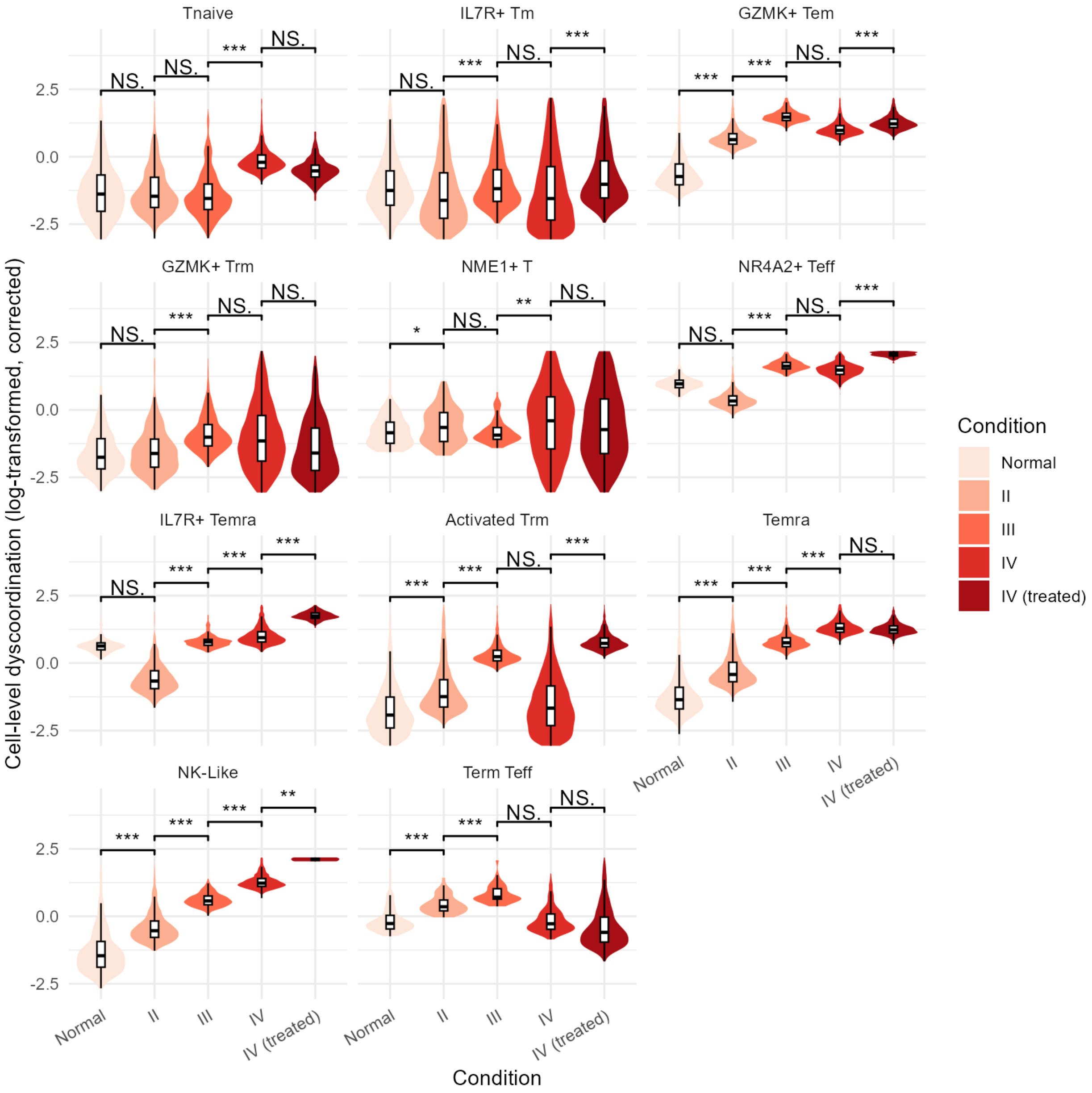


**Supplementary Figure 50 Distribution of corrected cell-level dyscoordination in glioma CD8+ T cells.**

Violin and box plots show the distribution of log-transformed, depth-corrected cell-level dyscoordination values separately for each subset and tumor condition. Values in the top and bottom 2.5% quantiles are considered outliers and are removed from the illustration. Pairwise comparisons between tumor conditions were tested using one-sided Wilcoxon rank-sum test (alternative hypothesis: more severe tumor condition has higher dyscoordination). Significance levels are annotated above each comparison (NS = not significant, * $p < 0.05$, ** $p < 0.01$, *** $p < 0.001$).

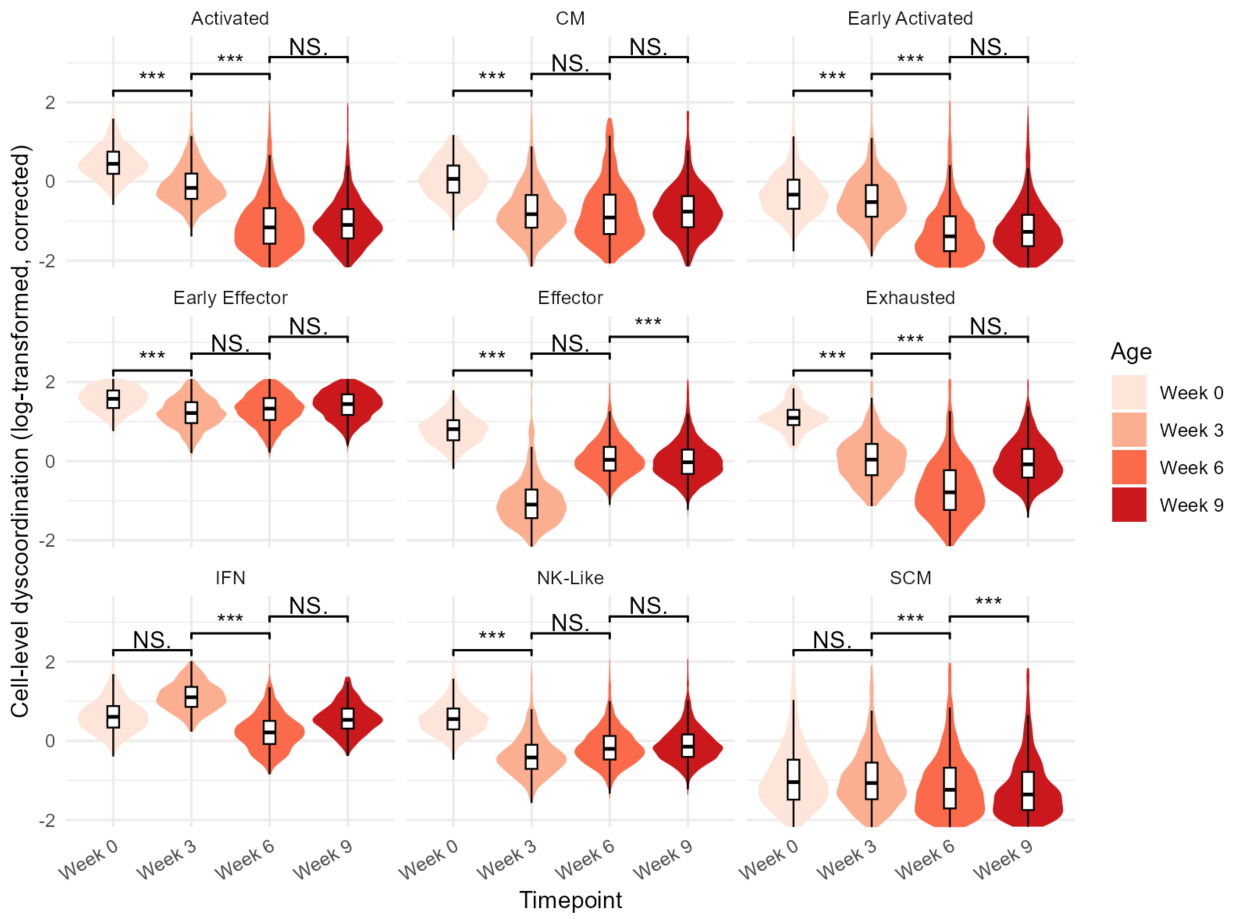


**Supplementary Figure 51 Distribution of corrected cell-level dyscoordination in treated melanoma CD8+ T cells.**

Violin and box plots show the distribution of log-transformed, depth-corrected cell-level dyscoordination values separately for each subset and timepoint after therapy. Values in the top and bottom 2.5% quantiles are considered outliers and are removed from the illustration. Pairwise comparisons between timepoints were tested using one-sided Wilcoxon rank-sum test (alternative hypothesis: later timepoint has lower dyscoordination). Significance levels are annotated above each comparison (NS = not significant, * $p < 0.05$, ** $p < 0.01$, *** $p < 0.001$).

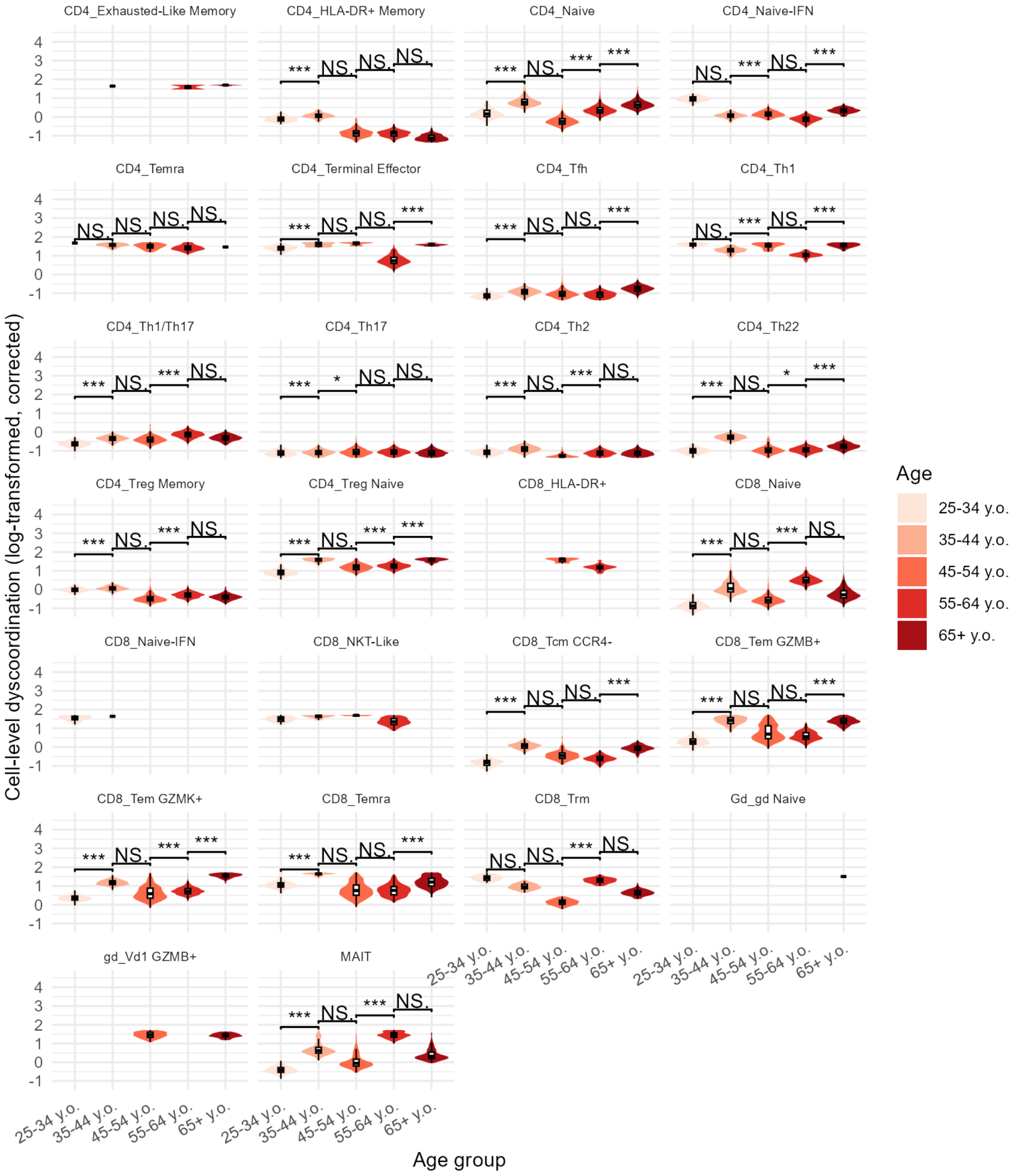

CD4_Exhausted-Like Memory
CD4_HLA-DR+ Memory
CD4_Naive
CD4_Naive-IFN
CD4_Temra
CD4_Terminal Effector
CD4_Tfh
CD4_Th1
CD4_Th1/Th17
CD4_Th17
CD4_Th2
CD4_Th22
CD4_Treg Memory
CD4_Treg Naive
CD8_HLA-DR+
CD8_Naive
CD8_Naive-IFN
CD8_NKT-Like
CD8_Tcm CCR4-
CD8_Tem GZMB+
CD8_Tem GZMK+
CD8_Temra
CD8_Trm
Gd_gd Naive
gd_Vd1 GZMB+
MAIT
Cell-level dyscoordination (log-transformed, corrected)
Age group
25-34 y.o.
35-44 y.o.
45-54 y.o.
55-64 y.o.
65+ y.o.
Age

**Supplementary Figure 52 Distribution of corrected cell-level dyscoordination in aging human T cells (Terekhova et al. 2023).**

Violin and box plots show the distribution of log-transformed, depth-corrected cell-level dyscoordination values separately for each subset and age group. Values in the top and bottom 2.5% quantiles are considered outliers and are removed from the illustration. Pairwise comparisons between age groups were tested using one-sided Wilcoxon rank-sum test (alternative hypothesis: older age group has higher dyscoordination). Significance levels are annotated above each comparison (NS = not significant, * $p < 0.05$, ** $p < 0.01$, *** $p < 0.001$).

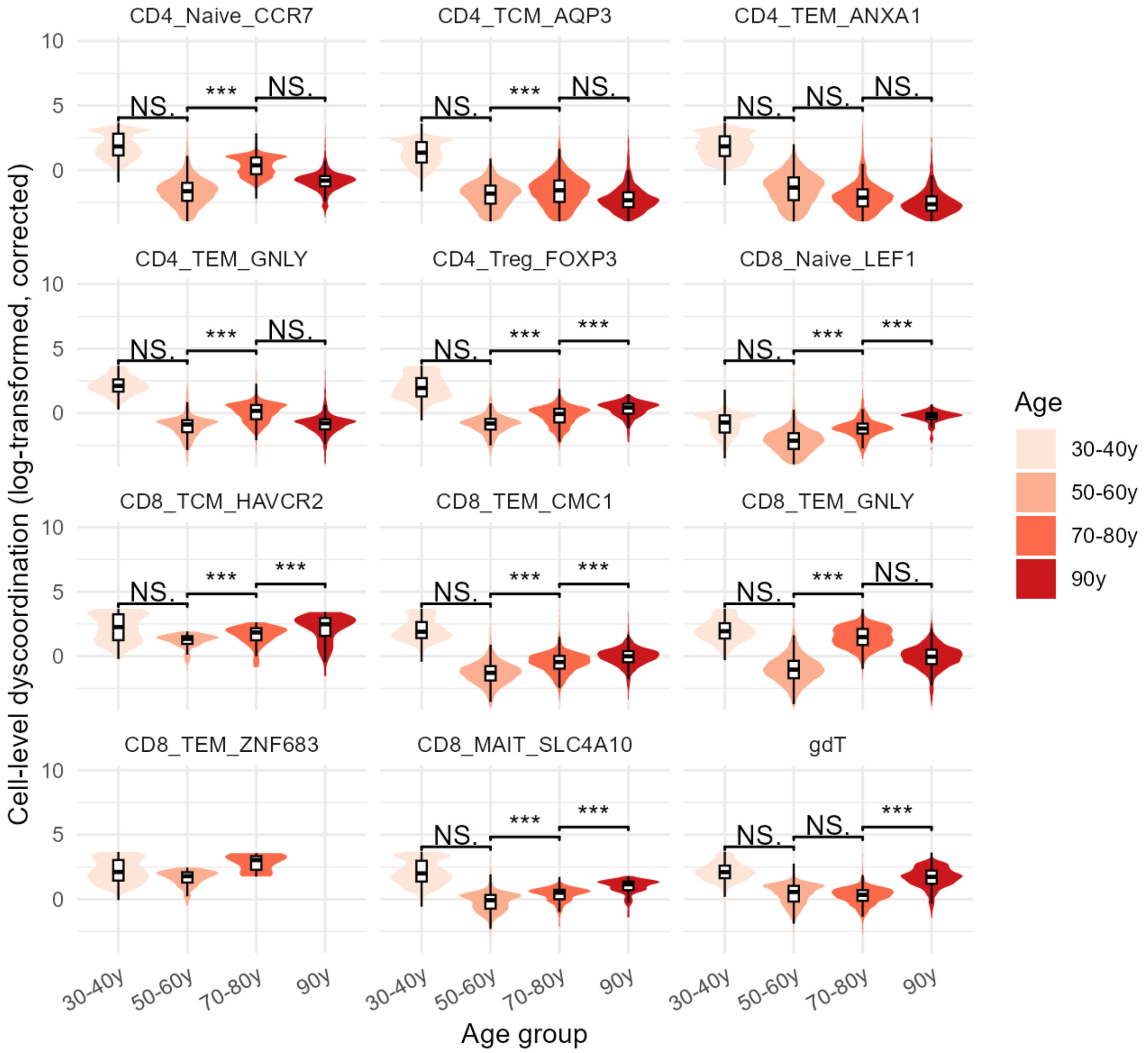


**Supplementary Figure 53 Distribution of corrected cell-level dyscoordination in aging human T cells (Wang et al. 2025).**

Violin and box plots show the distribution of log-transformed, depth-corrected cell-level dyscoordination values separately for each subset and age group. Values in the top and bottom 2.5% quantiles are considered outliers and are removed from the illustration. Pairwise comparisons between age groups were tested using one-sided Wilcoxon rank-sum test (alternative hypothesis: older age group has higher dyscoordination). Significance levels are annotated above each comparison (NS = not significant, * $p < 0.05$, ** $p < 0.01$, *** $p < 0.001$).

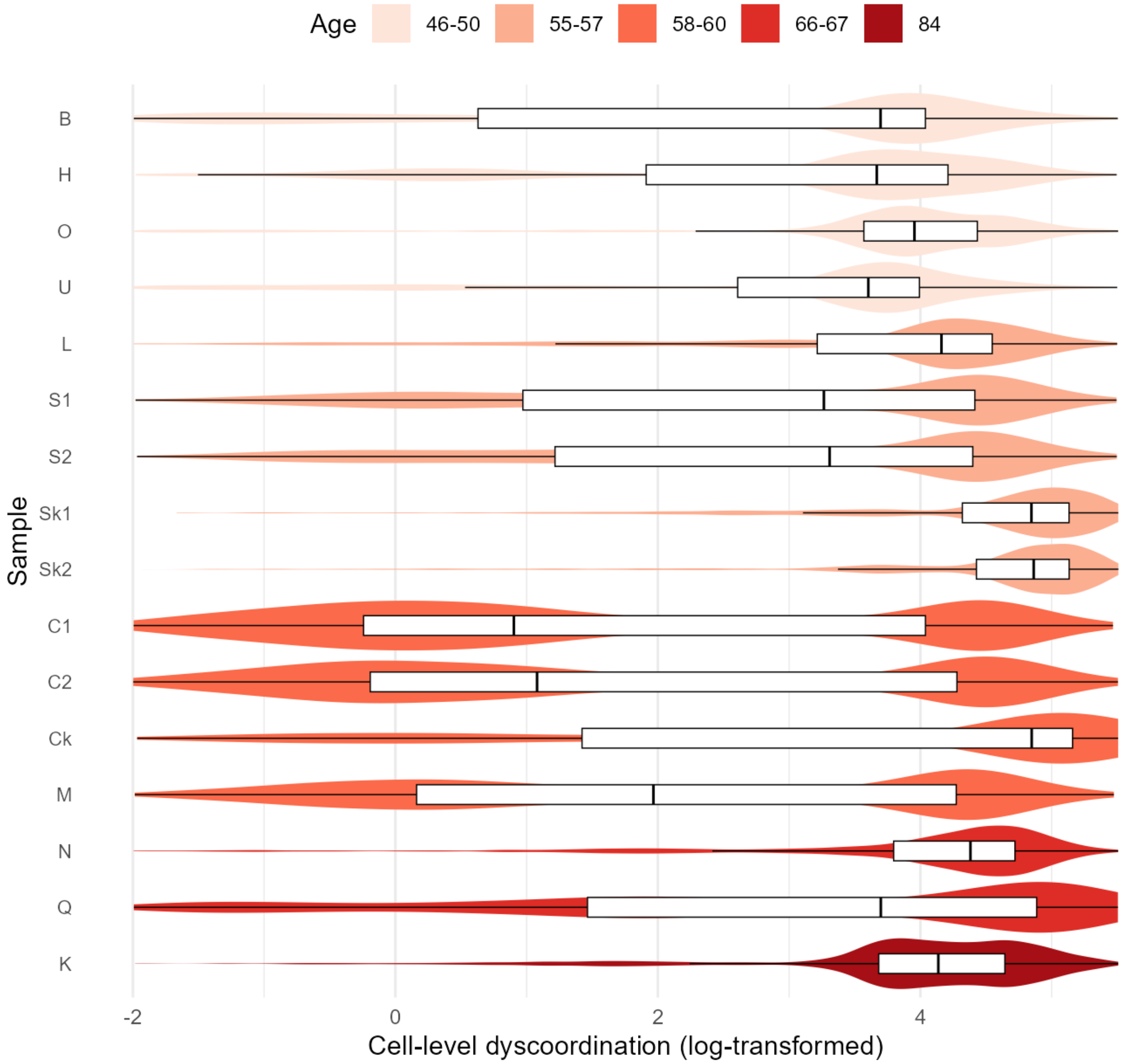


**Supplementary Figure 54 Distribution of cell-level dyscoordination across biological replicates in glioma CD8+ T cells.**

Violin and box plots show the distribution of log-transformed cell-level dyscoordination for each individual patient, ordered and colored by tumor condition. Values in the top and bottom 2.5% quantiles are considered outliers and are removed from the illustration.

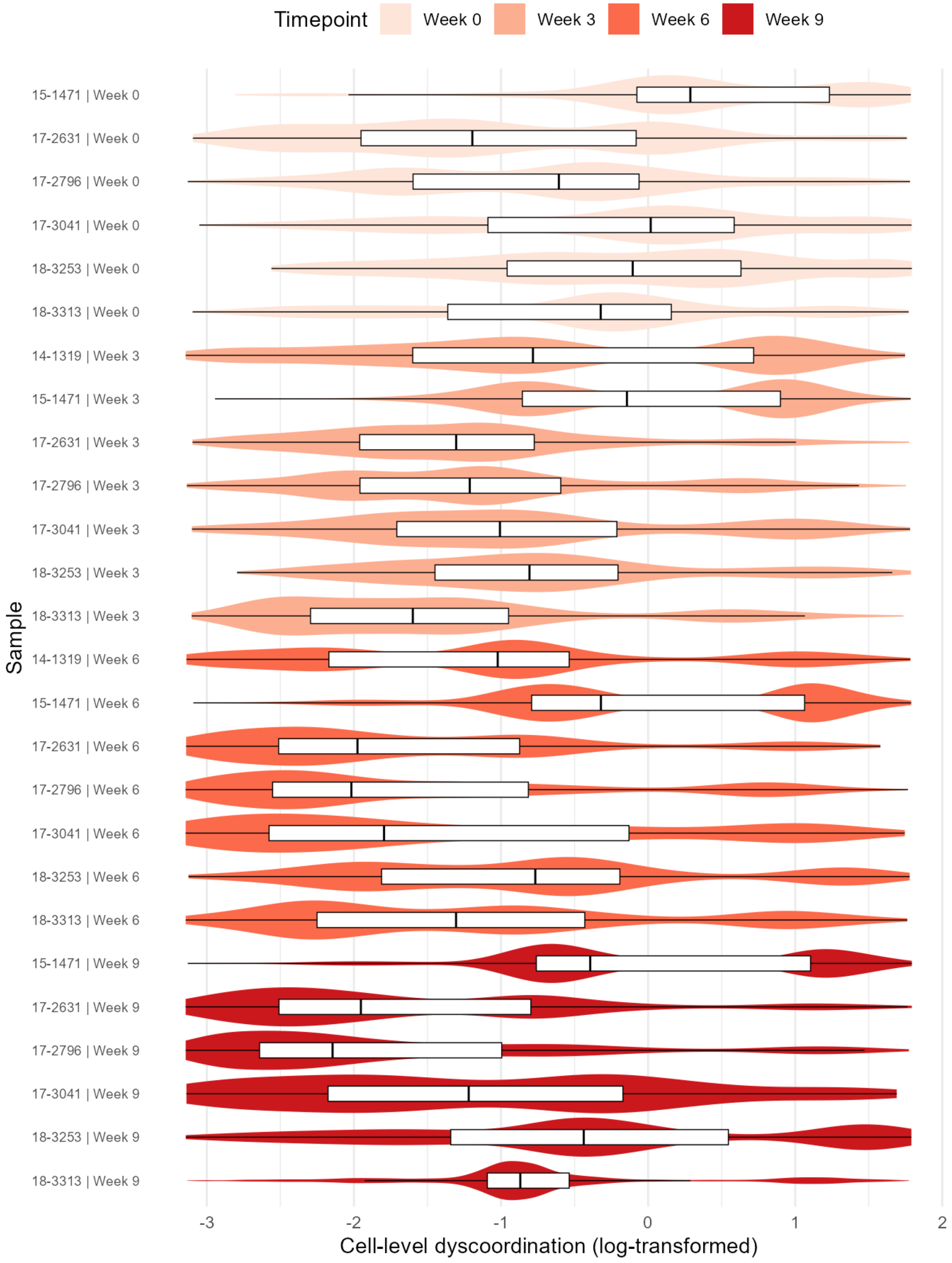
Timepoint
Week 0
Week 3
Week 6
Week 9
Sample
15-1471 | Week 0
17-2631 | Week 0
17-2796 | Week 0
17-3041 | Week 0
18-3253 | Week 0
18-3313 | Week 0
14-1319 | Week 3
15-1471 | Week 3
17-2631 | Week 3
17-2796 | Week 3
17-3041 | Week 3
18-3253 | Week 3
18-3313 | Week 3
14-1319 | Week 6
15-1471 | Week 6
17-2631 | Week 6
17-2796 | Week 6
17-3041 | Week 6
18-3253 | Week 6
18-3313 | Week 6
15-1471 | Week 9
17-2631 | Week 9
17-2796 | Week 9
17-3041 | Week 9
18-3253 | Week 9
18-3313 | Week 9
-3
-2
-1
0
1
2
Cell-level dyscoordination (log-transformed)

**Supplementary Figure 55 Distribution of cell-level dyscoordination across biological replicates in treated melanoma CD8+ T cells.**

Violin and box plots show the distribution of log-transformed cell-level dyscoordination for each individual patient and timepoint, ordered and colored by timepoint after therapy. Values in the top and bottom 2.5% quantiles are considered outliers and are removed from the illustration.

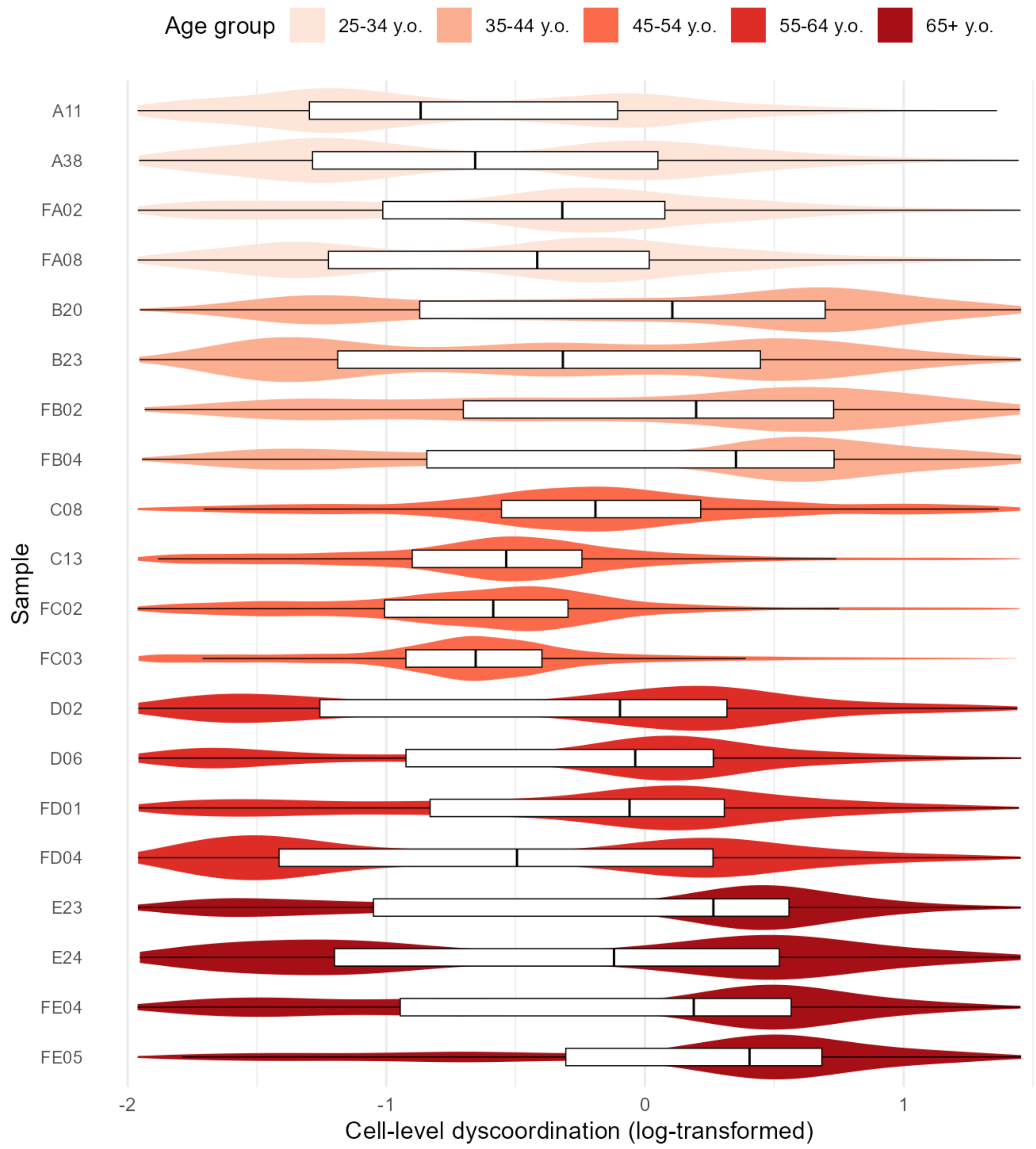


**Supplementary Figure 56 Distribution of cell-level dyscoordination across biological replicates in aging human T cells (Terekhova et al. 2023).**

Violin and box plots show the distribution of log-transformed cell-level dyscoordination for each individual donor, ordered and colored by age group. Values in the top and bottom 2.5% quantiles are considered outliers and are removed from the illustration.

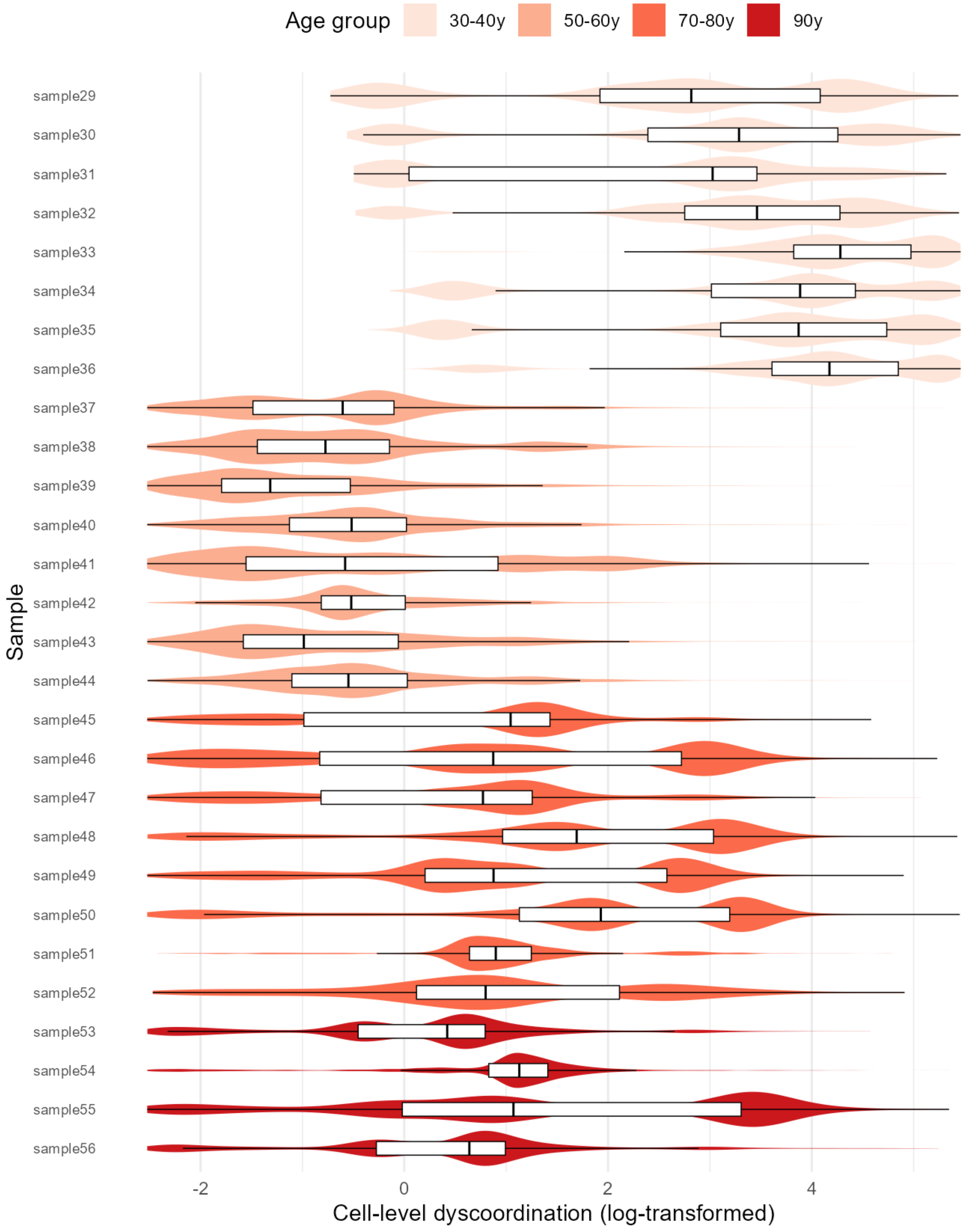
Age group
30-40y
50-60y
70-80y
90y
Sample
sample29
sample30
sample31
sample32
sample33
sample34
sample35
sample36
sample37
sample38
sample39
sample40
sample41
sample42
sample43
sample44
sample45
sample46
sample47
sample48
sample49
sample50
sample51
sample52
sample53
sample54
sample55
sample56
-2
0
2
4
Cell-level dyscoordination (log-transformed)

**Supplementary Figure 57 Distribution of cell-level dyscoordination across biological replicates in aging human T cells (Wang et al. 2025).**

Violin and box plots show the distribution of log-transformed cell-level dyscoordination for each individual sample, ordered and colored by age group. Values in the top and bottom 2.5% quantiles are considered outliers and are removed from the illustration.

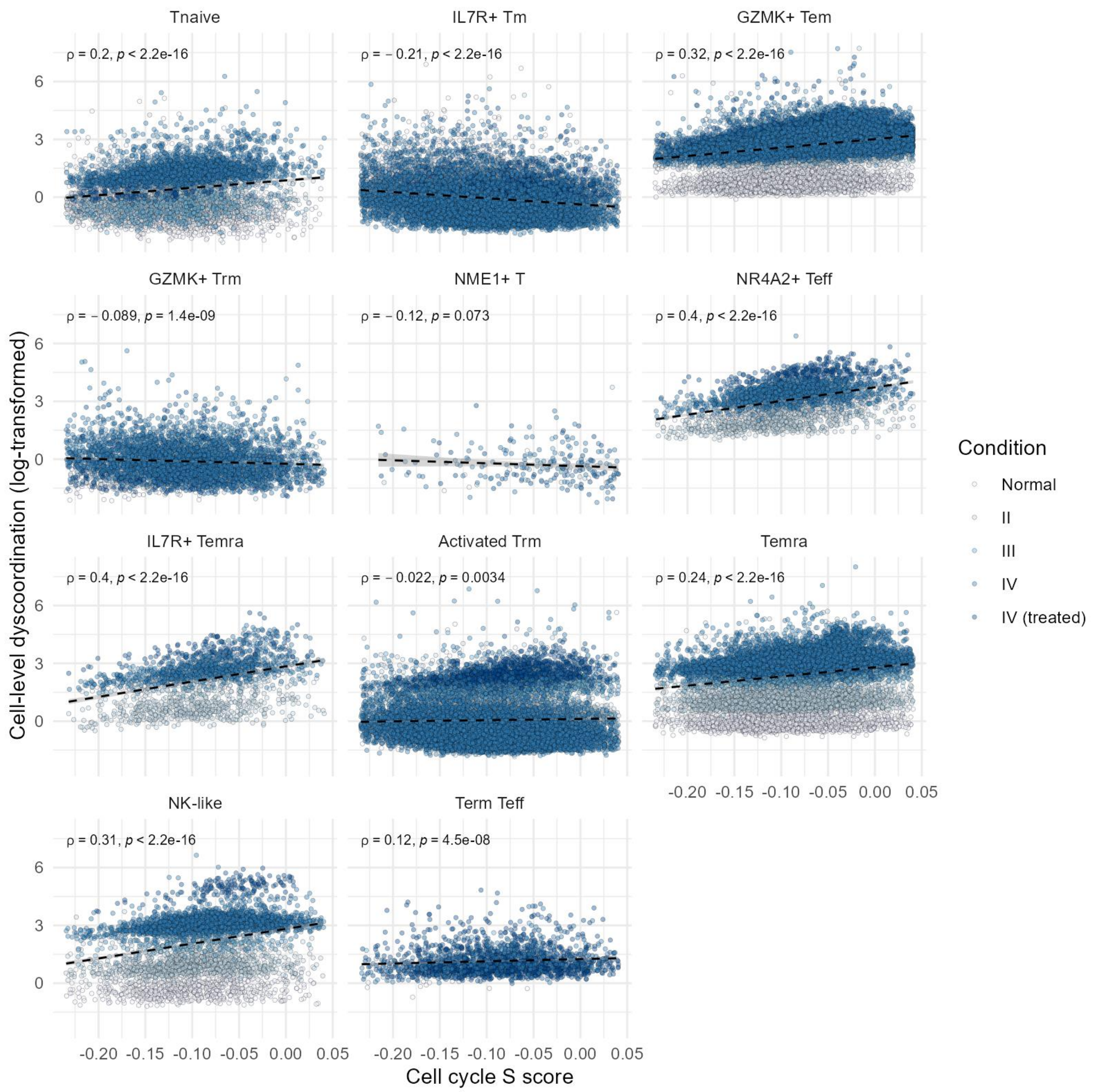


**Supplementary Figure 58 Correlation between cell cycle S phase score and cell-level dyscoordination in glioma CD8+ T cells.**

Scatter plots show the relationship between the cell cycle S phase score and log-transformed cell-level dyscoordination for each subset. Black dashed lines indicate linear regression fits. Spearman's correlation coefficients and p-values are shown on the top left corner of each panel.

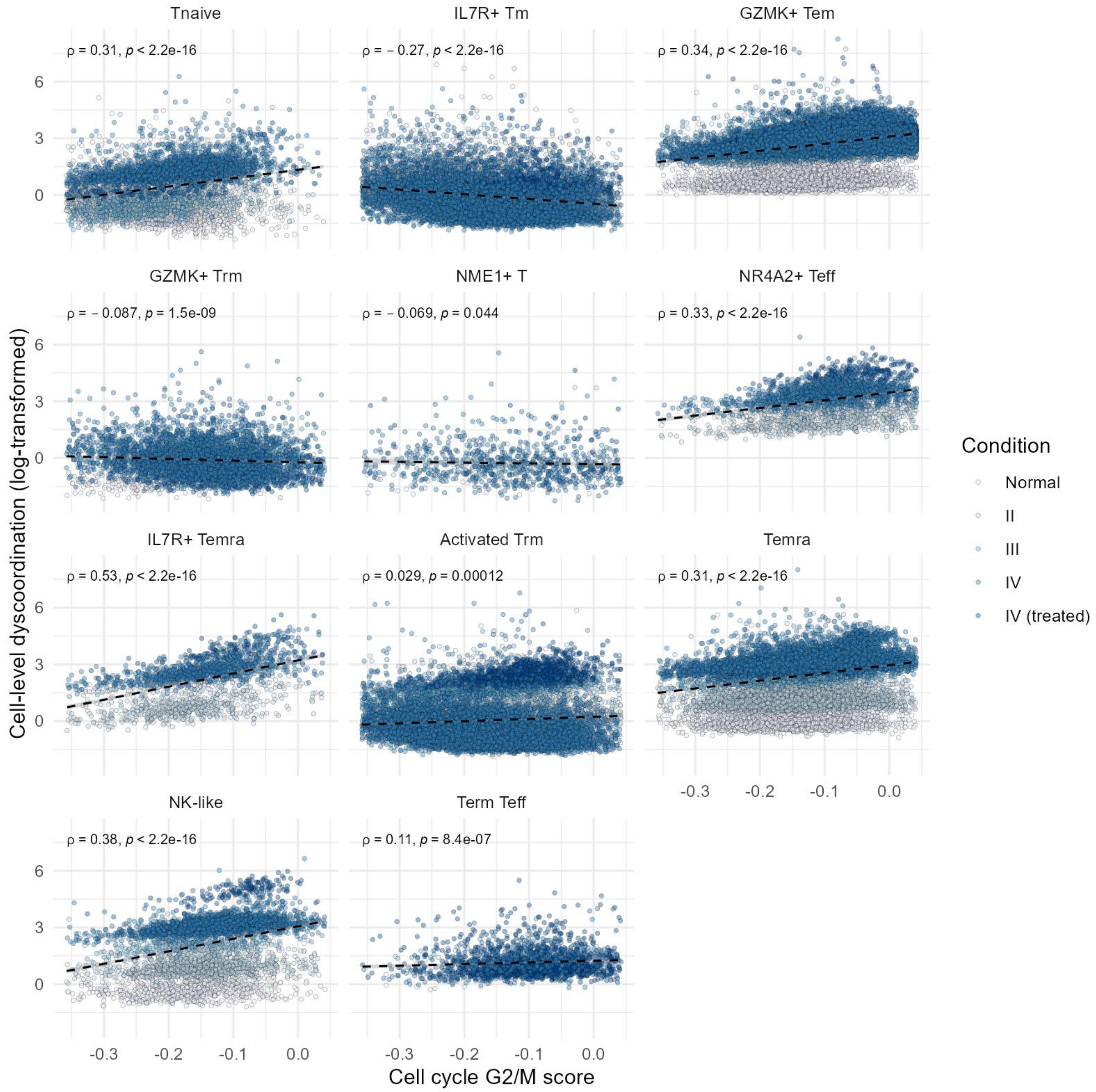


**Supplementary Figure 59 Correlation between cell cycle G2/M score and cell-level dyscoordination in glioma CD8+ T cells.**

Scatter plots show the relationship between the cell cycle G2/M phase score and log-transformed cell-level dyscoordination for each subset. Black dashed lines indicate linear regression fits. Spearman's correlation coefficients and p-values are shown on the top left corner of each panel.

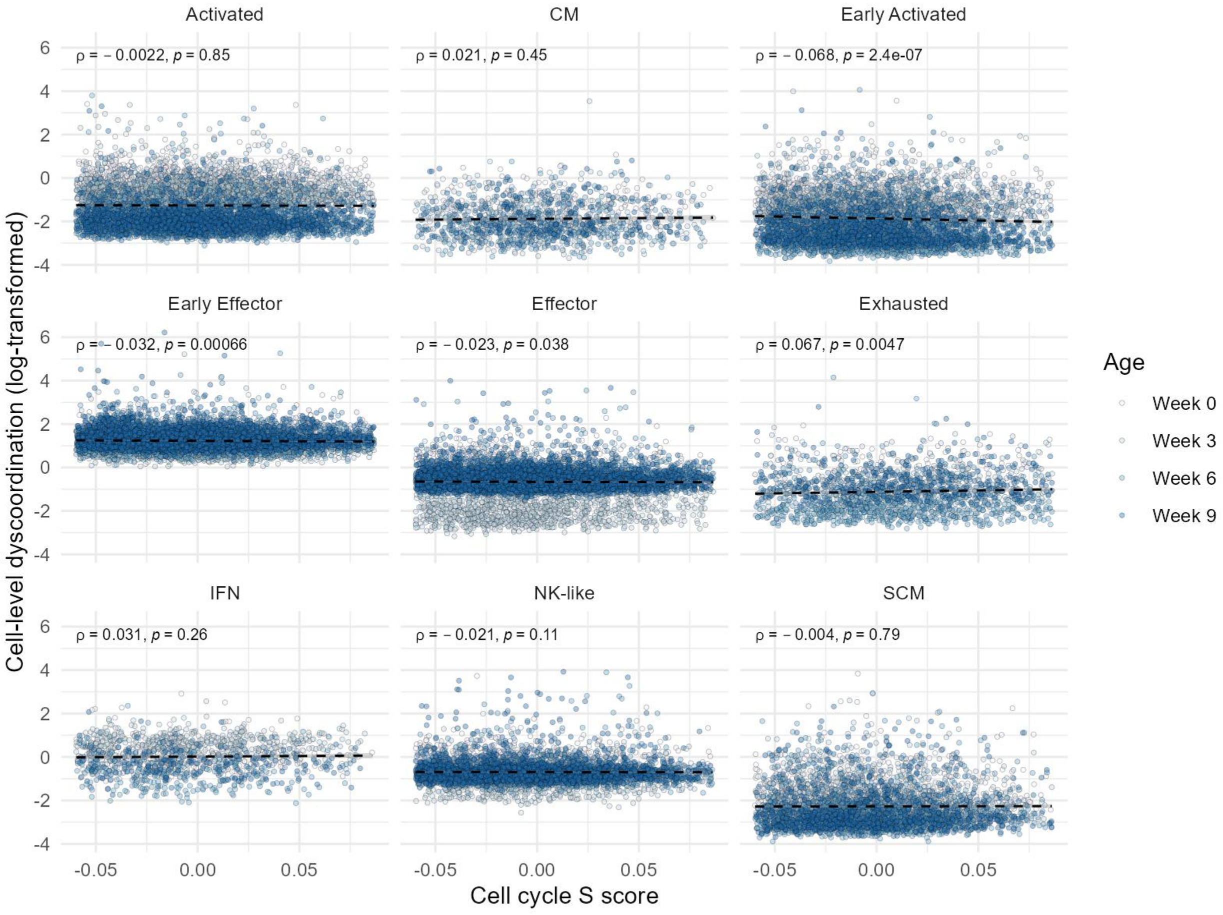


**Supplementary Figure 60 Correlation between cell cycle S phase score and cell-level dyscoordination in treated melanoma CD8+ T cells.**

Scatter plots show the relationship between the cell cycle S phase score and log-transformed cell-level dyscoordination for each subset. Black dashed lines indicate linear regression fits. Spearman’s correlation coefficients and p-values are shown on the top left corner of each panel.

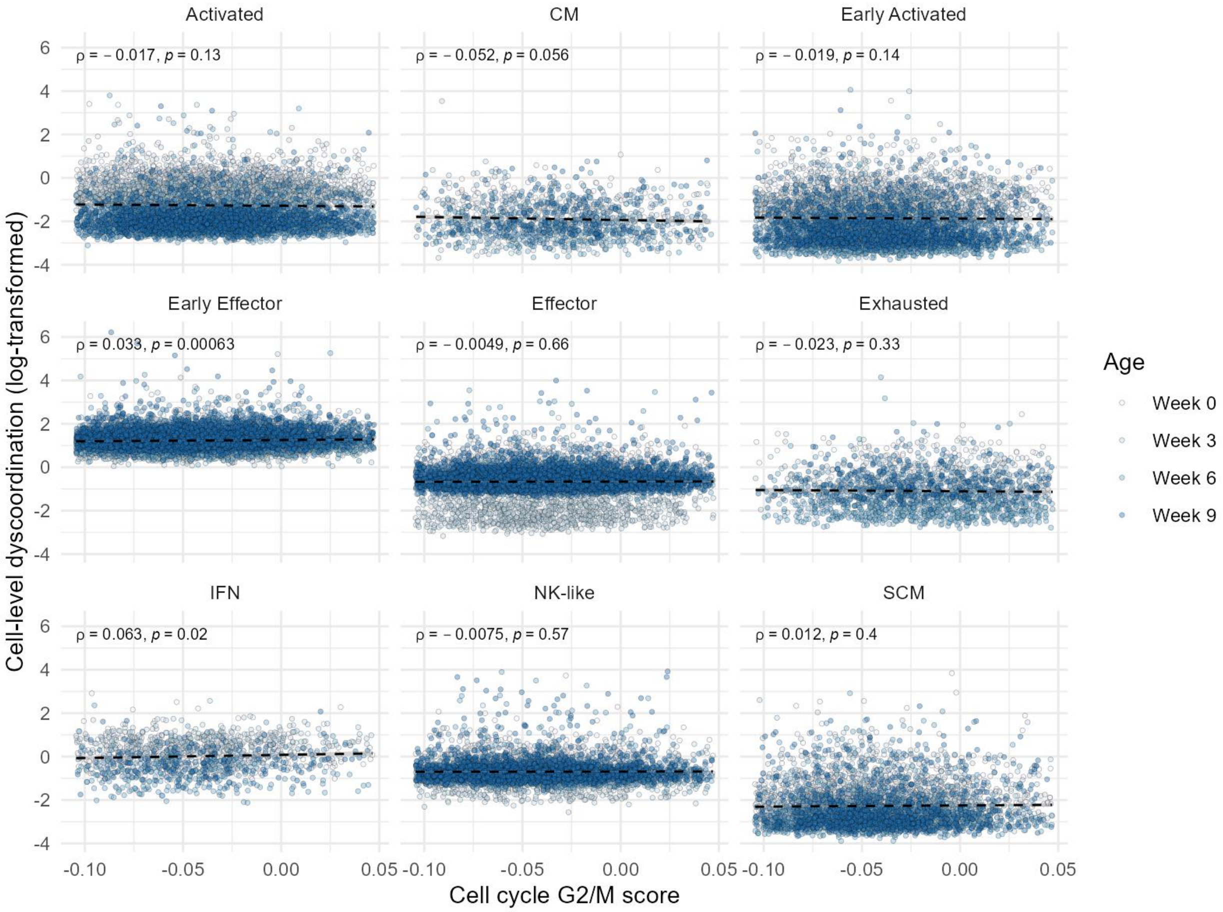


**Supplementary Figure 61 Correlation between cell cycle G2/M score and cell-level dyscoordination in treated melanoma CD8+ T cells.**

Scatter plots show the relationship between the cell cycle G2/M phase score and log-transformed cell-level dyscoordination for each subset. Black dashed lines indicate linear regression fits. Spearman's correlation coefficients and p-values are shown on the top left corner of each panel.

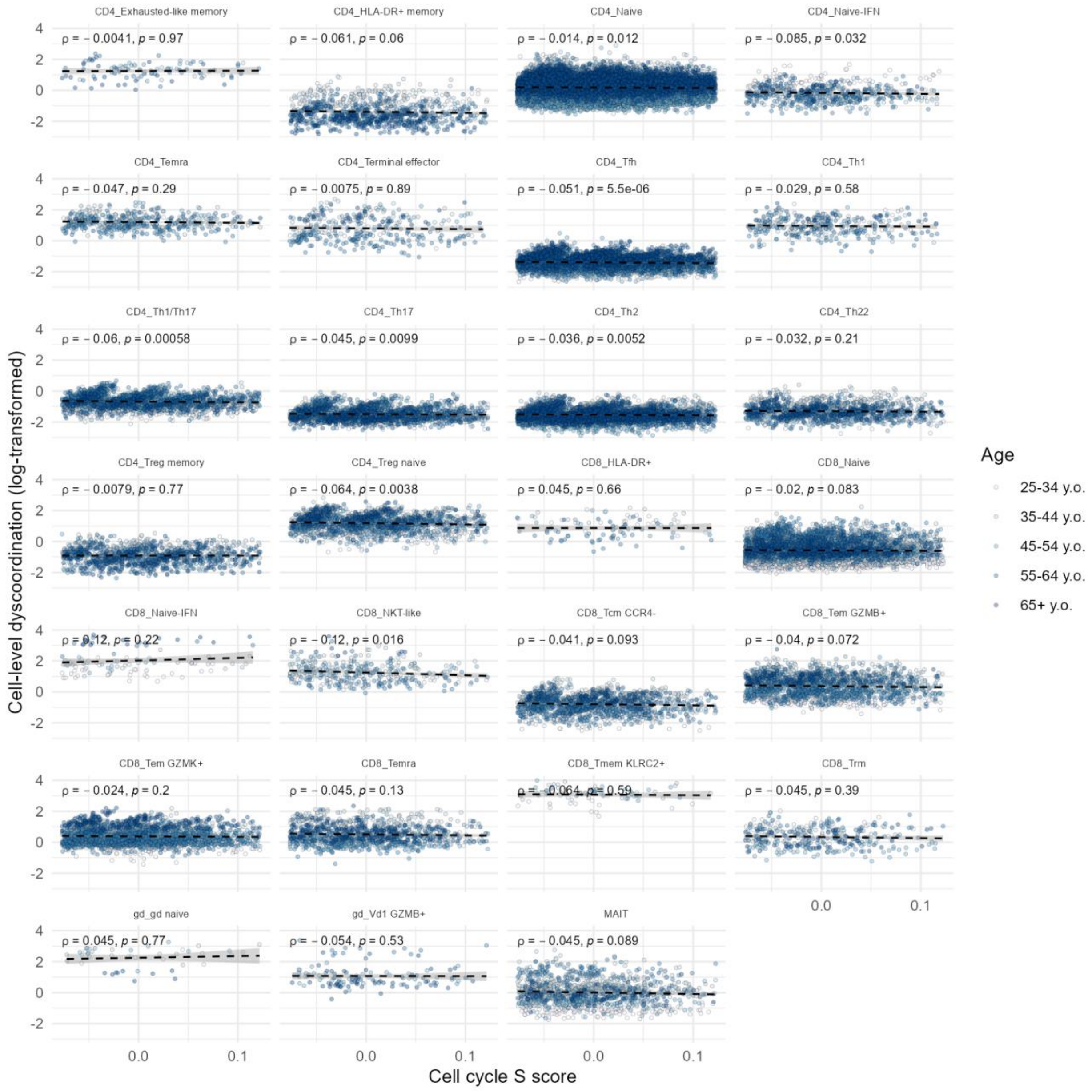


**Supplementary Figure 62 Correlation between cell cycle S phase score and cell-level dyscoordination in aging human T cells (Terekhova et al. 2023).**

Scatter plots show the relationship between the cell cycle S phase score and log-transformed cell-level dyscoordination for each subset. Black dashed lines indicate linear regression fits. Spearman's correlation coefficients and p-values are shown on the top left corner of each panel.

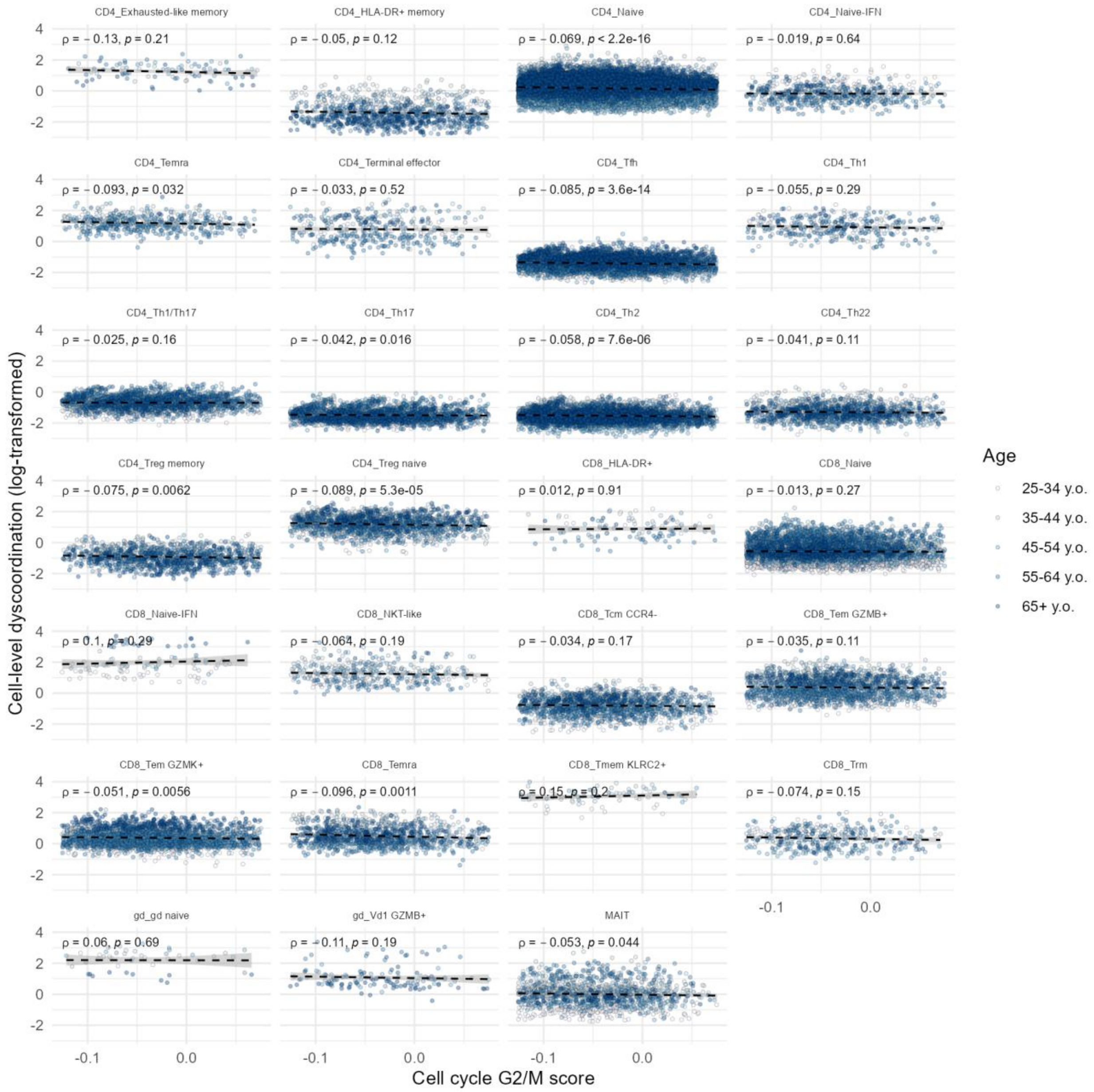


**Supplementary Figure 63 Correlation between cell cycle G2/M score and cell-level dyscoordination in aging human T cells (Terekhova et al. 2023).**

Scatter plots show the relationship between the cell cycle G2/M phase score and log-transformed cell-level dyscoordination for each subset. Black dashed lines indicate linear regression fits. Spearman’s correlation coefficients and p-values are shown on the top left corner of each panel.

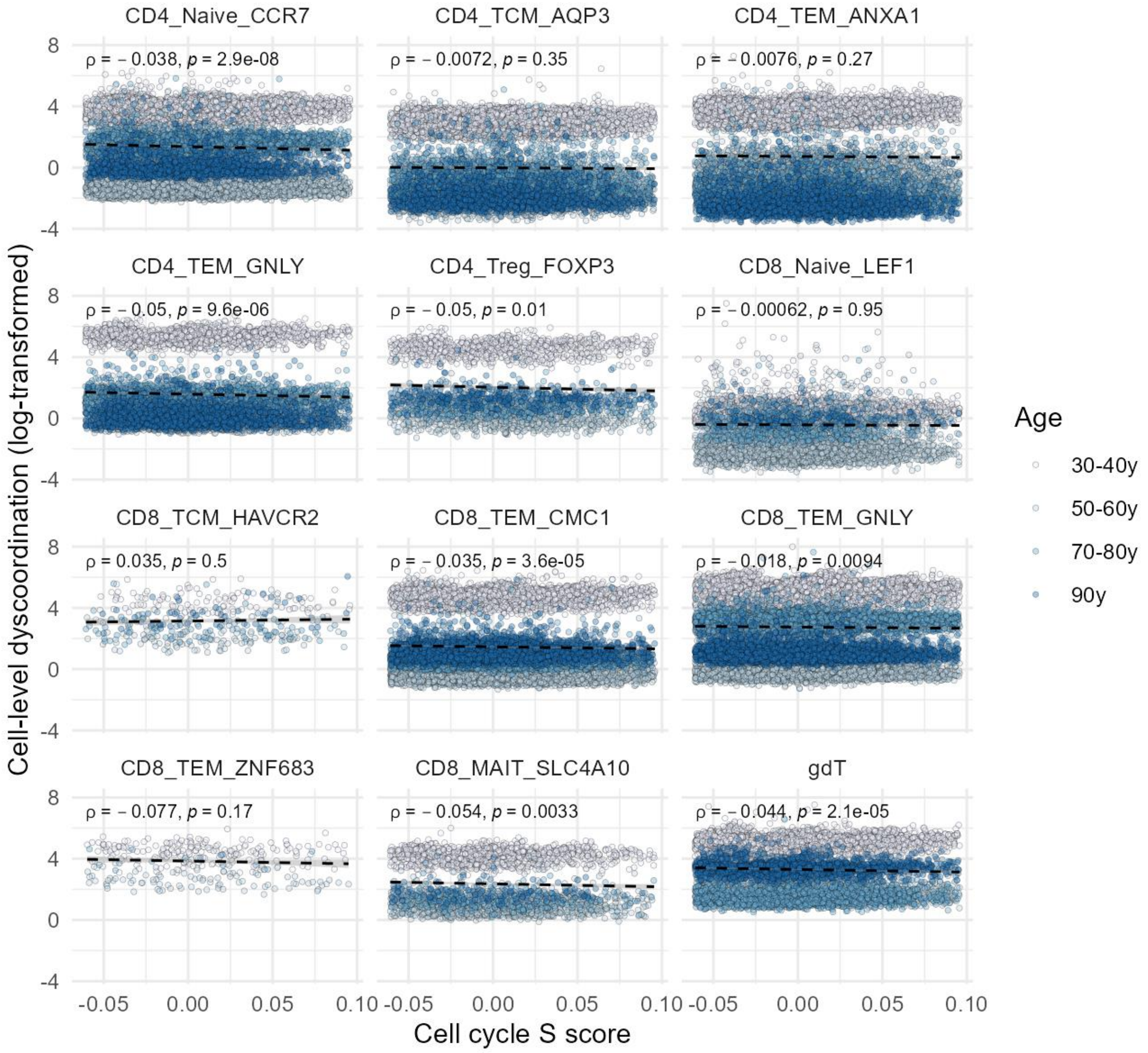


**Supplementary Figure 64 Correlation between cell cycle S phase score and cell-level dyscoordination in aging human T cells (Wang et al. 2025).**

Scatter plots show the relationship between the cell cycle S phase score and log-transformed cell-level dyscoordination for each subset. Black dashed lines indicate linear regression fits. Spearman’s correlation coefficients and p-values are shown on the top left corner of each panel.

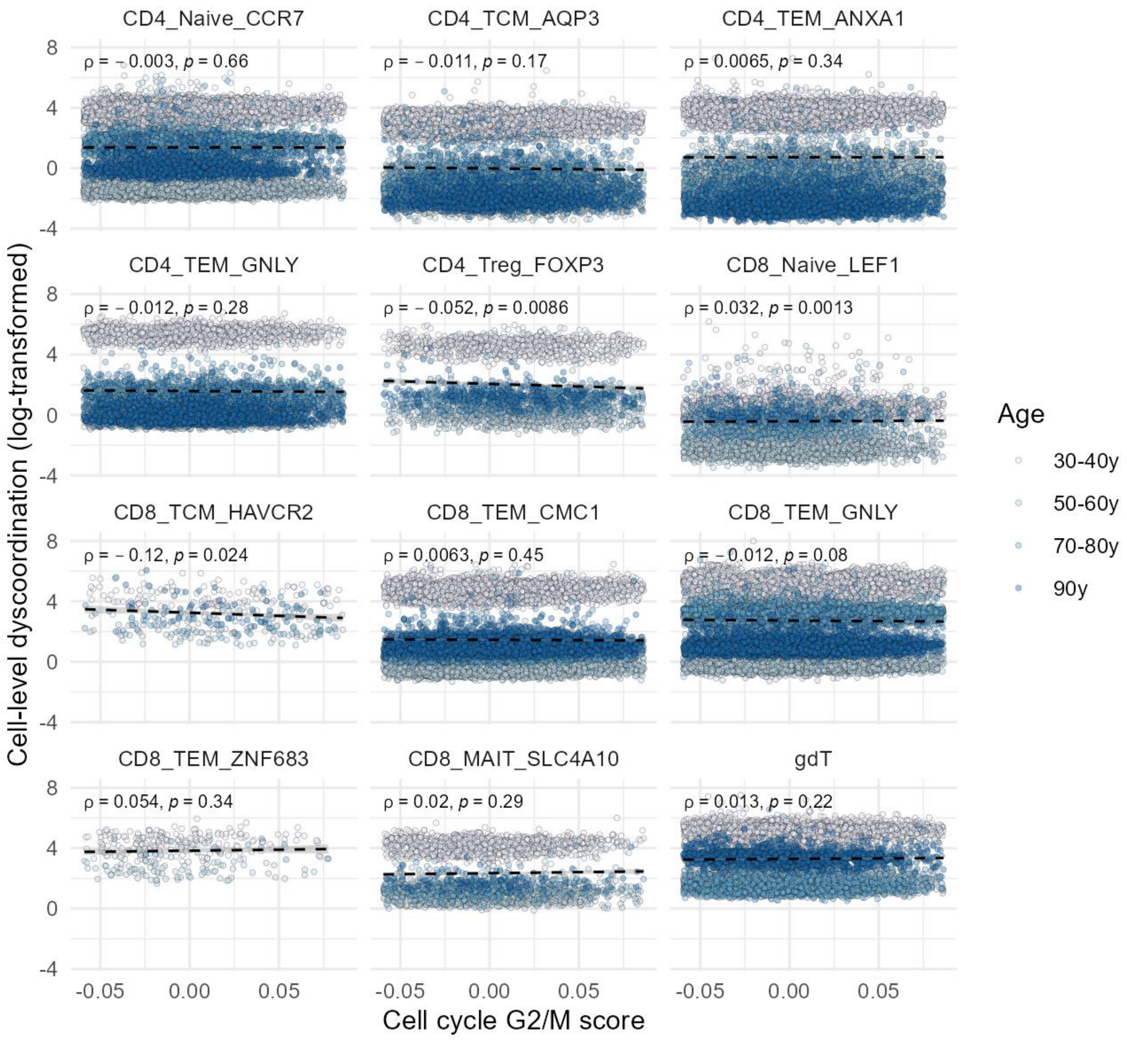


**Supplementary Figure 65 Correlation between cell cycle G2/M score and cell-level dyscoordination in aging human T cells (Wang et al. 2025).**

Scatter plots show the relationship between the cell cycle G2/M phase score and log-transformed cell-level dyscoordination for each subset. Black dashed lines indicate linear regression fits. Spearman's correlation coefficients and p-values are shown on the top left corner of each panel.

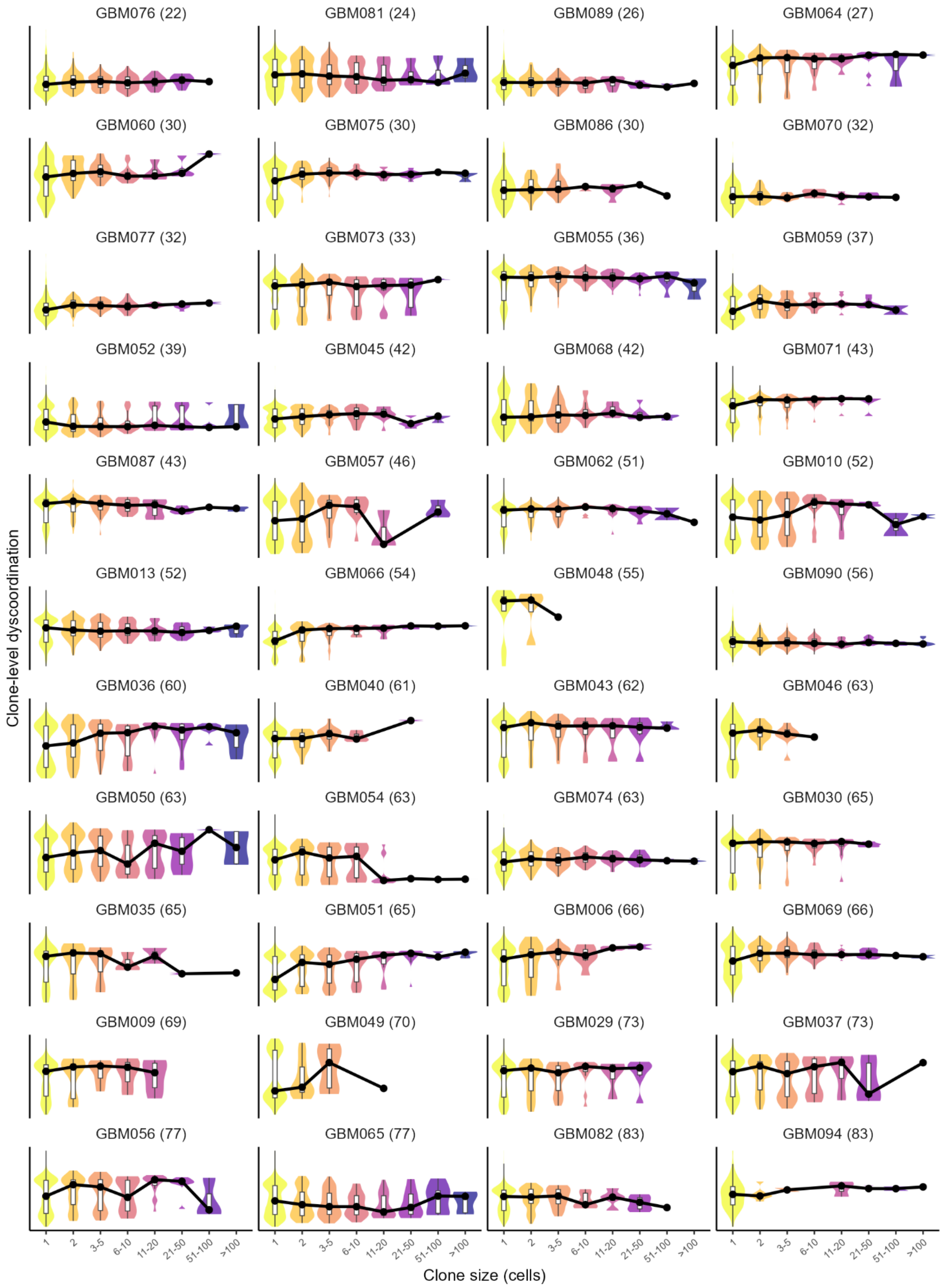

GBM076 (22)
GBM081 (24)
GBM089 (26)
GBM064 (27)
GBM060 (30)
GBM075 (30)
GBM086 (30)
GBM070 (32)
GBM077 (32)
GBM073 (33)
GBM055 (36)
GBM059 (37)
GBM052 (39)
GBM045 (42)
GBM068 (42)
GBM071 (43)
GBM087 (43)
GBM057 (46)
GBM062 (51)
GBM010 (52)
GBM013 (52)
GBM066 (54)
GBM048 (55)
GBM090 (56)
GBM036 (60)
GBM040 (61)
GBM043 (62)
GBM046 (63)
GBM050 (63)
GBM054 (63)
GBM074 (63)
GBM030 (65)
GBM035 (65)
GBM051 (65)
GBM006 (66)
GBM069 (66)
GBM009 (69)
GBM049 (70)
GBM029 (73)
GBM037 (73)
GBM056 (77)
GBM065 (77)
GBM082 (83)
GBM094 (83)
Clone-level dyscoordination
1
2
3-5
6-10
11-20
21-50
51-100
>100
Clone size (cells)

**Supplementary Figure 66 Distribution of within-donor clone-level dyscoordination by clone size in glioma CD8⁺ T cells.**

Violin and box plots show the distribution of clone-level dyscoordination, or the mean cell-level dyscoordination across cells in the same TCR clone, by clone size. Values are displayed separately for each patient, with the patient's age indicated in parentheses in each subplot title. Values in the top and bottom 1% quantiles within each patient are considered outliers and are removed from the illustration. Clone size is grouped into expansion strata along the x-axis, and the black line connects per-stratum medians within each patient (44 patients). Only patients whose clones span at least two clone-size strata are displayed.

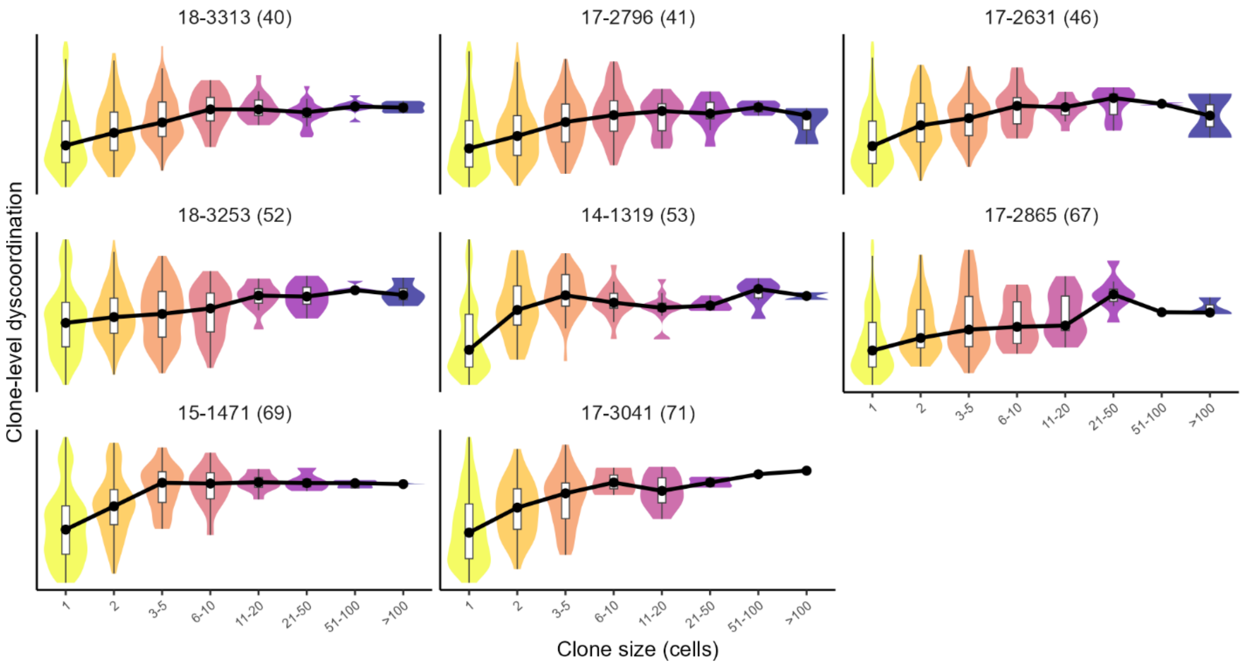


**Supplementary Figure 67 Distribution of within-donor clone-level dyscoordination by clone size in treated melanoma CD8⁺ T cells.**

Violin and box plots show the distribution of clone-level dyscoordination, or the mean cell-level dyscoordination across cells in the same TCR clone, by clone size. Values are displayed separately for each patient, with the patient's age indicated in parentheses in each subplot title. Values in the top and bottom 1% quantiles within each patient are considered outliers and are removed from the illustration. Clone size is grouped into expansion strata along the x-axis, and the black line connects per-stratum medians within each patient (8 patients). Only patients whose clones span at least two clone-size strata are displayed.

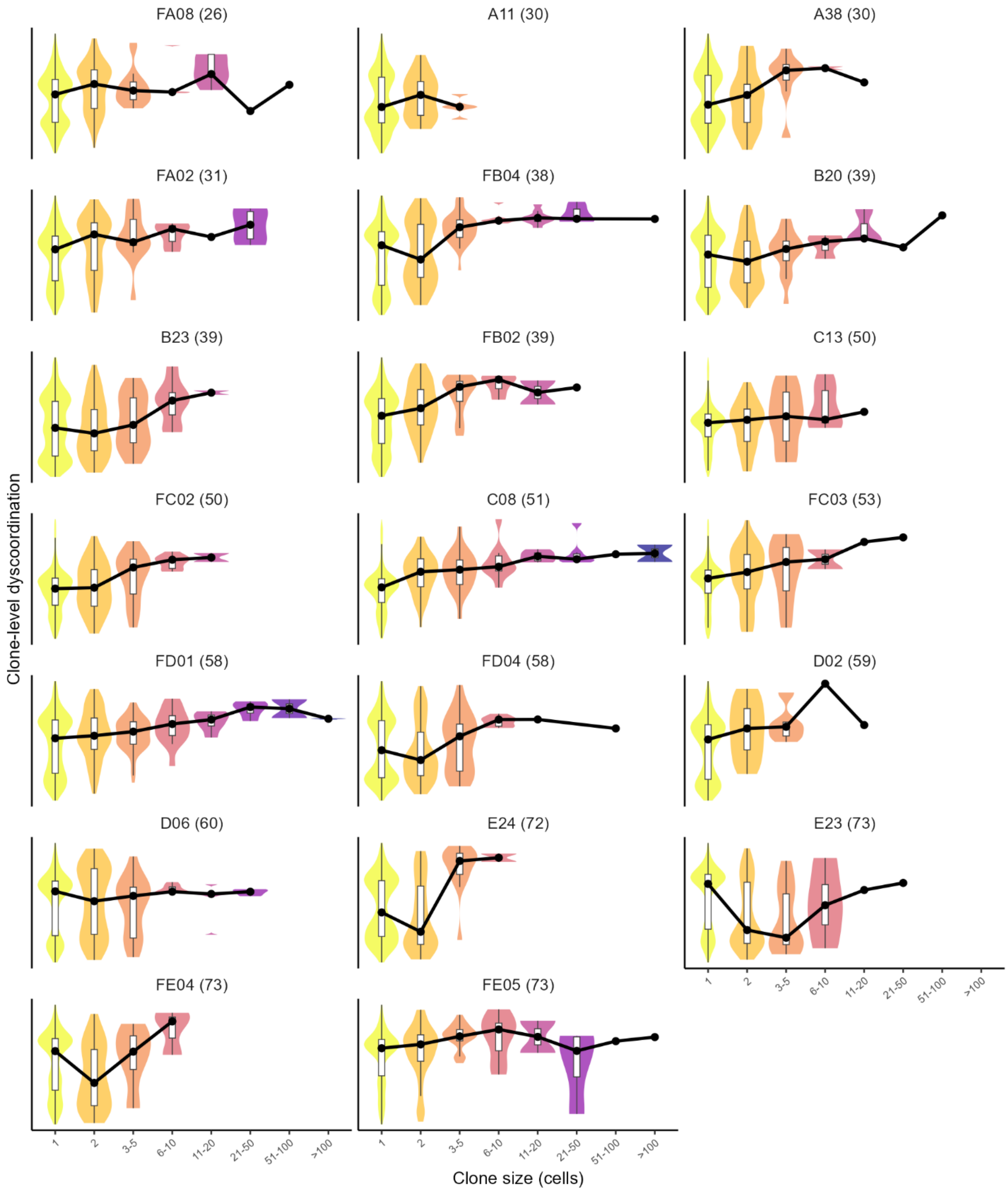

FA08 (26)
A11 (30)
A38 (30)
FA02 (31)
FB04 (38)
B20 (39)
B23 (39)
FB02 (39)
C13 (50)
FC02 (50)
C08 (51)
FC03 (53)
FD01 (58)
FD04 (58)
D02 (59)
D06 (60)
E24 (72)
E23 (73)
FE04 (73)
FE05 (73)
Clone-level dyscoordination
1
2
3-5
6-10
11-20
21-50
51-100
>100
Clone size (cells)

**Supplementary Figure 68 Distribution of within-donor clone-level dyscoordination by clone size in aging human T cells (Terekhova et al. 2023).**

Violin and box plots show the distribution of clone-level dyscoordination, or the mean cell-level dyscoordination across cells in the same TCR clone, by clone size. Values are displayed separately for each patient, with the patient's age indicated in parentheses in each subplot title. Values in the top and bottom 1% quantiles within each patient are considered outliers and are removed from the illustration. Clone size is grouped into expansion strata along the x-axis, and the black line connects per-stratum medians within each patient (20 patients). Only patients whose clones span at least two clone-size strata are displayed.

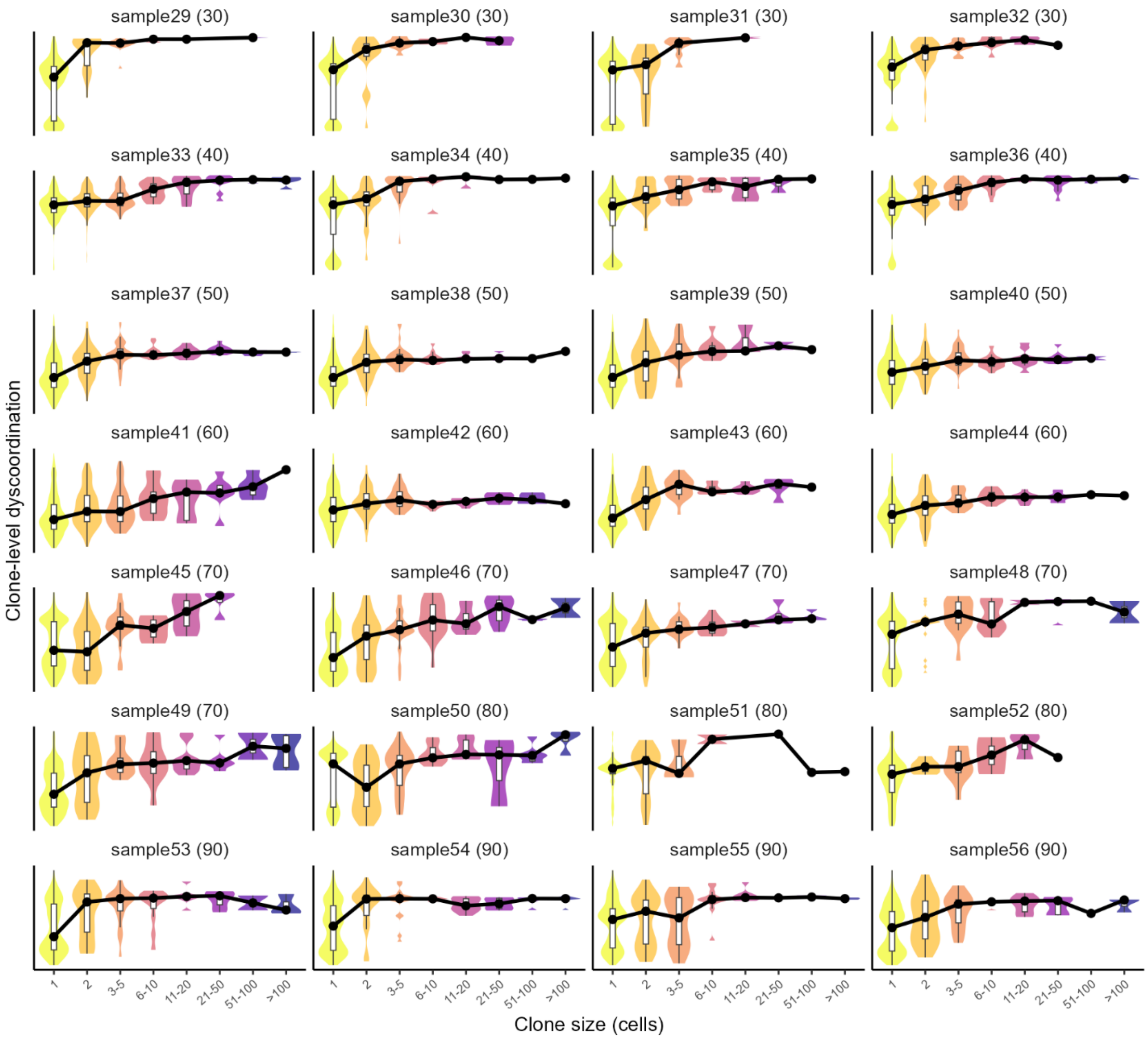


**Supplementary Figure 69 Distribution of within-donor clone-level dyscoordination by clone size in aging human T cells (Wang et al. 2025).**

Violin and box plots show the distribution of clone-level dyscoordination, or the mean cell-level dyscoordination across cells in the same TCR clone, by clone size. Values are displayed separately for each patient, with the patient's age indicated in parentheses in each subplot title. Values in the top and bottom 1% quantiles within each patient are considered outliers and are removed from the illustration. Clone size is grouped into expansion strata along the x-axis, and the black line connects per-stratum medians within each patient (28 patients). Only patients whose clones span at least two clone-size strata are displayed.

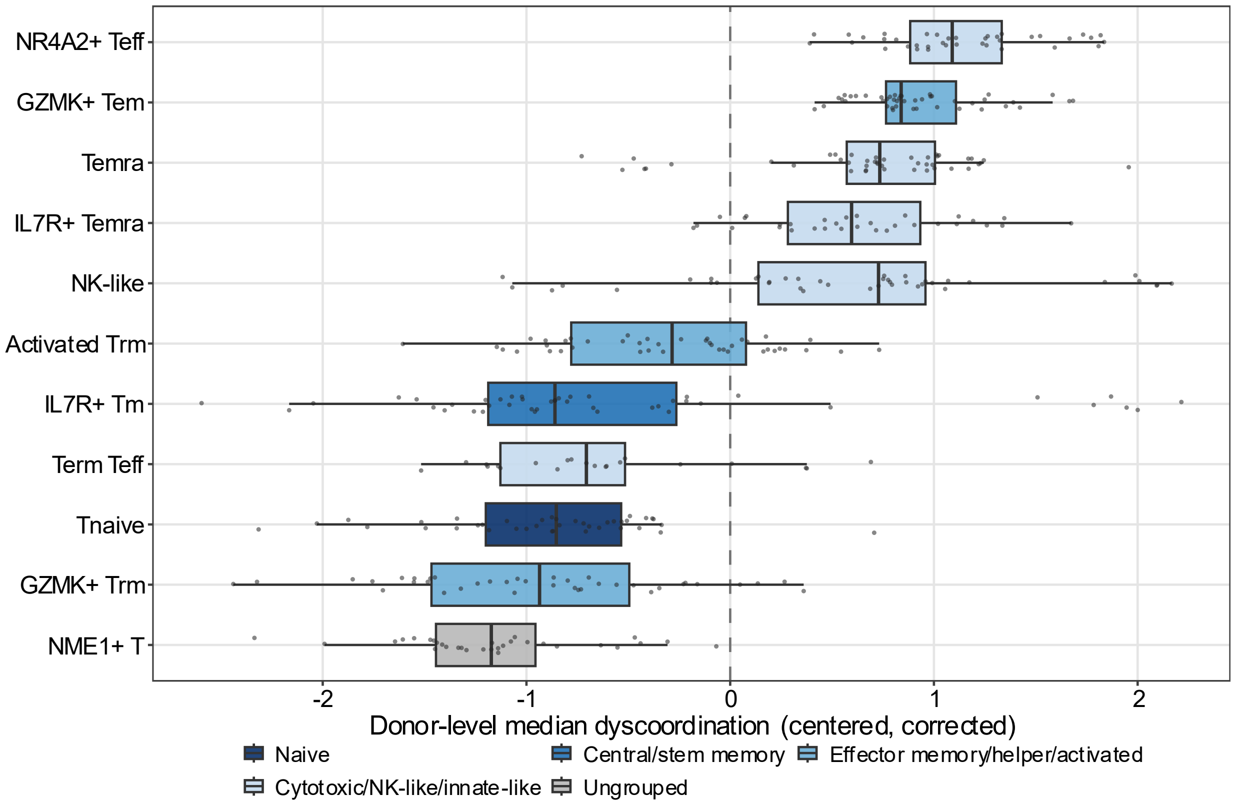


**Supplementary Figure 70 Transcriptional dyscoordination across annotated T cell subsets in glioma CD8$^{+}$ T cells.**

Box plots show the distribution across donors of subset-level transcriptional dyscoordination in glioma tumor-infiltrating CD8+ T cells (44 patients, 11 subsets). For each donor and each annotated subset, cells were summarized by the median depth-corrected cell-level dyscoordination, with at least 10 cells per donor and subset. Each value was then centered within its donor by subtracting that donor's mean across its own subsets so that no between-donor covariate can produce the ordering. Each point represents one donor, and subsets are ordered by their mean centered values. Subsets present in at least 5 donors are shown. Fill color denotes the coarse subset groups representing distinct lineages or transient states left ungrouped (gray).

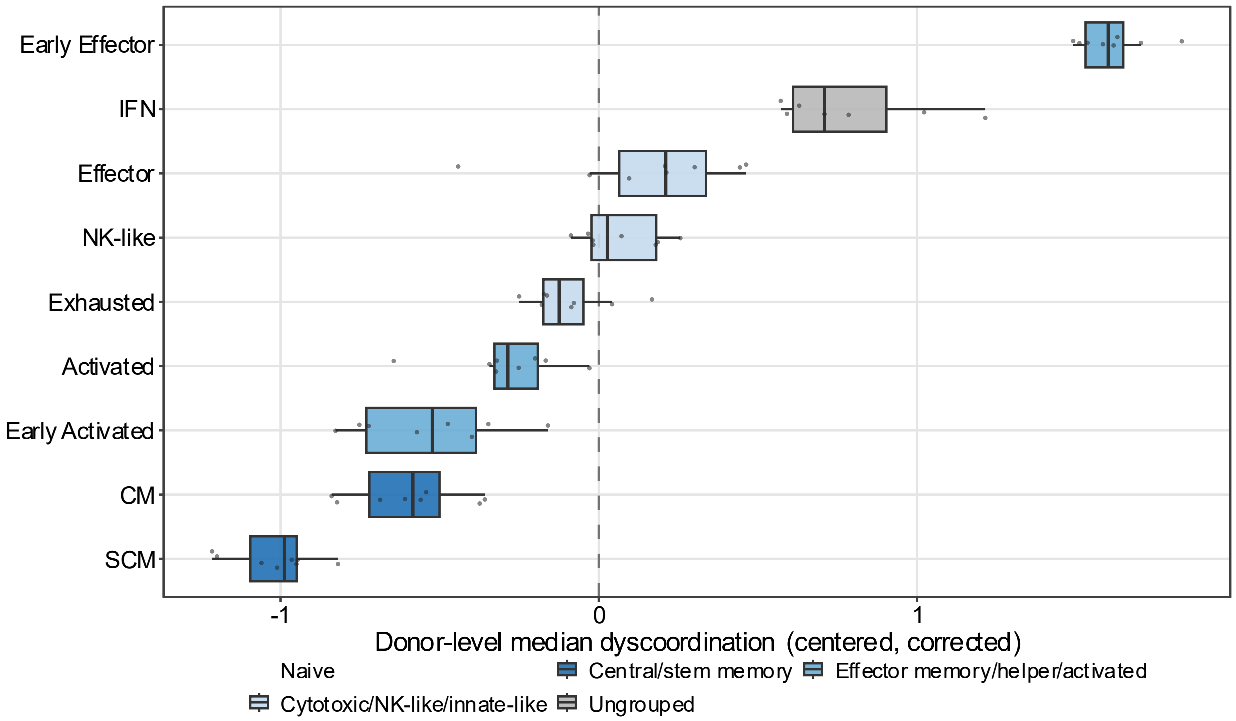


**Supplementary Figure 71 Transcriptional dyscoordination across annotated T cell subsets in treated melanoma CD8⁺ T cells.**

Box plots show the distribution across donors of subset-level transcriptional dyscoordination in treated melanoma CD8+ T cells (8 patients, 9 subsets). For each donor and each annotated subset, cells were summarized by the median depth-corrected cell-level dyscoordination, with at least 10 cells per donor and subset. Each value was then centered within its donor by subtracting that donor's mean across its own subsets. Each point represents one donor, and subsets are ordered by their mean centered value. Subsets present in at least 5 donors are shown. Fill color denotes the coarse subset groups representing distinct lineages or transient states left ungrouped (gray).

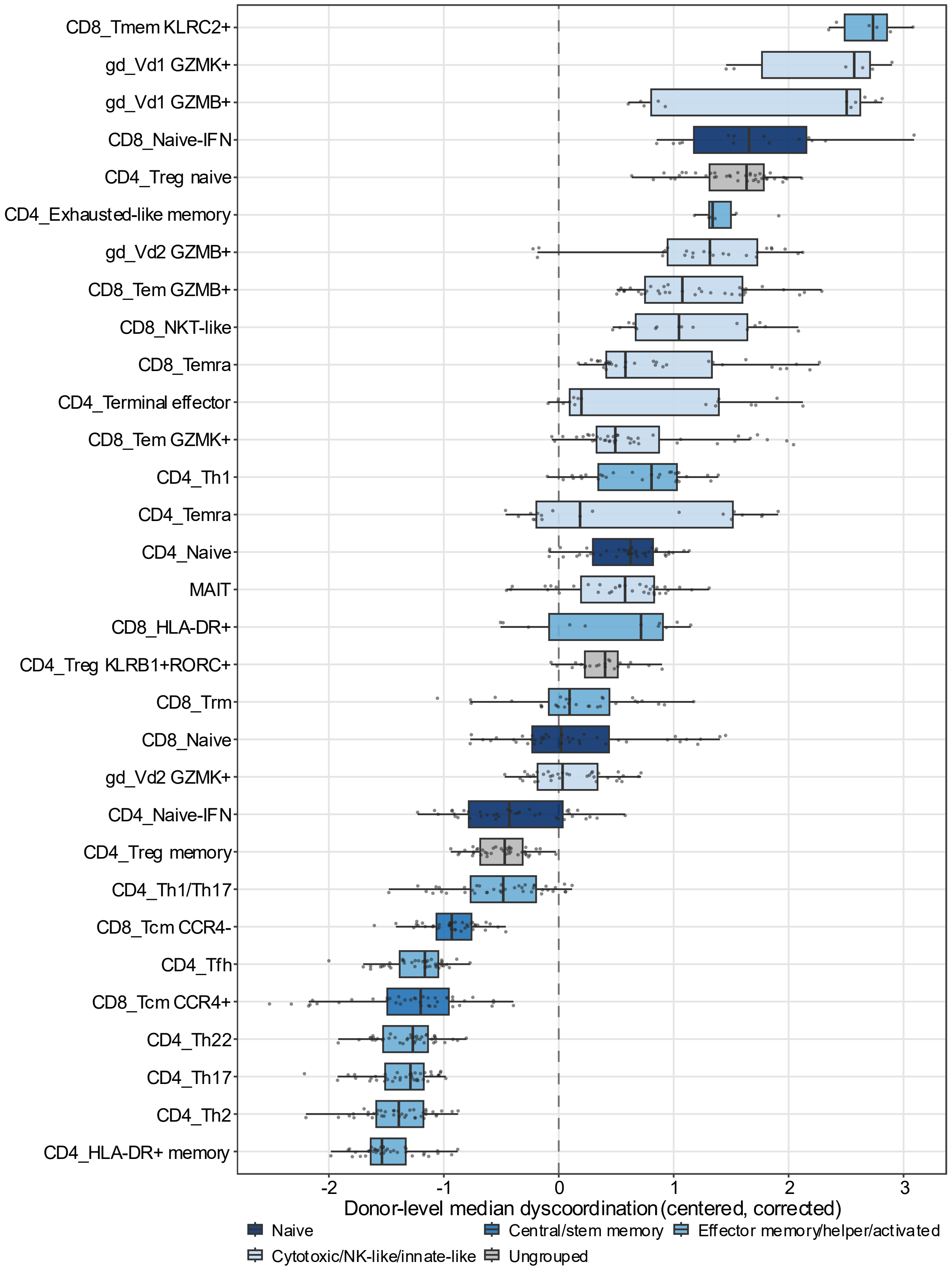
CD8_Tmem KLRC2+
gd_Vd1 GZMK+
gd_Vd1 GZMB+
CD8_Naive-IFN
CD4_Treg naive
CD4_Exhausted-like memory
gd_Vd2 GZMB+
CD8_Tem GZMB+
CD8_NKT-like
CD8_Temra
CD4_Terminal effector
CD8_Tem GZMK+
CD4_Th1
CD4_Temra
CD4_Naive
MAIT
CD8_HLA-DR+
CD4_Treg KLRB1+RORC+
CD8_Trm
CD8_Naive
gd_Vd2 GZMK+
CD4_Naive-IFN
CD4_Treg memory
CD4_Th1/Th17
CD8_Tcm CCR4-
CD4_Tfh
CD8_Tcm CCR4+
CD4_Th22
CD4_Th17
CD4_Th2
CD4_HLA-DR+ memory
-2
-1
0
1
2
3
Donor-level median dyscoordination (centered, corrected)
Naive
Central/stem memory
Effector memory/helper/activated
Cytotoxic/NK-like/innate-like
Ungrouped

**Supplementary Figure 72 Transcriptional dyscoordination across annotated T cell subsets in aging human T cells (Terekhova et al. 2023).**

Box plots show the distribution across donors of subset-level transcriptional dyscoordination in healthy aging blood T cells (40 donors, 31 subsets). For each donor and each annotated subset, cells were summarized by the median depth-corrected cell-level dyscoordination, with at least 10 cells per donor and subset. Each value was then centered within its donor by subtracting that donor's mean across its own subsets. Each point represents one donor, and subsets are ordered by their mean centered value. Subsets present in at least 5 donors are shown. Fill color denotes the coarse subset groups representing distinct lineages or transient states left ungrouped (gray).

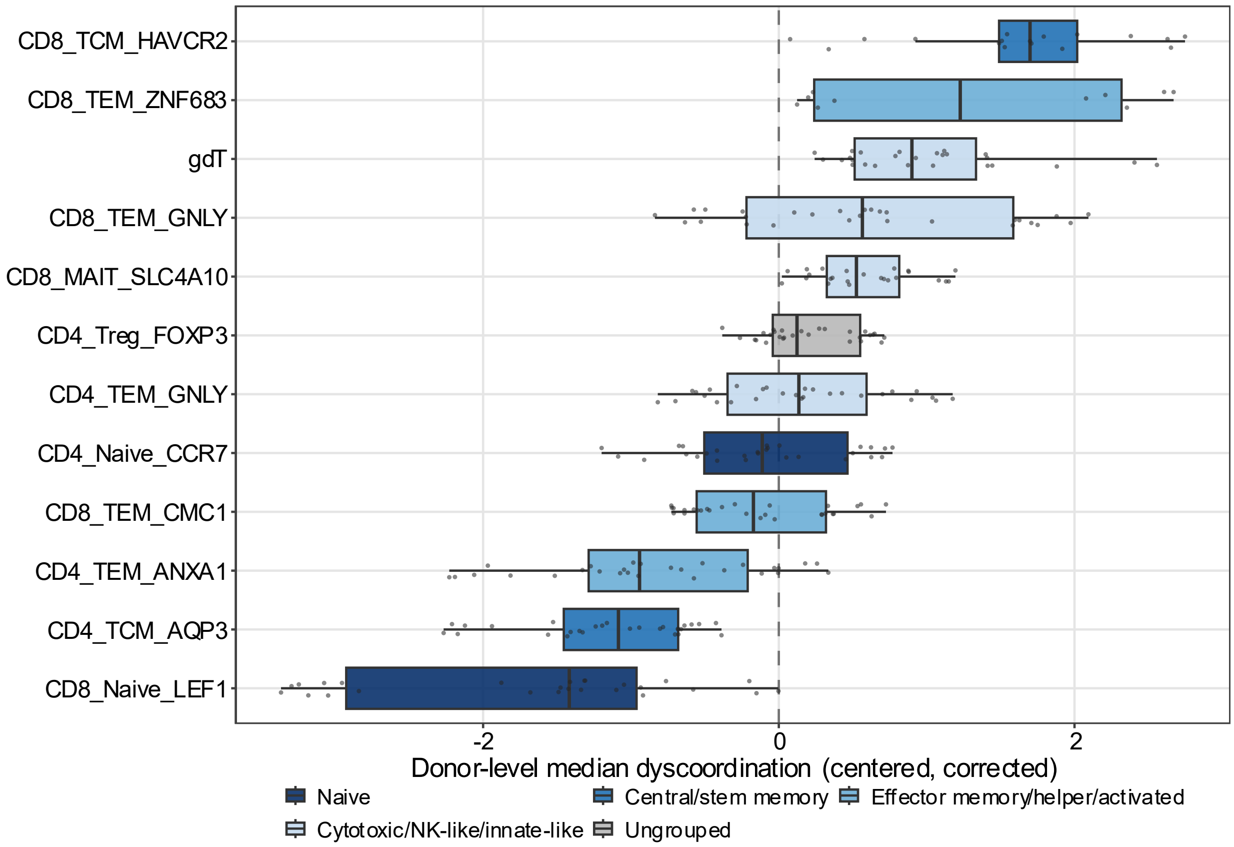


**Supplementary Figure 73 Transcriptional dyscoordination across annotated T cell subsets in aging human T cells (Wang et al. 2025).**

Box plots show the distribution across donors of subset-level transcriptional dyscoordination in healthy lifespan blood T cells (28 donors, 12 subsets). For each donor and each annotated subset, cells were summarized by the median depth-corrected cell-level dyscoordination, with at least 10 cells per donor and subset. Each value was then centered within its donor by subtracting that donor's mean across its own subsets, which removes the donor's overall level. Each point represents one donor, and subsets are ordered by their mean centered value. Subsets present in at least 5 donors are shown. Fill color denotes the coarse subset groups representing distinct lineages or transient states left ungrouped (gray).

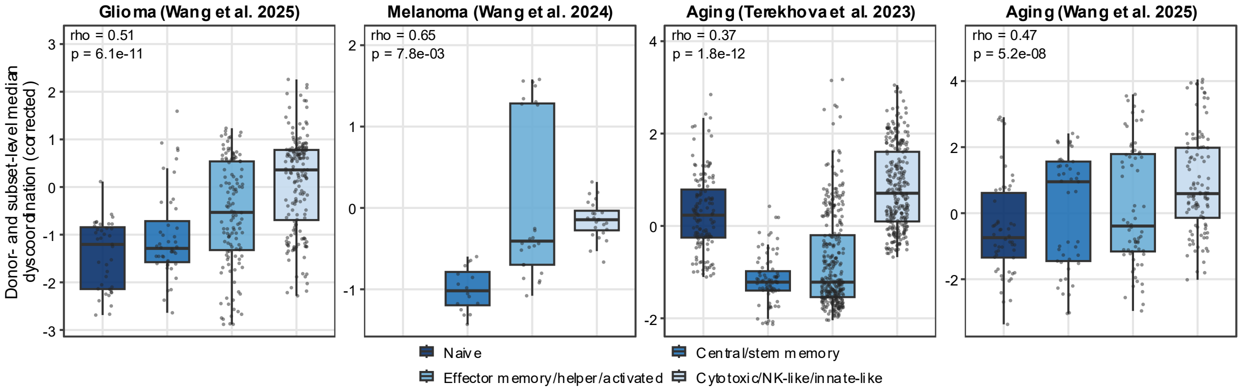


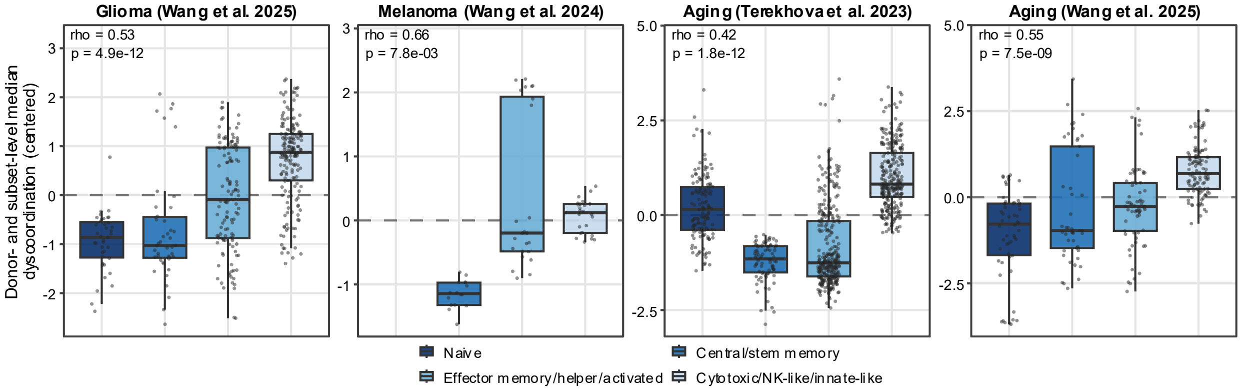


**Supplementary Figure 74 Transcriptional dyscoordination across coarse T cell subset groups without within-donor centering or depth correction.**

Box plots show the donor × subset median log-transformed cell-level dyscoordination without within-donor centering (top) or global depth correction (bottom). Each point represents one donor × subset pair, and fill color denotes the coarse subset groups representing distinct lineages.

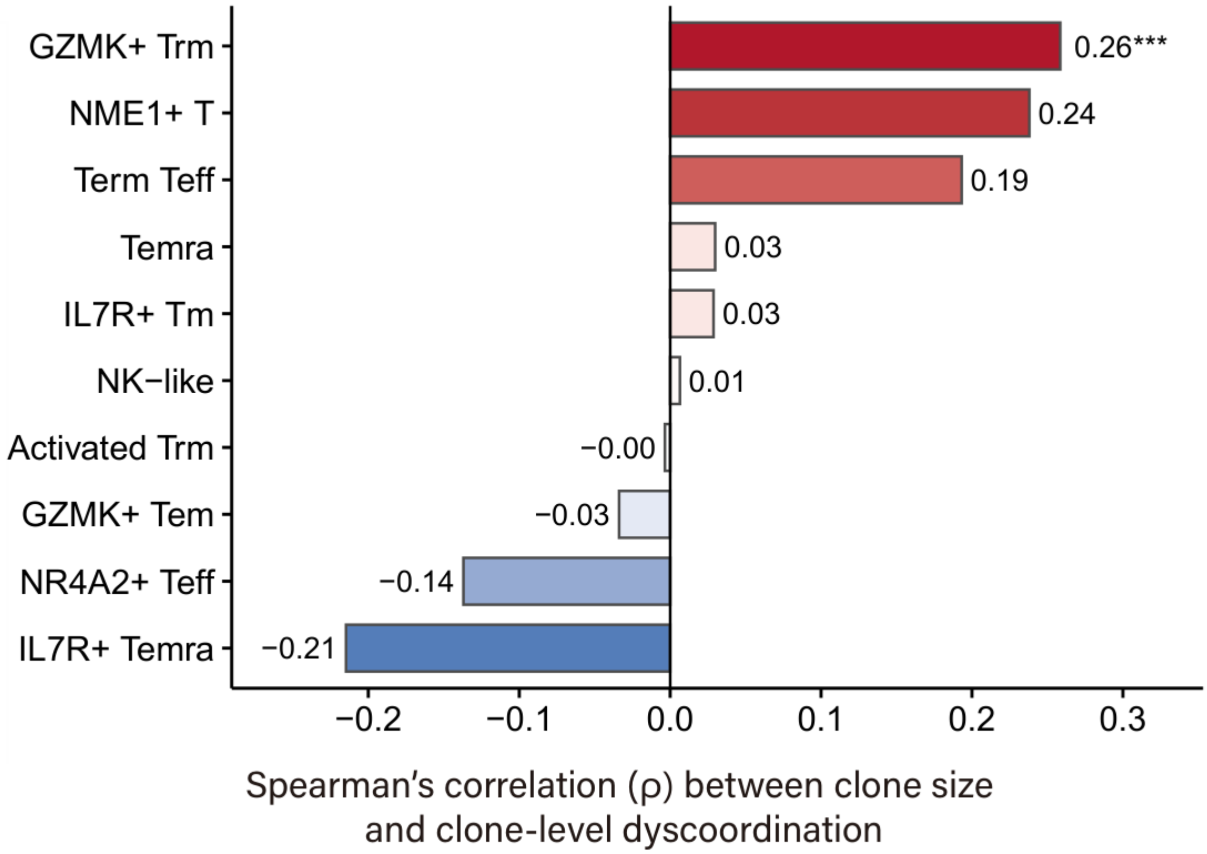


**Supplementary Figure 75 Subset-specific association between clone size and clone-level transcriptional dyscoordination in glioma CD8$^{+}$ T cells.**

Bar plots show the Spearman's correlation (ρ) between clone size and clone-level dyscoordination across all TCR clones assigned to a given dominant T cell subset. Cell types with over 20 qualifying clones are shown. Bars are colored and ordered by ρ, with asterisks denoting p-values after FDR control (* FDR < 0.05, ** < 0.01, *** < 0.001).

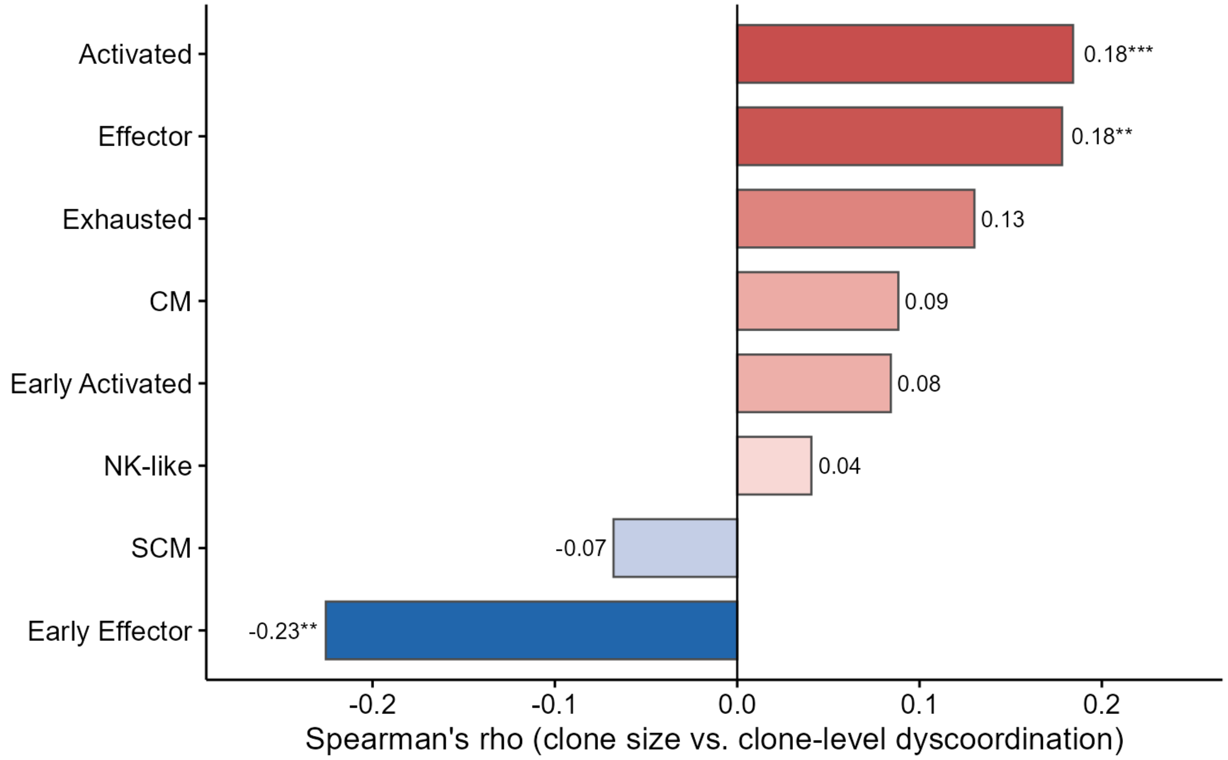


**Supplementary Figure 76 Subset-specific association between clone size and clone-level transcriptional dyscoordination in treated melanoma CD8⁺ T cells.**

Bar plots show the Spearman's correlation (ρ) between clone size and clone-level dyscoordination across all TCR clones assigned to a given dominant T cell subset. Cell types with over 20 qualifying clones are shown. Bars are colored and ordered by ρ, with asterisks denoting p-values after FDR control (* FDR < 0.05, ** < 0.01, *** < 0.001).

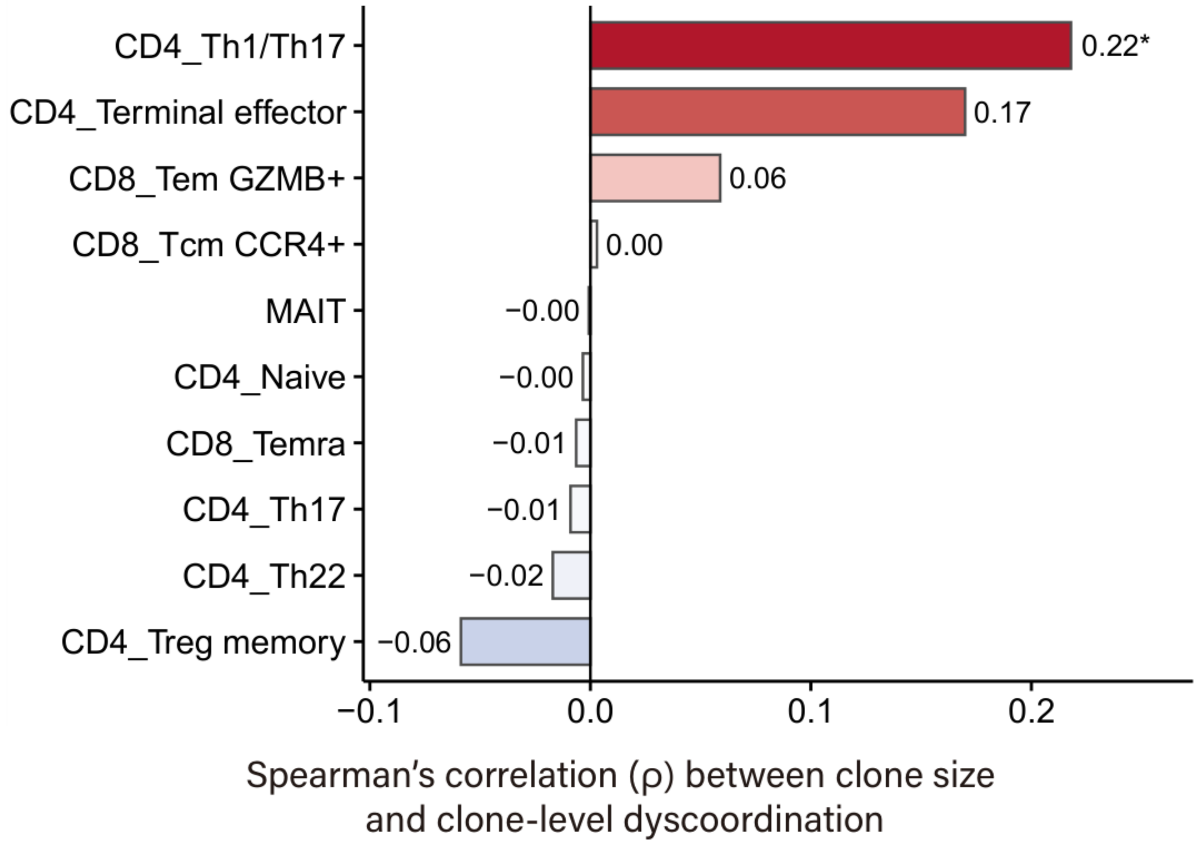


**Supplementary Figure 77 Subset-specific association between clone size and clone-level transcriptional dyscoordination in aging human T cells (Terekhova et al. 2023).**

Bar plots show the Spearman's correlation (ρ) between clone size and clone-level dyscoordination across all TCR clones assigned to a given dominant T cell subset. Cell types with over 20 qualifying clones are shown. Bars are colored and ordered by ρ, with asterisks denoting p-values after FDR control (* FDR < 0.05, ** < 0.01, *** < 0.001).

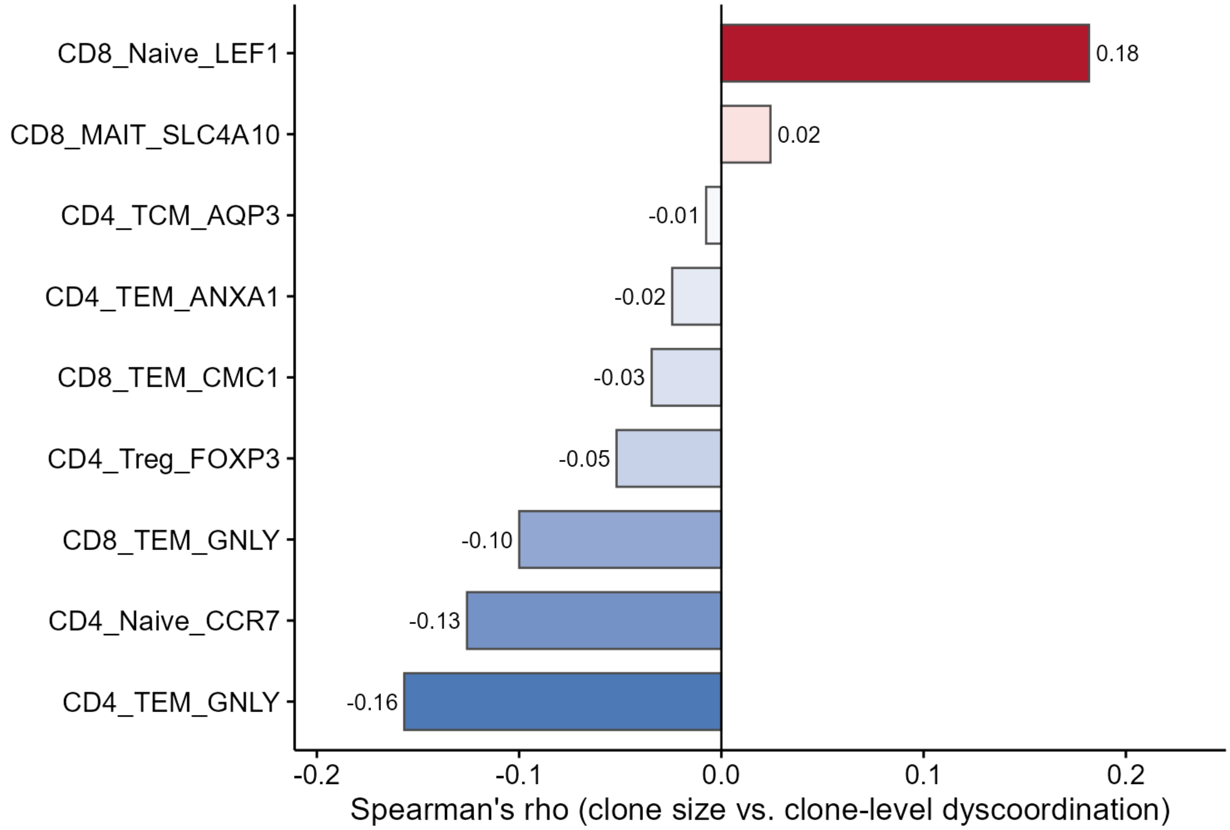


**Supplementary Figure 78 Subset-specific association between clone size and clone-level transcriptional dyscoordination in aging human T cells (Wang et al. 2025).**

Bar plots show the Spearman's correlation (ρ) between clone size and clone-level dyscoordination across all TCR clones assigned to a given dominant T cell subset. Cell types with over 20 qualifying clones are shown. Bars are colored and ordered by ρ, with asterisks denoting p-values after FDR control (* FDR < 0.05, ** < 0.01, *** < 0.001).

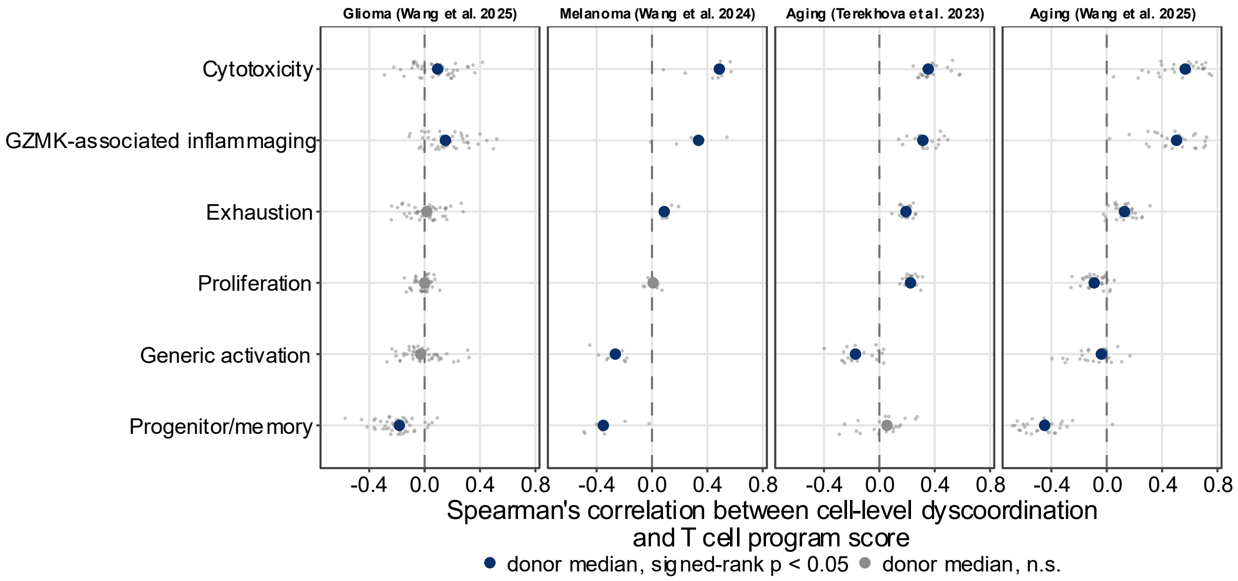


**Supplementary Figure 79 Correlation between transcriptional dyscoordination and T cell gene program scores without depth correction.**

Dot plots show the donor-level Spearman's correlations between cell-level dyscoordination and gene program score computed without the global depth correction. Each gray dot represents one correlation per donor, restricted to donors with at least 100 cells, and the colored point represents the median across donors, colored by whether p < 0.05 in the donor-level signed-rank test.

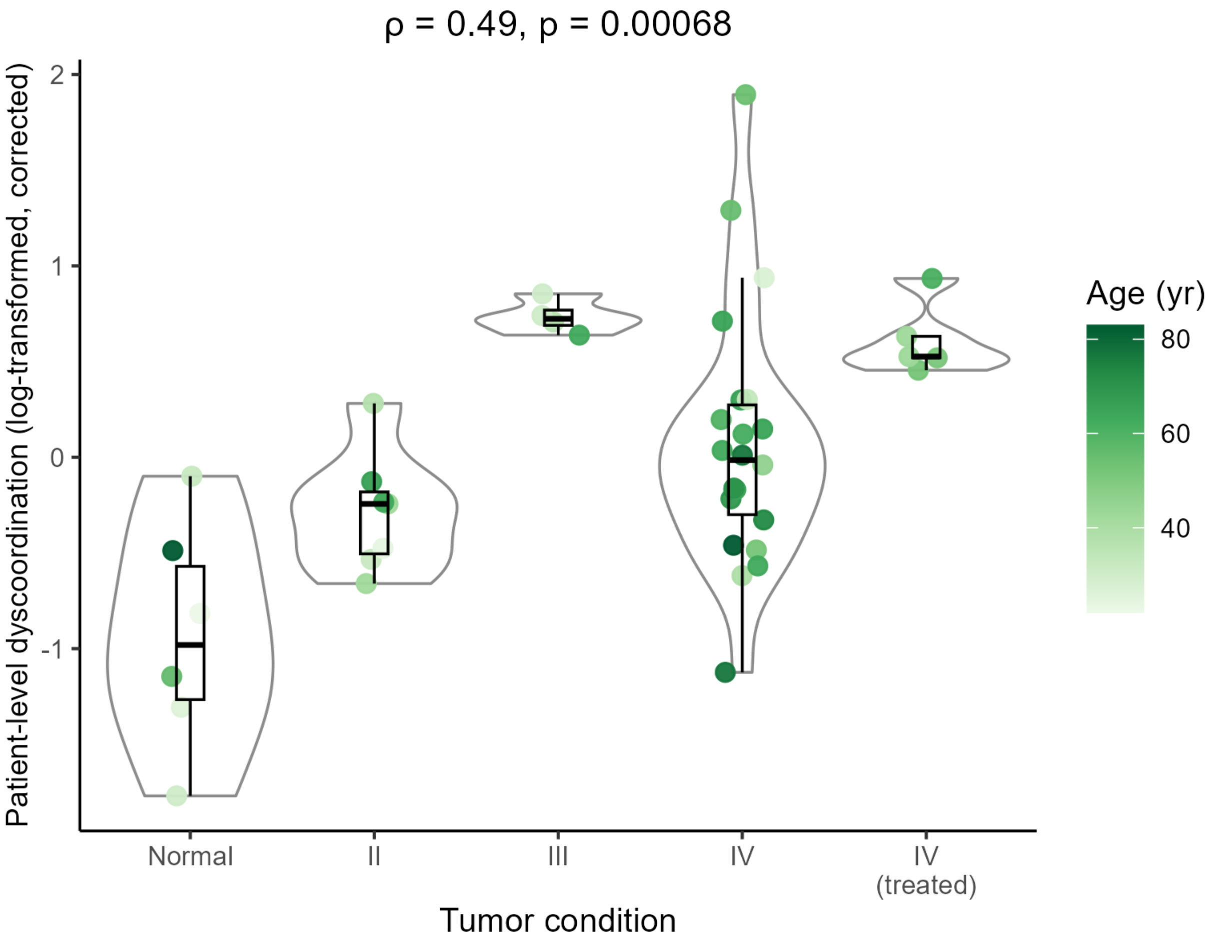


**Supplementary Figure 80 Distribution of patient-level transcriptional dyscoordination across tumor conditions in glioma CD8⁺ T cells.**

Violin and box plots show the distribution of patient-level transcriptional dyscoordination, or the mean of the log-transformed cell-level dyscoordination within each patient, for each tumor condition, after regressing out patient age, patient-mean log-transformed TCR clone size, and patient-averaged cytotoxicity and progenitor/memory gene program scores. Each point represents one patient, with color indicating their age. Spearman's correlation (ρ) and corresponding *p*-value are shown on the top. Gene program scores are per-cell means of z-scored expression averaged per patient, with the following genes surveyed: *GZMB*, *PRF1*, *NKG7*, *GNLY*, *KLRG1* for cytotoxicity programs; *TCF7*, *IL7R*, *CCR7*, *LEF1*, *SELL* for progenitor/memory programs.

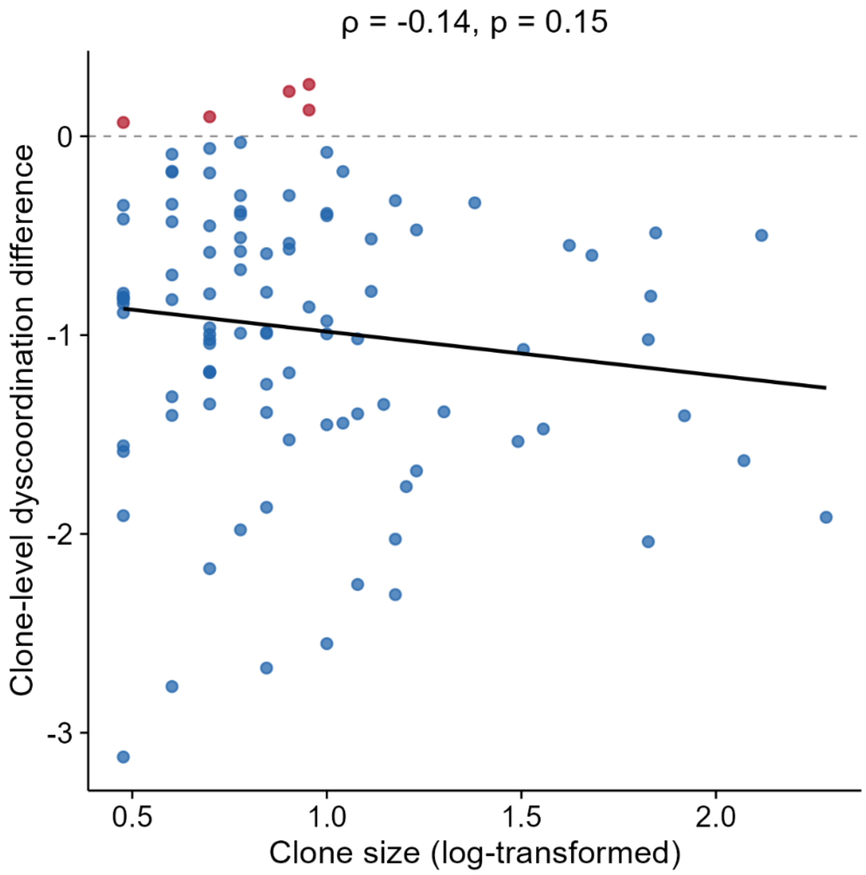


**Supplementary Figure 81 Correlation between clone size and clone-level dyscoordination difference in treated melanoma CD8$^+$ T cells.**

Scatter plot shows the relationship between log-transformed clone size and clone-level dyscoordination difference across TCR clones tracked at both baseline and the first follow-up (week 3). Each point is one clone, colored by the direction of change (blue: decrease; red: increase). The black dashed line marks no change, and the solid line is the linear fit. Spearman's correlation (ρ) and corresponding *p*-value are shown on the top.

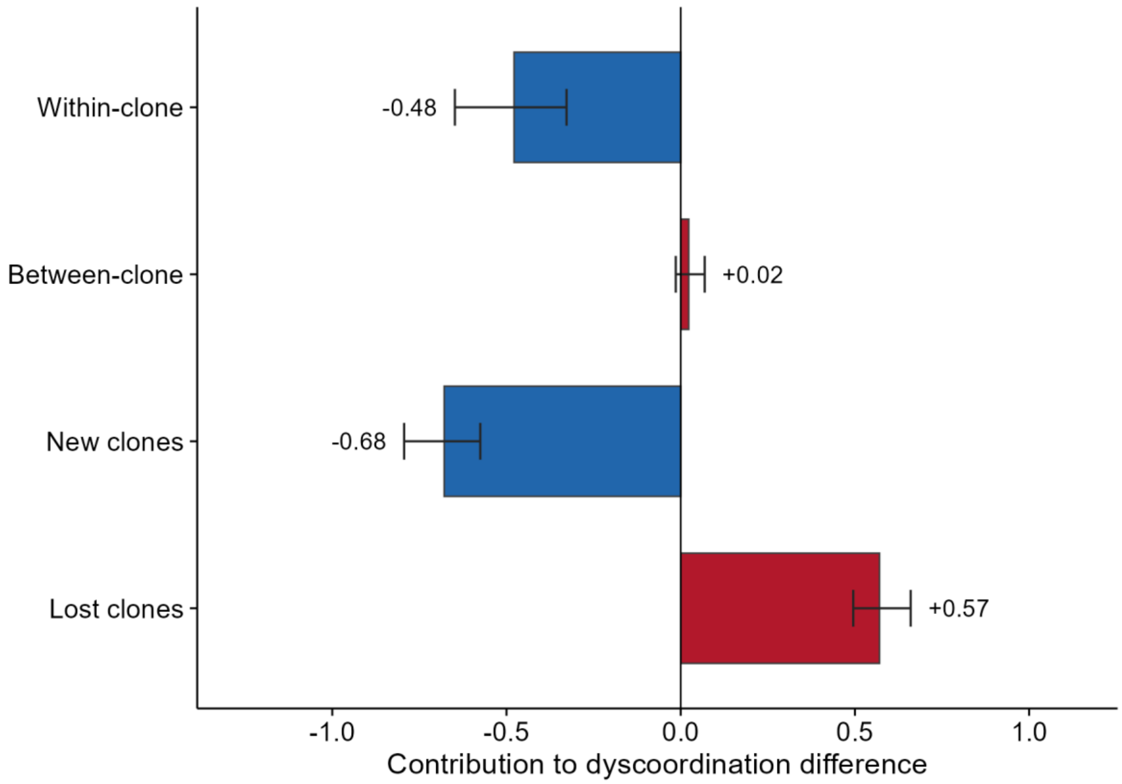


**Supplementary Figure 82 Shift-share decomposition of clone-level dyscoordination difference in treated melanoma CD8$^+$ T cells.**

Bar plots show the shift in the clone-level transcriptional dyscoordination from baseline to the first follow-up (week 3) across TCR clones tracked between the two timepoints, weighted by clone size and decomposed by a Marshall-Edgeworth shift-share into four additive components. Within-clone captures persisting clones changing their own dyscoordination; between-clone captures the re-weighting of persisting clones through expansion or contraction; new clones capture clones present only at follow-up; and lost-clones capture clones present only at baseline. Bars show each component's contribution to the net change of -0.56 log units, colored by sign, with whiskers representing bootstrap 95% confidence intervals.

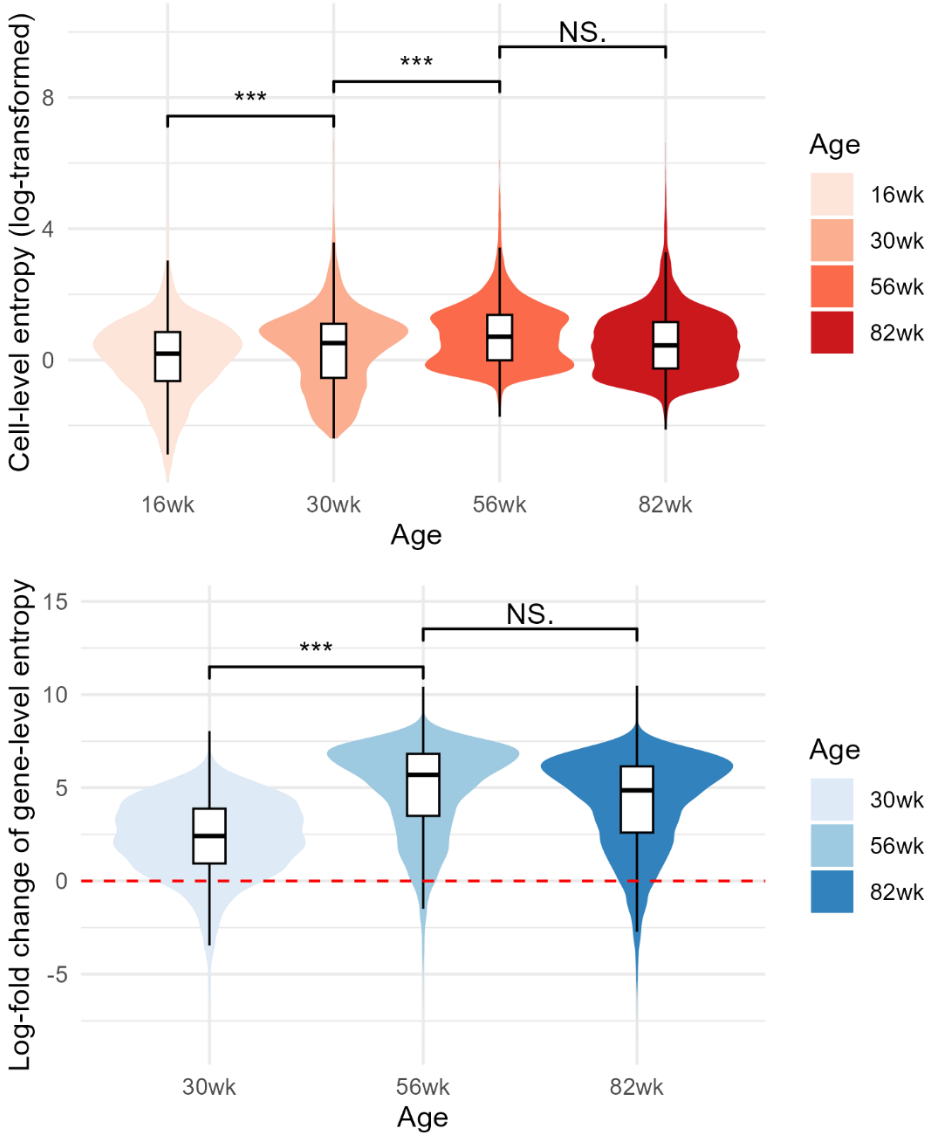


**Supplementary Figure 83 Distribution of transcriptional dyscoordination across rat kidney PT cells of different ages.**

Violin and box plots show the distribution of log-transformed cell-level dyscoordination (top) and log-fold change of gene-level dyscoordination (with respect to 16 weeks, bottom) values separately for each age group. Values in the top and bottom 1% quantiles are considered outliers and are removed from the illustration. Pairwise comparisons between age groups were tested using one-sided Wilcoxon rank-sum test (alternative hypothesis: older age group has higher dyscoordination). Significance levels are annotated above each comparison (NS = not significant, * $p < 0.05$, ** $p < 0.01$, *** $p < 0.001$).

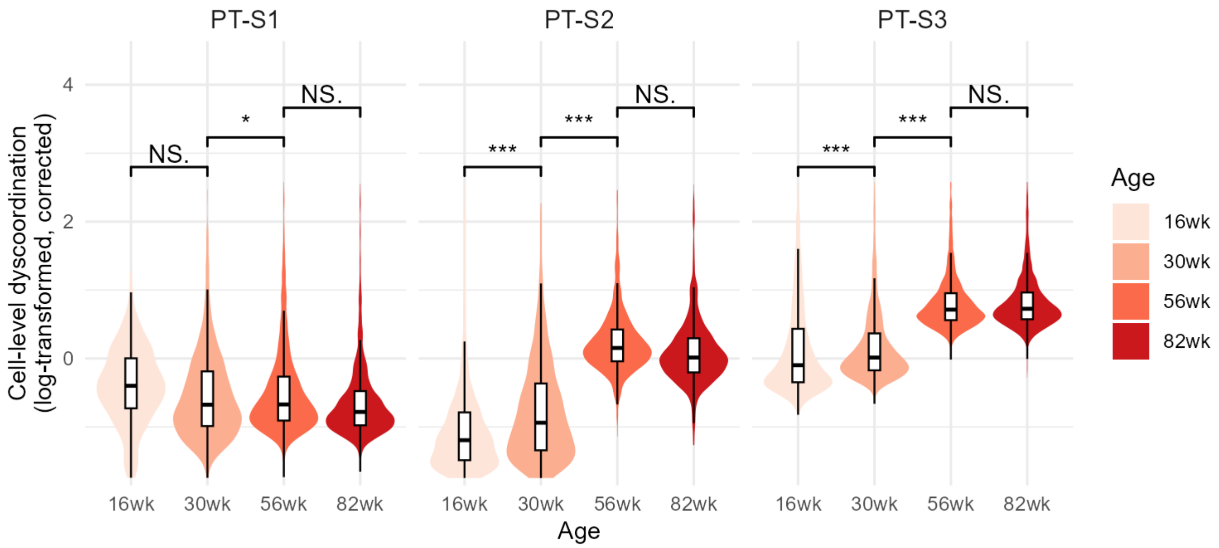


**Supplementary Figure 84 Distribution of corrected cell-level dyscoordination in rat kidney PT cells.**

Violin and box plots show the distribution of log-transformed, depth-corrected cell-level dyscoordination values separately for each cell type and age group. Values in the top and bottom 2.5% quantiles are considered outliers and are removed from the illustration. Pairwise comparisons between age groups were tested using one-sided Wilcoxon rank-sum test (alternative hypothesis: older age group has higher dyscoordination). Significance levels are annotated above each comparison (NS = not significant, * $p < 0.05$, ** $p < 0.01$, *** $p < 0.001$).

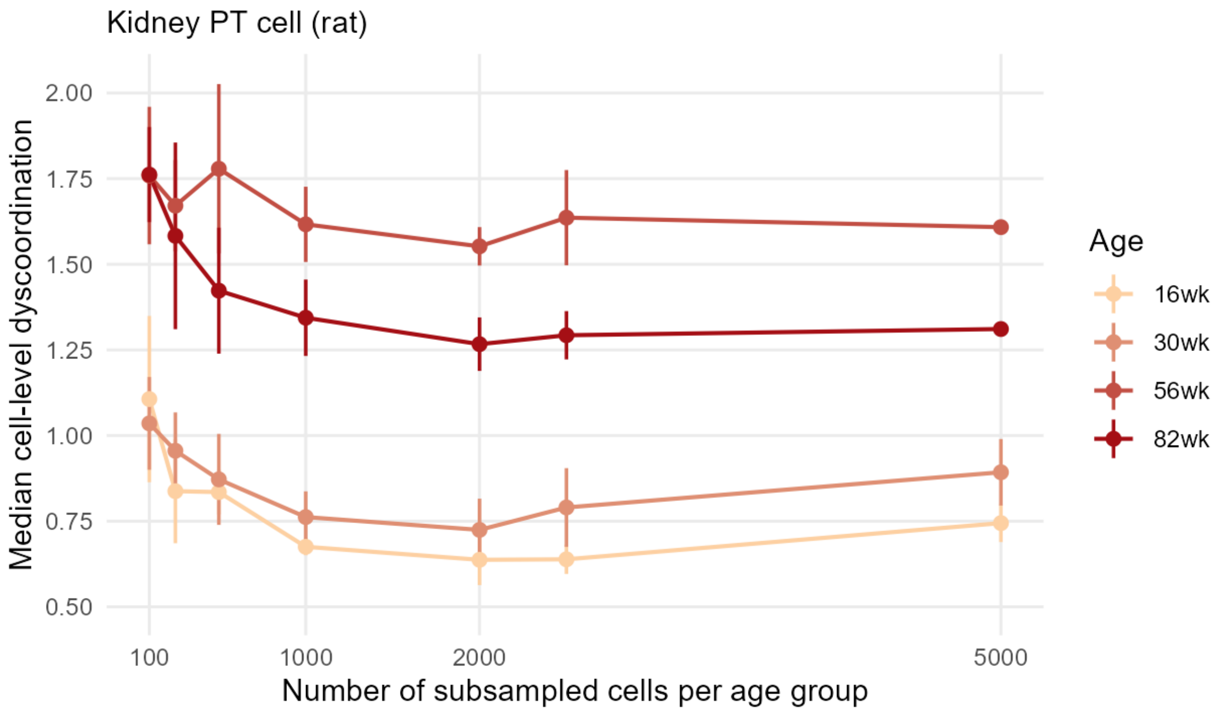


**Supplementary Figure 85 Median cell-level dyscoordination across subsampled age groups in rat kidney PT cells.**

Line plots show the median cell-level dyscoordination for each age group across random subsamples of increasing size (100 to 5,000 cells per age group). Each subsample is summarized by its median value, and each point shows the mean of these medians across 5 independent draws, with vertical lines indicating the 95% confidence interval across draws.

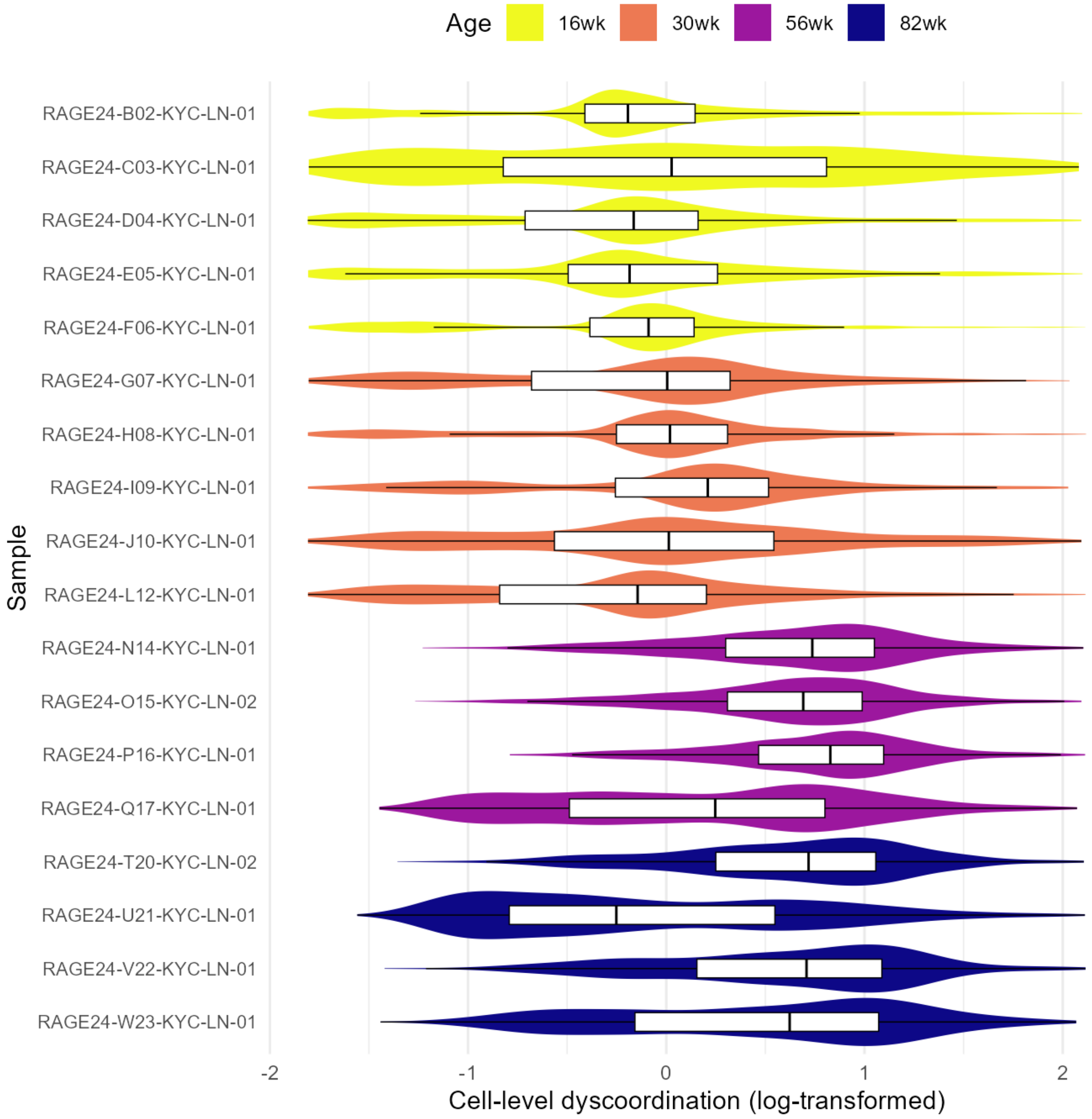


**Supplementary Figure 86 Distribution of cell-level dyscoordination across biological replicates in rat kidney PT cells.**

Violin and box plots show the distribution of log-transformed cell-level dyscoordination for each individual animal, ordered and colored by age group. The distributions are consistent within each age group, showing that the reported trend is not driven by any individual sample or batch.

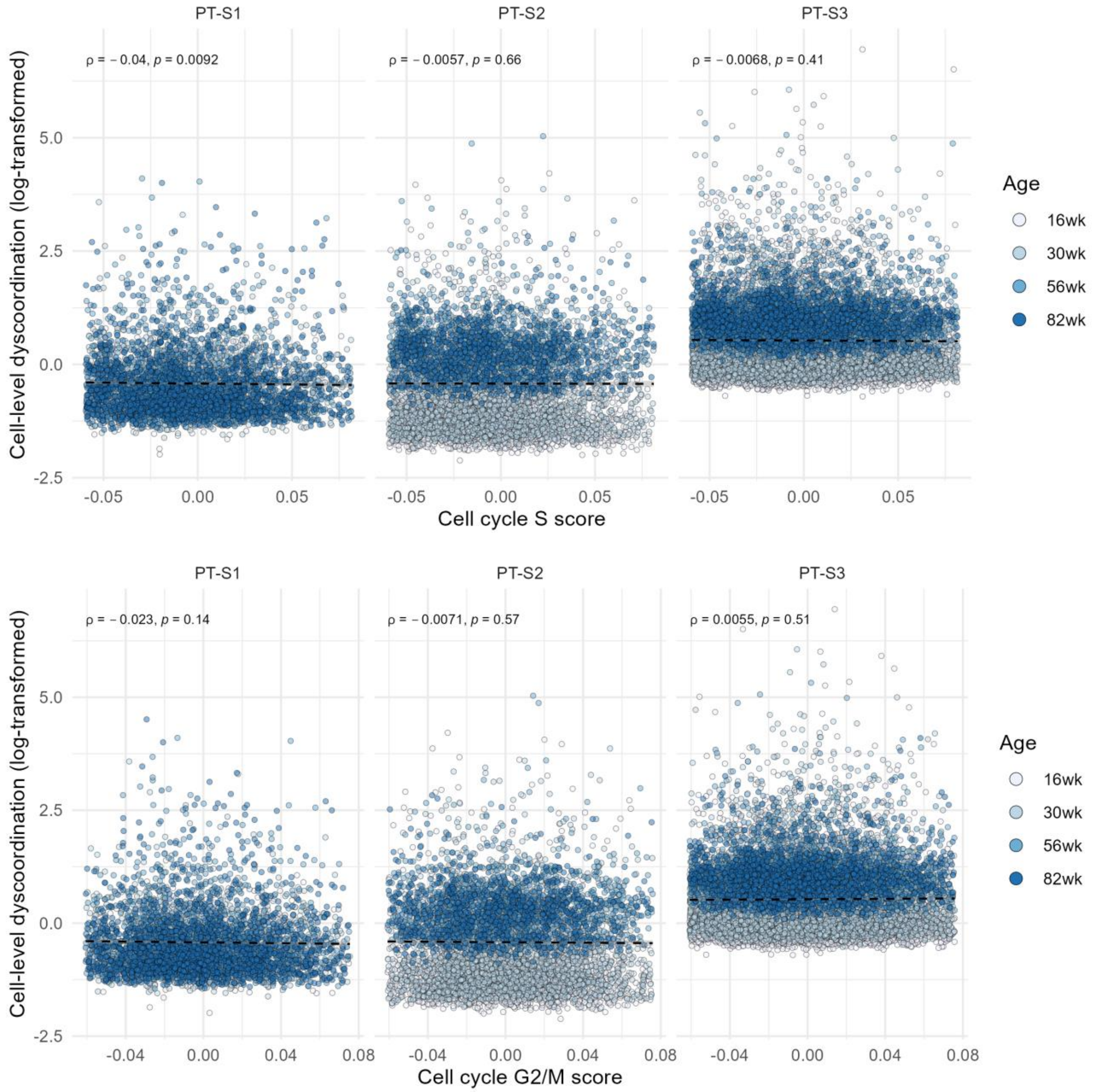


**Supplementary Figure 87 Correlation between cell cycle S phase and G2/M score and cell-level dyscoordination in rat kidney PT cells.**

Scatter plots show the relationship between the cell cycle S phase (top) and G2/M (bottom) score and log-transformed cell-level dyscoordination for each cell type. Black dashed lines indicate linear regression fits. Spearman's correlation coefficients and p-values are shown on the top left corner of each panel.

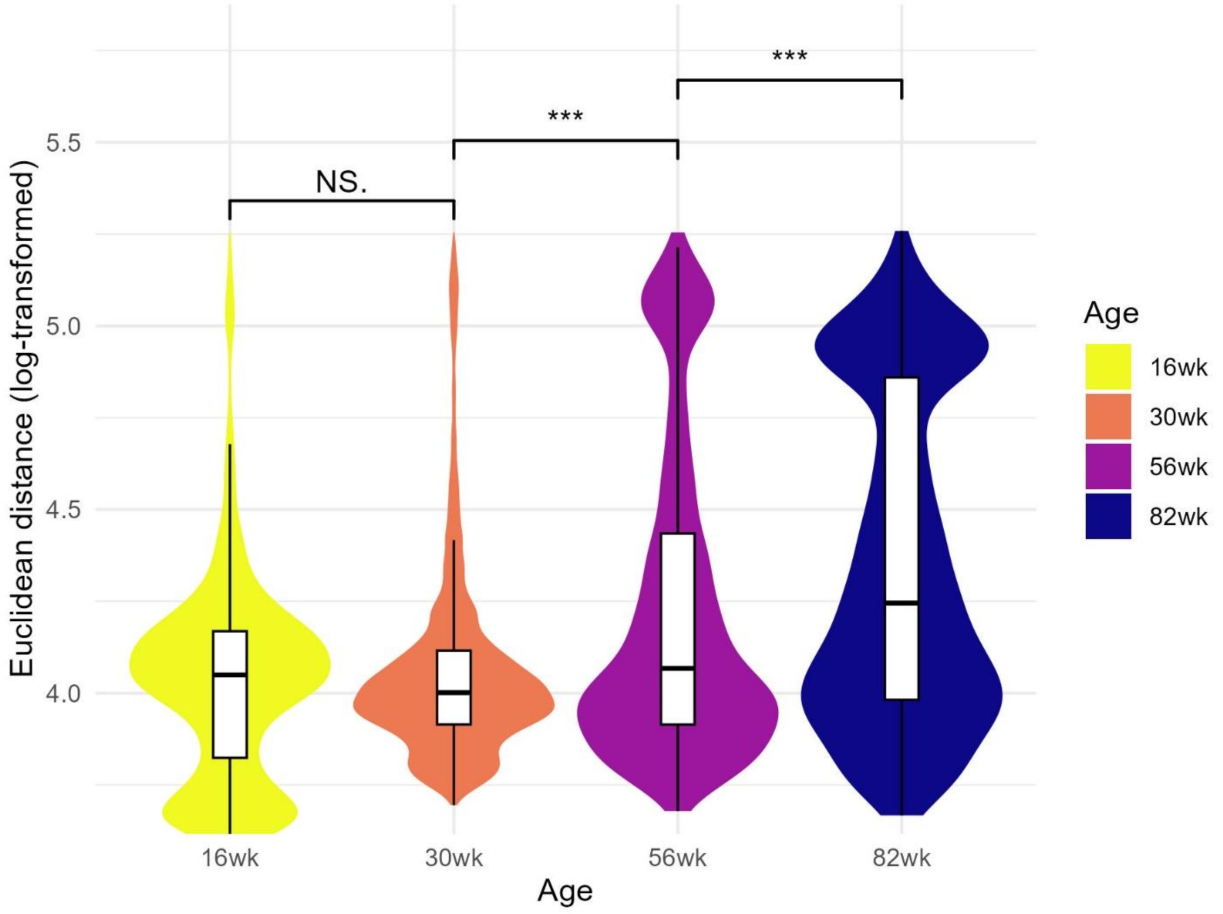


**Supplementary Figure 88 Distribution of Euclidean distance to cell type average in rat kidney PT cells.**

Violin and box plots show the distribution of log-transformed Euclidean distance to cell type average across age groups. Pairwise comparisons between age groups were tested using one-sided Wilcoxon rank-sum test (alternative hypothesis: older age group has higher distance). Significance levels are annotated above each comparison (NS = not significant, * $p < 0.05$, ** $p < 0.01$, *** $p < 0.001$).

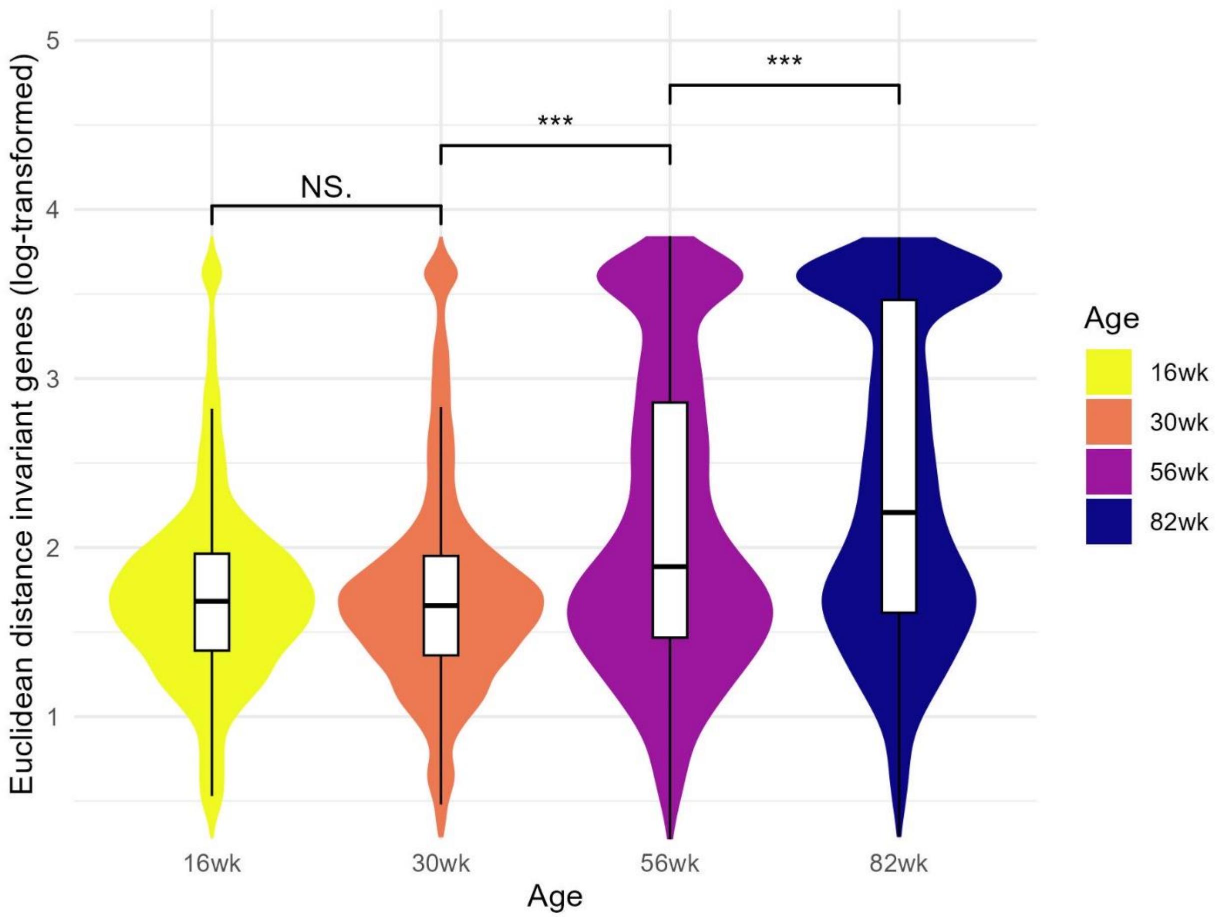


**Supplementary Figure 89 Distribution of Euclidean distance to tissue average using invariant genes in rat kidney PT cells.**

Violin and box plots show the distribution of log-transformed Euclidean distance to tissue average using invariant genes across age groups. Pairwise comparisons between age groups were tested using one-sided Wilcoxon rank-sum test (alternative hypothesis: older age group has higher distance). Significance levels are annotated above each comparison (NS = not significant, * $p < 0.05$, ** $p < 0.01$, *** $p < 0.001$).

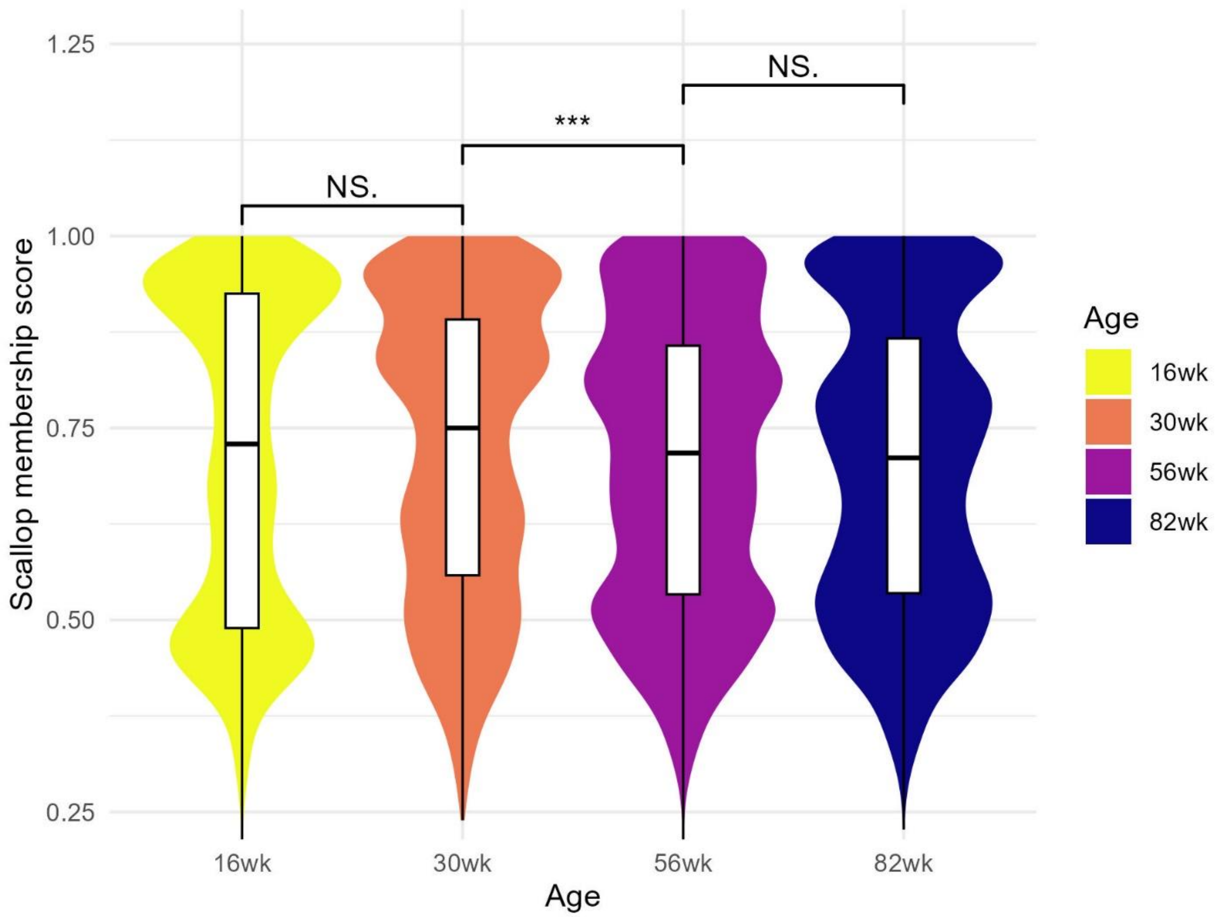


**Supplementary Figure 90 Distribution of Scallop membership score in rat kidney PT cells.**

Violin and box plots show the distribution of Scallop membership score across age groups. Scallop computes it as the frequency with which each cell is assigned to its most frequently assigned cluster. Since this metric reflects transcriptional stability, it is inversely related to our transcriptional dyscoordination measure. Pairwise comparisons between age groups were tested using one-sided Wilcoxon rank-sum test (alternative hypothesis: older age group has lower membership score). Significance levels are annotated above each comparison (NS = not significant, * $p < 0.05$, ** $p < 0.01$, *** $p < 0.001$).

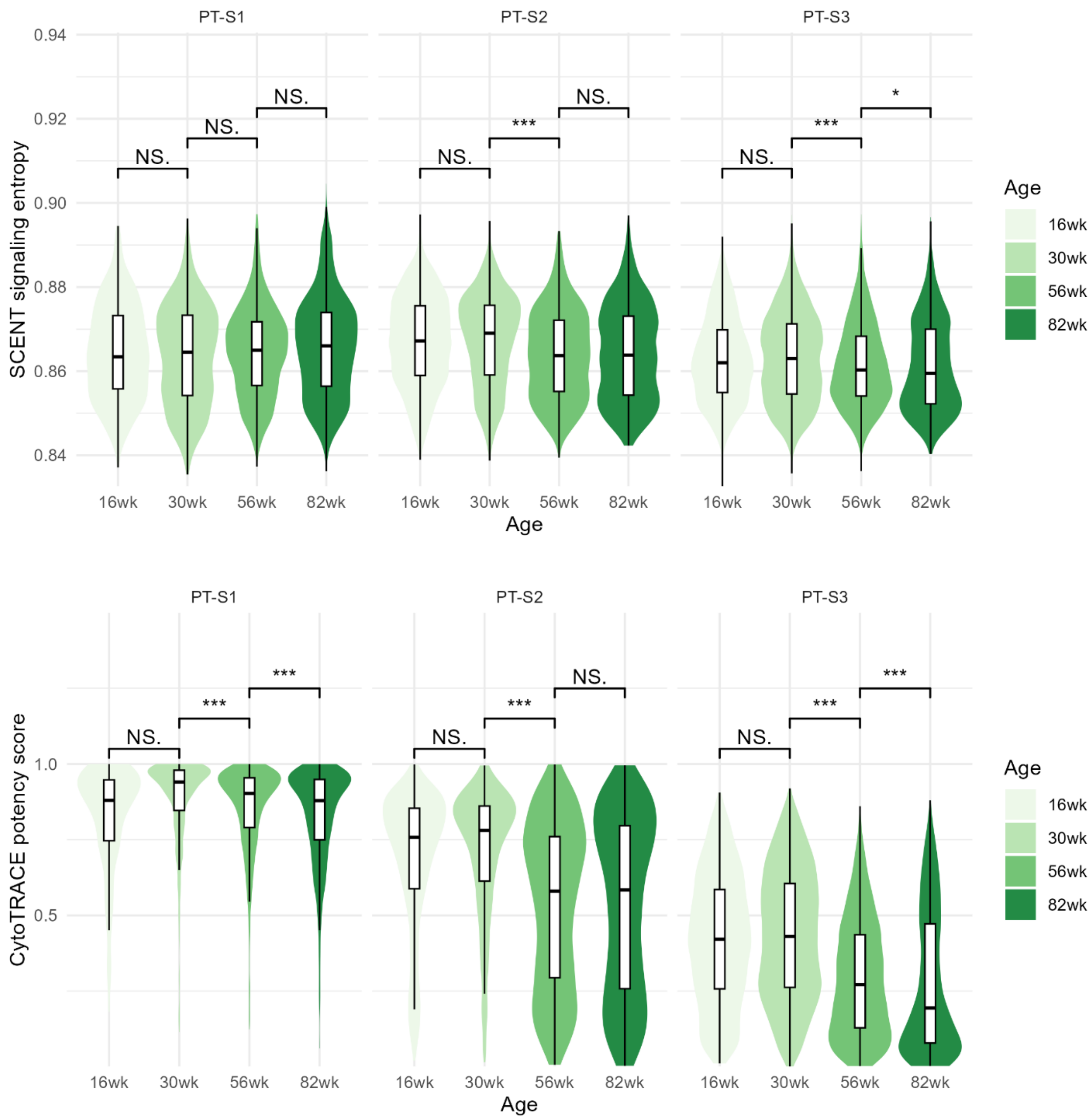


**Supplementary Figure 91 Distribution of SCENT signaling entropy and CytoTRACE potency score in rat kidney PT cells.**

Violin and box plots show the distribution of SCENT signaling entropy and CytoTRACE potency score across rat kidney PT cell subtypes and age groups. Pairwise comparisons between age groups were tested using one-sided Wilcoxon rank-sum test (alternative hypothesis: older age group has lower entropy or potency score). Significance levels are annotated above each comparison (NS = not significant, * $p < 0.05$, ** $p < 0.01$, *** $p < 0.001$).

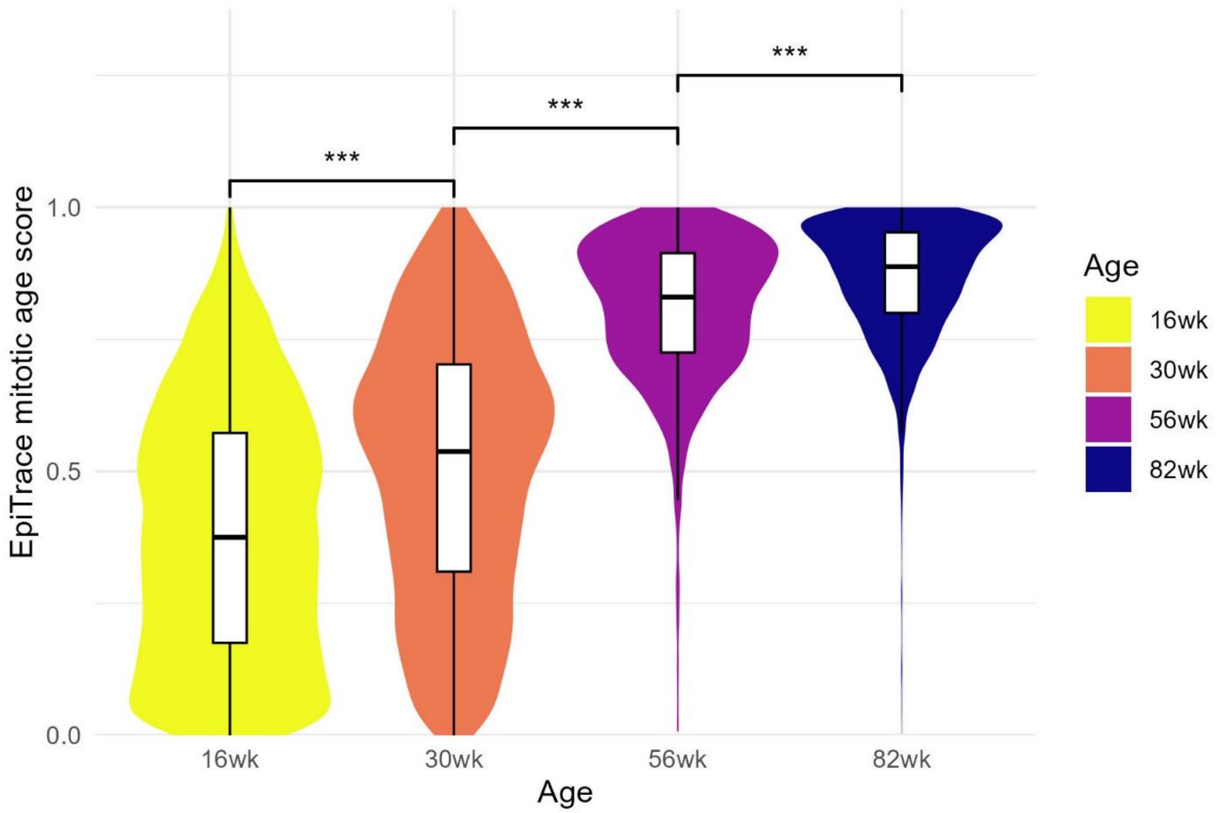


**Supplementary Figure 92 Distribution of EpiTrace mitotic age score in rat kidney PT cells.**

Violin and box plots show the distribution of EpiTrace mitotic age score across age groups. EpiTrace estimates the mitotic age by measuring the total openings of the reference "clock-like" genomic loci in single-cell ATAC data. Pairwise comparisons between age groups were tested using one-sided Wilcoxon rank-sum test (alternative hypothesis: older age group has higher EpiTrace score). Significance levels are annotated above each comparison (NS = not significant, * $p < 0.05$, ** $p < 0.01$, *** $p < 0.001$).

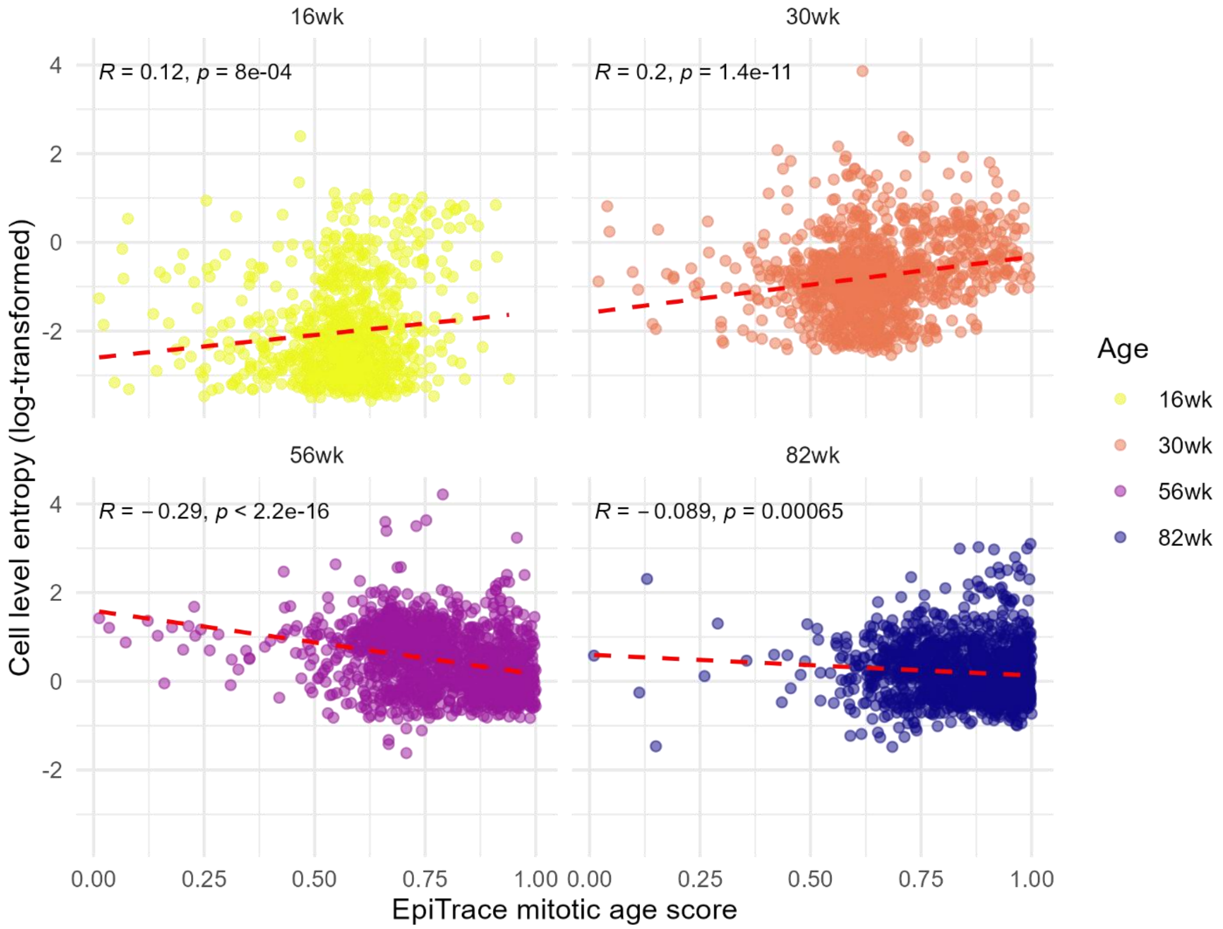


**Supplementary Figure 93 Correlation between EpiTrace mitotic age score and cell-level dyscoordination in rat kidney PT cells.**

Scatter plots show the relationship between EpiTrace mitotic age score and log-transformed cell-level dyscoordination across age groups. Red dashed lines indicate linear regression fits. Pearson correlation coefficients and corresponding p-values are shown in the top left corner of each panel. Significant positive correlations were primarily observed in 16-week and 30-week PT cells.

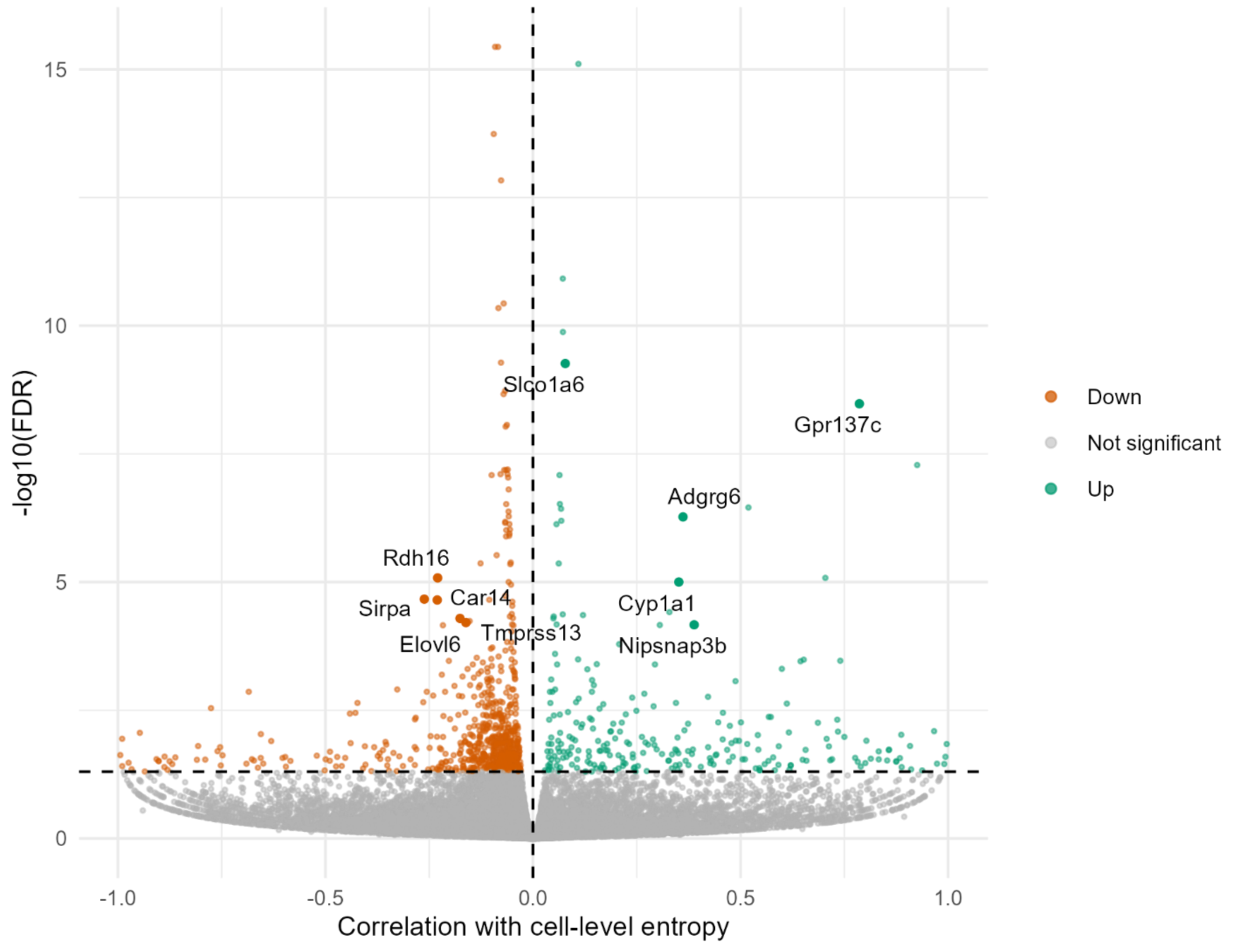


**Supplementary Figure 94 Correlation of gene expression with cell-level dyscoordination in rat kidney PT cells.**

Volcano plots show the correlation of expression levels with cell-level dyscoordination for each individual gene. Positively and negatively correlated genes are separately colored. Top 5 significantly correlated genes in each direction with -log10(FDR) > 4 are highlighted.

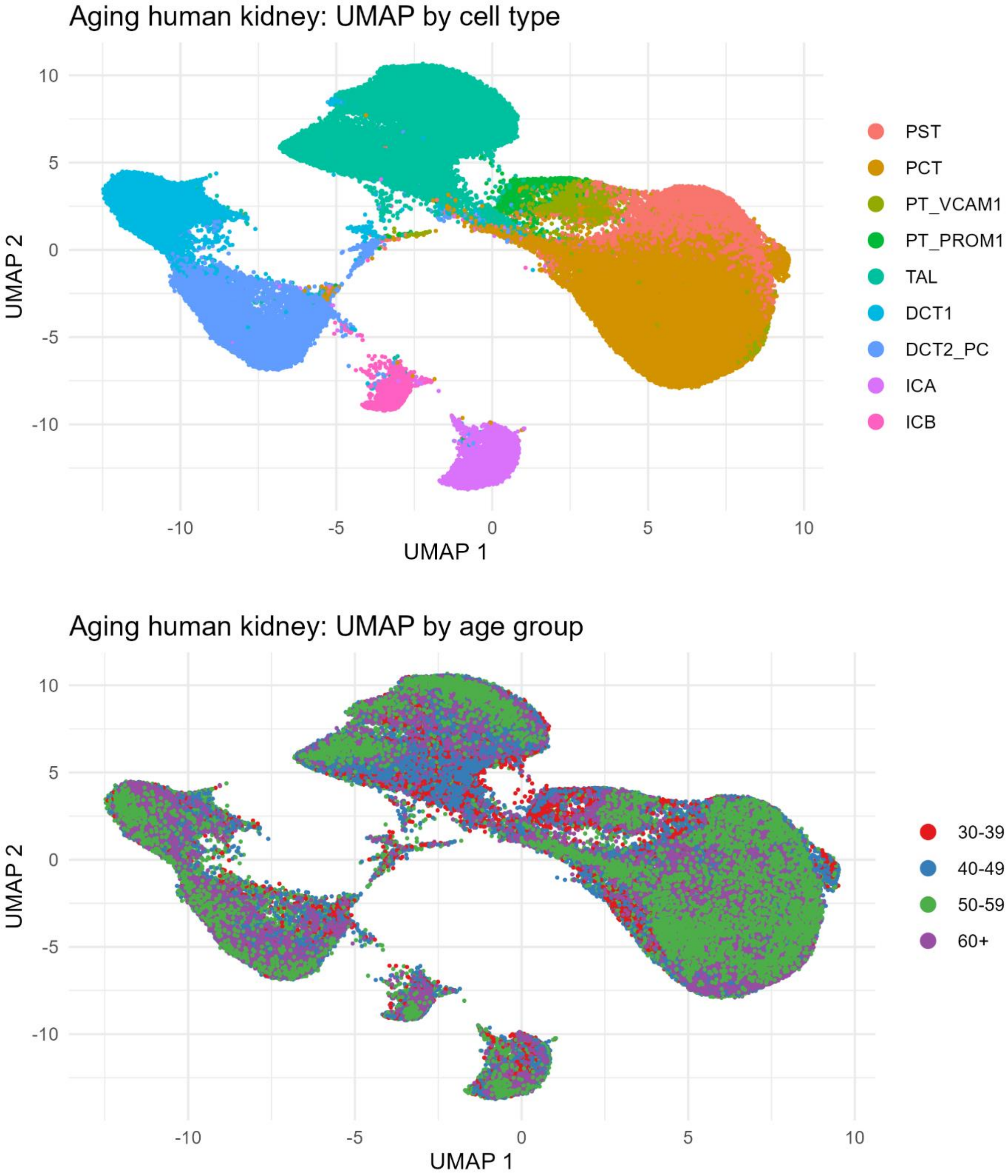


**Supplementary Figure 95 UMAP visualization of epithelial cells from aging human kidney.**

Cells are embedded using the first two UMAP dimensions and colored by annotated cell type (top) and age group (bottom).

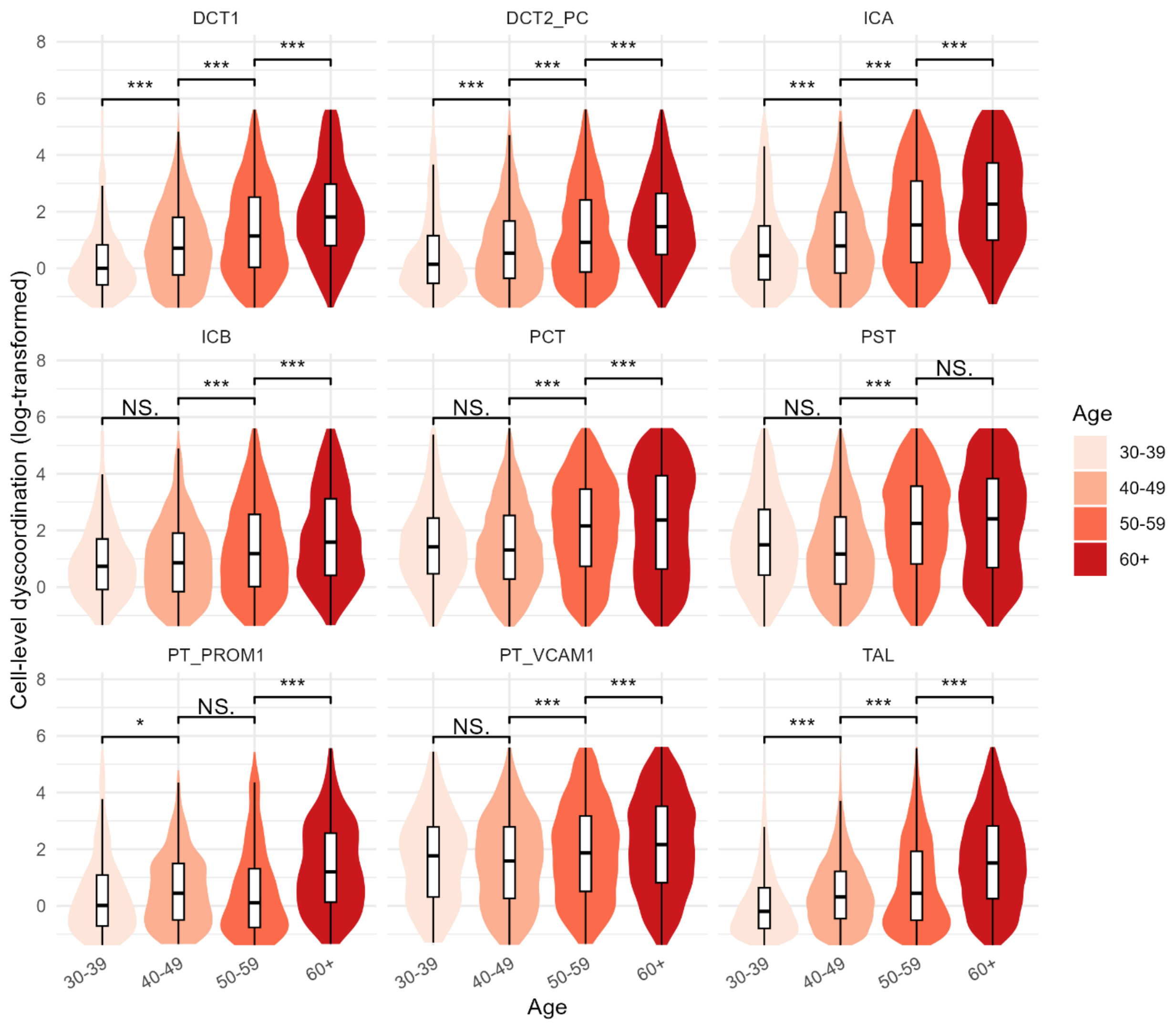


**Supplementary Figure 96 Distribution of cell-level dyscoordination in human kidney epithelial cells.**

Violin and box plots show the distribution of log-transformed cell-level dyscoordination values separately for each cell type and age group. Values in the top and bottom 2.5% quantiles are considered outliers and are removed from the illustration. Pairwise comparisons between age groups were tested using one-sided Wilcoxon rank-sum test (alternative hypothesis: older age group has higher dyscoordination). Significance levels are annotated above each comparison (NS = not significant, * $p < 0.05$, ** $p < 0.01$, *** $p < 0.001$).

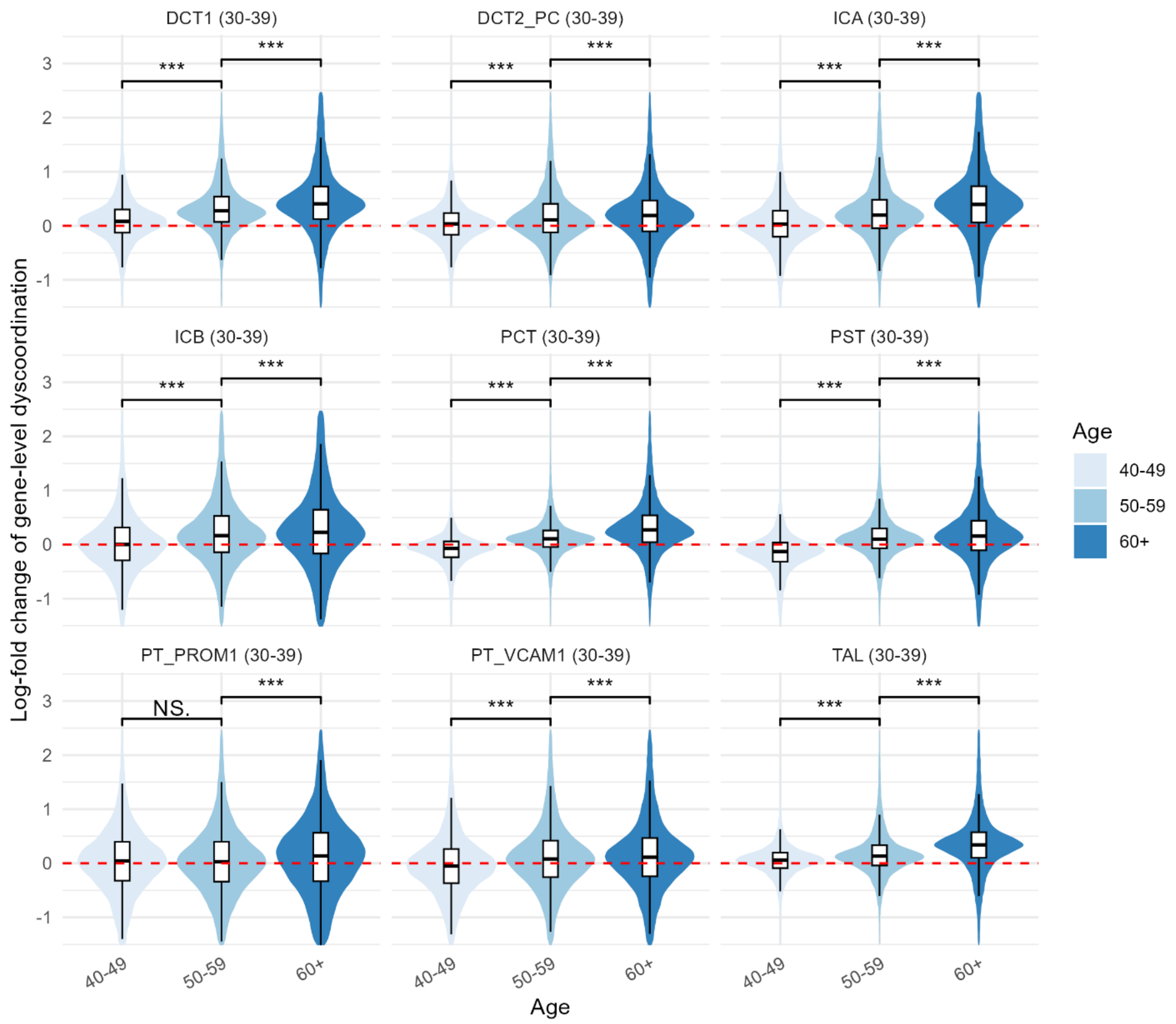


**Supplementary Figure 97 Distribution of gene-level dyscoordination in human kidney epithelial cells.**

Violin and box plots show the distribution of log-fold changes in dyscoordination relative to the baseline age group. Values are displayed separately for each cell type and age group, with the baseline age group indicated in parentheses in each subplot title. Values in the top and bottom 2.5% quantiles are considered outliers and are removed from the illustration. Red dashed line denotes zero log-fold change. Pairwise comparisons between age groups were tested using one-sided Wilcoxon rank-sum test (alternative hypothesis: older age group has higher dyscoordination). Significance levels are annotated above each comparison (NS = not significant, * $p < 0.05$, ** $p < 0.01$, *** $p < 0.001$).

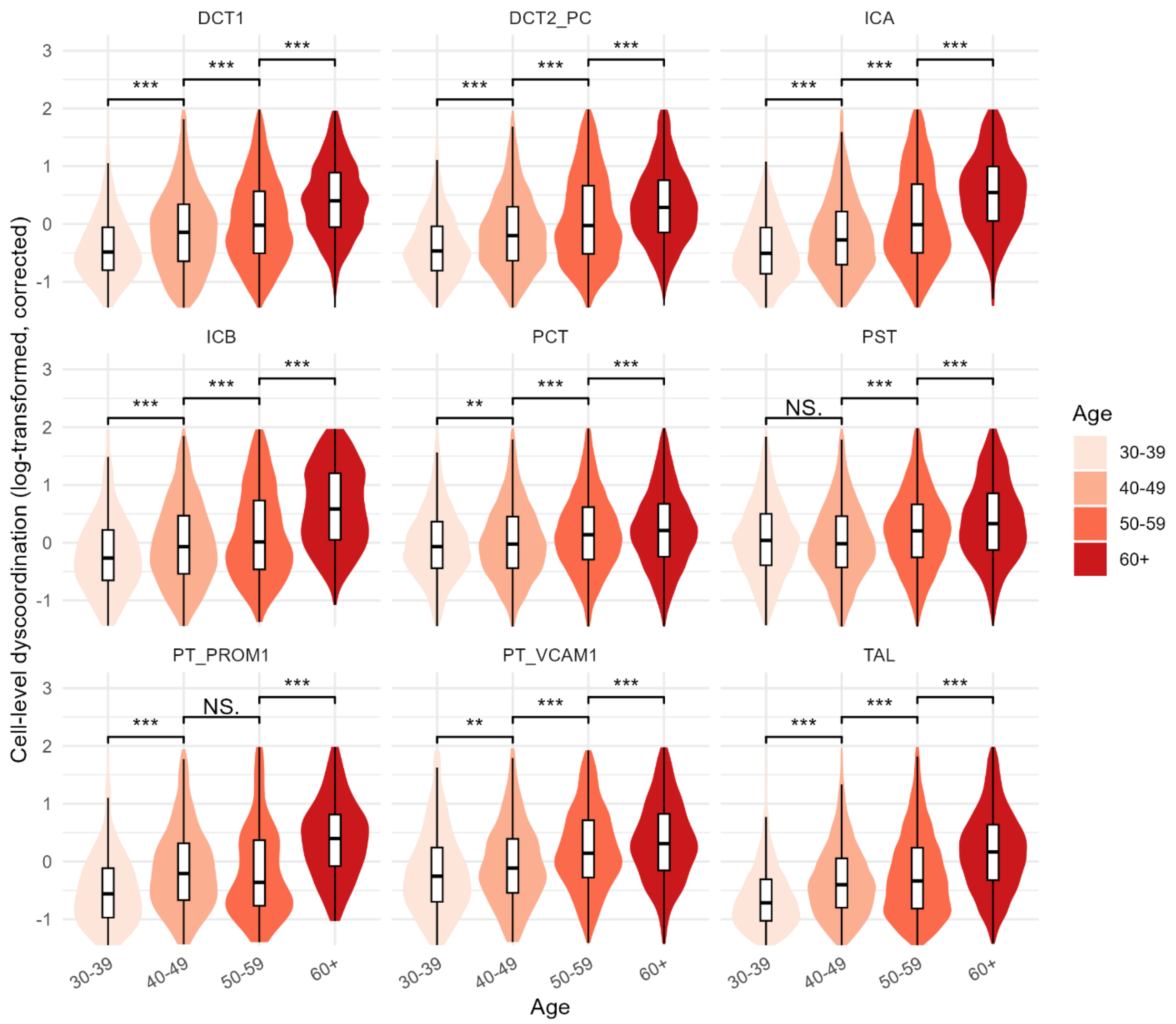


**Supplementary Figure 98 Distribution of corrected cell-level dyscoordination in human kidney epithelial cells.**

Violin and box plots show the distribution of log-transformed, depth-corrected cell-level dyscoordination values separately for each cell type and age group. Values in the top and bottom 2.5% quantiles are considered outliers and are removed from the illustration. Pairwise comparisons between age groups were tested using one-sided Wilcoxon rank-sum test (alternative hypothesis: older age group has higher dyscoordination). Significance levels are annotated above each comparison (NS = not significant, * $p < 0.05$, ** $p < 0.01$, *** $p < 0.001$).

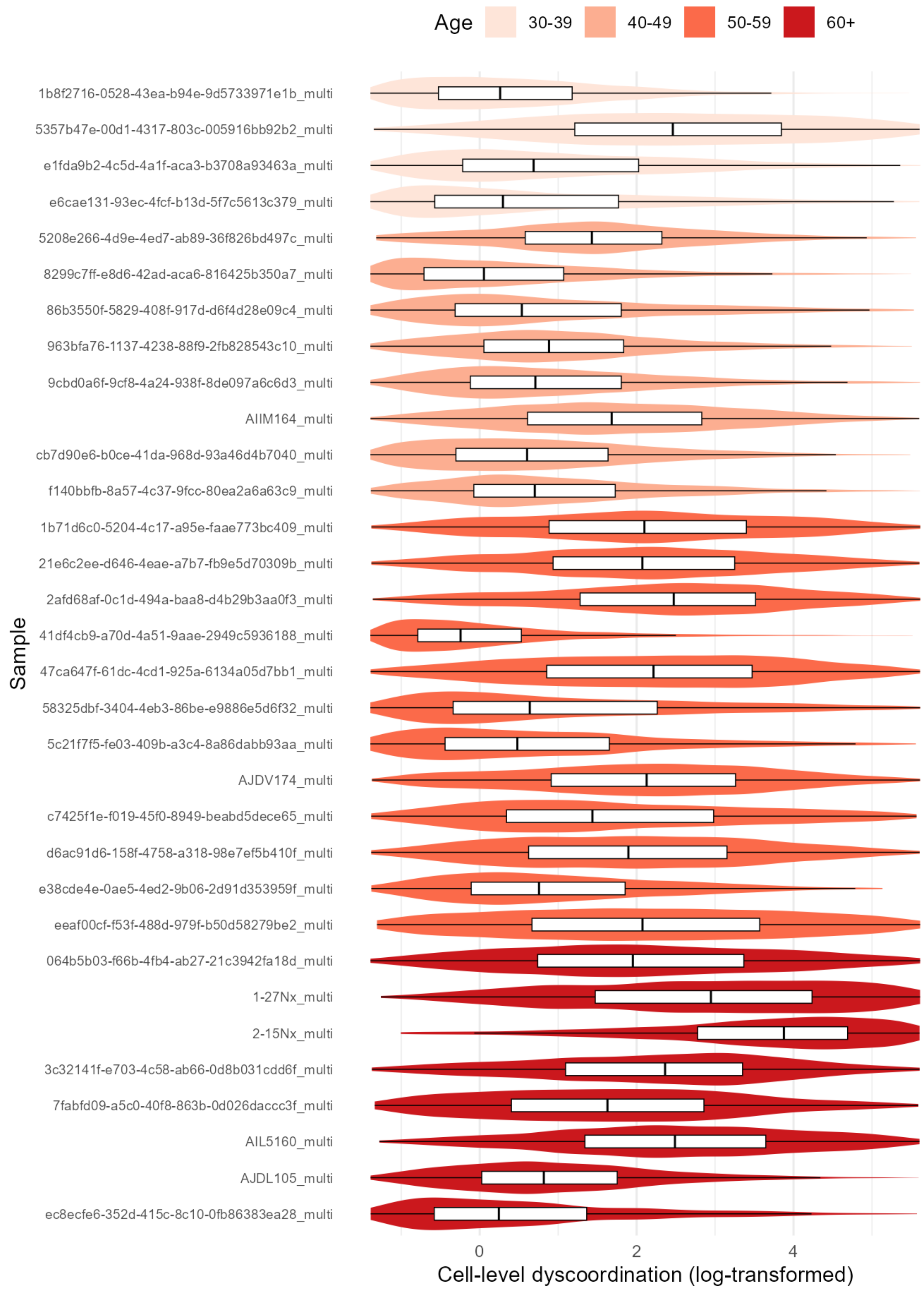
Age
30-39
40-49
50-59
60+
Sample
1b8f2716-0528-43ea-b94e-9d5733971e1b_multi
5357b47e-00d1-4317-803c-005916bb92b2_multi
e1fda9b2-4c5d-4a1f-aca3-b3708a93463a_multi
e6cae131-93ec-4fcf-b13d-5f7c5613c379_multi
5208e266-4d9e-4ed7-ab89-36f826bd497c_multi
8299c7ff-e8d6-42ad-aca6-816425b350a7_multi
86b3550f-5829-408f-917d-d6f4d28e09c4_multi
963bfa76-1137-4238-88f9-2fb828543c10_multi
9cbd0a6f-9cf8-4a24-938f-8de097a6c6d3_multi
AIIM164_multi
cb7d90e6-b0ce-41da-968d-93a46d4b7040_multi
f140bbfb-8a57-4c37-9fcc-80ea2a6a63c9_multi
1b71d6c0-5204-4c17-a95e-faae773bc409_multi
21e6c2ee-d646-4eae-a7b7-fb9e5d70309b_multi
2afd68af-0c1d-494a-baa8-d4b29b3aa0f3_multi
41df4cb9-a70d-4a51-9aae-2949c5936188_multi
47ca647f-61dc-4cd1-925a-6134a05d7bb1_multi
58325dbf-3404-4eb3-86be-e9886e5d6f32_multi
5c21f7f5-fe03-409b-a3c4-8a86dabb93aa_multi
AJDV174_multi
c7425f1e-f019-45f0-8949-beabd5dece65_multi
d6ac91d6-158f-4758-a318-98e7ef5b410f_multi
e38cde4e-0ae5-4ed2-9b06-2d91d353959f_multi
eeaf00cf-f53f-488d-979f-b50d58279be2_multi
064b5b03-f66b-4fb4-ab27-21c3942fa18d_multi
1-27Nx_multi
2-15Nx_multi
3c32141f-e703-4c58-ab66-0d8b031cdd6f_multi
7fabfd09-a5c0-40f8-863b-0d026daccc3f_multi
AIL5160_multi
AJDL105_multi
ec8ecfe6-352d-415c-8c10-0fb86383ea28_multi
0
2
4
Cell-level dyscoordination (log-transformed)

**Supplementary Figure 99 Distribution of cell-level dyscoordination across biological replicates in human kidney epithelial cells.**

Violin and box plots show the distribution of log-transformed cell-level dyscoordination for each individual sample, ordered and colored by age group. Values in the top and bottom 2.5% quantiles are considered outliers and are removed from the illustration.

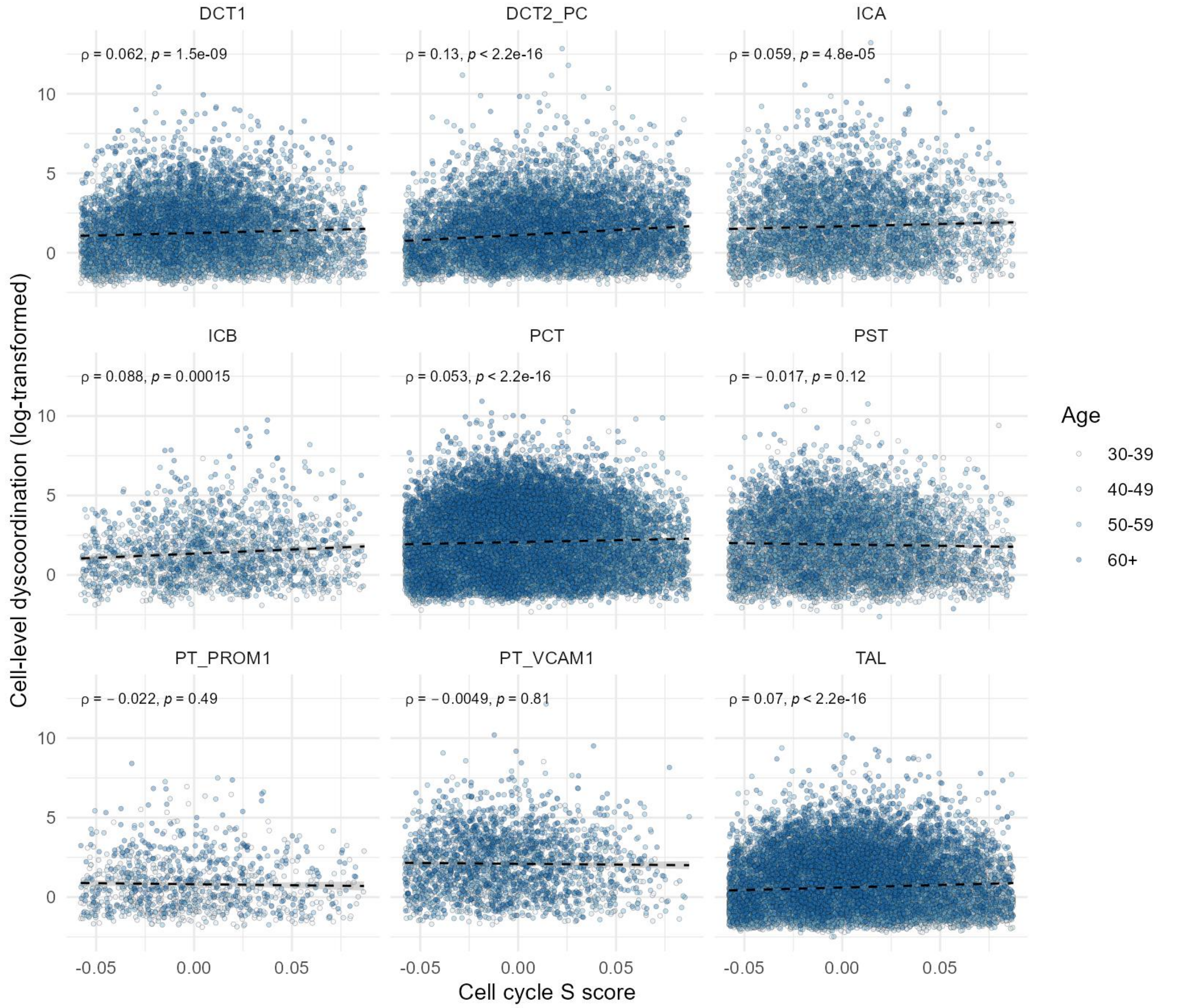


**Supplementary Figure 100 Correlation between cell cycle S phase score and cell-level dyscoordination in human kidney epithelial cells.**

Scatter plots show the relationship between the cell cycle S phase score and log-transformed cell-level dyscoordination for each cell type. Black dashed lines indicate linear regression fits. Spearman's correlation coefficients and p-values are shown on the top left corner of each panel.

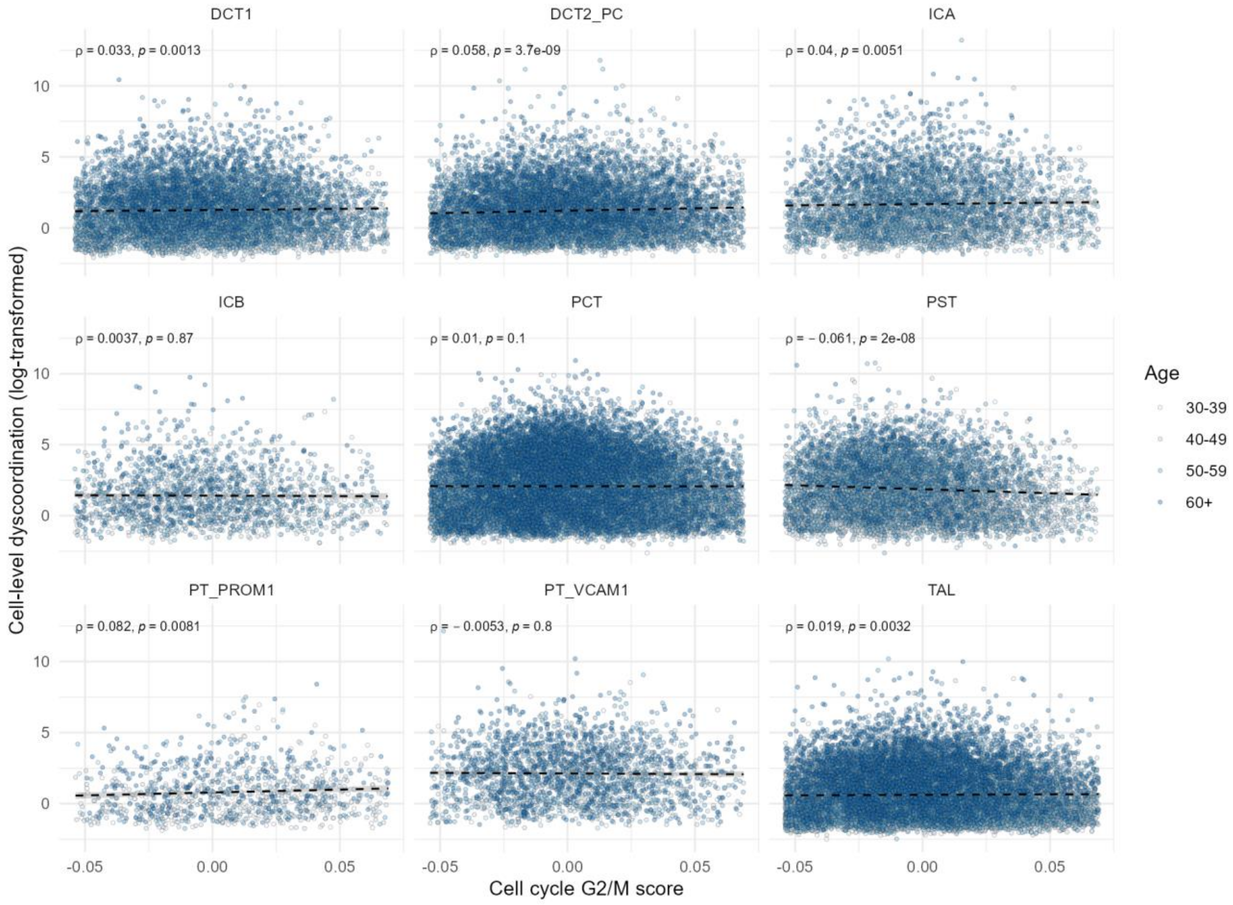


**Supplementary Figure 101 Correlation between cell-cycle G2/M score and cell-level dyscoordination in human kidney epithelial cells.**

Scatter plots show the relationship between the cell cycle G2/M phase score and log-transformed cell-level dyscoordination for each cell type. Black dashed lines indicate linear regression fits. Spearman's correlation coefficients and p-values are shown on the top left corner of each panel.

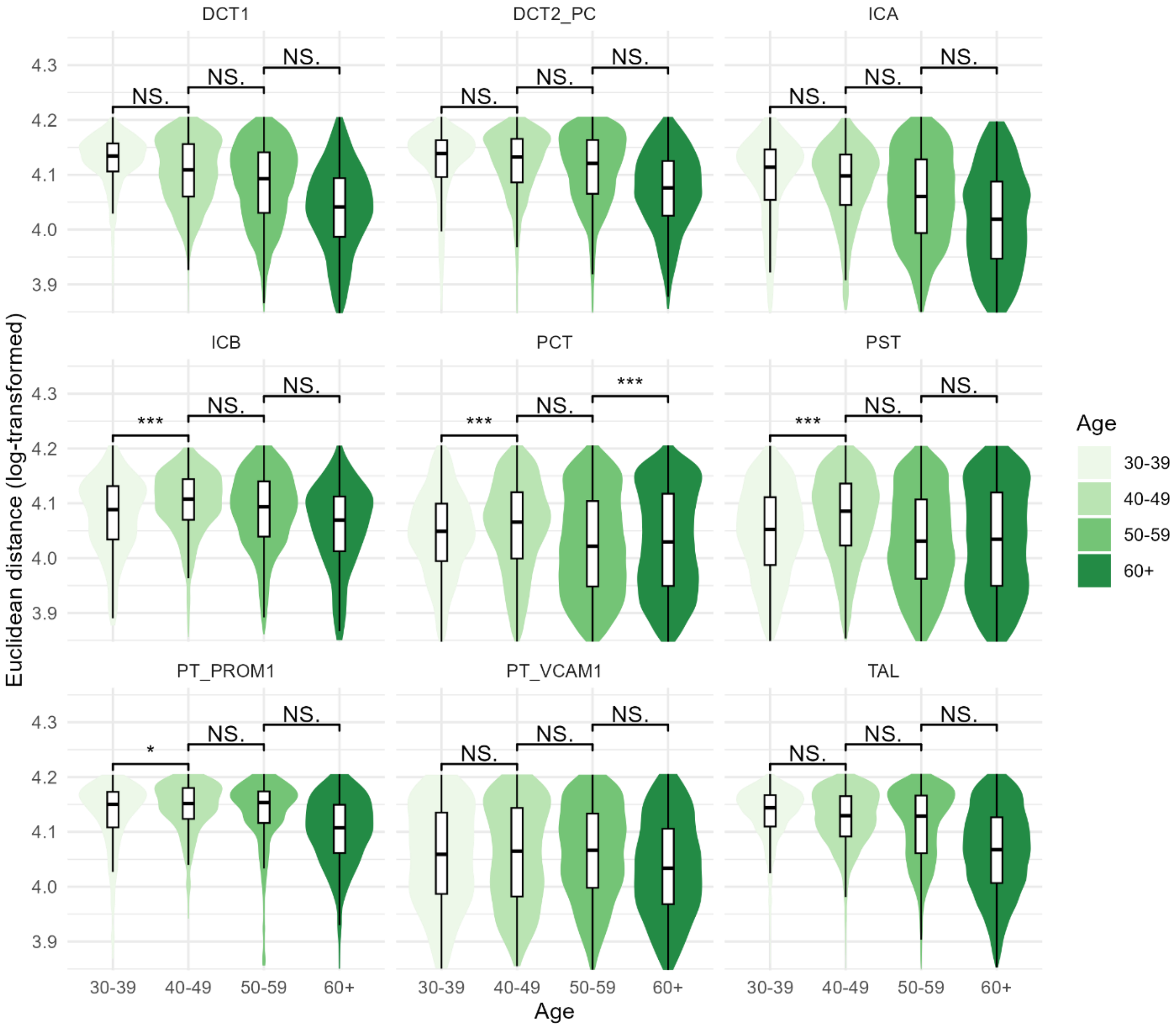


**Supplementary Figure 102 Distribution of Euclidean distance to cell type average in human kidney epithelial cells.**

Violin and box plots show the distribution of log-transformed Euclidean distance to cell type average across human kidney epithelial cell types and age groups. This method is adopted from decibel R package, which computes the Euclidean distance between each cell and the mean expression of its cell type across all genes. Pairwise comparisons between age groups were tested using one-sided Wilcoxon rank-sum test (alternative hypothesis: older age group has higher distance). Significance levels are annotated above each comparison (NS = not significant, * $p < 0.05$, ** $p < 0.01$, *** $p < 0.001$).

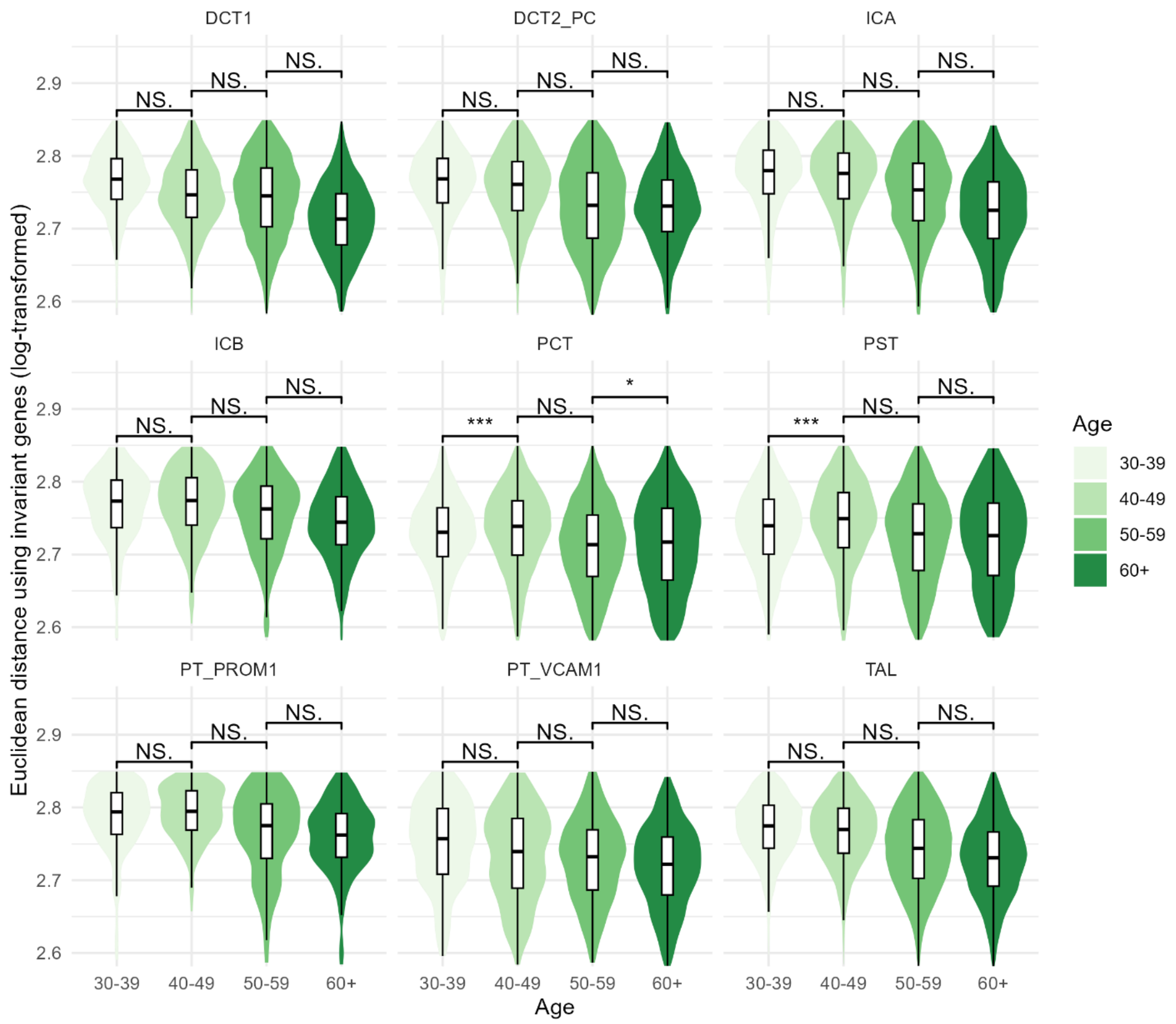


**Supplementary Figure 103 Distribution of Euclidean distance to tissue average using invariant genes in human kidney epithelial cells.**

Violin and box plots show the distribution of log-transformed Euclidean distance to tissue average using invariant genes across human kidney epithelial cell types and age groups. This method is adopted from decibel R package, which computes the Euclidean distance between each cell and the mean expression of all cells in the tissue across all genes using a set of invariant genes. Details for how invariant genes are selected can be found in the package documentation. Pairwise comparisons between age groups were tested using one-sided Wilcoxon rank-sum test (alternative hypothesis: older age group has higher distance). Significance levels are annotated above each comparison (NS = not significant, * $p < 0.05$, ** $p < 0.01$, *** $p < 0.001$)

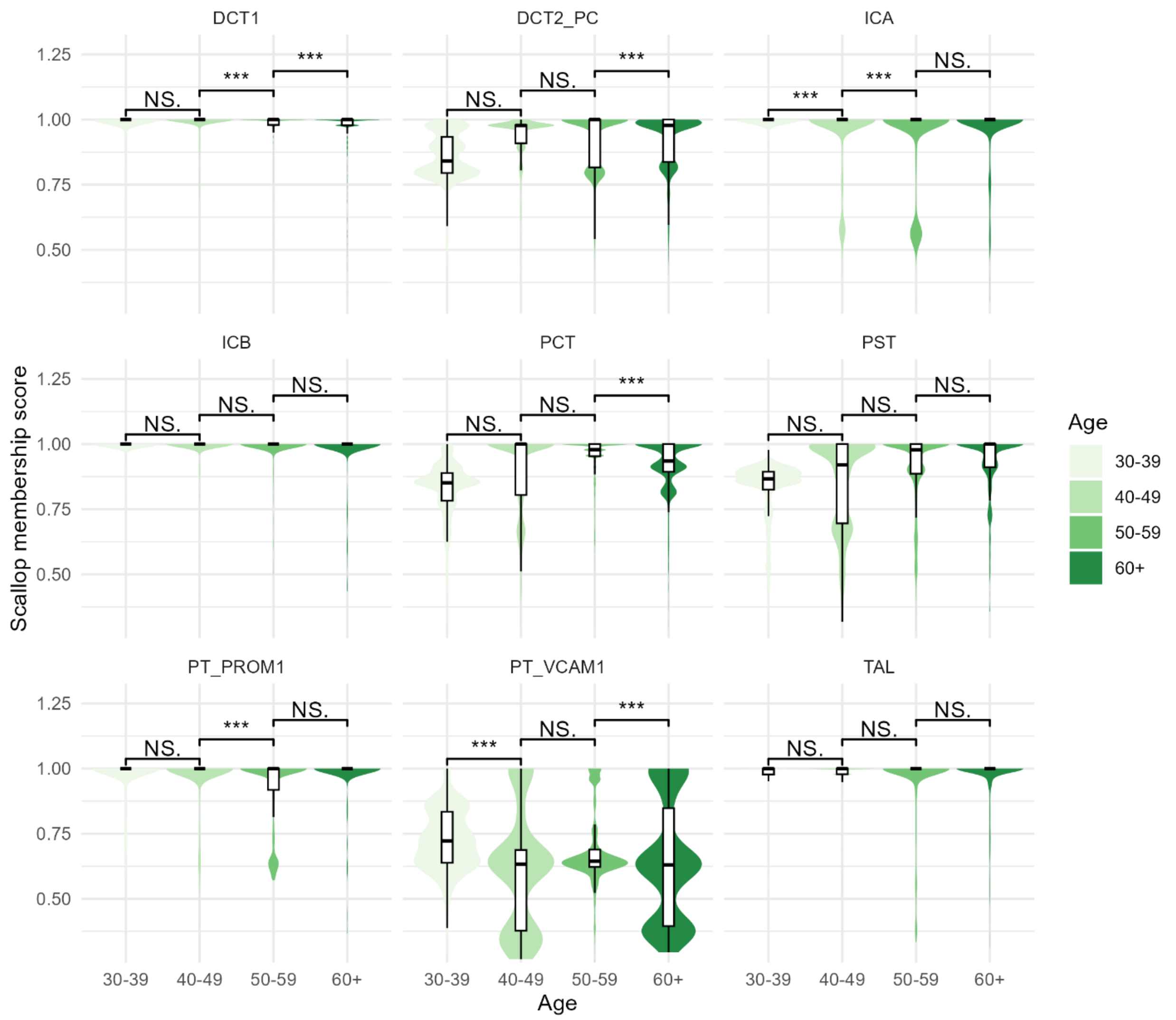


**Supplementary Figure 104 Distribution of Scallop membership score in human kidney epithelial cells.**

Violin and box plots show the distribution of Scallop membership score across human kidney epithelial cell types and age groups. Scallop computes it as the frequency with which each cell is assigned to its most frequently assigned cluster. Since this metric reflects transcriptional stability, it is inversely related to our transcriptional dyscoordination measure. Pairwise comparisons between age groups were tested using one-sided Wilcoxon rank-sum test (alternative hypothesis: older age group has lower membership score). Significance levels are annotated above each comparison (NS = not significant, * p < 0.05, ** p < 0.01, *** p < 0.001).

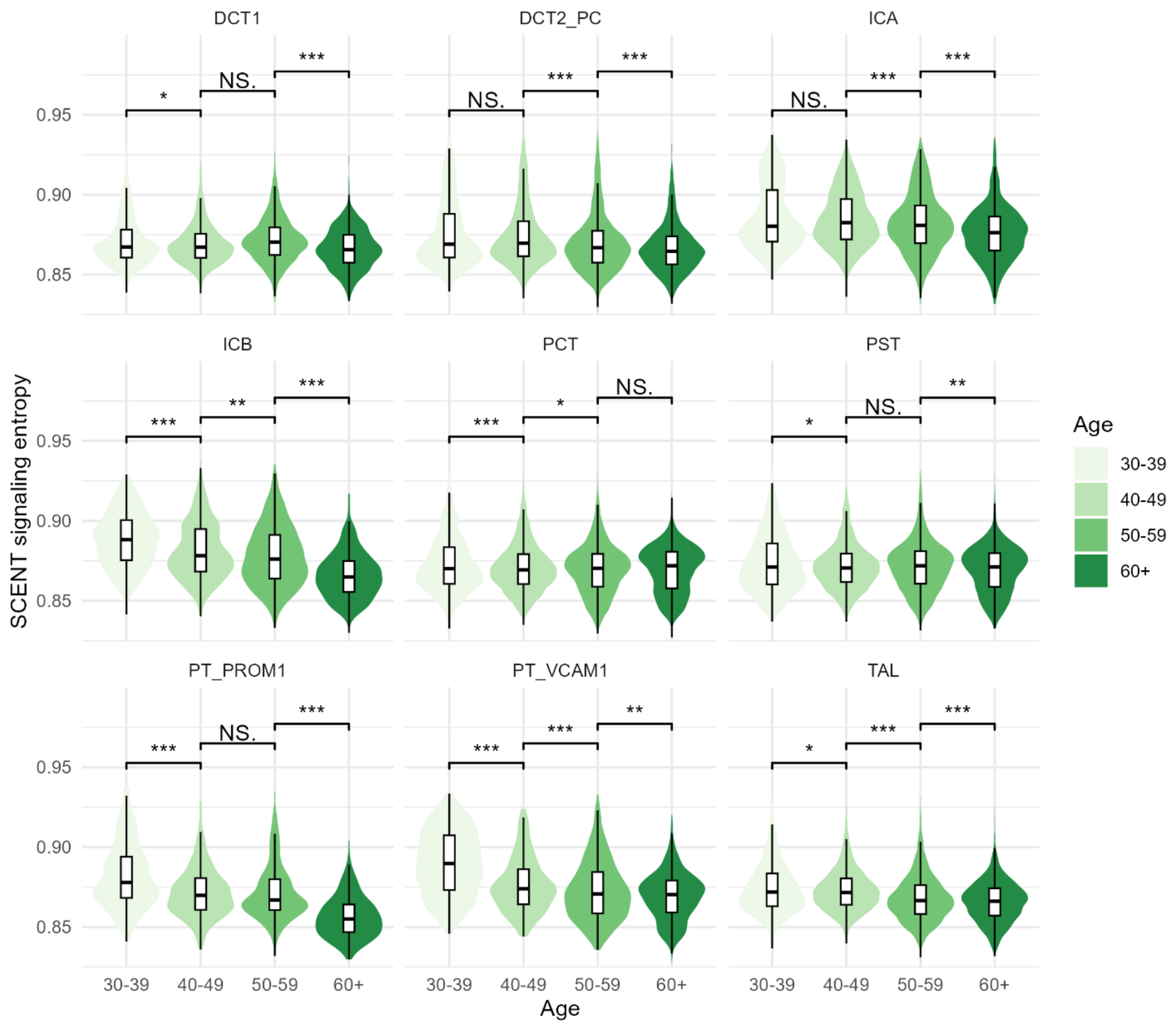


**Supplementary Figure 105 Distribution of SCENT signaling entropy in human kidney epithelial cells.**

Violin and box plots show the distribution of SCENT signaling entropy across human kidney epithelial cell types and age groups. SCENT computes it by integrating each cell's gene expression with a PPI network and calculating the entropy rate of a random walk over the network. Since this metric reflects differentiation potency, it is inversely related to our transcriptional dyscoordination measure. Pairwise comparisons between age groups were tested using one-sided Wilcoxon rank-sum test (alternative hypothesis: older age group has lower signaling entropy). Significance levels are annotated above each comparison (NS = not significant, * $p < 0.05$, ** $p < 0.01$, *** $p < 0.001$).

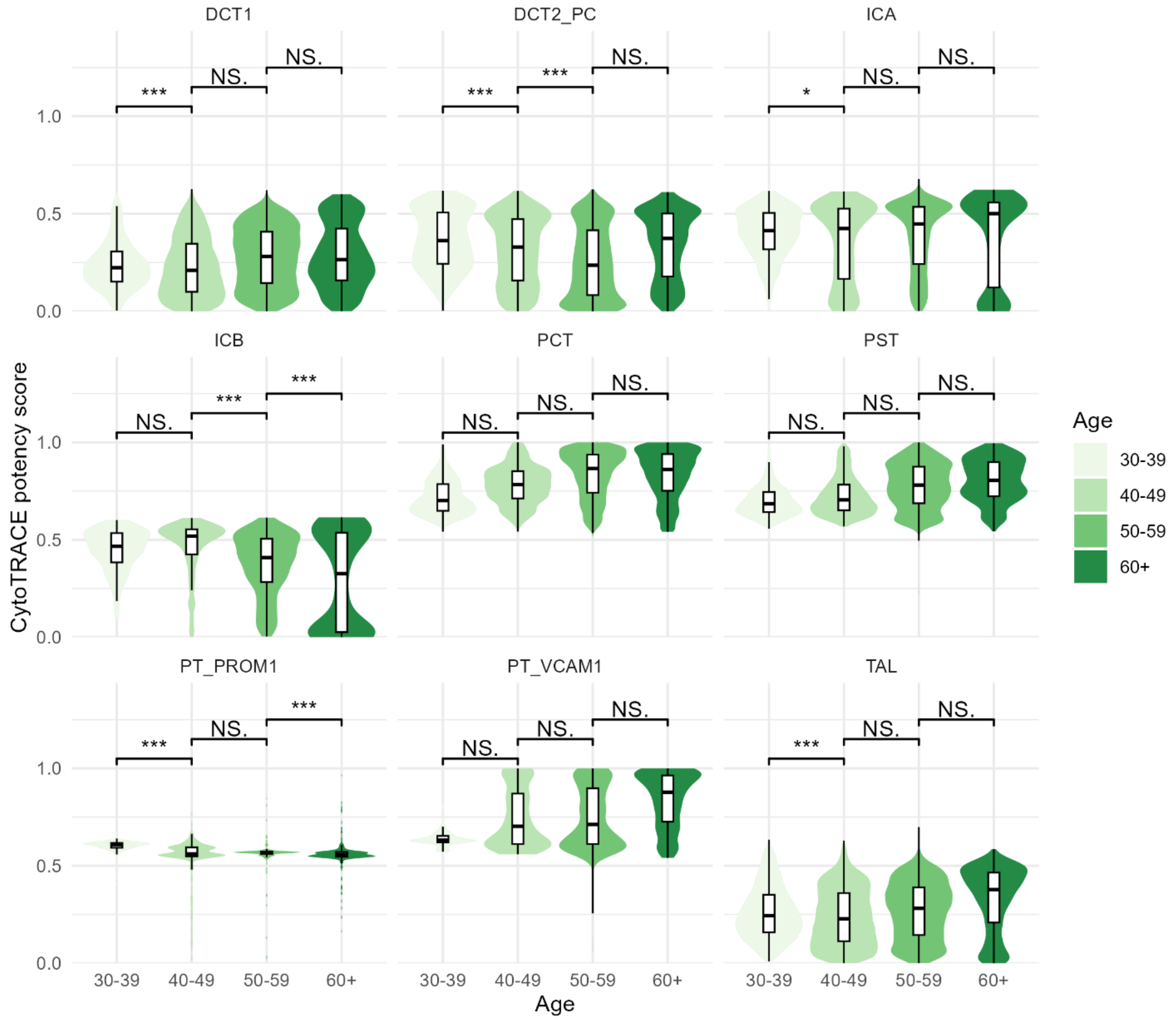


**Supplementary Figure 106 Distribution of CytoTRACE potency score in human kidney epithelial cells.**

Violin and box plots show the distribution of CytoTRACE potency score across human kidney epithelial cell types and age groups. CytoTRACE computes it from the number of distinctly expressed genes per cell, with more diverse transcriptome indicating higher potency. Since this metric reflects differentiation potency, it is inversely related to our transcriptional dyscoordination measure. Pairwise comparisons between age groups were tested using one-sided Wilcoxon rank-sum test (alternative hypothesis: older age group has lower potency score). Significance levels are annotated above each comparison (NS = not significant, * $p < 0.05$, ** $p < 0.01$, *** $p < 0.001$).

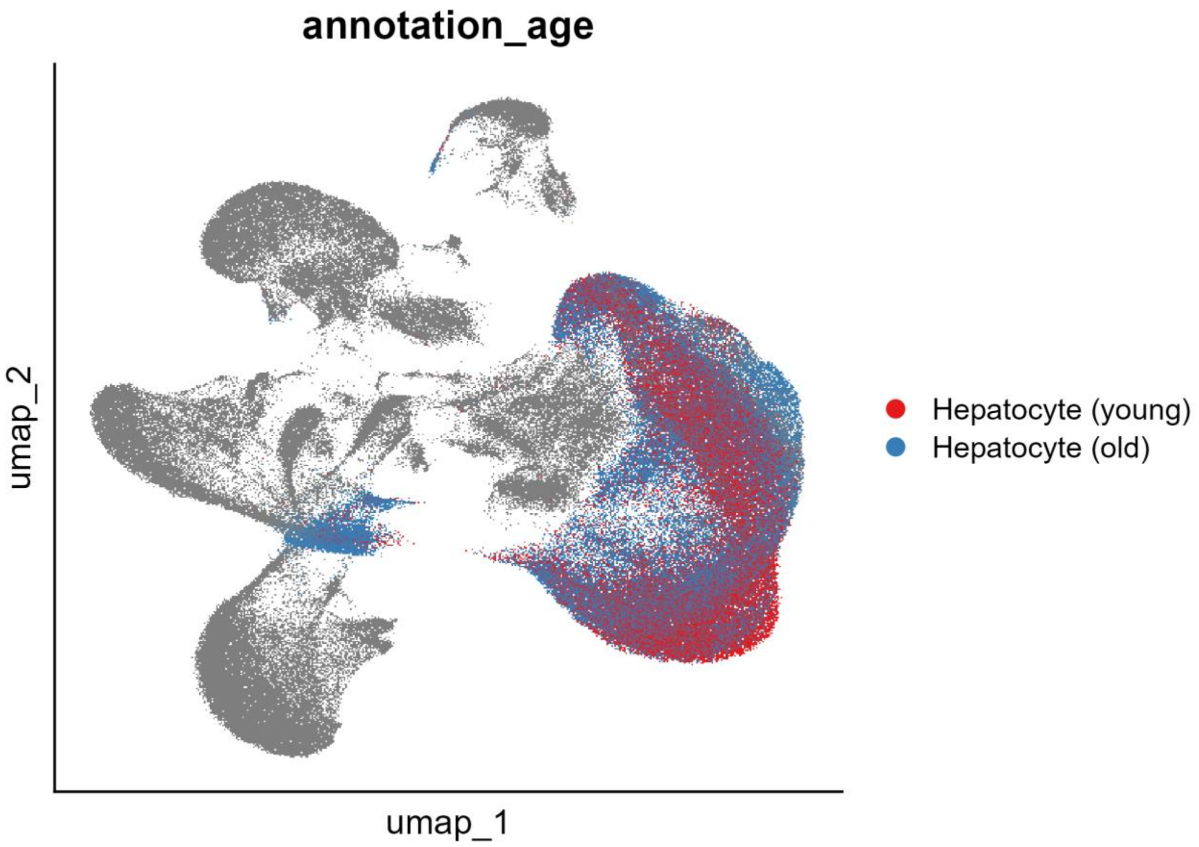


**Supplementary Figure 107 UMAP visualization of hepatocytes from aging mouse liver.**

Cells are embedded using the first two UMAP dimensions and colored by age group.

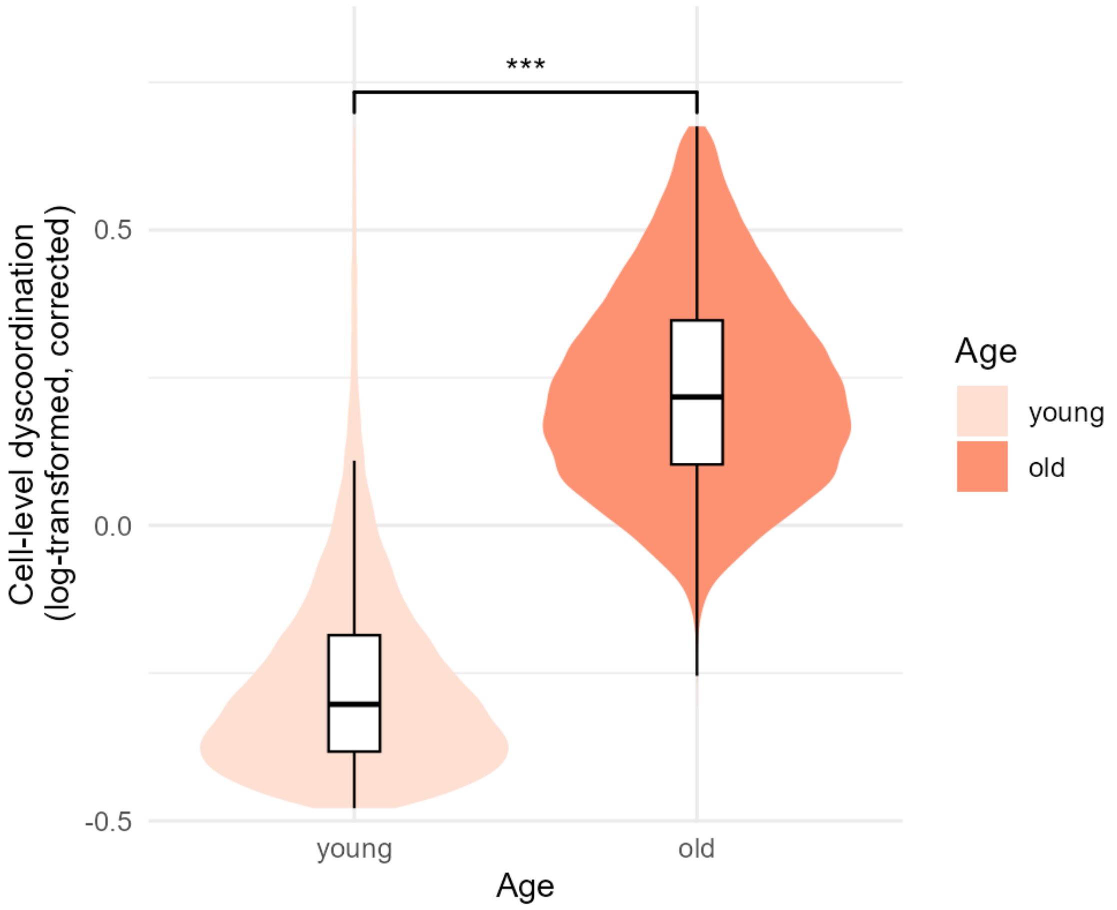


**Supplementary Figure 108 Distribution of corrected cell-level dyscoordination in mouse liver hepatocytes.**

Violin and box plots show the distribution of log-transformed, depth-corrected cell-level dyscoordination values separately for each cell type and age group. Values in the top and bottom 2.5% quantiles are considered outliers and are removed from the illustration. Pairwise comparisons between age groups were tested using one-sided Wilcoxon rank-sum test (alternative hypothesis: older age group has higher dyscoordination). Significance levels are annotated above each comparison (NS = not significant, * $p < 0.05$, ** $p < 0.01$, *** $p < 0.001$).

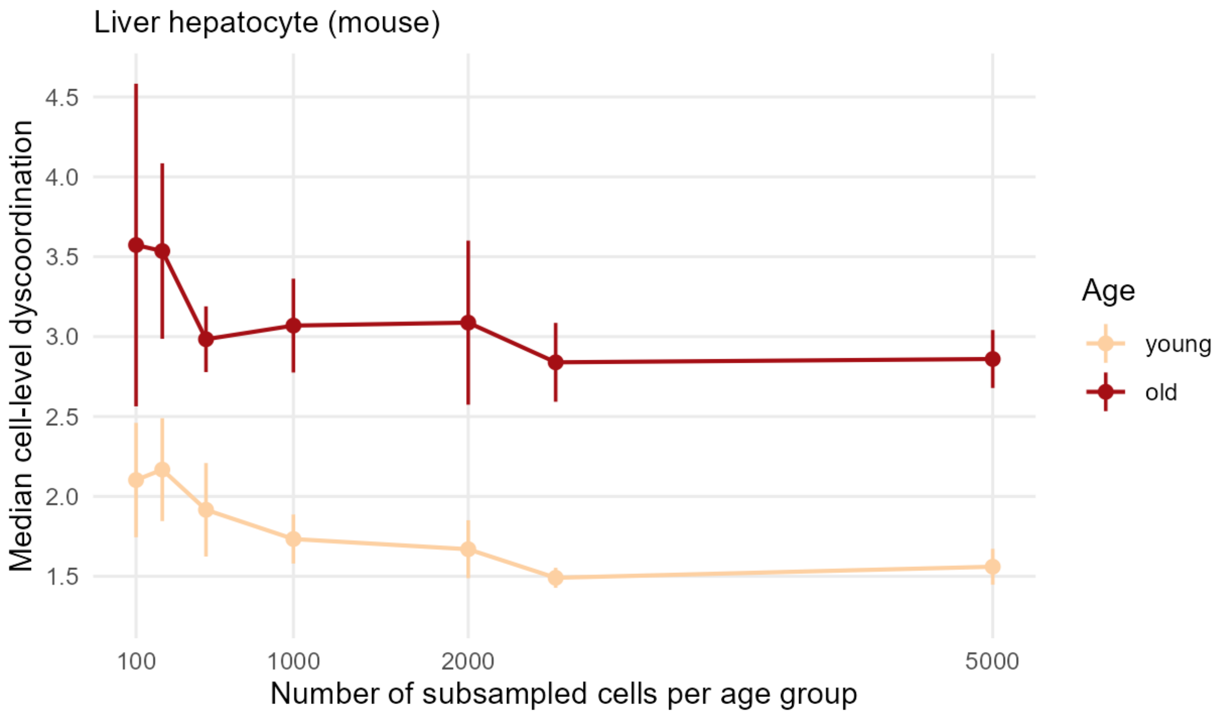


**Supplementary Figure 109 Median cell-level dyscoordination across subsampled age groups in aging mouse liver hepatocytes.**

Line plots show the median cell-level dyscoordination for each age group across random subsamples of increasing size (100 to 5,000 cells per age group). Each subsample is summarized by its median value, and each point shows the mean of these medians across 5 independent draws, with vertical lines indicating the 95% confidence interval across draws.

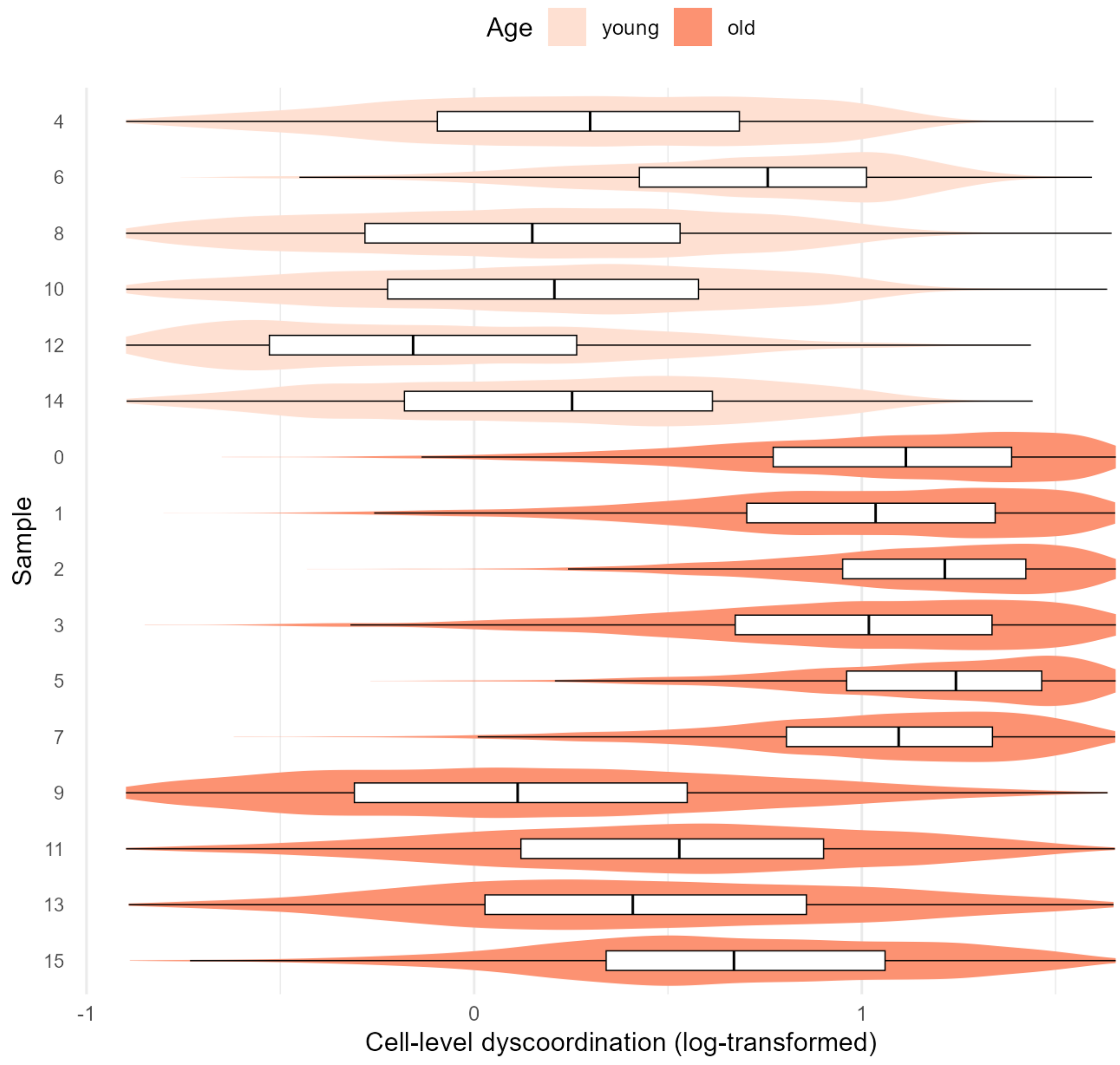


**Supplementary Figure 110 Distribution of cell-level dyscoordination across biological replicates in aging mouse liver hepatocytes.**

Violin and box plots show the distribution of log-transformed cell-level dyscoordination for each individual sample, ordered and colored by age group. Values in the top and bottom 2.5% quantiles are considered outliers and are removed from the illustration.

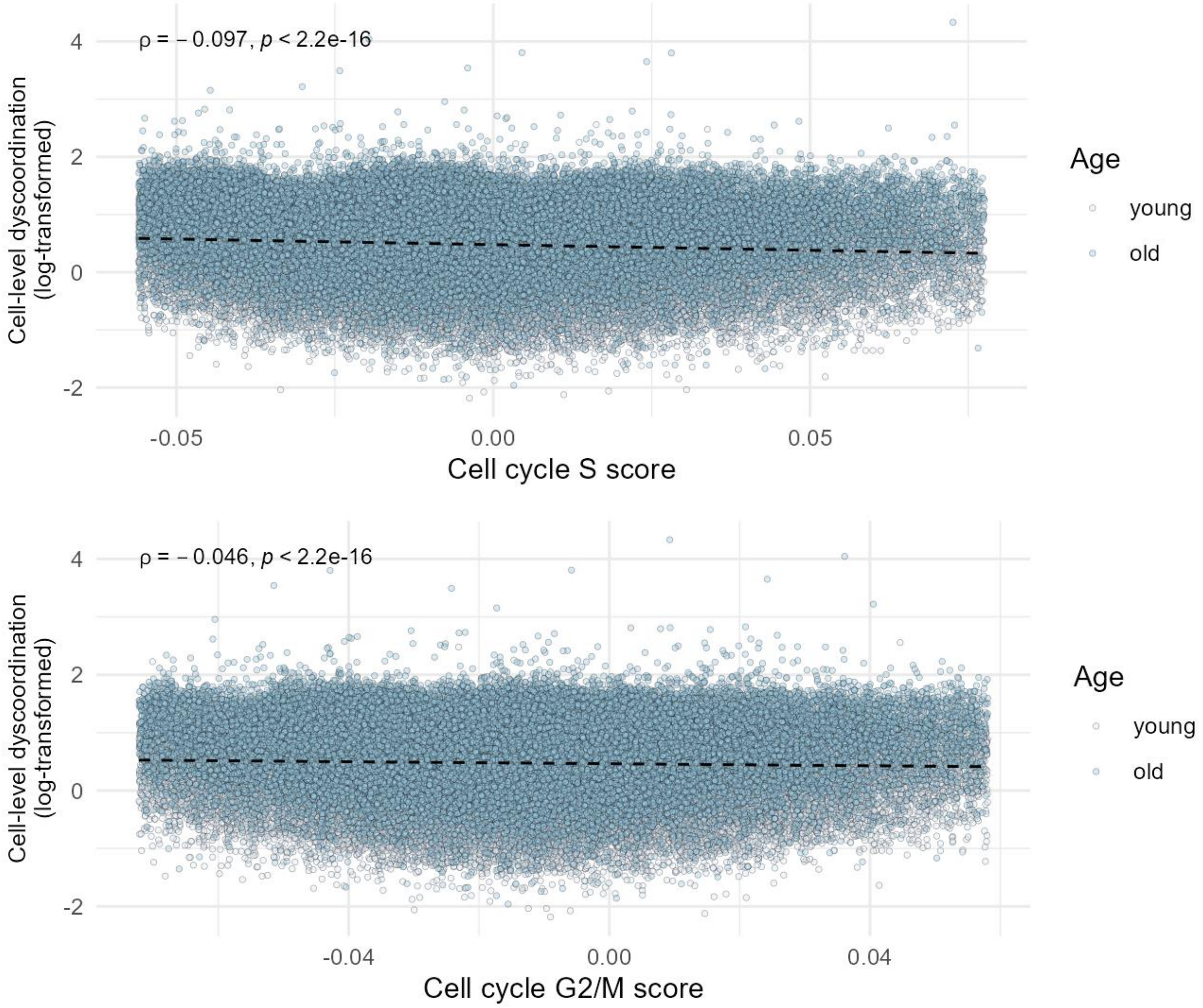


**Supplementary Figure 111 Correlation between cell cycle S phase and G2/M score and cell-level dyscoordination in aging mouse liver hepatocytes.**

Scatter plots show the relationship between the cell cycle S phase (top) and G2/M (bottom) score and log-transformed cell-level dyscoordination for each cell type. Black dashed lines indicate linear regression fits. Spearman's correlation coefficients and p-values are shown on the top left corner of each panel.

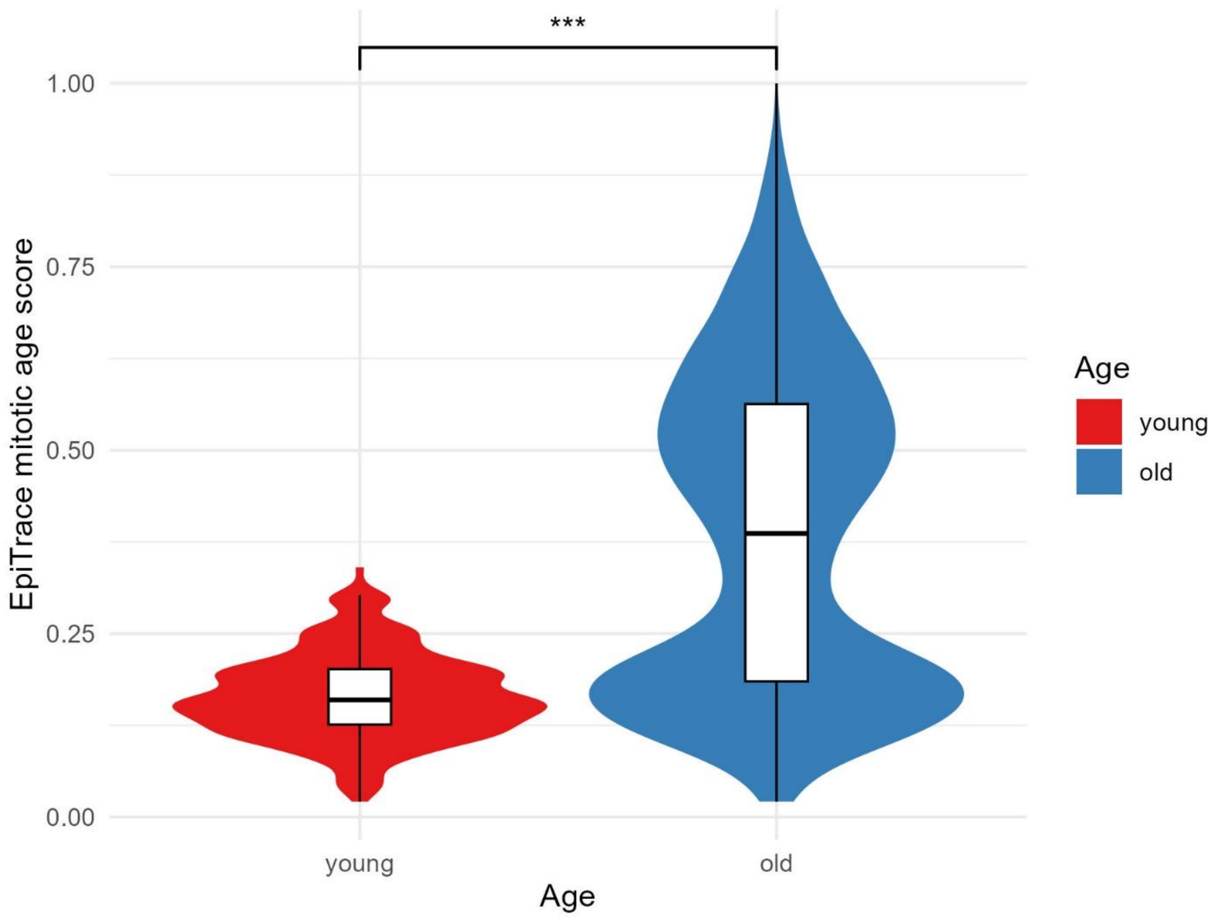


**Supplementary Figure 112 Distribution of EpiTrace mitotic age score in hepatocytes.**

Violin and box plots show the distribution of EpiTrace mitotic age score across age groups. Pairwise comparisons between age groups were tested using one-sided Wilcoxon rank-sum test (alternative hypothesis: older age group has higher EpiTrace score). Significance levels are annotated above each comparison (NS = not significant, * $p < 0.05$, ** $p < 0.01$, *** $p < 0.001$).

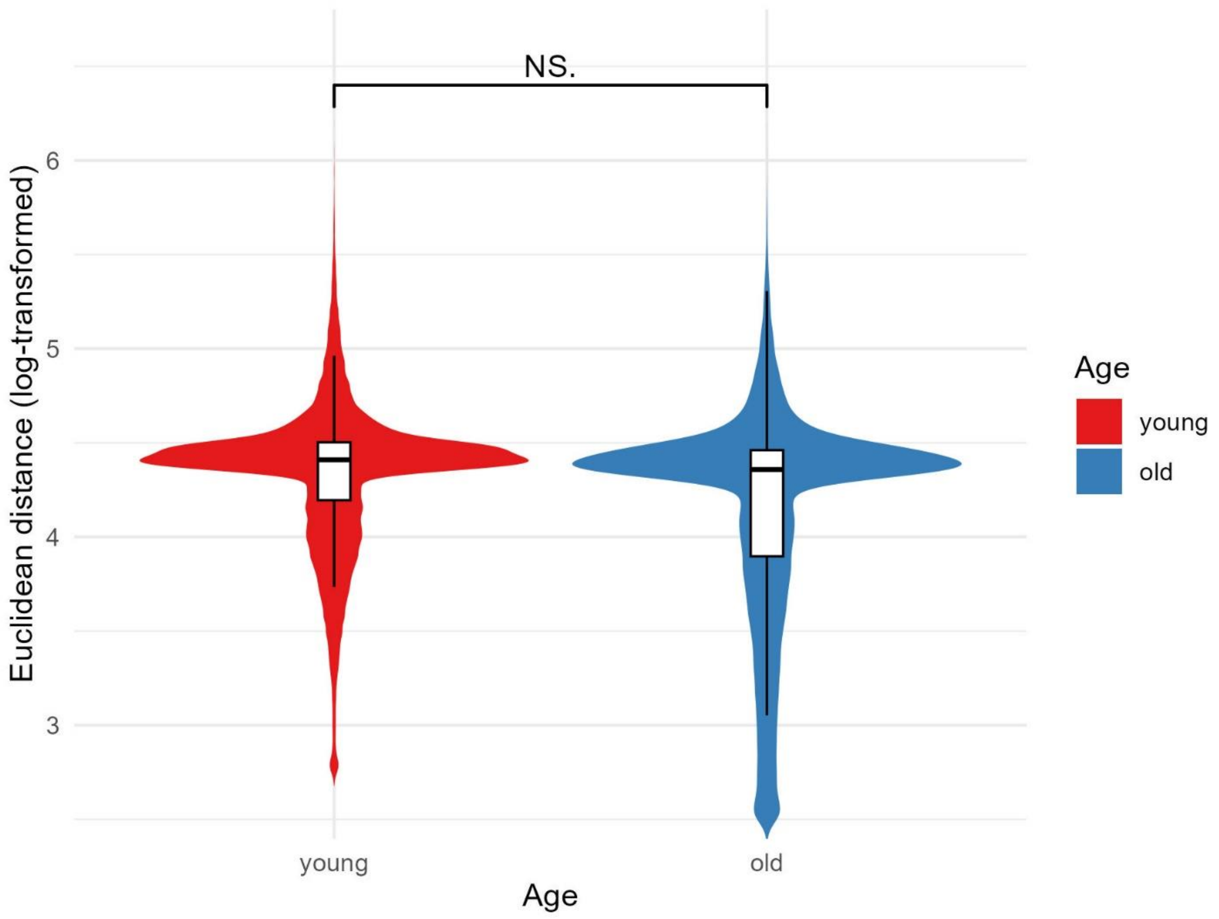


**Supplementary Figure 113 Distribution of Euclidean distance to cell type average from published methods in hepatocytes**.

Violin and box plots show the distribution of log-transformed Euclidean distance to cell type average across age groups. Pairwise comparisons between age groups were tested using one-sided Wilcoxon rank-sum test (alternative hypothesis: older age group has higher distance). Significance levels are annotated above each comparison (NS = not significant, * $p < 0.05$, ** $p < 0.01$, *** $p < 0.001$).

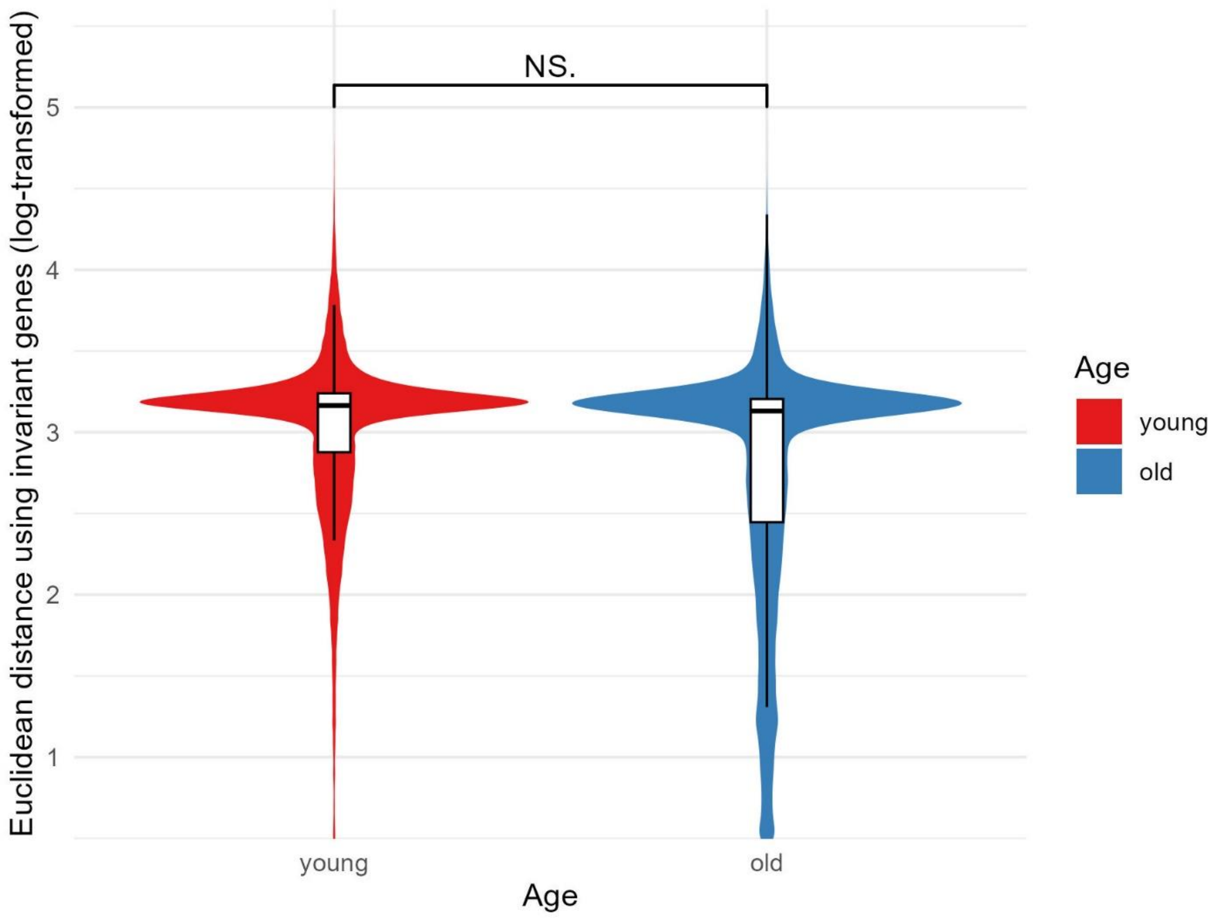


**Supplementary Figure 114 Distribution of Euclidean distance to tissue average using invariant genes in hepatocytes**.

Violin and box plots show the distribution of log-transformed Euclidean distance to tissue average using invariant genes across age groups. Pairwise comparisons between age groups were tested using one-sided Wilcoxon rank-sum test (alternative hypothesis: older age group has higher distance). Significance levels are annotated above each comparison (NS = not significant, * $p < 0.05$, ** $p < 0.01$, *** $p < 0.001$).

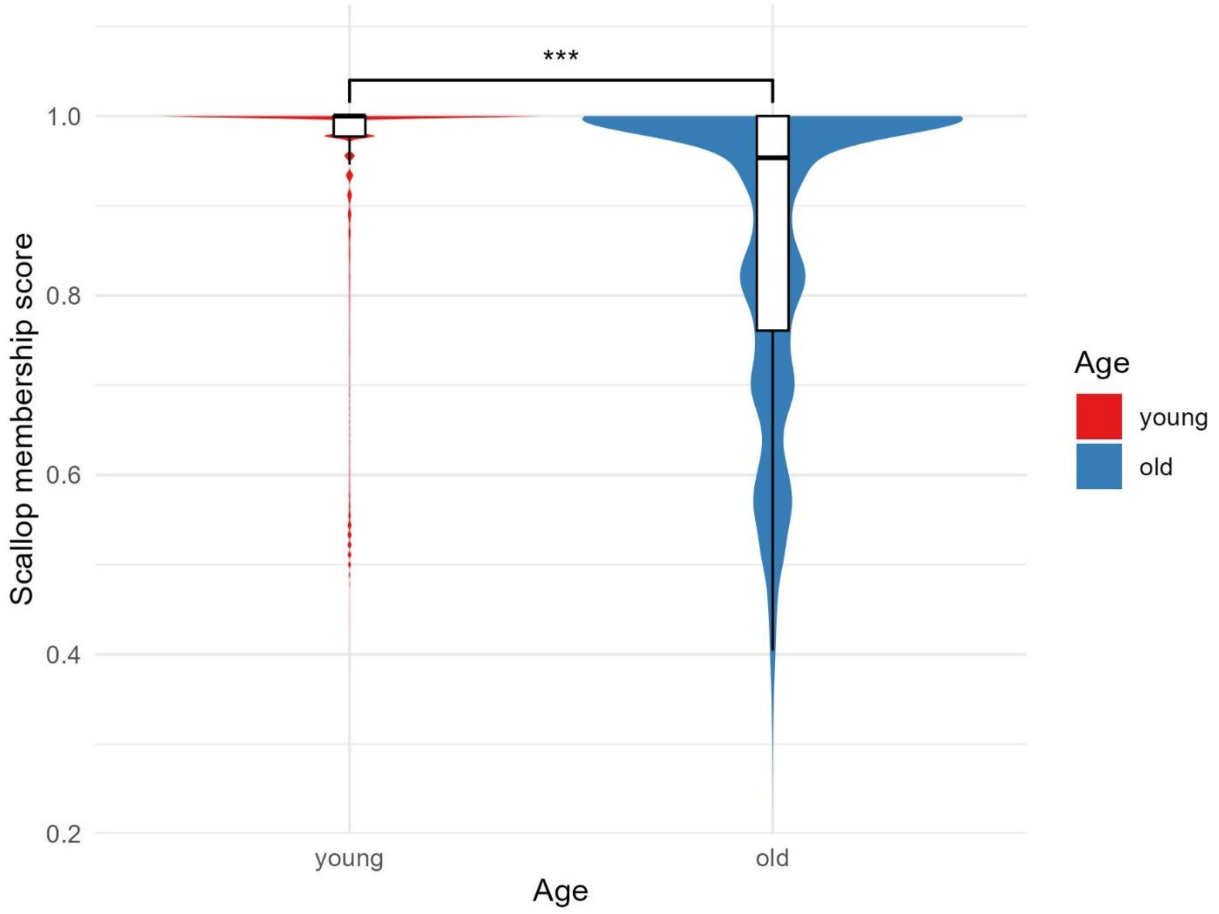


**Supplementary Figure 115 Distribution of Scallop membership score in hepatocytes**.

Violin and box plots show the distribution of Scallop membership score across age groups. Pairwise comparisons between age groups were tested using one-sided Wilcoxon rank-sum test (alternative hypothesis: older age group has lower membership score). Significance levels are annotated above each comparison (NS = not significant, * $p < 0.05$, ** $p < 0.01$, *** $p < 0.001$).

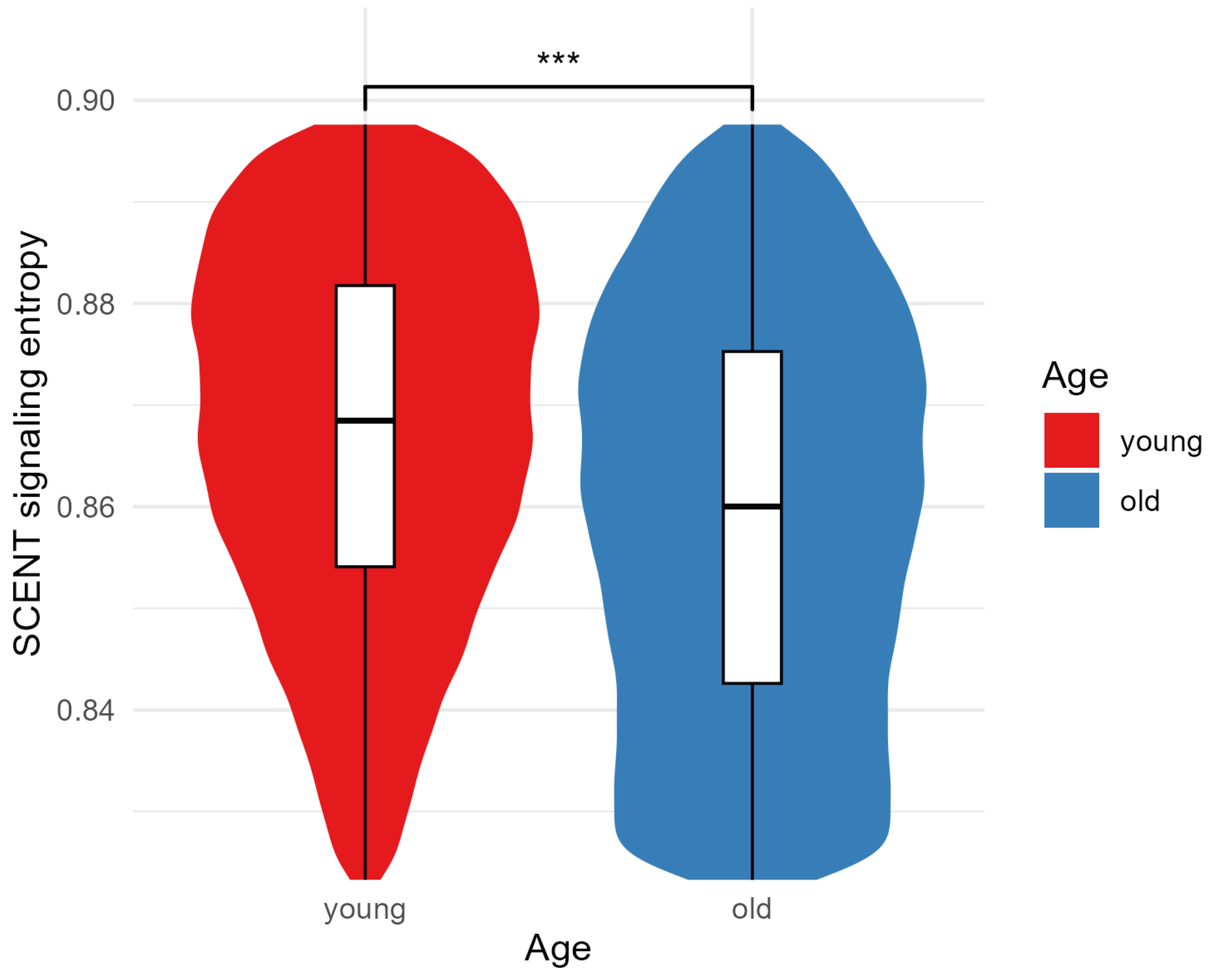


**Supplementary Figure 116 Distribution of SCENT signaling entropy in hepatocytes**.

Violin and box plots show the distribution of SCENT signaling entropy across age groups. Pairwise comparisons between age groups were tested using one-sided Wilcoxon rank-sum test (alternative hypothesis: older age group has lower signaling entropy). Significance levels are annotated above each comparison (NS = not significant, * $p < 0.05$, ** $p < 0.01$, *** $p < 0.001$).

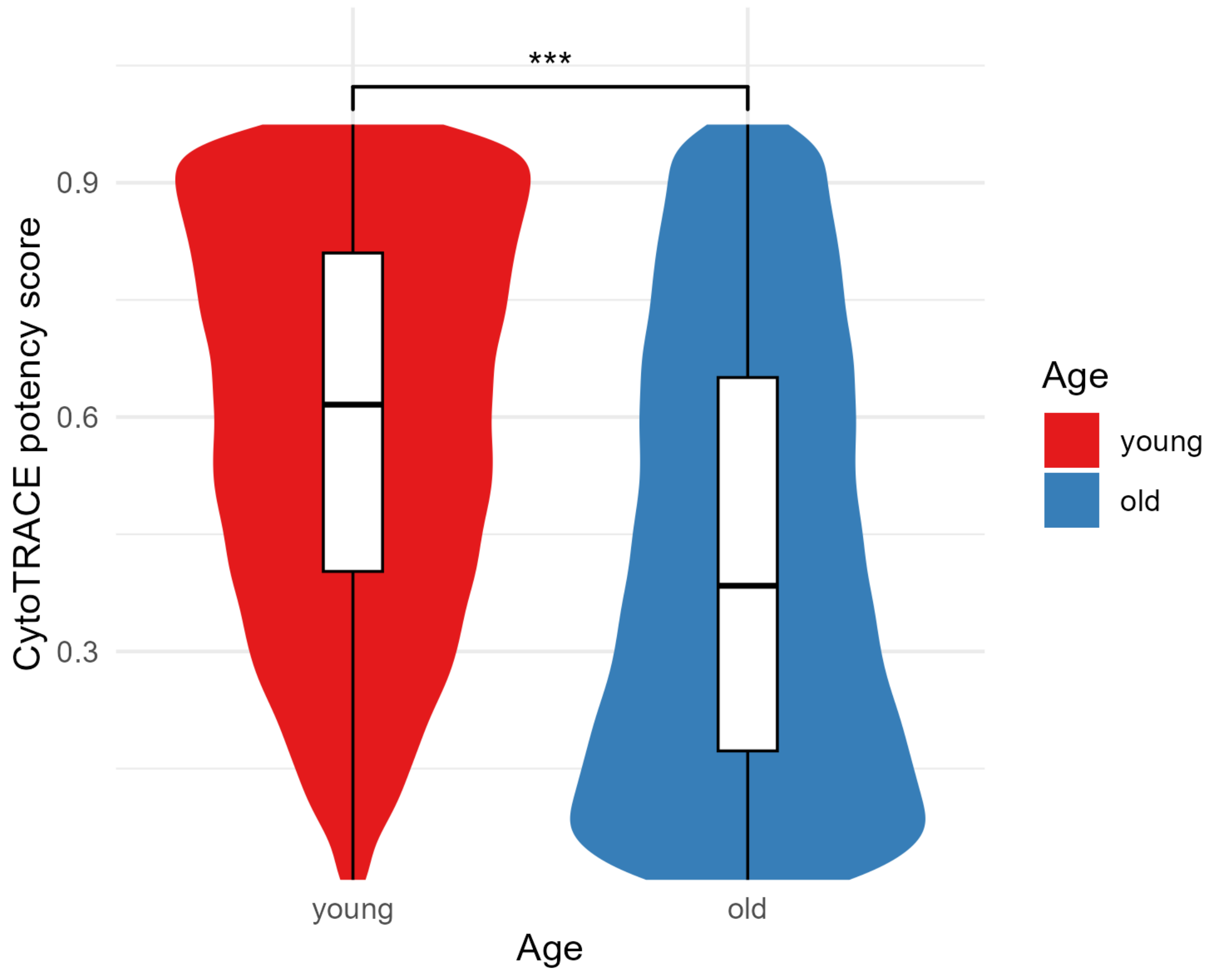


**Supplementary Figure 117 Distribution of CytoTRACE potency score in hepatocytes**.

Violin and box plots show the distribution of CytoTRACE potency score across age groups. Pairwise comparisons between age groups were tested using one-sided Wilcoxon rank-sum test (alternative hypothesis: older age group has lower potency score). Significance levels are annotated above each comparison (NS = not significant, * $p < 0.05$, ** $p < 0.01$, *** $p < 0.001$).

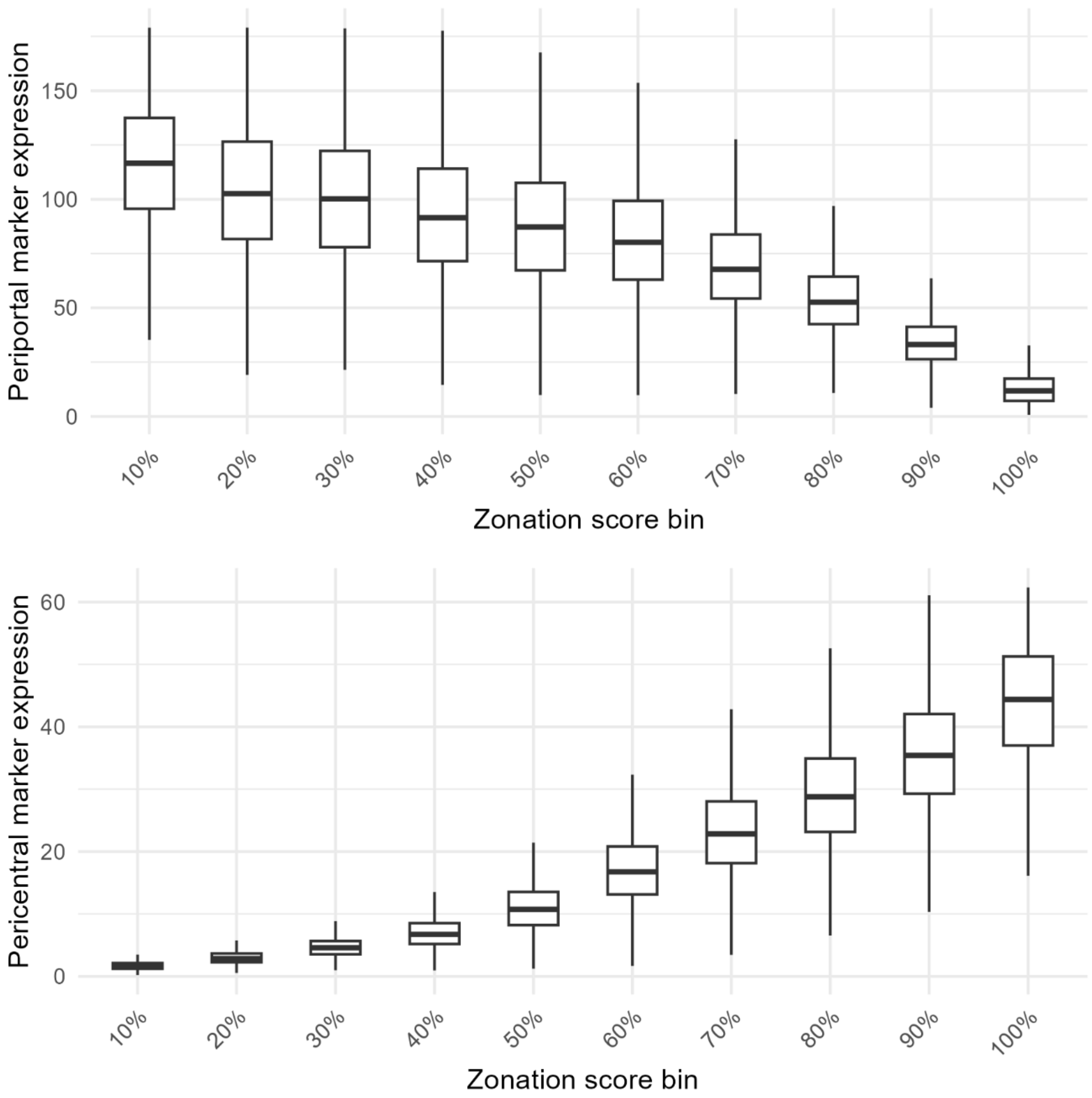


**Supplementary Figure 118 Distribution of periportal and pericentral marker expression by zonation score.**

Box plots show distribution of periportal (top) and pericentral (bottom) marker expression within each zonation score bin (x-axis indicating the upper thresholds). Values in the top 1% quantile are removed from the illustration.

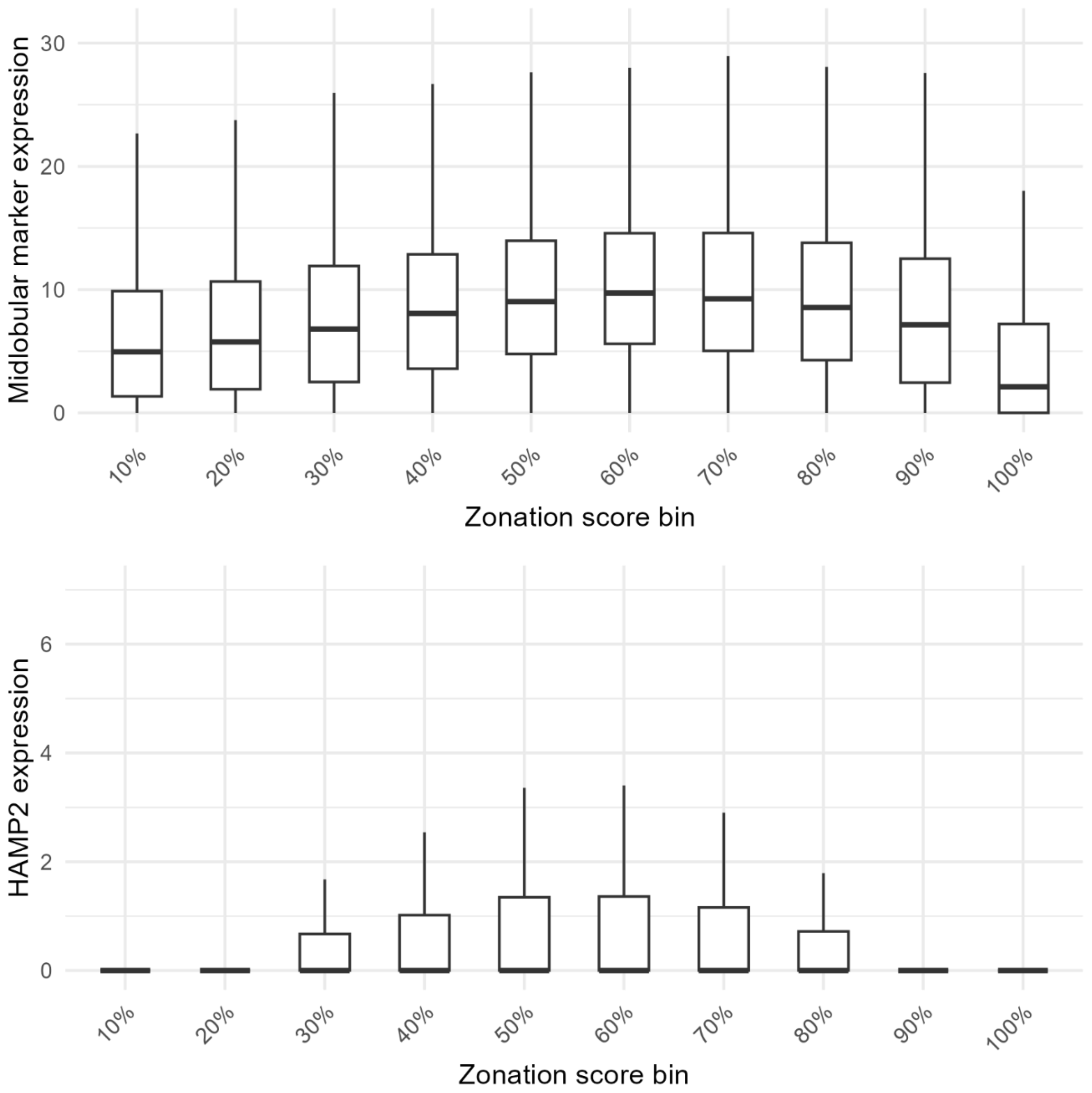


**Supplementary Figure 119 Distribution of midlobular marker and HAMP2 expression by zonation score.**

Box plots in the top panel show distribution of total expressions of 4 midlobular markers (HAMP, HAMP2, CCND1, CYP8B1) within each zonation score bin (x-axis indicating the upper thresholds). Box plots in the bottom panel show distribution of HAMP2 expression only within each zonation score bin. Values in the top 1% quantile are removed from the illustration.

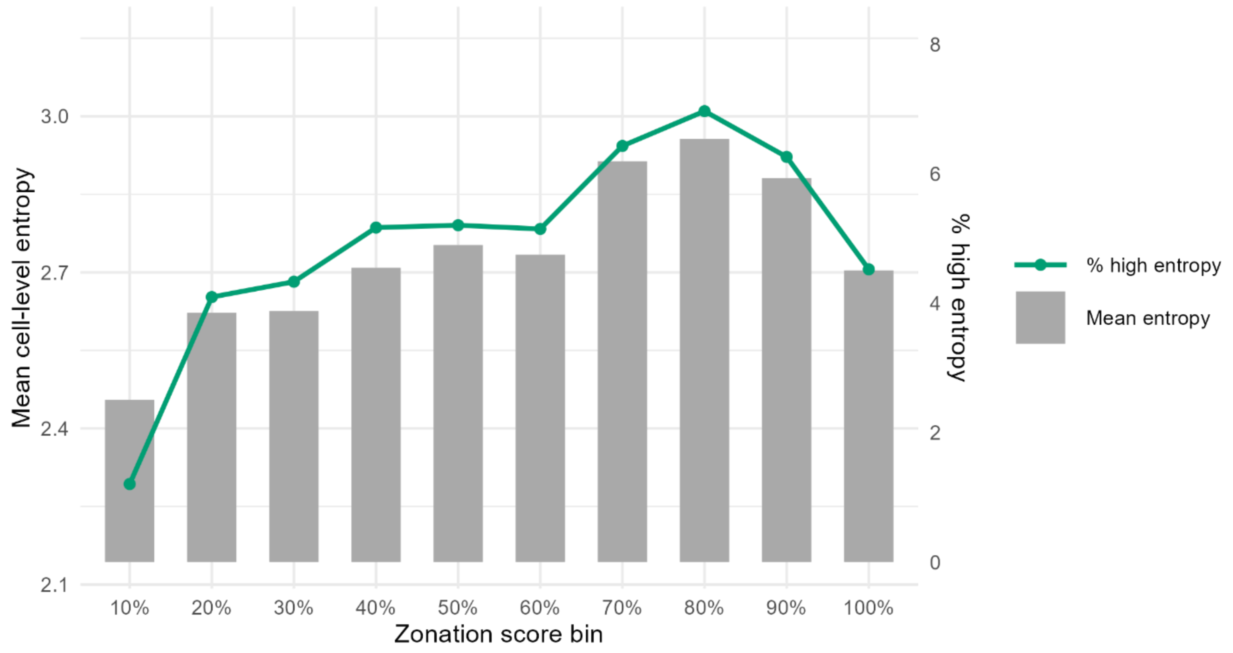


**Supplementary Figure 120 Distribution of mean cell-level dyscoordination and percentage of high-dyscoordination hepatocytes by zonation score**.

Grey bar plots represent the mean cell-level dyscoordination (left axis) within each zonation score bin (x-axis indicating the upper thresholds), while green line plots represent the percentage of high-dyscoordination cells (right axis), defined as those exceeding the 95$^{th}$ quantile of dyscoordination across all cells. These two measures follow a broadly similar trend along the zonation axis.

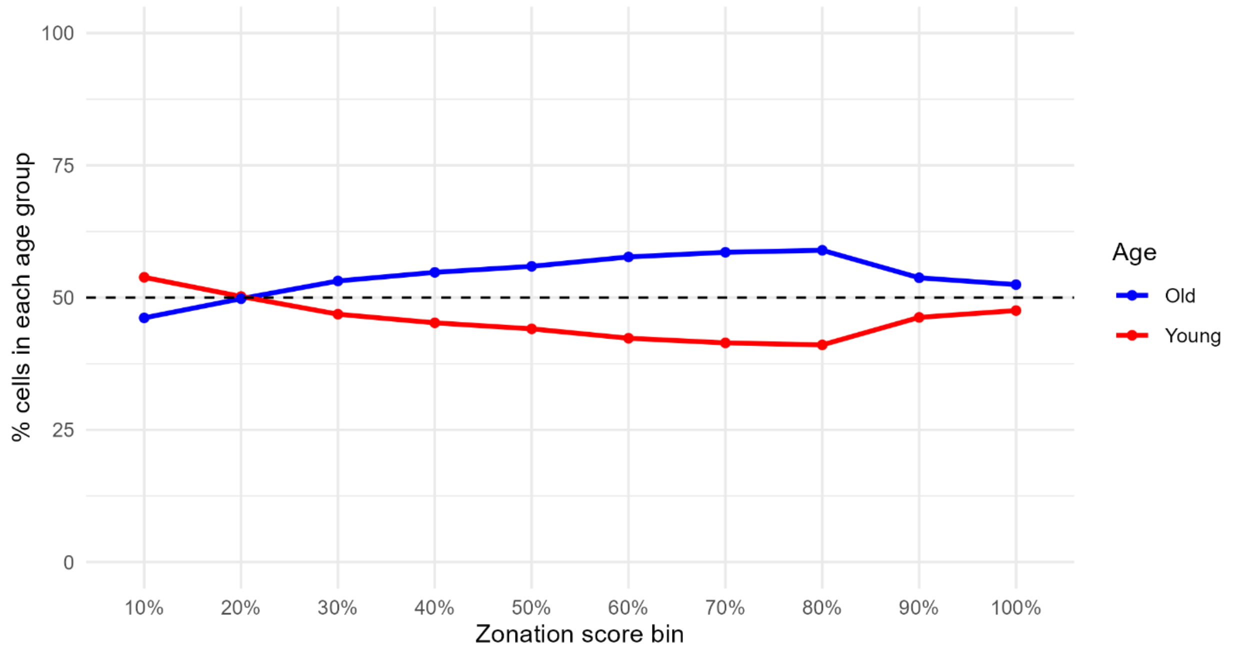


**Supplementary Figure 121 Distribution of hepatocytes in different age groups by zonation score.**

Solid lines show the percentage of cells in each age group within each zonation score bin (x-axis indicating the upper thresholds), while black dashed line represents 50%. Across all zonation score bins, these percentages remain mostly around 50:50, with the most unbalanced one being 60:40.

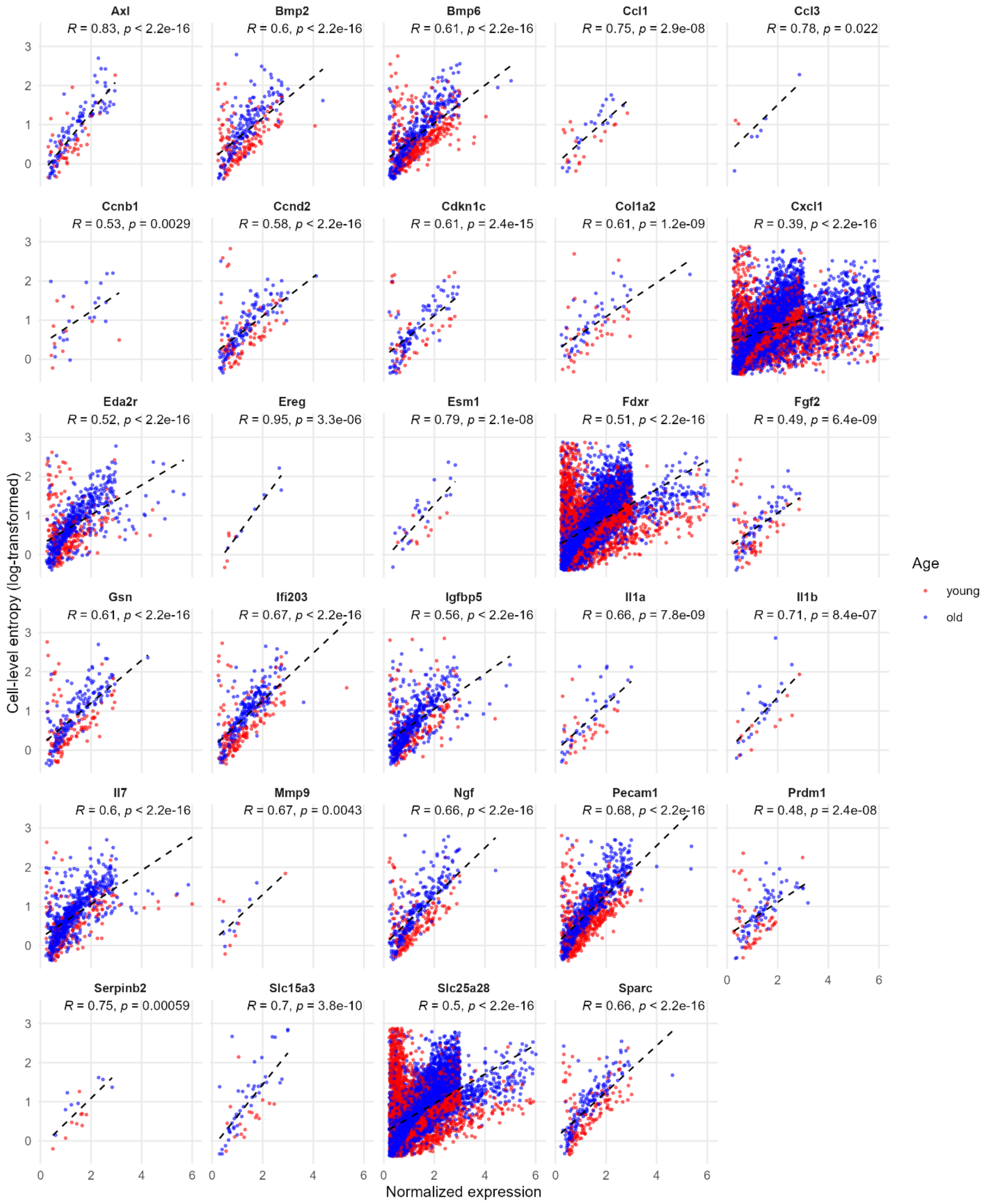
Axl
R = 0.83, p < 2.2e-16
Bmp2
R = 0.6, p < 2.2e-16
Bmp6
R = 0.61, p < 2.2e-16
Ccl1
R = 0.75, p = 2.9e-08
Ccl3
R = 0.78, p = 0.022
Ccnb1
R = 0.53, p = 0.0029
Ccnd2
R = 0.58, p < 2.2e-16
Cdkn1c
R = 0.61, p = 2.4e-15
Col1a2
R = 0.61, p = 1.2e-09
Cxcl1
R = 0.39, p < 2.2e-16
Eda2r
R = 0.52, p < 2.2e-16
Ereg
R = 0.95, p = 3.3e-06
Esm1
R = 0.79, p = 2.1e-08
Fdxr
R = 0.51, p < 2.2e-16
Fgf2
R = 0.49, p = 6.4e-09
Gsn
R = 0.61, p < 2.2e-16
Ifi203
R = 0.67, p < 2.2e-16
Igfbp5
R = 0.56, p < 2.2e-16
Il1a
R = 0.66, p = 7.8e-09
Il1b
R = 0.71, p = 8.4e-07
Il7
R = 0.6, p < 2.2e-16
Mmp9
R = 0.67, p = 0.0043
Ngf
R = 0.66, p < 2.2e-16
Pecam1
R = 0.68, p < 2.2e-16
Prdm1
R = 0.48, p = 2.4e-08
Serpinb2
R = 0.75, p = 0.00059
Slc15a3
R = 0.7, p = 3.8e-10
Slc25a28
R = 0.5, p < 2.2e-16
Sparc
R = 0.66, p < 2.2e-16
Cell-level entropy (log-transformed)
Normalized expression
Age
young
old

**Supplementary Figure 122 Scatter plots of cell-level dyscoordination versus expression of selected senescent genes in hepatocytes.**

29 senescent genes with positive correlations between their expressions and cell-level dyscoordination were selected for plotting. Cell-level dyscoordination (y-axis, log-transformed) and expression values (x-axis) were both filtered to remove zero values as well as the top and bottom 1% of nonzero values. Each point represents an individual cell colored its age group. A dashed black line shows the linear regression fit within each facet. Pearson correlation coefficients (R) and their respective p-values are shown in the top-right corners.

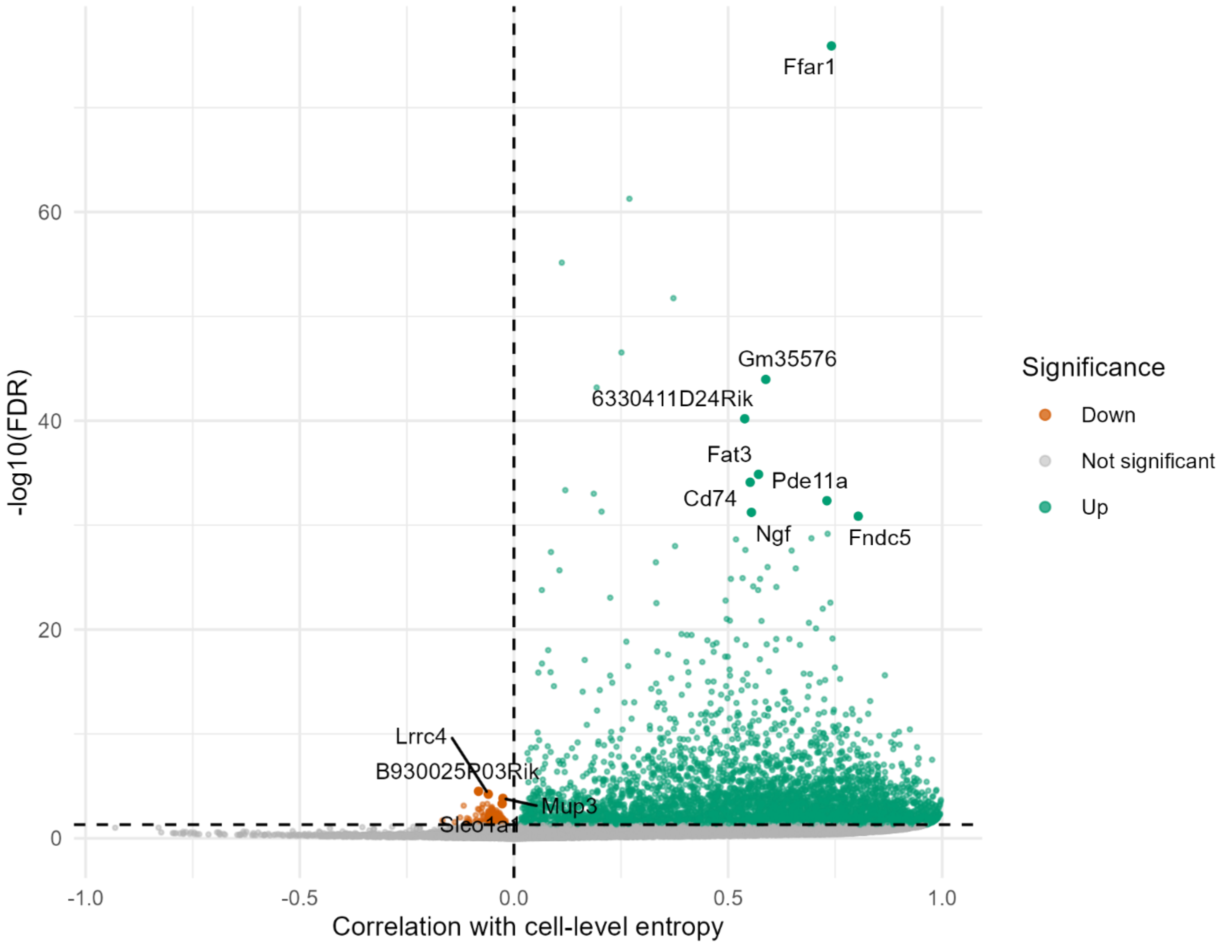


**Supplementary Figure 123 Correlation of gene expression with cell-level dyscoordination in hepatocytes.**

Volcano plots show the correlation of expression levels with cell-level dyscoordination for each individual gene. Positively and negatively correlated genes are separately colored. Top genes by their p-values with significantly positive correlations (> 0.5) or negative correlations (< 0) are highlighted.

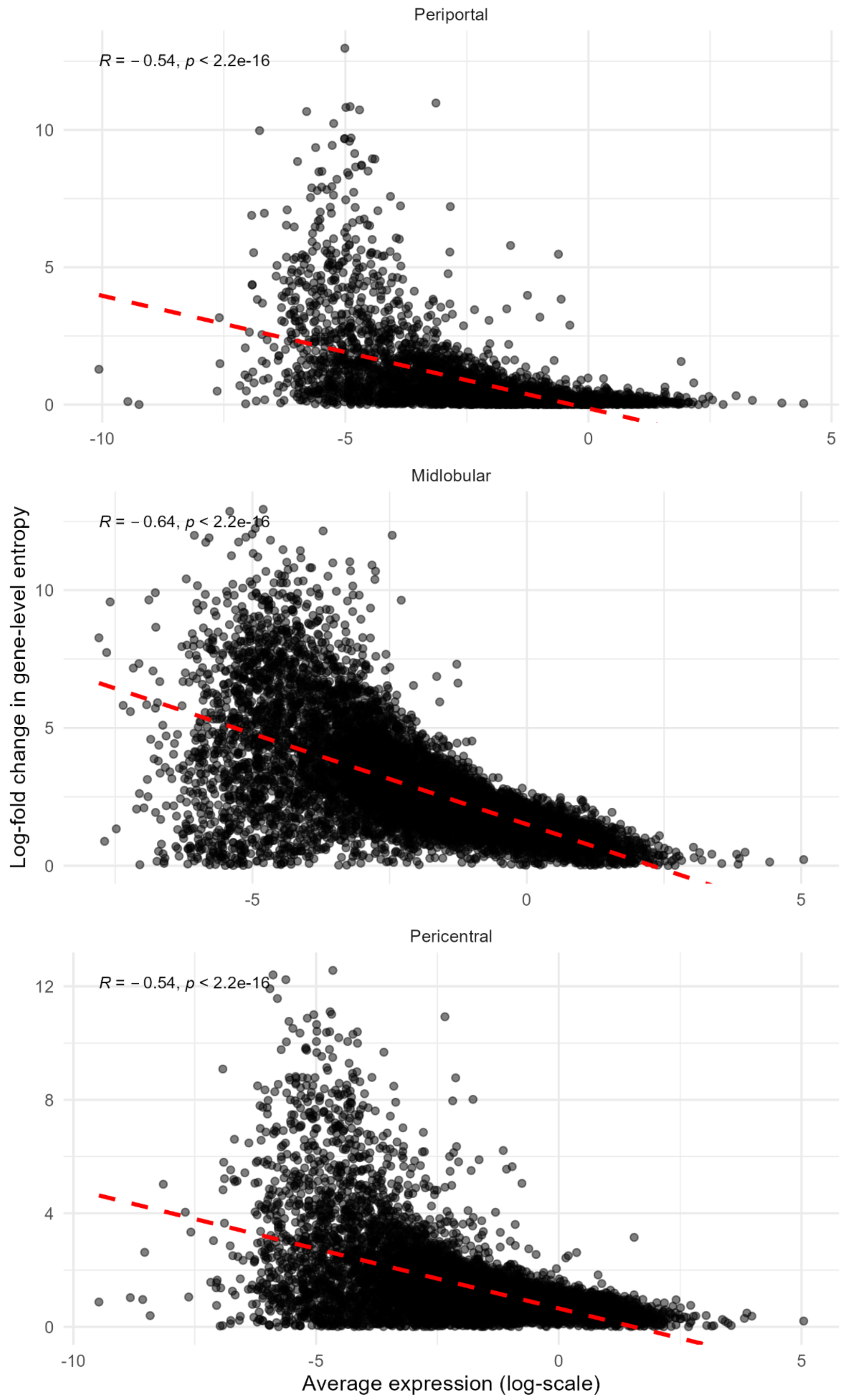

Periportal
R = − 0.54, p < 2.2e-16
Midlobular
R = − 0.64, p < 2.2e-16
Pericentral
R = − 0.54, p < 2.2e-16
Log-fold change in gene-level entropy
Average expression (log-scale)

**Supplementary Figure 124 Correlation between average expression and change in gene-level dyscoordination in hepatocytes.**

Scatter plots show the relationship between log-transformed average expression and log-fold change in gene-level dyscoordination (old vs. young) for each gene expressed in cells from each lobule. Red dashed lines indicate linear regression fits. Pearson correlation coefficients and corresponding p-values are shown in the top left corner of each panel.

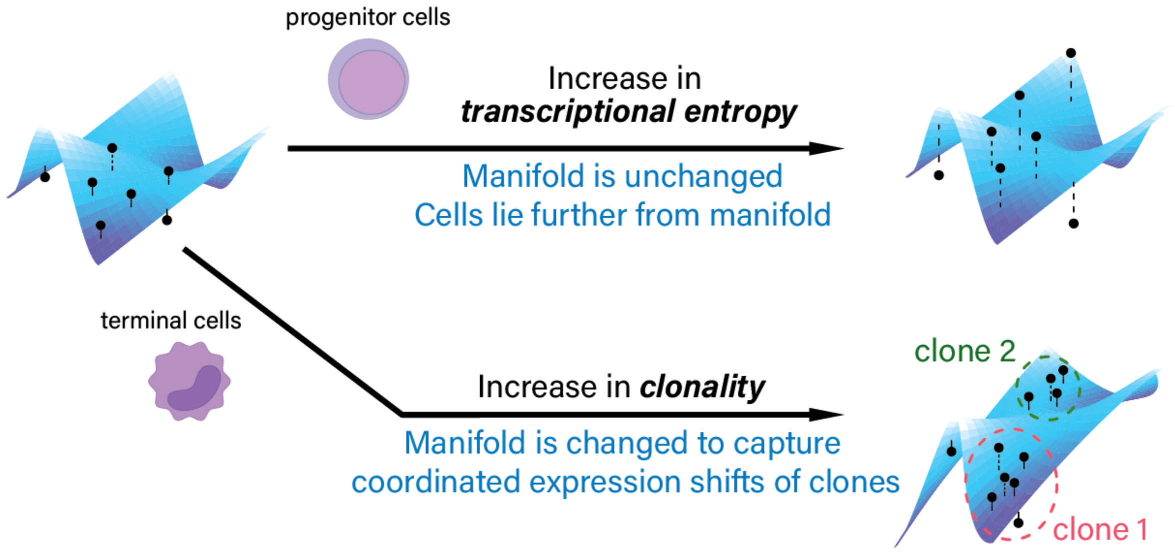


**Supplementary Figure 125 Transcriptional dysregulation vs. clonal evolution**

The diagram displays two distinct ways by which cells accumulate molecular damage over time. **Top**: As progenitor cells experience transcriptional dysregulation, the underlying manifold remains stable, but individual cells deviate further from it due to increased transcriptional dyscoordination. **Bottom**: As terminal cells undergo clonal evolution, the manifold shifts to accommodate shared expression changes of clones, reflecting increased clonality. This illustrates that transcriptional dyscoordination captures intrinsic molecular disorder rather than structured heterogeneity driven by clonal expansion.
